# Orthogonal IMiD-Degron Pairs Induce Selective Protein Degradation in Cells

**DOI:** 10.1101/2024.03.15.585309

**Authors:** Patrick J. Brennan, Rebecca E. Saunders, Mary Spanou, Marta Serafini, Liang Sun, Guillaume P. Heger, Agnieszka Konopacka, Ryan D. Beveridge, Laurie Gordon, Shenaz B. Bunally, Aurore Saudemont, Andrew B. Benowitz, Carlos Martinez-Fleites, Markus A. Queisser, Heeseon An, Charlotte M. Deane, Michael M. Hann, Lewis L. Brayshaw, Stuart J. Conway

**Affiliations:** Department of Chemistry, Chemistry Research Laboratory, University of Oxford; Oxford, UK; Department of Chemistry & Biochemistry, University of California, Los Angeles; Los Angeles, USA; GSK, Medicines Research Centre; Stevenage, UK; PerkinElmer; Beaconsfield, UK; Chemical Biology Program, Memorial Sloan Kettering Cancer Center; New York, USA; Virus Screening Facility, Weatherall Institute of Molecular Medicine, University of Oxford; Oxford, UK; Department of Statistics, University of Oxford; Oxford, UK

## Abstract

Immunomodulatory imide drugs (IMiDs) including thalidomide, lenalidomide, and pomalidomide, can be used to induce degradation of a protein of interest that is fused to a short zinc finger (ZF) degron motif. These IMiDs, however, also induce degradation of endogenous neosubstrates, including IKZF1 and IKZF3. To improve degradation selectivity, we took a bump-and-hole approach to design and screen bumped IMiD analogs against 8380 ZF mutants. This yielded a bumped IMiD analog that induces efficient degradation of a mutant ZF degron, while not affecting other cellular proteins, including IKZF1 and IKZF3. In proof-of-concept studies, this system was applied to induce efficient degradation of TRIM28, a disease-relevant protein with no known small molecule binders. We anticipate that this system will make a valuable addition to the current arsenal of degron systems for use in target validation.

**One-Sentence Summary:** Engineered zinc-finger-based degrons enable targeted protein degradation induced by selective molecular glues.

## Introduction

Chemically induced degradation of proteins is a powerful complement to traditional, occupancy-based, small molecule modulation of a biological target. This strategy allows all functions of a protein to be neutralized, including scaffolding roles, and the action of undruggable domains that would otherwise be difficult to inhibit. Within this approach, two distinct small molecule-based strategies have been developed: proteolysis targeting chimeras (PROTACs) and molecular glues (*1*). PROTACs are bifunctional molecules that induce proximity between the target protein of interest (POI) and an E3 ligase to promote POI degradation (*1*, *2*). Alternatively, molecular glues are small molecules that bind at the interface of two proteins, enhancing or inducing their interaction (*3*, *4*). Molecules that function as molecular glues include the natural products auxin, cyclosporin, FK506, and rapamycin, and synthetic stabilizers of the hub protein 14-3-3 with its binding partners (*5*). Immunomodulatory imide drugs (IMiDs) are a well-studied family of molecular glues that includes thalidomide, lenalidomide, pomalidomide, and CC-220 (*6–8*). IMiDs possess a glutarimide ring that binds to a tri-tryptophan pocket in cereblon (CRBN), a substrate adaptor of the Cullin4 RING E3 ligase (CRL4^CRBN^) (*9*, *10*). Binding of the IMiD remodels the CRL4^CRBN^ protein surface, inducing affinity between the CRBN-IMiD binary complex and a series of β-hairpin-containing proteins, known as neosubstrates. Subsequent ubiquitination by the CRL4^CRBN^ machinery results in proteasomal degradation of the neosubstrate. Recent work by Heim *et al*. and Ichikawa *et al*. has shown that the true endogenous targets of CRBN are proteins that possess similar cyclic imides as a C-terminal post-translational modification (PTM), formed by intramolecular cyclization of glutamine or asparagine residues (*11*, *12*).

At least ten CRL4^CRBN^ neosubstrates are known, including Ikaros family zinc finger protein 1 (IKZF1; Ikaros) and Ikaros family zinc finger protein 3 (IKZF3; Aiolos), ZFP91 zinc finger protein (ZFP91), G1 to S phase transition 1 protein (GSPT1), casein kinase 1 alpha 1 (CK1α), and spalt-like transcription factor 4 (SALL4); the latter of which is thought to be the effector of the teratogenic properties of thalidomide (*13–23*). Although these neosubstrates do not possess a specific consensus sequence, they all share a β-hairpin motif with a conserved glycine residue which binds at the CRBN-IMiD interface (*6*, *24*), and is frequently part of a C2H2 zinc finger (ZF). This motif is an example of a degron, broadly defined as a targeting signal that confers metabolic instability on some, or all, of the peptide bonds in a protein (*25–27*). This definition encompasses inducible degrons, which require the presence of a small molecule to promote protein degradation. Genetic knock-in methods enable a degron to be used in a complementary manner to PROTACs, allowing induced degradation of POIs that lack a small molecule ligand (*25*, *28*). A number of these technologies are well established, including the auxin inducible degron system (AID), small molecule assisted shutoff (SMASh-tag), a destabilizing domain (DD) stabilized by small molecule Shld1, a degron based on methyl guanine methyltransferase (MGMT), and systems that utilize bifunctional small molecules, including dTAG (degron = 11.9 kDa), HaloPROTAC (degron = 33.6 kDa), Bromotag (degron = 14.9 kDa), and an approach based on NanoLuciferase (*28–37*). While these techniques have been elegantly applied in a numerous studies, a potential drawback is the use of large degron motifs, which can negatively affect feasibility of CRISPR knock-in and function of the POI (*25*, *34*). Recent work by Tsang *et al*. offers a potential solution to this using HiBiT-SpyTag and SpyCatcher to add a degron motif to a POI (*38*). However, whilst this approach does allow minimal alteration of the POI, it also requires further transfection stages (*39–41*).

A combination of IMiD small molecules and ZF-based degrons offer an attractive alternative as inducible degrons; the low molecular weight inducers employed can readily enter cells and do not display a hook effect. In addition, the ZF degron motif can be as small as 23 amino acid residues, but works optimally at 60 amino acids (*6*, *24*), minimally perturbing the POI (degron = 7.0 kDa). Existing implementations of such systems have used degron motifs derived from IKZF1/3, SALL4, and hybrid sequences comprising halves of two different neosubstrate ZF degron sequences, such as Superdegron and iTAG (*24*, *42–45*), These IMiD-degron systems have been used to degrade chimeric antigen receptors (CARs) in T-cells, and modulate CRISPR Cas9 genetic editing (*44–47*). While this elegant approach overcomes some limitations of other inducible degrons, the use of existing small molecule IMiDs presents important selectivity-related limitations. Sievers *et al*. identified 11 zinc finger-containing transcription factors that were degraded in the presence of thalidomide, lenalidomide, or pomalidomide (*6*). Computational analysis suggested that more than 150 zinc fingers can bind to the drug-CRBN complex. Therefore, the use of IMiDs that can bind to the wild-type (WT) ZF degrons in CAR T-cell therapy, or target validation, risks degradation of additional neosubstrates leading to unwanted effects.

To overcome the above limitations, we employed a bump-and-hole approach developed IMiD-degron pairs with orthogonality to the existing IMiD-WT ZF combinations (Figure 1). This involves making a mutant protein with a ‘hole’ that can accommodate a ‘bumped’ ligand that can bind to this mutant, but not the original wild type (WT) protein. This approach was pioneered by Schreiber using FKBP12 and cyclophilin, and Shokat who applied this approach to generating selective kinase-ligand pairs (*48–51*). More recently, Ciulli has published an elegant application of the ‘bump and hole’ approach to distinguish between the structurally similar first and second bromodomains of BRD4 (*28*, *52*).

**Figure 1.**
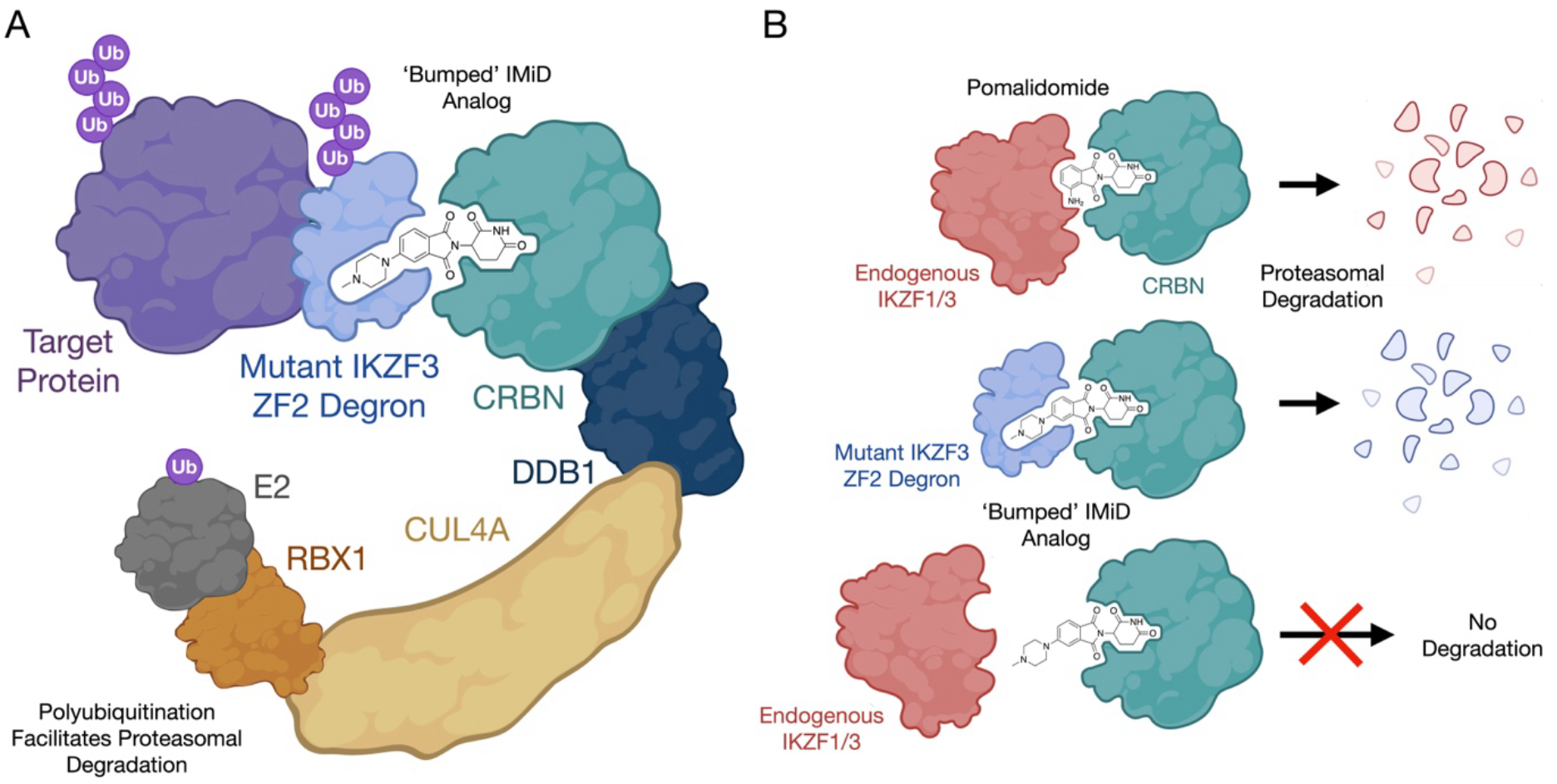
Schematic depicting the mechanism of action for the bumped IMiD degron system. **A.** The IMiD molecule binds to CRBN, and the resulting binary complex binds to the mutant IKZF3 ZF2-derived degron. This brings the degron-tagged target protein into proximity to the CRBN-recruited ubiquitination machinery, effecting proteasomal degradation. Ub, ubiquitin. **B.** The bump and hole strategy: traditional IMiDs such as pomalidomide induce degradation of neosubstrates such as IKZF1 and IKZF3; a bumped IMiD analog will recruit a degron with a suitable ‘hole’ mutation, but will ignore endogenous neosubstrates.

Here, we designed a series of bumped IMiD derivatives and selected compounds that showed only modest degradation of the WT IKZF3-derived degron attached to enhanced green fluorescent protein (EGFP). We then designed a library of 8380 ZF mutants, using a similar approach to Sievers *et al*. and Jan *et al*. (*6*, *44*), and screened this against the bumped IMiDs. This approach resulted in the discovery of an IMiD analog-mutant ZF degron pairing that efficiently degrades a tagged POI. Proteomics studies revealed this pairing has exquisite selectivity over other cellular proteins, including known CRL4^CRBN^ neosubstrates.

### Bumped IMiD analogs preferentially degrade the Q147A mutant of the IKZF3-EGFP fusion over the WT IKZF3-EGFP fusion

Initial work focused on the identification of appropriate ZF residues to mutate, based on the crystal structure of pomalidomide bound to CRBN and the IKZF1 ZF (PDB ID: 6H0F, Fig. 1C) (*6*). IKZF3 residues Q147, N149, Q150, C151, and G152 form the interface with pomalidomide-bound CRBN (Figure 2A; IKZF1 and IKZF3 possess identical ZF2 sequences with numbering schemes that differ by one residue, for consistency IKZF3-ZF2 numbering is used here). While residues N149, Q150 and C151 are proximal to pomalidomide, their backbones rather than their side chains are oriented towards pomalidomide, again making them unsuitable candidates for mutation in a bump-and hole approach. The conserved G152 residue is also close to pomalidomide, but cannot be mutated to an amino acid with a smaller side chain. We, therefore, decided to focus on a Q147A mutation (Figure 2B). This residue is located close to pomalidomide and is a key component of the CRBN-IKZF1 interface, forming a water-mediated hydrogen bond with E378 from CRBN.

**Figure 2.**
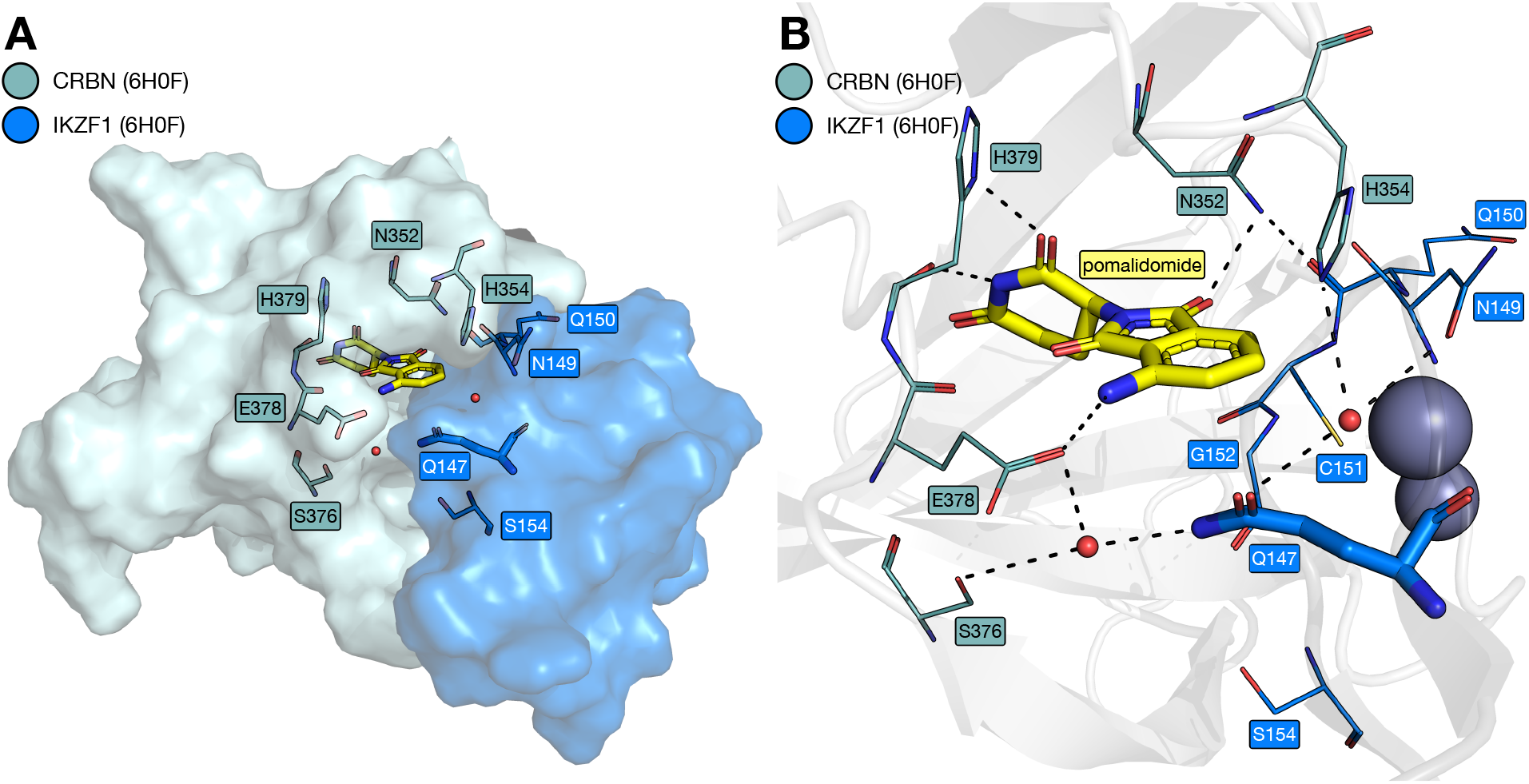
Structural details of the CRBN-IKZF1 ZF2 interface. **A**. Key residues at the interface between CRBN (teal) and IKZF1 ZF2 (blue) with pomalidomide (carbon = yellow) and two structured water molecules (red spheres) bound at the interface (PDB ID: 6H0F). **B.** Key predicted hydrogen bonds (dashed lines) formed between pomalidomide, two structured water molecules, CRBN (carbon = teal) and IKZF1-ZF2 (carbon = blue; PDB ID: 6H0F) (*6*).

Next, we designed and synthesized a set of 36 IMiD analogs. These compounds are modified at either the 4- or 5-position and were intended to probe the degradation structure-activity relationships (SAR) in this region of the molecule. We assessed the ability of each of these compounds to induce degradation of the mutant degrons using a ratiometric fluorescence-based assay (*6*). The degron spans residues 130-189 of IKZF3, which incorporates the minimal degron ZF2 (residues 146-168) and regions of the flanking ZFs on either side. The degron was fused to EGFP upstream from a red fluorescent protein (mCherry), separated by an internal ribosomal entry site (IRES). This construct was expressed in Jurkat cells, the cells incubated with a given compound for 18 h, and then analyzed using flow cytometry. EGFP:mCherry ratios from treated cell populations were then normalized against the EGFP:mCherry ratio from an untreated cell population to determine the level of compound-induced degradation. A reduction in EGFP, relative to mCherry, indicates that degradation is being induced (Figure 3).

**Figure 3.**
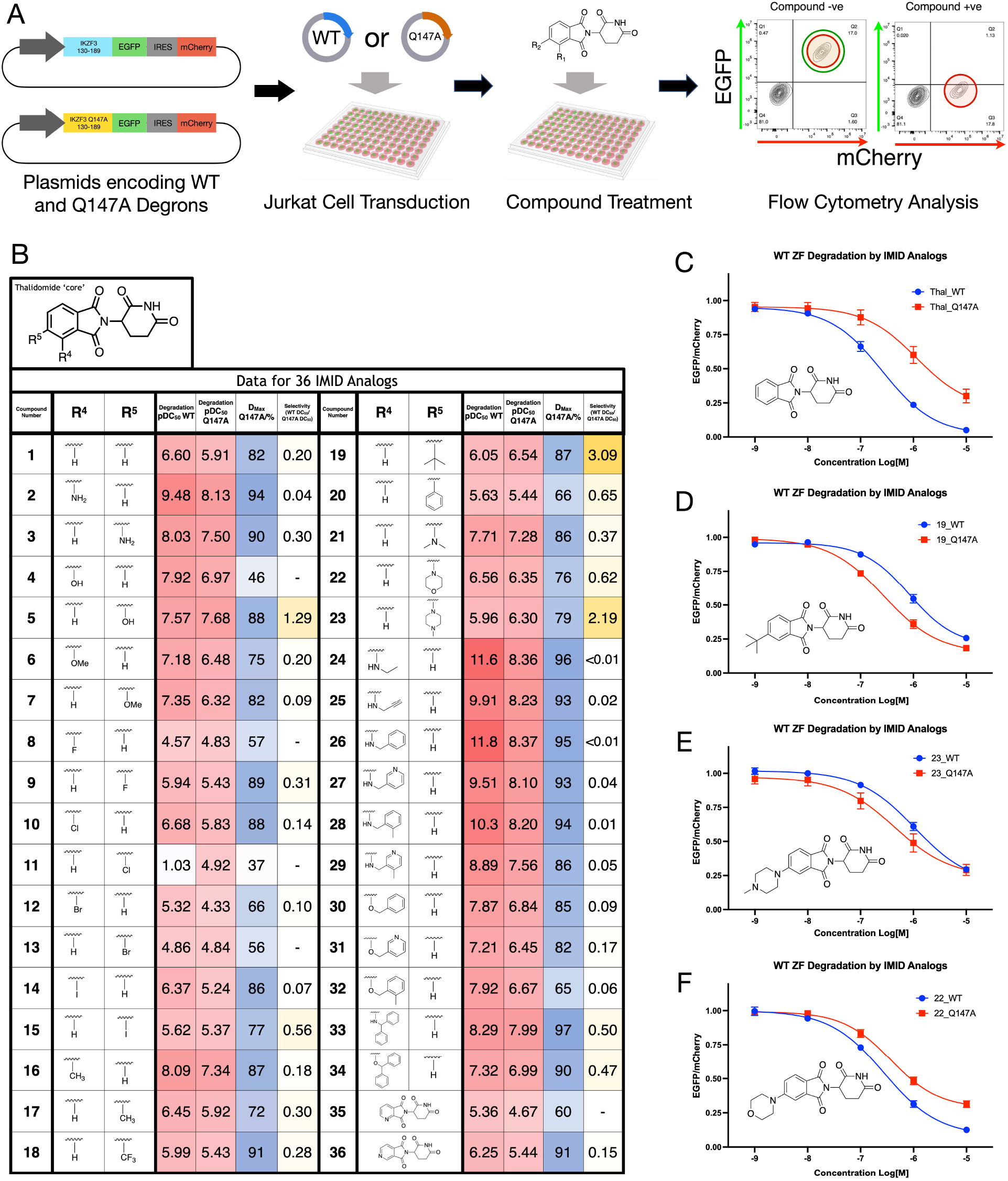
Degradation studies on the Q147A mutant. **A.** Schematic for the ratiometric flow cytometry assay used to assess degradation properties of 36 IMiD analogs against the WT IKZF3-derived degron and a Q147A mutant degron. **B.** Table showing pDC_50_ (WT), pDC_50_ (Q147A), D_MAX_ (Q147A) and selectivity values for 36 IMiD analogs. Selectivity value = DC_50_(WT)/DC_50_(Q147A). **C-F.** Jurkat cells stably expressing either WT or Q147A degrons fused to EGFP were treated with a concentration range of thalidomide (**C**), compound **19 (D)**, compound **23 (E)**, or compound **22 (F)** to yield degradation curves. n=3.

pDC_50_ values (−log_10_DC_50_, where DC_50_ = the compound concentration that induces 50% protein degradation) obtained for both the WT and Q147A mutant were compared for every compound exhibiting a Q147A D_max_ of >60% (D_max_ is the compound concentration at which maximum protein degradation is observed). This analysis revealed that the 4-position-modified IMiD analogs generally induced higher degradation of the WT degron compared to thalidomide. However, most of the 5-position-modified analogs were less potent degraders of the WT degron, compared to thalidomide, which is the directionality of change that we required (Figure 3B). Of particular interest were the 5-hydroxy (**5**), 5-*tert*-butyl (**19**; Figure 3D), and 5-*N*-methylpiperazine (**23**; Figure 3E) analogs, which all induced greater degradation of the Q147A-containing degron, compared to the WT degron. It is notable that compound **22**, which possesses a similarly sized ‘bump’ group to compounds **19** and **23**, preferentially degraded the WT degron over the Q147A mutant (Figure 3F). This indicates that protein-ligand interactions beyond simple steric bulk could be important for inducing the selective degradation observed. As 5-hydroxythalidomide (**5**) is a known degrader of SALL4, our studies focused on **19** and **23** (*17*).

This initial work served as a proof-of-concept that a bump and hole approach could be applied to the CRBN-IMiD-ZF system. Importantly, compounds **19** and **23** showed only moderate degradation of the WT degron, with pDC_50_ values of 6.05 and 5.96, respectively, compared to 6.60 for thalidomide and 9.48 for pomalidomide. Despite this, the selectivity gained from a single Q147A mutation was modest. We therefore decided to explore ZF mutations at a wider range of positions with the aim of identifying more selective IMiD-degron pairs.

### Design of a sequence library identifies 8380 mutants for evaluation

Inspection of the X-ray crystal structure of pomalidomide bound to CRBN and IKZF1 (PDB ID: 6H0F) shows that, in addition to Q147, residues N149, Q150, G152, A153, S154 F155, and L167 form important components of the IKZF1 protein interface with CRBN. The side chain of residue S154 is proximal to that of Q147. Residues N149, Q150, A153 and L167 interface directly with CRBN; residue A153 also forms part of a three-residue motif with residues F146 and F155, providing a ‘core’ to the ZF structure. Residue G152 is notable as being common to all IMiD-binding ZF degrons (*44*). The full library design is summarized in Figure 4A.

**Figure 4.**
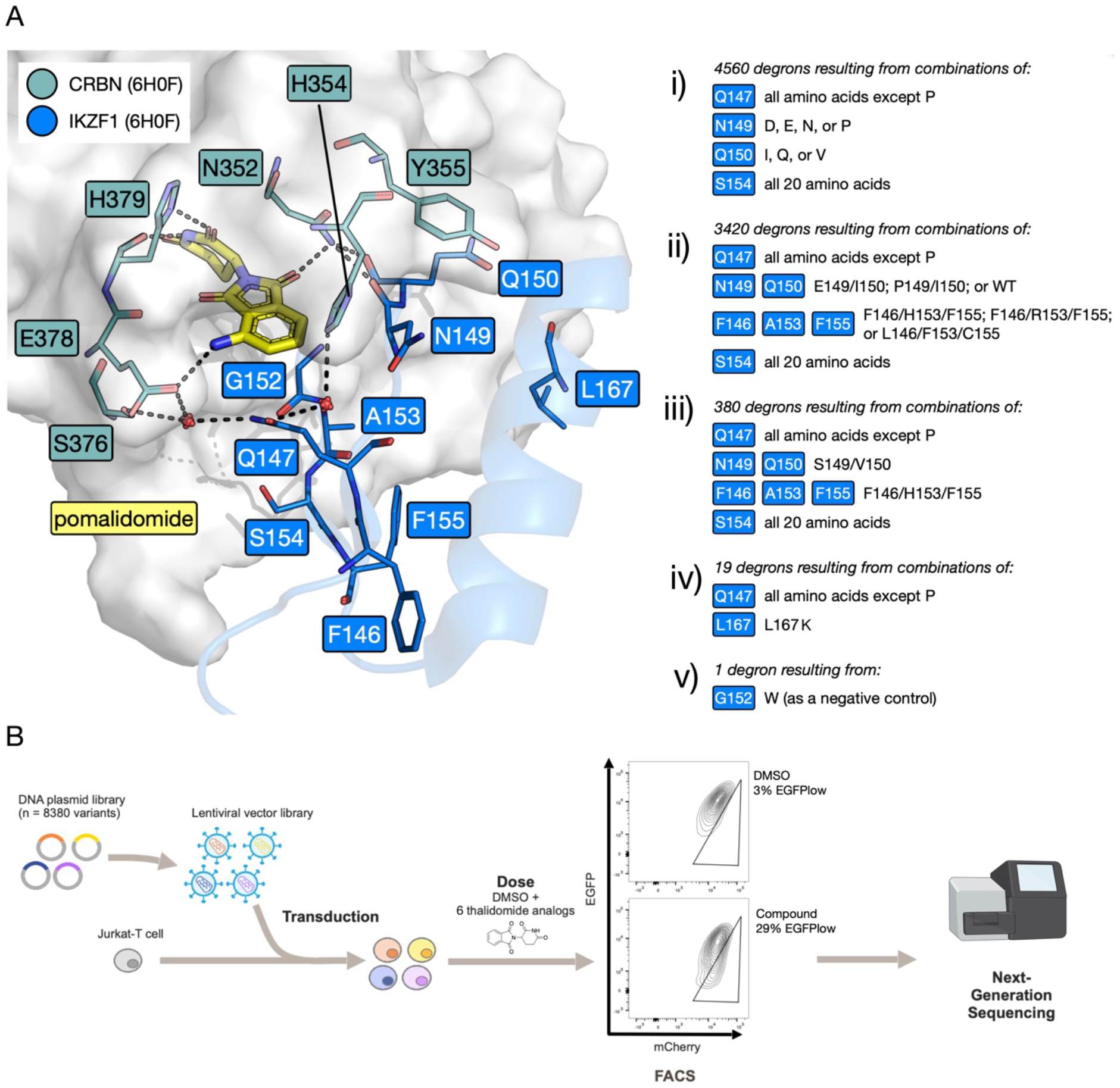
Design of the mutant library and the library screening workflow. **A.** Key residues at the interface between CRBN (grey surface; teal sticks) and IKZF1 ZF2 (blue cartoon and sticks) with bridging pomalidomide (yellow sticks) and waters (red spheres) (PDB ID: 6H0F) (*6*); key interactions shown as dashed lines. **i-v)** Description of mutations comprising the mutant ZF library. **B.** Schematic for the ZF degron library screen: plasmids encoding 8380 mutant ZFs were transduced into a population of Jurkat cells, cells were treated with one of 6 compounds – **1** (thalidomide), **19**, **22**, **23**, **33** or lenalidomide – or left untreated (DMSO control), then sorted by FACS into EGFP_+_ and EGFP_low_ populations; Next generation sequencing (NGS) was then performed on both populations.

Exploring all possible combinations of mutations at these nine positions would result in 512 billion combinations and was therefore impractical, so steps were taken to narrow down the number of mutants explored. Given our focus on position 147 we decided to explore all possible mutations, however, proline was not included as the backbone angles observed for position 147 in the X-ray crystal structure (Φ of −111.3°, Ψ of 148.4°) did not fall within the acceptable ranges for proline (Φ angle ≈ −60°; Ψ angle ≈ −45° or 135°). Similarly, position 154 was significant enough for all 20 proteinogenic amino acids to be included at this position.

To determine the ability of mutations at residues 149, 150, 153 and 167 to help stabilize the CRBN-ZF protein-protein interaction (PPI), Rosetta and Foldx softwares were employed to perform an *in silico* screen using a region of the CRBN-IMiD-ZF complex crystal structure (PDB ID: 6H0F) (*6*) in the absence of the IMiD (Figure S4) (*53*, *54*). These data, and inspection of known endogenous neosubstrate sequences, predicted that residues D, E, N, or P at position 149, and I, Q, or V at position 150 (Figure 4Ai) might help to stabilize the PPI. Using this approach, no stabilizing mutations were predicted at positions 153 and 167. However, mutations of residue 153 were considered in combination with mutations at positions 146 and 155. IKZF3 possesses the F146/A153/F155 (FAF) motif, but inspection of endogenous neosubstrate sequences showed that the FHF, FRF, and LFC motifs are also found at these positions. Consequently, these motifs were also explored in the mutant library (Figure 4Aii), with the idea that they might lead to ZF sequences showing enhanced interactions with CRBN (*6*, *44*). Another subset of mutations was included that focused on positions 147 and 154, while keeping residues 146, 149, 150, 153 and 155 the same as the equivalent positions in the ZF degron of SALL4 (F, S, V, H and F respectively) (Figure 4Aiii) (*16*).

Mutations at position 167 were deprioritized as no CRBN-ZF stabilizing mutations were identified in the *in silico* screen, but a small subset of mutations exploring 19 mutations at 147 (all except proline) alongside a L167K mutation was included, as inspection of endogenous neosubstrate sequences suggested a high frequency of lysine at position 167 in naturally occurring degrons (Figure 4Aiv) (*6*). Finally, a single G152W mutant was included as a negative control, as any residue other than glycine at this position interferes with IMiD-induced degradation (Figure 4Av) (*16*). Therefore, the total number of ZF sequences explored was 8380.

### A ratiometric fluorescence-based mutant library screen identifies degrons that undergo bumped IMiD-induced degradation

As we sought to screen a large number of mutant ZF degrons with different mutation combinations, we pursued a library screen approach similar to that previously employed by Sievers *et al*. and Jan *et al*. (*6*, *44*). Only the most promising compounds from the original set of 36 were used for the library screen. To help narrow down the candidates, a subset of compounds was tested for endogenous IKZF1 degradation in Jurkat cells using a fluorescent antibody flow cytometry assay (Figure S5). Consistent with the initial work, compounds **19**, **22**, **23** did not induce degradation of endogenous IKZF1 in this assay, and so were taken forward. While the diphenyl derivative **33** showed modest degradation of IKZF1 in this assay, it was progressed due to both its relatively high selectivity value in the initial screen, and its structural difference from compounds **19**, **22** and **23**. Thalidomide and lenalidomide were also included as controls. All compounds taken forward were observed to display suitable CRBN IC_50_ values and acceptable solubility in PBS buffer (Figure S3).

The library screen was carried out by transducing Jurkat cells with a lentiviral vector pool containing all 8380 sequences, using the same plasmid construct as the previous screen. Low level transduction (<30%) was carried out to maximize the number of cells with a single integration. An initial round of fluorescence-activated cell sorting (FACS) was used to isolate cells expressing mCherry; these mCherry positive (mCherry+) cells were then treated with one of the selected compounds for 18 h, and then sorted into 2 populations by FACS: EGFP_low_ and EGFP_+_. The EGFP_low_ population contains cells in which the EGFP-degron fusion protein is depleted due to compound-induced degradation. Next generation sequencing (NGS) was then conducted to determine which mutant ZF sequences were overrepresented in the EGFP_low_ population compared to the rest of the EGFP_+_ population. Data for all screens are a mean of 3 biological replicates (Figure 4B).

Raw library data (Figures S6 and S7) show that the three repeats of the compound screen correlate well, while the three DMSO control repeats exhibit lower correlation with each other. This observation indicates that highly similar sets of sequences are enriched for each repeat of the same compound. High correlation is also observed between screens of compounds with similar structures, such as between the three compounds with ‘bump’ groups at the 5 position (compounds **19**, **22** and **23**), or between thalidomide and lenalidomide. This observation indicates that structurally-related compounds induce degradation of similar sequence spaces.

Volcano plots for thalidomide and lenalidomide screens show that the WT degron sequence is overrepresented in the EGFP_low_ population indicating that, as expected, WT degron degradation is induced relatively efficiently by these compounds, compared to the rest of the library sequences (Figure 5A). However, the opposite is observed for the four bumped compounds (**19**, **22**, **23** and **33**), where the WT sequence is underrepresented in the EGFP_low_ population (Figures 5A and S8). As the −log10 *p*-values are generally high, it was decided that log_2_ fold change (log_2_FC) values could be used to assess all sequences going forward.

**Figure 5.**
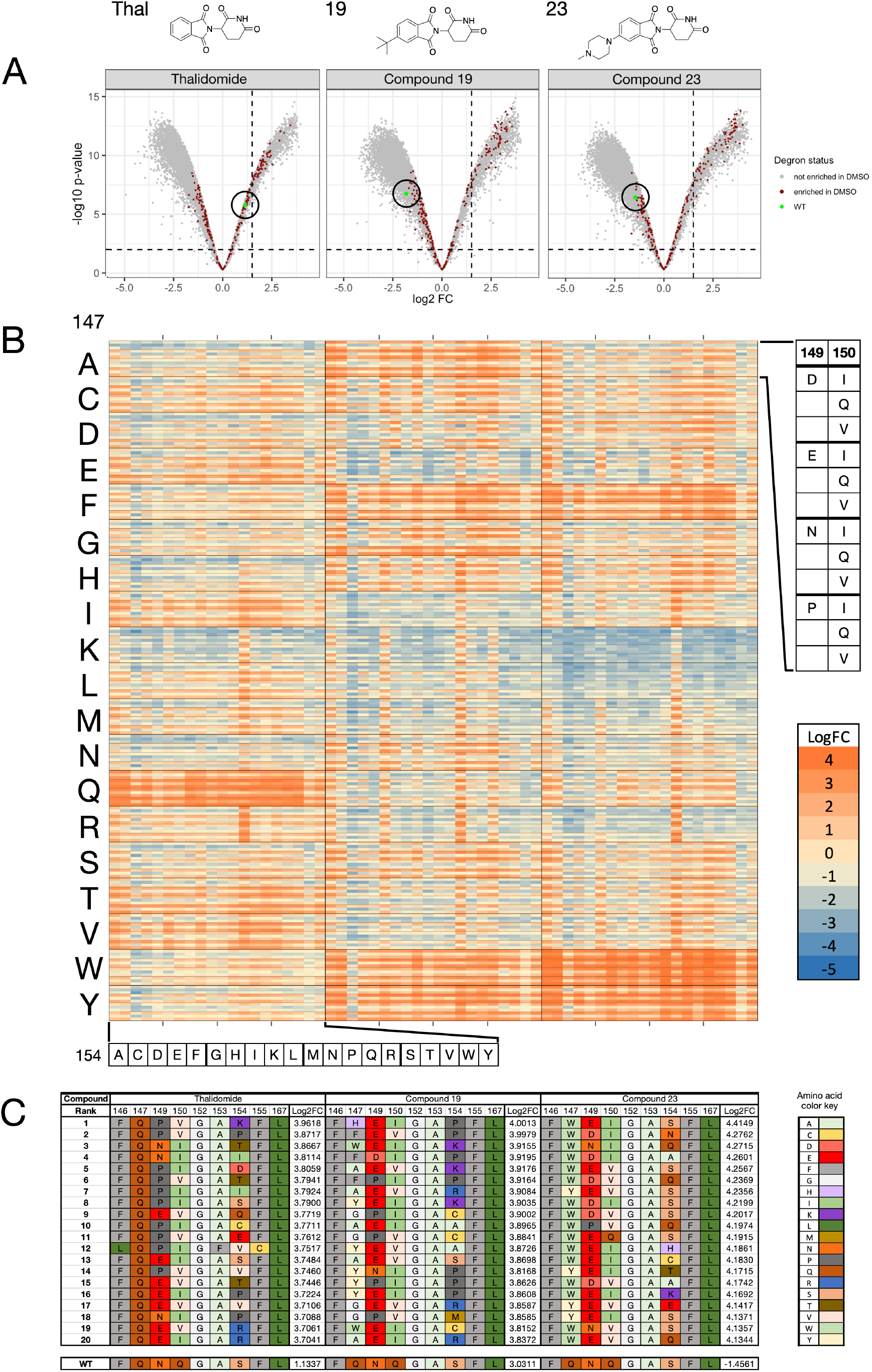
Data obtained from the library screen. **A.** Volcano plots for thalidomide (**1**), **19**, and **23** tested against the mutant library. Brown data points represent sequences that were enriched in the DMSO control; the green data point in each plot represents the WT sequence (no change in sequence from the original IKZF1/3 ZF degron). **B.** Log_2_FC data for thalidomide (**1**), **19**, and **23** tested against the 4560 library mutant ZF degrons in section i of the library. Log_2_FC values are arranged according to compound and residue 154 on the x-axis, and values are arranged according to residues 147, 149, and 150 on the y-axis. All other degron residues are identical to WT IKZF1/3 degron. Color scale shows high log_2_FC as orange, representing high sequence enrichment in the EGFP_low_ population, and low log_2_FC as blue, representing low sequence occurrence. A log_2_FC score of zero, colored as tan, represents equal representation of a sequence in the ‘degraded’ cell population and the remaining cell population after FACS. **C.** Table showing the 20 highest ranked mutant ZF sequences by log_2_FC, for compounds thalidomide (**1**), **19**, or **23**; the WT (IKZF1/3 ZF2) sequence and log_2_FC for each of the 3 compound screens are also shown for comparison.

We have conceptually divided the library into three sections, section i features the ‘FAF’ motif at positions 146, 153 and 155, while sections ii and iii have the ‘FHF’, ‘FRF’ and ‘LFC’ motifs these positions. Inspection of the full library data for library sections i, ii, and iii highlights several key observations (Figure S9). Library section i encompasses more sequences with high log_2_FC values compared to sections ii and iii, indicating that these sequences are effectively degraded when treated with the given IMiD. Although the ‘LFC’ section does have a large number of sequences with higher log_2_FC values, these sequences also tend have higher log_2_FC values for the DMSO control. This observation suggests that these sequences either show poor expression levels, or are susceptible to degradation in the absence of an IMiD derivative.

Within library section i thalidomide and lenalidomide preferentially degrade sequences that have the WT glutamine at position 147 (Figures 5B and S9). The *^t^*Bu derivative **19** exhibits the strongest preferences for ZFs with alanine, phenylalanine, tryptophan, or tyrosine at position 147. Smaller numbers of sequences with cysteine, glycine, histidine, or serine at this position are also efficiently degraded by this compound. Compounds **22** and **23** share similar strong overall preferences for ZFs with phenylalanine, tryptophan, or tyrosine at position 147. Interestingly, the *N*-methyl piperazine derivative **23** is especially efficient at degrading degrons with tryptophan at position 147. The diphenyl derivative **33** does not, however, exhibit a clear preference for a specific residue at position 147 (Figures 5B and S9).

For all compounds screened, there is a clear preference for either glutamic acid or proline over the WT asparagine at position 149; likewise, both isoleucine and valine are favored over the WT glutamine at position 150. However, the residue preference at position 154 across all compounds appears to be context dependent, and the trends are less straightforward. Residue preference trends for all compounds can be seen mirrored in the top 20 sequences ranked by log_2_FC for each compound screen (Figures 5C and S10).

Having identified the most efficiently degraded ZF degron sequences for each compound, the most selective compound-sequence pairs were chosen for validation using the same ratiometric fluorescence flow cytometry assay as previously used. Only sequences degraded by compounds **19** and **23** were progressed as these compound-degron pairings were expected to show the highest selectivity over the WT.

Sequences tested at this stage only have mutations at positions 147, 149, 150 or 154, and so are named based on their residues at these positions, for example, the Q147A mutant has A at 147, N at 149, Q at 150 and S at 154, and so is called ANQS.

Compound **19** degrades the HEIP and AEVK sequences most effectively, with these compound-degron pairs showing 25-fold and 28-fold selectivity over WT, respectively (Figures 6A, 6B and S11E). It is notable that all five the library sequences degraded most efficiently by compound **19**, ranked by log_2_FC, possess an acidic residue (either D or E) at position 149. To confirm our earlier results, compound **19** was also tested against the single Q147A mutant sequence ANQS, with only 1.4-fold selectivity over WT observed in this case.

**Figure 6.**
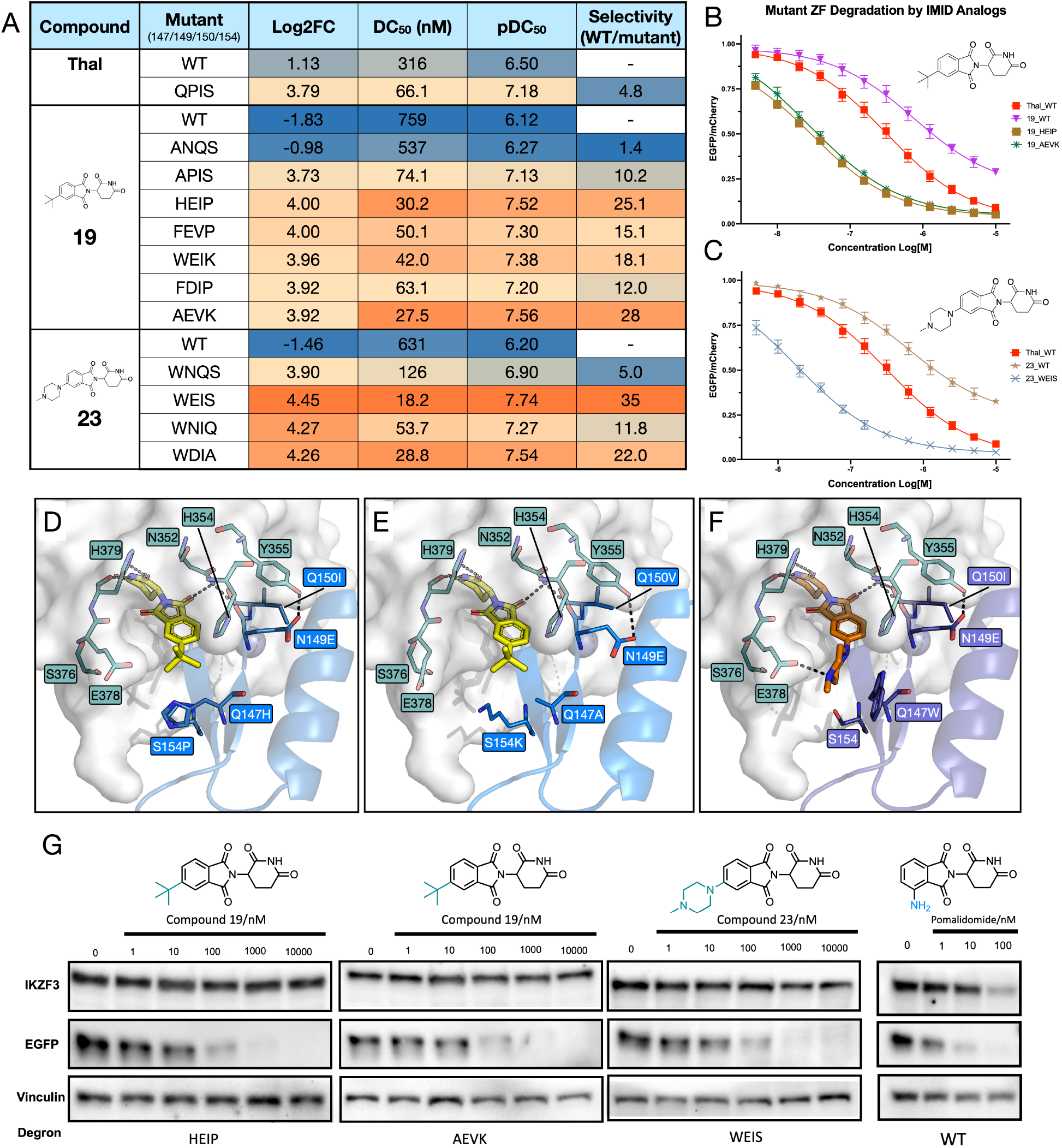
Investigation of the selectivity for certain degrons. **A.** Table showing library log_2_FC, DC_50_, pDC_50_ and selectivity values for different compound-mutant ZF degron combinations; selectivity = DC_50_(WT)/DC_50_(mutant). **B.** Jurkat cells stably expressing either WT or mutant (HEIP or AEVK) ZF degrons fused to EGFP were treated with a concentration curve of thalidomide or compound **19** to yield degradation curves. **C.** Jurkat cells stably expressing either WT or WEIS mutant ZF degrons fused to EGFP were treated with a concentration curve of thalidomide or compound **23** to yield degradation curves. **D.** Docking model of compound **19** at the interface between CRBN and the HEIP mutant ZF. **E.** Docking model of compound **19** at the interface between CRBN and the AEVK mutant ZF. **F.** Docking model of compound **23** at the interface between CRBN and the WEIS mutant ZF; all three models derived from crystal structure of CRBN-pomalidomide-IKZF1 ZF2 (PDB ID: 6H0F) (*6*). **G.** Western blots performed on transduced Jurkat cells expressing a ZF-EGFP fusion protein with either the HEIP, AEVK, WEIS or WT degron, for five concentrations of compound **19** or compound **23**, or 3 concentrations of pomalidomide, and an untreated DMSO control. Bands show levels of IKZF3, EGFP and vinculin loading control.

The *N*-methyl piperazine derivative **23** degrades the triple mutant sequence WEIS most effectively, with this ligand-degron pair showing the highest selectivity we observed in this study; 35-fold over WT. Interestingly, most sequences for which **23** efficiently induced degradation possess a tryptophan residue at position 147. Even the single Q147W mutant, WNQS, shows 5-fold selectivity compared to WT, indicating that incorporation of this residue at position 147 is favorable for **23** stabilizing the CRBN-**23**-IKZF1/3 ternary complex (Figures 6A and S11D). Like the sequences above, WEIS also incorporates an acidic glutamate residue at position 149, again suggesting that this is beneficial for CRBN-compound-IKZF1/3 ternary complex stability. As W147 is a relatively large residue, this suggests that compound **23** forms a favorable interaction with W147 that stabilizes the CRBN-IMiD-ZF, resulting in enhanced degradation of the ZF-fusion protein.

Thalidomide was tested against an N149P-Q150I double mutant (QPIS), to assess the effect of including mutations that were predicted to be favorable at these positions. Thalidomide induced degradation of the QPIS degron with 4.8-fold selectivity over WT (Figures 6A and S11B). Interestingly, a similar result was observed when compound **19** was tested against the Q147A-N149P-Q150I triple mutant, APIS. Compound **19** induced degradation of the APIS degron 10-fold more effectively compared to the WT, which is a substantial improvement over the 1.4-fold selectivity displayed by ANQS. This demonstrates that these mutations increase degradation induced by the IMiD, despite these residues not being located immediately proximal to the IMiD binding site.

Docking studies were carried out to assess the potential structure of the CRBN-IMiD-ZF interfaces for the three most selective compound-mutant pairings: **19**-HEIP, **19**-AEVK, and **23**-WEIS (Figures 6D, 6E and 6F). FoldX 5 was used to generate the mutant structures from the DDB1-CRBN-pomalidomide complex bound to IKZF1(ZF2) (PDB ID: 6H0F). The bumped ligands were docked into these structures using GOLD.

In all three cases, a hydrogen bond is predicted to form between E149 in the ZF and Y355 of CRBN. The WT N149 is not observed to form this interaction in PDB ID: 6H0F, suggesting that this additional interaction stabilizes the CRBN-ZF protein-protein interaction (PPI). WT Q150 does not form any polar interactions with CRBN, but its side chain forms part of the hydrophobic interface between the ZF and CBRN. HEIP and WEIS possess a Q150I mutation at this position, while AEVK has a Q150V mutation. This observation suggests that the I150 or V150 residues replace the WT Q150 hydrophobic interactions, again helping to stabilize the CRBN-ZF PPI.

AEVK possesses a Q147A mutation, which seems to have a classic bump and hole effect, where the smaller A147 residue can accommodate the larger *^t^*Bu moiety of compound **19**. This degron incorporates a S154K mutation, which is spatially adjacent to position 147, with K154 predicted to form an ionic interaction with CRBN E378. This interaction places the lysine side chain away from the bumped IMiD and also likely further stabilizes the CRBN-ZF PPI. It interesting to note that the three additional mutations are having a substantial effect as the AEVK degron has 28-fold selectivity *vs* WT, compared to only 1.4-fold selectivity for the simple Q147A mutant ANQS. The HEIP degron has Q147H and S154P mutations. Docking studies predict that the conformationally restricted P154 residue enables H147 to move, accommodating the *^t^*Bu bump of compound **19**. The WEIS degron, which is selectively degraded by compound **23** possesses only three mutations, as S154 is the same as WT. The E149 and I150 mutations are thought to have the same effect as in the HEIP degron above, and serve to stabilize the CRBN-ZF PPI. At pH 7.4 *N*-methyl-piperazine amine of **23** is predicted to be >93% protonated on the methylated piperazine nitrogen (Chemicalize), and docking suggests that this moiety will form a salt bridge with CRBN E378. This salt bridge orients the piperazine ring so that it can form a cation-π interaction with W147. This observation explains why most of the mutants that are preferentially degraded by compound **23** possess a tryptophan at position 147.

It is interesting to note that in the WT DDB1-CRBN-pomalidomide complex bound to IKZF1(ZF2) (PDB ID: 6H0F) two structured water molecules are observed at the CRBN-ZF interface, forming hydrogen bonds with CRBN S376 and E378, and IKZF1 Q147 and N149. The structures predicted by our docking studies indicate that these water molecules will not be accommodated in the CRBN-IMiD-mutant ZF ternary complexes. It is therefore possible that, in addition to forming new interactions with the mutant degrons, the bumped ligands are being accommodated by the loss of two water molecules at the CRBN-ZF interface. We note that this is, to the best of our knowledge, the first example of a bump and hole approach on a ternary complex, and therefore the effect of the mutant combinations are harder to predict than in a ligand-protein binary complex setting. Consequently, while we do have an example of a Q147A mutant, where a smaller residue is included at position 147, we also have an example of a Q147W mutant, where W is at least the same size as Q, if not larger.

Analysis using western blotting, on a pre-sorted population of mCherry+ cells, confirmed that compounds **19** or **23** induced complete degradation of EGFP tagged with either the HEIP (**19**), AEVK (**19**), or WEIS (**23**) mutant degrons in transduced Jurkat cells, while endogenous IKZF3 levels remained unaffected (Figures 6G, S12, S13 and S14). In Jurkat cells transduced with the WT degron-tagged EGFP, pomalidomide induced degradation of EGFP but also depleted endogenous IKZF3. In untransduced Jurkat cells, compounds **19** and **23** do not induce degradation of endogenous IKZF3, but, as expected, pomalidomide did cause IKZF3 degradation (Figures S15 and S16).

As the **23**-WEIS compound-mutant pairing had exhibited the highest selectivity value, we conducted tandem mass tag (TMT) mass spectrometry proteomics to determine the whole cell selectivity of compound **23**. Quantitative proteomics screens were carried out in untransduced Jurkat cells for compound **23** (10 μM), thalidomide (10 μM), lenalidomide (1 μM) and pomalidomide (1 μM) for 16 h using a 15-plex TMTpro SPS-MS (*55*) workflow (Figure S19). Compound **23** (10 μM) had no major effect on any of the 9644 quantified proteins (Figure 6A), while 1 μM of pomalidomide induced significant (Welch’s t-test) degradation of at least 20 proteins (Figures 7A and 7B). Thalidomide did not show any major proteome activity and lenalidomide induced significant (Welch’s t-test) degradation of at least 6 proteins (Figure S17).

**Figure 7.**
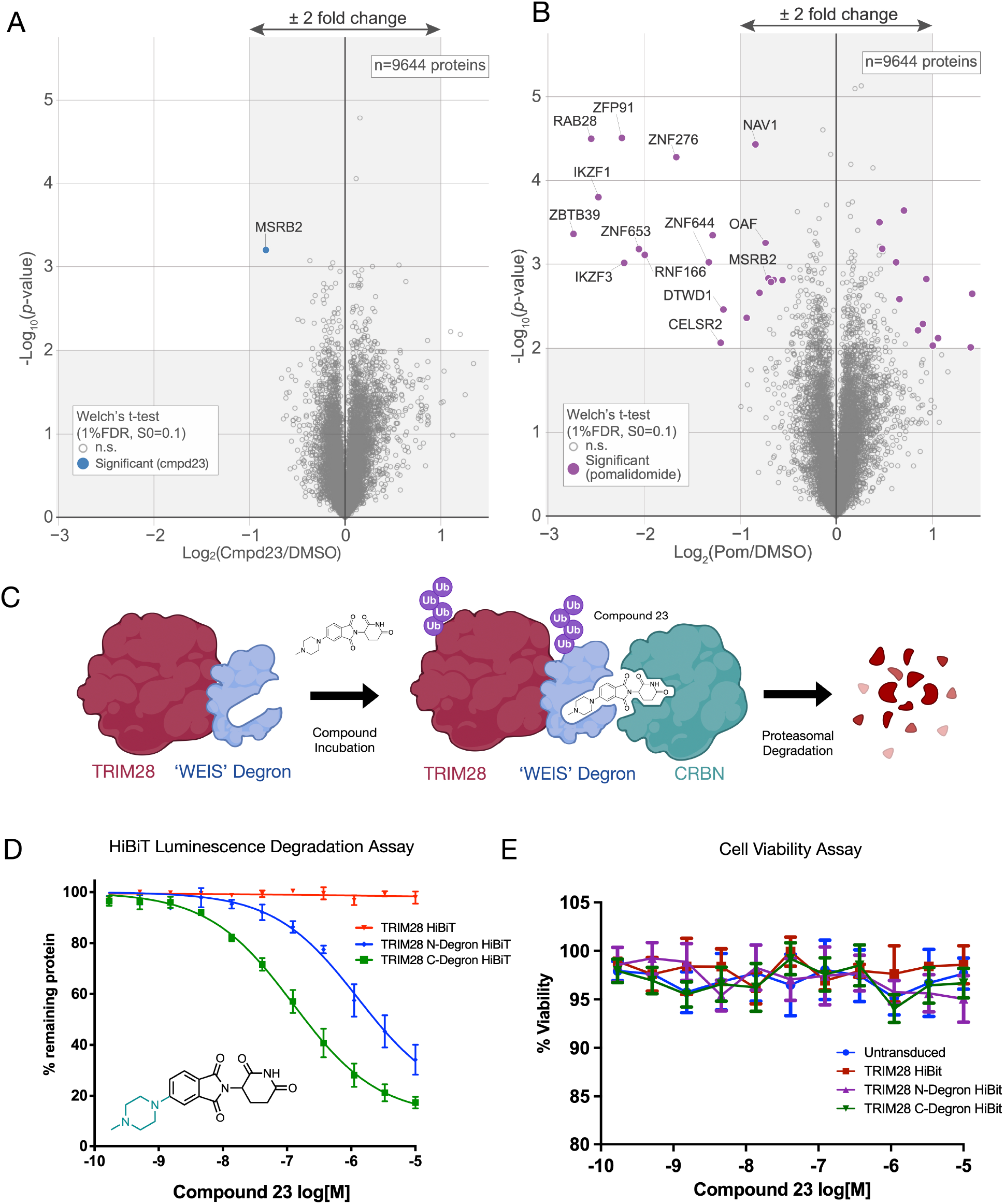
Proteomics data and application of the WEIS degron to TRIM28 degradation. **A-B.** Quantitative proteomics profiling of compound **23** (10 μM (**A**) and pomalidomide (1 μM (**B**). Jurkat cells were treated for 16 h with the corresponding compound or DMSO control, and protein abundance was analyzed using multiplexed TMTpro quantification mass spectrometry. log_2_FC is shown on the x-axis, and −log_10_(*p*-value) is shown on the y-axis. The values shown are an average of three biological replicates. **C.** Schematic of degradation of TRIM28 tagged with the WEIS degron by compound **23**. **D.** Jurkat cells were transduced with TRIM28 with either N-terminal or C-terminal WEIS degron tag (or untagged control) and HiBiT peptide tag, then incubated with a concentration curve of compound **23** (100 pM – 10 μM) for 24 h and analyzed using a HiBiT luminescence assay. **E.** Cell Titre Glo assay on transduced Jurkat cells shows cell viability.

Having demonstrated the selectivity of the **23**-WEIS degron system, we decided to apply it to induce degradation of TRIM28, a disease-relevant bromodomain- and PHD-containing protein, for which there are no ligands (*56*). Jurkat cells were CRISPR edited to knockout the endogenous TRIM28 gene. Genes for TRIM28 tagged with the WEIS mutant ZF degron at either the N- or C-terminus (or untagged), and also tagged with HiBiT at the other terminus, were then transduced into Jurkat cells and incubated with compound **23** for 24 h. The linker region used was the same as for the previously tested EGFP fusion. Degradation levels were then assessed using a HiBiT luminescence assay. Compound **23** induced degradation of both the N- and C-terminus tagged fusion protein, with DC_50_ values of 2340 nM and 190 nM, respectively. For the C-terminal tagged protein a D_max_ value of >80% was observed with 10 μM of **23**. This result demonstrates that our technology can be used to efficiently induce degradation of an unliganded protein. The difference in degradation observed between the N- and C-terminal tagged fusions proteins hints that ubiquitination is occurring on TRIM28 and not the degron or linker. We note that no optimization of the TRIM28-ZF linker was undertaken, and more potent degradation could likely be achieved through investigating the linker region.

## Conclusion

Induced protein degradation has emerged as a powerful tool to assist validation of a putative therapeutic target. Key considerations in ensuring this approach is as effective as possible are the size of the degron tag, the physicochemical properties of the small molecule inducer of degradation, and the selectivity for induced degradation for the POI over other cellular proteins. Here we report a new chemically-inducible degron system with distinct advantages over the current state-of-the-art, that addresses these key points. The most selective degron-IMiD pair that we have identified is WEIS combined with compound **23**. This degron is only 60 amino acids, with a molecular weight of 7.0 kDa, and can be easily attached to the N- or C-terminus of a POI. Compound **23** has a molecular weight of 356 and solubility forecast index (SFI) of 0.27, correlating with high solubility and cell permeability (*57*). The smaller degron and chemical properties of the IMiD analogs described here offer advantages over systems such as dTAG (degron = 11.9 kDa), HaloTag (degron = 33.6 kDa), Bromotag (degron = 14.9 kDa), which employ much larger bifunctional molecules and degrons. The small size of both the mutant WEIS degron (7.0 kDa) and the bumped IMiD analog **23** offer solutions to challenges associated with low efficiency of endogenous degron tagging using CRISPR Cas9 knock-in, or effects of large tags on protein function. In addition the small molecule inducers have higher cell penetration than the bifunctional molecules affiliated with systems such as dTAG and HaloPROTAC.

The bump-and-hole approach that we have employed minimizes degradation of endogenous neosubstrates that are affected by established IMiDs such as thalidomide, lenalidomide, and pomalidomide. TMT proteomics studies have confirmed the high selectivity of the bumped IMiD analog **23** in Jurkat cells, a model cancer T cell line. The library screen approach we adopted has not only identified a degron design that is complementary to compound **23**, but has also afforded insight into the nature of the CRBN-IMiD-ZF interface that is central to the mode of action of all IMiDs.

We anticipate that this new degron system will be a powerful addition to the existing approaches available for target validation studies. We have shown that this degron system can be used to efficiently degrade TRIM28, a disease-relevant protein for which no small molecule ligands are known, even without optimization of the degron size or nature of the linker region. This approach can be applied to other POIs that do not possess small molecule ligands, as a method of testing their suitability as therapeutic targets. Importantly, the small size of our degron tag makes it less likely to be ubiquitinated and drive degradation independently from the POI, which is common for larger tags. This attribute makes it suitable for evaluating POI degradability. This system could be especially useful in scenarios where degradation of typical IMiD neosubstrates, such as IKZF1 or IKZF3, is undesirable, such as in CAR T cell therapy. Application of this system to the CAR degron approaches reported by Jan *et al*. and Carboneau *et al*. (*44*, *46*) could mitigate any unwanted effects observed from the use of established IMiD molecules, while retaining the ability to rapidly and reversibly degrade CARs, and therefore control cytotoxicity of infused CAR T cells in patients (*44*, *46*).

## Acknowledgments

MS thanks Corpus Christi College, Oxford, for research support. SJC is grateful to Michael and Alice Jung for endowing the Jung Chair in Medicinal Chemistry and Drug Discovery at UCLA, which partially supported this work. We thank A. Ordureau for proteomics support and expertise.

## Author contributions

Conceptualization: SJC, LLB, MAQ, MMH

Methodology: PJB, LLB, SJC, MMH, RES, MSp, AK, LG, SBB, AS, CMF, ABB, MAQ, HA, LS

Investigation: PJB, LLB, LS, MSp, AK, LG, SBB, RDB, MSe Visualization: PJB, SJC, RES, GPH, LLB

Data curation: PJB, CMD, LLB Funding acquisition: SJC, HA

Supervision: SJC, LLB, MMH, CMD, HA Project administration: SJC, LLB

Writing – original draft: PJB, LLB

Writing – review & editing: SJC, PJB, LLB, MMH

## Data and materials availability

All data are available in the main text or the supplementary materials.

## Supplementary Materials

Figs. S1 to S19

Chemistry Experimental Section

## Materials and Methods

Plasmid Information

Sequences for Next Generation Sequencing

Computational Methods

NMR Spectra for Compounds

HPLC Traces for Compounds

Degron Screen Data

