## Supplementary Materials for "Orthogonal IMiD-Degron Pairs Induce Selective Protein Degradation in Cells"

**The PDF file includes:**

Figs. S1 to S17
Materials and Methods
Supplementary Text

**Other Supplementary Materials for this manuscript include the following:**

Figs. S18 and S19
Data S1 (Separate File)

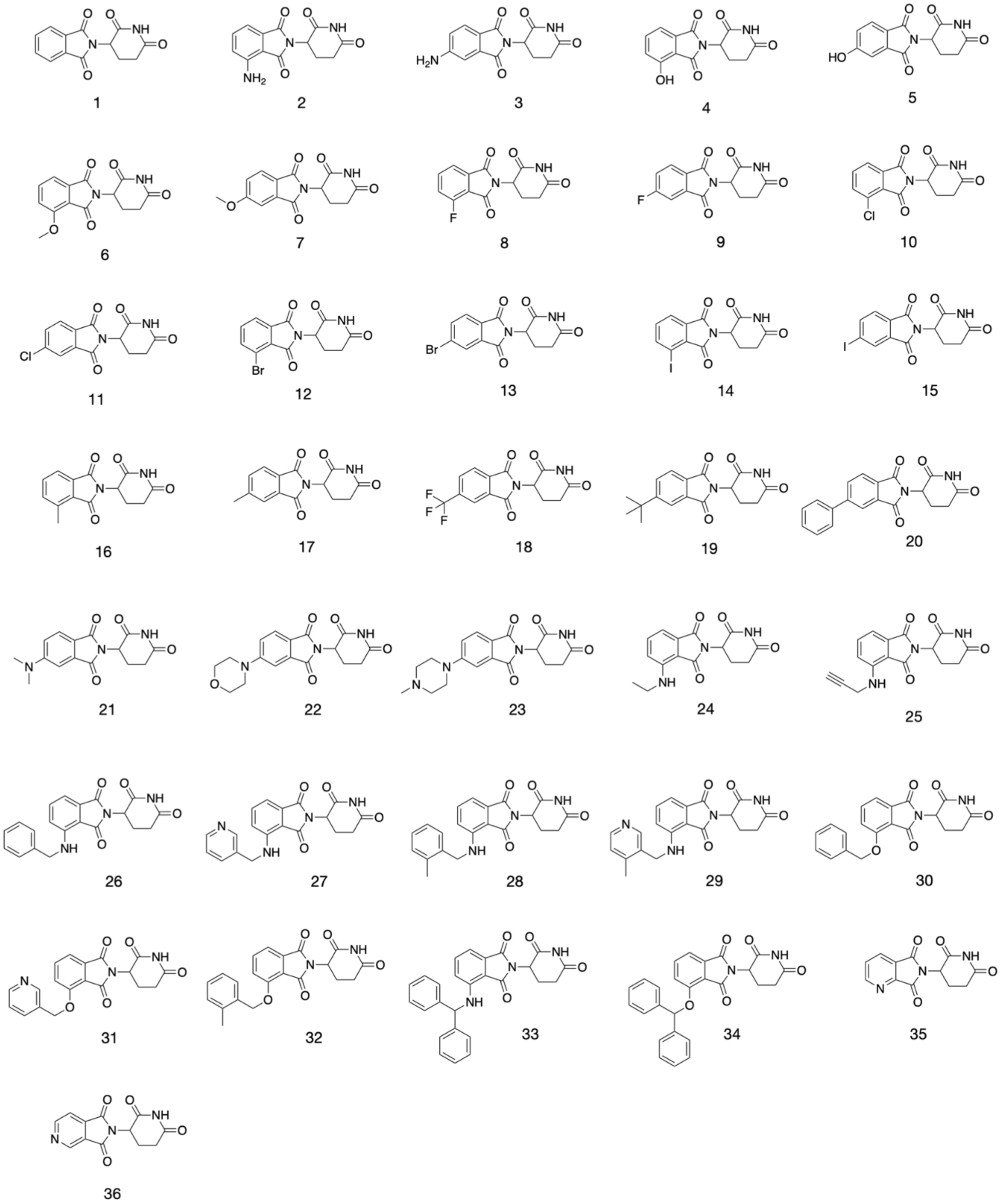

**Figure S1.** Full compound library of 36 IMiD analogs used in biological testing.

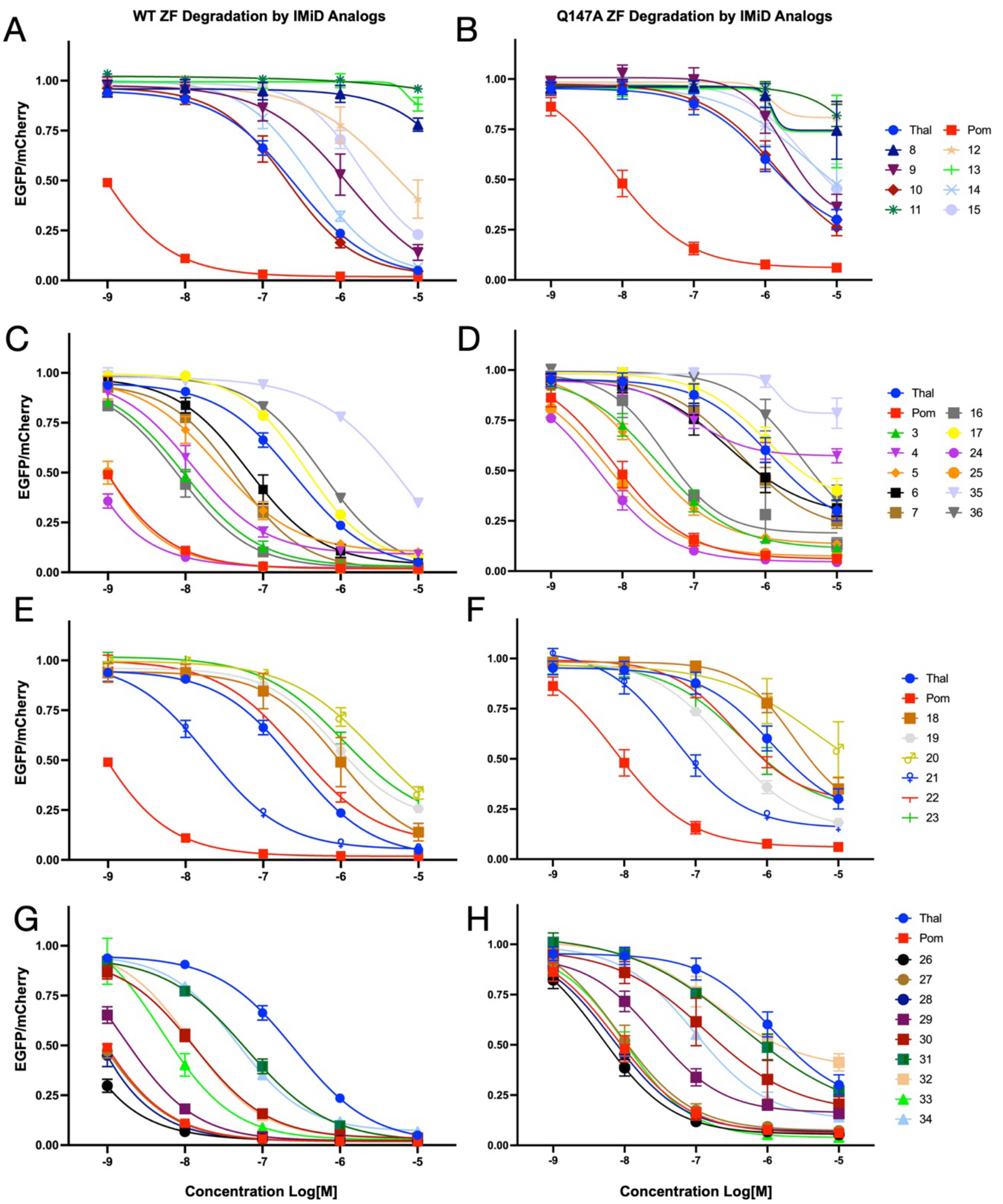

**Figure S2.**

**A.** Degradation curves for halogenated IMiD analogs against WT ZF degron. **B.** Degradation curves for halogenated IMiD analogs against Q147A ZF degron. **C.** Degradation curves for IMiD analogs with small bump groups against WT ZF degron. **D.** Degradation curves for IMiD analogs with small bump groups against Q147A ZF degron. **E.** Degradation curves for IMiD analogs with large bump groups at the 5-position against WT ZF degron. **F.** Degradation curves for IMiD analogs with large bump groups at the 5-position against Q147A ZF degron. **G.** Degradation curves for analogs of compound 26 against WT ZF degron. **H.** Degradation curves for analogs of compound 26 against Q147A ZF degron.

Thalidomide 'core'

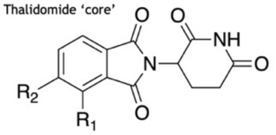

**Data for 36 IMID Analogs**

| Coumpound Number | $R_1$ | $R_2$ | Cereblon $pIC_{50}$ | Solubility /µM | Coumpound Number | $R_1$ | $R_2$ | Cereblon $pIC_{50}$ | Solubility /µM |
|---|---|---|---|---|---|---|---|---|---|
| 1 | H | H | 6.90 | 321 | 19 | H | 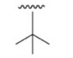 | 7.07 | 265 |
| 2 | $NH_2$ | H | 7.19 | 341 | 20 | H | 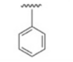 | 7.06 | 34 |
| 3 | H | $NH_2$ | 7.32 | 301 | 21 | H | 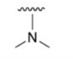 | 7.00 | 254 |
| 4 | OH | H | 7.57 | 342 | 22 | H | 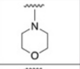 | 6.96 | 254 |
| 5 | H | OH | 7.39 | 348 | 23 | H | 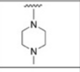 | 7.00 | 287 |
| 6 | OMe | H | 7.02 | 387 | 24 | 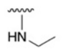 | H | 7.47 | 307 |
| 7 | H | OMe | 7.08 | 287 | 25 | 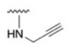 | H | 7.78 | 389 |
| 8 | F | H | 6.62 | 162 | 26 | 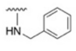 | H | 7.79 | 54 |
| 9 | H | F | 6.73 | 191 | 27 | 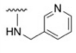 | H | 7.61 | 377 |
| 10 | Cl | H | 6.86 | 111 | 28 | 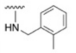 | H | 7.85 | 20 |
| 11 | H | Cl | 6.47 | 53 | 29 | 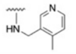 | H | 7.64 | 257 |
| 12 | Br | H | 7.10 | 23 | 30 | 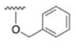 | H | 7.57 | 7 |
| 13 | H | Br | 6.58 | 9 | 31 | 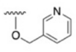 | H | 7.08 | 22 |
| 14 | I | H | 7.23 | 18 | 32 | 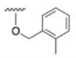 | H | 7.65 | 17 |
| 15 | H | I | 6.79 | 153 | 33 | 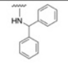 | H | 7.61 | 1 |
| 16 | $CH_3$ | H | 7.14 | 48 | 34 | 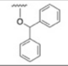 | H | 7.48 | 2 |
| 17 | H | $CH_3$ | 7.12 | 279 | 35 | 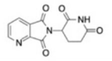 | | 6.00 | 2 |
| 18 | H | $CF_3$ | 6.32 | 69 | 36 | 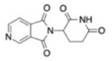 | | 6.05 | 1 |

**Figure S3.**

CRBN $pIC_{50}$ and solubility values (in PBS buffer) for all IMiD analog library compounds.

| | ΔΔG | | | | ΔΔG | | | | ΔΔG | | | | ΔΔG | | |
|---|---|---|---|---|---|---|---|---|---|---|---|---|---|---|---|
| Mutation | Foldx | Rosetta | Mean | Mutation | Foldx | Rosetta | Mean | Mutation | Foldx | Rosetta | Mean | Mutation | Foldx | Rosetta | Mean |
| N149A | -0.26 | -0.23 | -0.25 | Q150A | -0.22 | -1.51 | -0.87 | A153A | 0.01 | N/A | 0.01 | L167A | 0.41 | 1.61 | 1.01 |
| N149C | 0.02 | 1.76 | 0.89 | Q150C | 0.20 | 1.46 | 0.83 | A153C | 2.32 | N/A | 2.32 | L167C | 0.46 | 3.64 | 2.05 |
| N149D | -0.91 | -1.59 | -1.25 | Q150D | 0.53 | 1.30 | 0.92 | A153D | 3.76 | N/A | 3.76 | L167D | 1.34 | 2.91 | 2.13 |
| N149E | -1.28 | -0.71 | -0.99 | Q150E | -0.34 | -1.27 | -0.81 | A153E | 5.33 | N/A | 5.33 | L167E | 0.81 | 2.67 | 1.74 |
| N149F | -0.58 | 1.98 | 0.70 | Q150F | -1.11 | 4.65 | 1.77 | A153F | 23.77 | N/A | 23.77 | L167F | 0.29 | 3.04 | 1.67 |
| N149G | 0.45 | 2.50 | 1.48 | Q150G | 0.46 | 2.66 | 1.56 | A153G | 1.42 | 5.32 | 3.37 | L167G | 0.71 | 3.49 | 2.10 |
| N149H | 0.74 | 0.98 | 0.86 | Q150H | -0.59 | 1.44 | 0.43 | A153H | 49.24 | N/A | 49.24 | L167H | 0.85 | 2.26 | 1.55 |
| N149I | -0.23 | 1.89 | 0.83 | Q150I | -0.99 | -2.31 | -1.65 | A153I | 8.08 | N/A | 8.08 | L167I | 0.38 | 1.42 | 0.90 |
| N149K | -0.91 | -0.59 | -0.75 | Q150K | -0.92 | 1.36 | 0.22 | A153K | 8.20 | N/A | 8.20 | L167K | 0.53 | 0.96 | 0.75 |
| N149L | -0.58 | 3.55 | 1.48 | Q150L | -1.32 | 2.36 | 0.52 | A153L | 5.25 | N/A | 5.25 | L167L | 0.00 | N/A | 0.00 |
| N149M | -0.29 | 1.65 | 0.68 | Q150M | -1.30 | 1.63 | 0.16 | A153M | 4.46 | N/A | 4.46 | L167M | -0.05 | 1.32 | 0.63 |
| N149N | 0.00 | N/A | 0.00 | Q150N | 0.47 | 1.47 | 0.97 | A153N | 5.33 | N/A | 5.33 | L167N | 0.83 | 1.17 | 1.00 |
| N149P | -1.49 | -2.19 | -1.84 | Q150P | 0.60 | N/A | 0.60 | A153P | 5.46 | N/A | 5.46 | L167P | 0.50 | N/A | 0.50 |
| N149Q | -0.56 | -0.19 | -0.38 | Q150Q | 0.00 | N/A | 0.00 | A153Q | 7.81 | N/A | 7.81 | L167Q | 0.11 | 1.79 | 0.95 |
| N149R | -0.71 | -0.04 | -0.38 | Q150R | -0.27 | 1.71 | 0.72 | A153R | 15.58 | N/A | 15.58 | L167R | 0.23 | 0.89 | 0.56 |
| N149S | 0.41 | -0.25 | 0.08 | Q150S | 0.15 | -0.09 | 0.03 | A153S | 1.99 | 4.85 | 3.42 | L167S | 0.26 | 1.18 | 0.72 |
| N149T | 0.72 | 0.04 | 0.38 | Q150T | 0.02 | -1.06 | -0.52 | A153T | 4.97 | N/A | 4.97 | L167T | 0.04 | 2.25 | 1.14 |
| N149V | 0.12 | 0.51 | 0.32 | Q150V | -0.64 | -1.37 | -1.00 | A153V | 4.64 | N/A | 4.64 | L167V | 0.83 | 2.33 | 1.58 |
| N149W | 0.11 | 2.13 | 1.12 | Q150W | -0.94 | 3.42 | 1.24 | A153W | 22.72 | N/A | 22.72 | L167W | 0.56 | 0.93 | 0.74 |
| N149Y | 0.19 | 1.22 | 0.71 | Q150Y | 0.28 | 3.88 | 2.08 | A153Y | 25.84 | N/A | 25.84 | L167Y | 0.08 | 1.57 | 0.82 |

Favourable Mutation

Unfavourable Mutation

**Figure S4.**
Change in predicted free energies for binding of CRBN with the IKZF3 ZF2 degron when incorporating single mutations at positions 149, 150, 153, or 167 of the ZF, predicted by Foldx and Rosetta. Color scale highlights mutations resulting in a more stable complex in orange, and mutations resulting in less stable complex in blue.

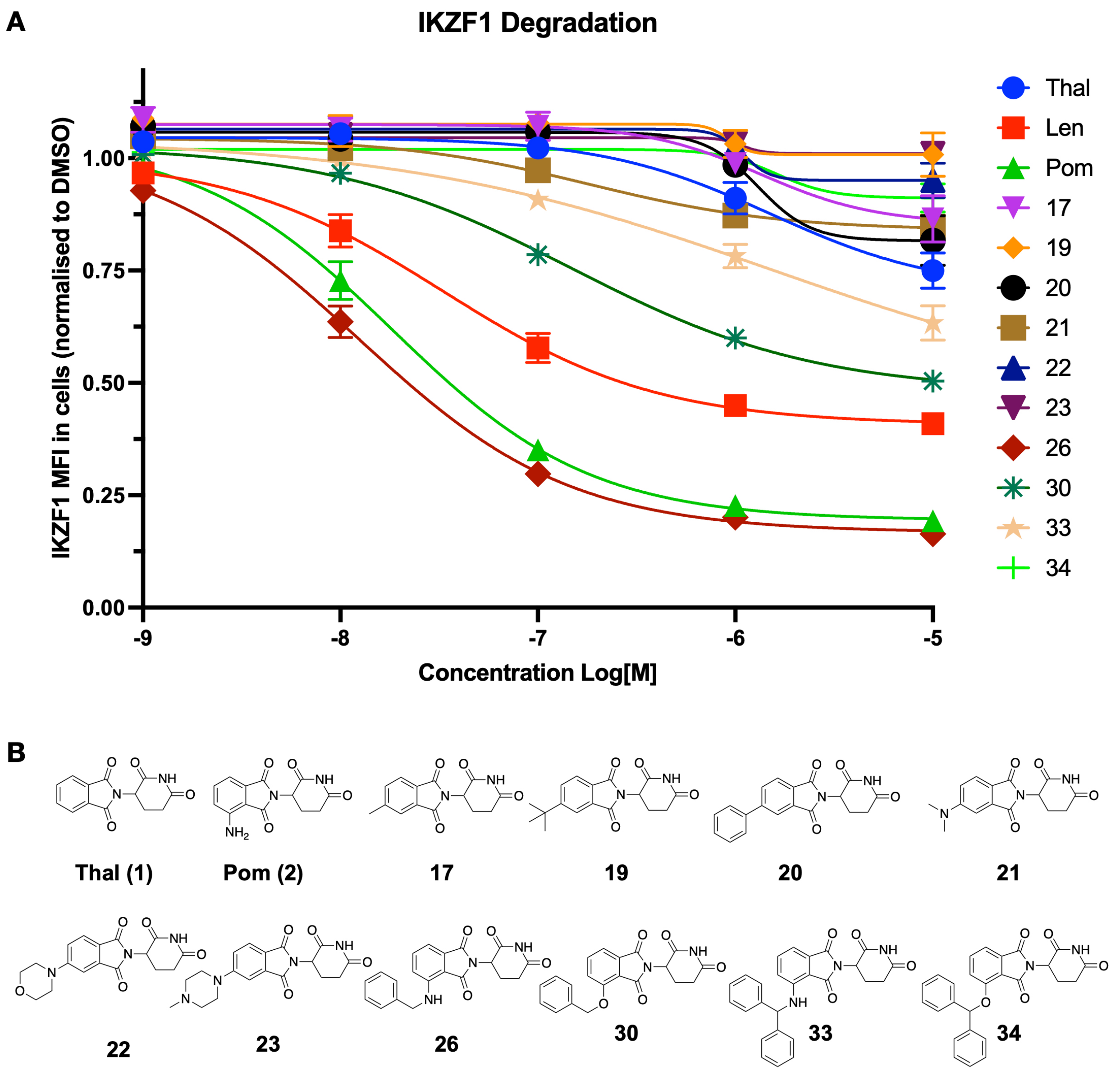

**Figure S5.**
**A.** Degradation curves for a subset of IMiD analog library compounds (**1**, **3**, **17**, **19**, **20**, **21**, **22**, **23**, **26**, **30**, **33**, **34** and commercially acquired lenalidomide) against endogenous IKZF1 determined using a fluorescent antibody flow cytometry assay. **B.** Chemical structures for compounds **1**, **3**, **17**, **19**, **20**, **21**, **22**, **23**, **26**, **30**, **33**, and **34**.

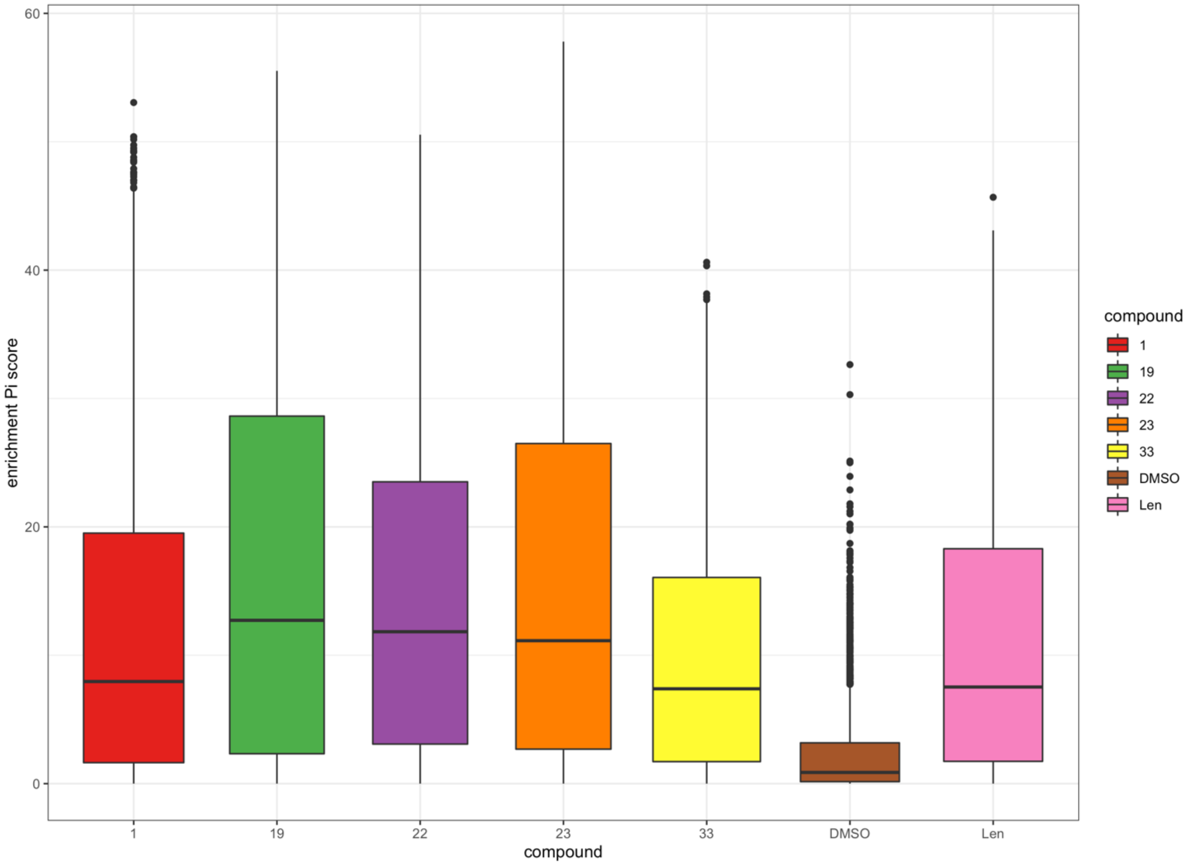

**Figure S6.**
Enrichment Pi score box plots for library screens for compounds **1** (thalidomide), **19**, **22**, **23**, **33** and lenalidomide, and a DMSO control. Pi score is calculated by $\log_2 FC \times \log10(p\text{-value})$, where *p*-value represents confidence in the result and $\log_2 FC$ represents fold-enrichment; pi score enables the identification of statistically significant fold-changes.

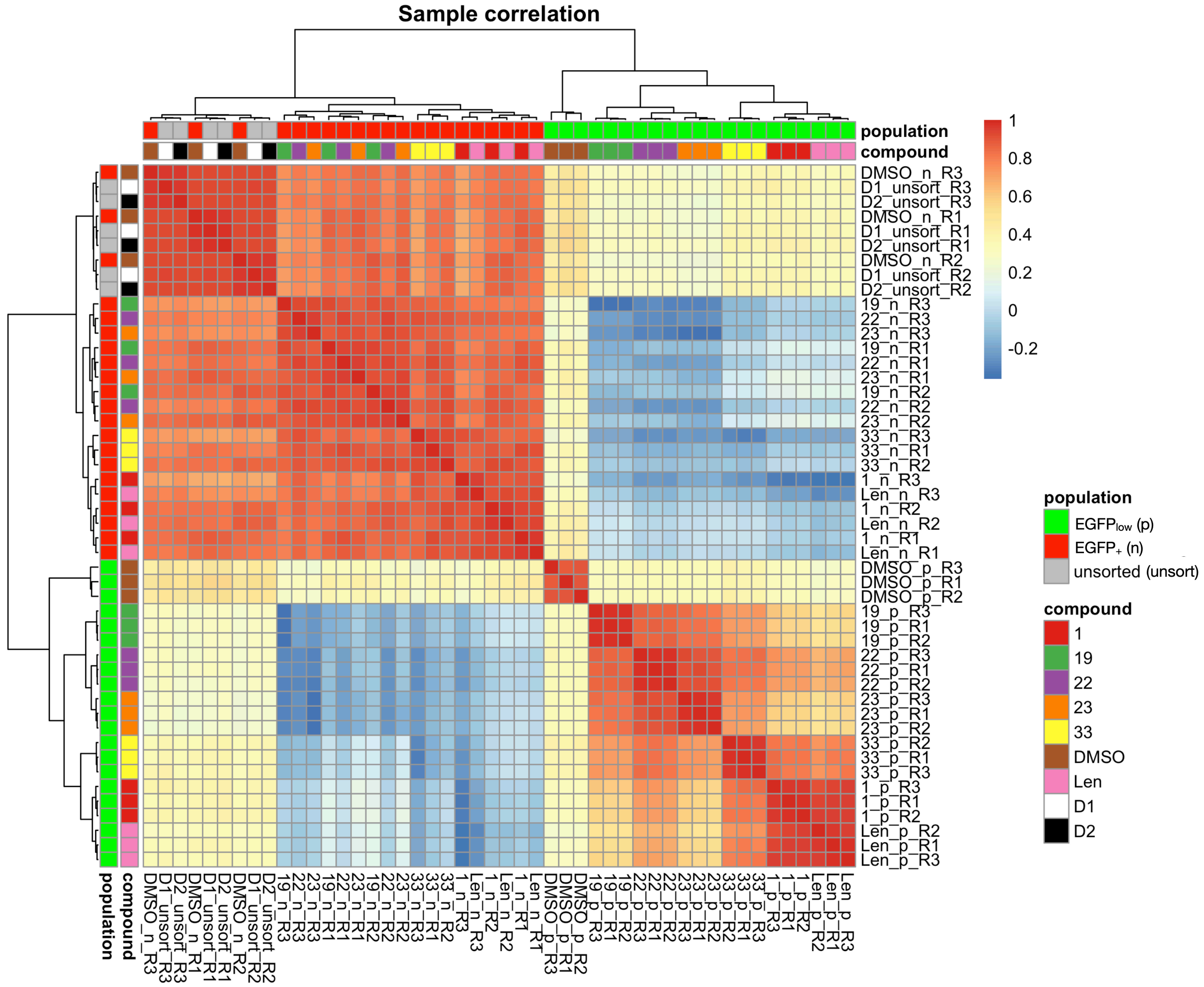

**Figure S7.**

A representation of the similarity of enriched sequences observed in populations of library ZF-EGFP-expressing Jurkat cell populations. Data are shown for three replicates of each condition. A value of 1 represents identical sequences; lower values represent increasing dissimilarity. The plot shows three replicates (R1, R2 and R3) for each condition: sorted EGFP$_{low}$ populations (p) for compound screens (**1** (thalidomide), **19**, **22**, **23**, **33** and lenalidomide) and an untreated control (DMSO); sorted EGFP$_{+}$ populations (n) for the same screens; and two populations of unsorted library cells (D1 and D2; unsort). The blue to red color scale represents sample correlation, with 1 (red) meaning complete sequence correlation.

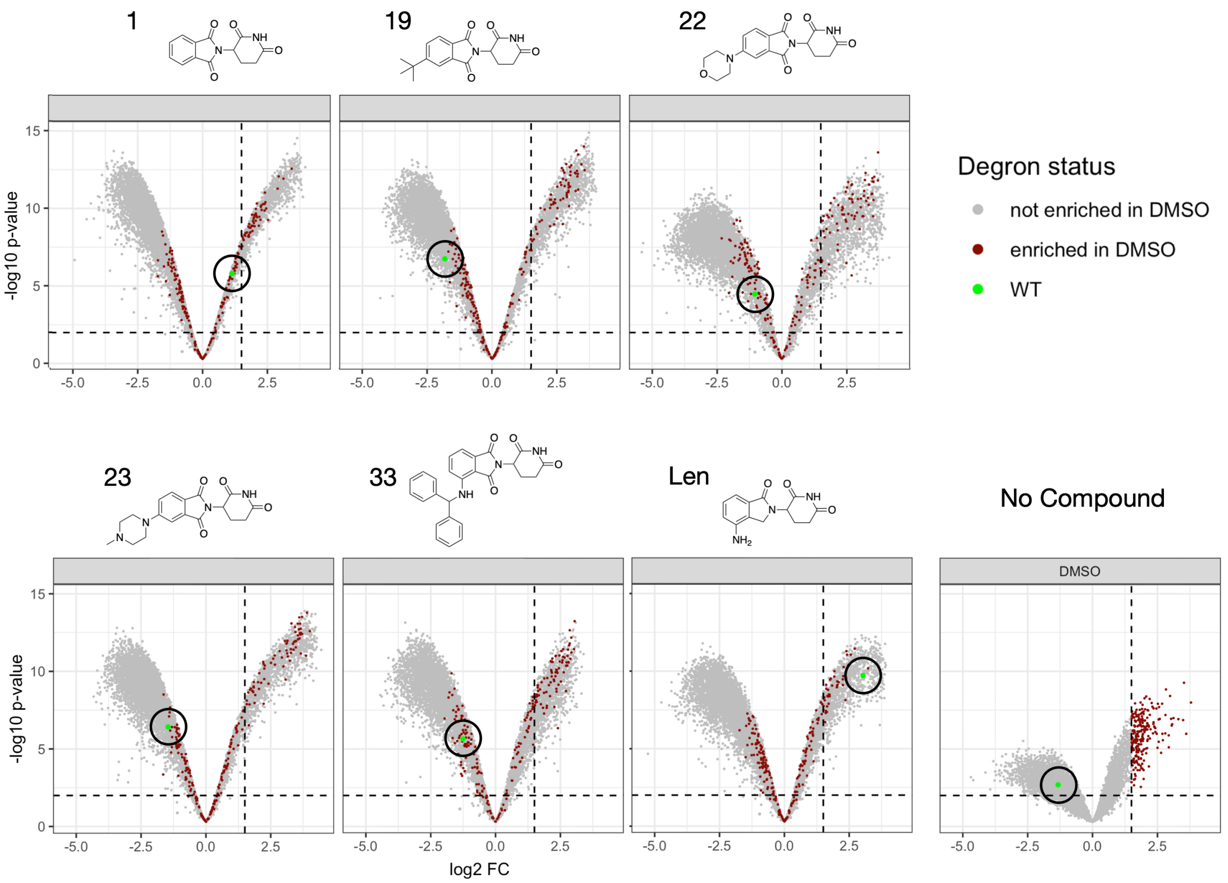

**Figure S8.**
Volcano plots for thalidomide (**1**), lenalidomide, **19**, **22**, **23** and **33** tested against the mutant library, and a DMSO control. Brown data points represent sequences that were enriched in the DMSO control; the green data point in each plot represents the WT sequence (i.e. no change in sequence from the original IKZF1/3 ZF degron).

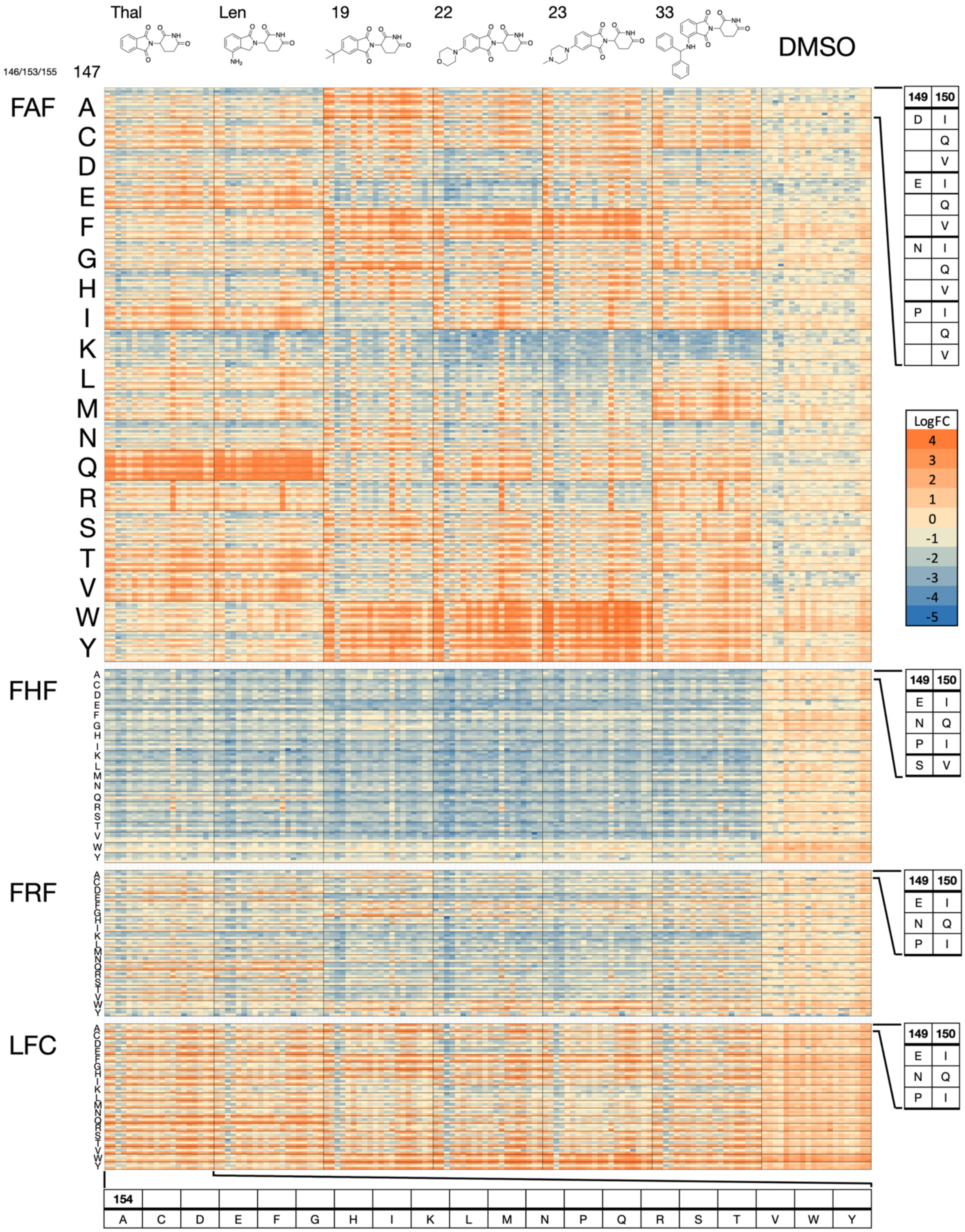

**Figure S9.**
$log_2FC$ data for six compounds and a DMSO control against 8360 mutant ZF degrons. $log_2FC$ values are arranged according to compound and residue 154 on the x-axis, and values are arranged according to residues 146/153/155, 147, 149, and 150 on the y-axis. Color scale shows high $log_2FC$ as orange, representing high sequence enrichment in the $EGFP_{low}$ population, and low $log_2FC$ as

blue, representing low sequence occurrence. A $log_2$FC score of zero, colored as tan, represents equal representation of a sequence in the ‘degraded’ cell population and the remaining cell population after FACS.

| Compound | Thalidomide | | | | | | | | | |
|---|---|---|---|---|---|---|---|---|---|---|
| Rank | 146 | 147 | 149 | 150 | 152 | 153 | 154 | 155 | 167 | Log2FC |
| 1 | F | Q | P | V | G | A | K | F | L | 3.9618 |
| 2 | F | Q | P | V | G | A | P | F | L | 3.8717 |
| 3 | F | Q | N | I | G | A | T | F | L | 3.8667 |
| 4 | F | Q | N | I | G | A | I | F | L | 3.8114 |
| 5 | F | Q | P | I | G | A | D | F | L | 3.8059 |
| 6 | F | Q | P | V | G | A | T | F | L | 3.7941 |
| 7 | F | Q | P | I | G | A | I | F | L | 3.7924 |
| 8 | F | Q | P | I | G | A | S | F | L | 3.7900 |
| 9 | F | Q | E | V | G | A | Q | F | L | 3.7719 |
| 10 | F | Q | P | I | G | A | C | F | L | 3.7711 |
| 11 | F | Q | P | V | G | A | E | F | L | 3.7612 |
| 12 | L | Q | P | I | G | F | V | C | L | 3.7517 |
| 13 | F | Q | E | I | G | A | S | F | L | 3.7484 |
| 14 | F | Q | P | V | G | A | V | F | L | 3.7460 |
| 15 | F | Q | E | V | G | A | T | F | L | 3.7446 |
| 16 | F | Q | E | I | G | A | P | F | L | 3.7224 |
| 17 | F | Q | E | V | G | A | V | F | L | 3.7106 |
| 18 | F | Q | N | I | G | A | P | F | L | 3.7088 |
| 19 | F | Q | E | V | G | A | R | F | L | 3.7061 |
| 20 | F | Q | E | I | G | A | R | F | L | 3.7041 |
| WT | F | Q | N | Q | G | A | S | F | L | 1.1337 |

| Compound | Lenalidomide | | | | | | | | | |
|---|---|---|---|---|---|---|---|---|---|---|
| Rank | 146 | 147 | 149 | 150 | 152 | 153 | 154 | 155 | 167 | Log2FC |
| 1 | F | Q | N | I | G | A | E | F | L | 3.9142 |
| 2 | F | Q | P | V | G | A | D | F | L | 3.8922 |
| 3 | F | Q | N | V | G | A | V | F | L | 3.8861 |
| 4 | F | Q | P | V | G | A | T | F | L | 3.8852 |
| 5 | F | Q | N | I | G | A | Q | F | L | 3.8639 |
| 6 | F | Q | P | V | G | A | N | F | L | 3.8156 |
| 7 | F | Q | P | I | G | A | L | F | L | 3.8345 |
| 8 | F | Q | N | I | G | A | T | F | L | 3.8290 |
| 9 | F | Q | N | I | G | A | A | F | L | 3.8170 |
| 10 | F | Q | E | V | G | A | A | F | L | 3.7915 |
| 11 | F | Q | N | I | G | A | I | F | L | 3.7833 |
| 12 | F | Q | E | I | G | A | R | F | L | 3.7803 |
| 13 | F | Q | E | Q | G | A | C | F | L | 3.7802 |
| 14 | F | Q | P | I | G | A | S | F | L | 3.7741 |
| 15 | F | Q | P | V | G | A | V | F | L | 3.7703 |
| 16 | F | Q | E | V | G | A | I | F | L | 3.7619 |
| 17 | F | Q | P | Q | G | A | V | F | L | 3.7592 |
| 18 | F | Q | P | Q | G | A | C | F | L | 3.7152 |
| 19 | F | Q | E | Q | G | A | V | F | L | 3.7519 |
| 20 | F | Q | P | Q | G | A | A | F | L | 3.7513 |
| WT | F | Q | N | Q | G | A | S | F | L | 3.0311 |

| Compound | Compound 19 | | | | | | | | | |
|---|---|---|---|---|---|---|---|---|---|---|
| Rank | 146 | 147 | 149 | 150 | 152 | 153 | 154 | 155 | 167 | Log2FC |
| 1 | F | H | E | I | G | A | P | F | L | 4.0013 |
| 2 | F | F | E | V | G | A | P | F | L | 3.9979 |
| 3 | F | W | E | I | G | A | K | F | L | 3.9155 |
| 4 | F | F | D | I | G | A | P | F | L | 3.9195 |
| 5 | F | A | E | V | G | A | K | F | L | 3.9176 |
| 6 | F | F | P | I | G | A | P | F | L | 3.9164 |
| 7 | F | A | E | V | G | A | R | F | L | 3.9084 |
| 8 | F | Y | E | I | G | A | K | F | L | 3.9035 |
| 9 | F | G | P | I | G | A | C | F | L | 3.9002 |
| 10 | F | A | E | I | G | A | A | F | L | 3.8965 |
| 11 | F | G | P | V | G | A | C | F | L | 3.8841 |
| 12 | F | Y | E | V | G | A | A | F | L | 3.8726 |
| 13 | F | A | E | V | G | A | S | F | L | 3.8698 |
| 14 | F | Y | N | I | G | A | P | F | L | 3.8168 |
| 15 | F | Y | P | I | G | A | P | F | L | 3.8626 |
| 16 | F | Y | E | I | G | A | P | F | L | 3.8608 |
| 17 | F | G | E | V | G | A | R | F | L | 3.8587 |
| 18 | F | G | P | V | G | A | M | F | L | 3.8585 |
| 19 | F | W | E | I | G | A | C | F | L | 3.8152 |
| 20 | F | A | E | I | G | A | R | F | L | 3.8372 |
| WT | F | Q | N | Q | G | A | S | F | L | -1.8291 |

| Compound | Compound 22 | | | | | | | | | |
|---|---|---|---|---|---|---|---|---|---|---|
| Rank | 146 | 147 | 149 | 150 | 152 | 153 | 154 | 155 | 167 | Log2FC |
| 1 | F | Y | E | V | G | A | T | F | L | 4.1392 |
| 2 | F | Y | E | I | G | A | T | F | L | 4.0154 |
| 3 | F | R | E | I | G | A | P | F | L | 3.9835 |
| 4 | F | I | P | V | G | A | P | F | L | 3.9370 |
| 5 | F | W | E | I | G | A | Q | F | L | 3.9298 |
| 6 | F | W | P | V | G | A | Q | F | L | 3.9190 |
| 7 | F | W | E | V | G | A | H | F | L | 3.9186 |
| 8 | F | W | E | V | G | A | A | F | L | 3.9041 |
| 9 | L | Y | E | I | G | F | R | C | L | 3.9018 |
| 10 | F | W | D | V | G | A | N | F | L | 3.8903 |
| 11 | F | Y | E | V | G | A | N | F | L | 3.8762 |
| 12 | F | Y | E | V | G | A | S | F | L | 3.8615 |
| 13 | F | W | E | V | G | A | K | F | L | 3.8571 |
| 14 | F | W | E | V | G | A | N | F | L | 3.8564 |
| 15 | F | W | P | V | G | A | N | F | L | 3.8155 |
| 16 | F | W | D | V | G | A | Q | F | L | 3.8513 |
| 17 | F | Y | E | V | G | A | C | F | L | 3.8457 |
| 18 | F | Y | E | I | G | A | K | F | L | 3.8391 |
| 19 | F | F | E | I | G | A | C | F | L | 3.8315 |
| 20 | F | W | E | I | G | A | A | F | L | 3.8306 |
| WT | F | Q | N | Q | G | A | S | F | L | -1.0370 |

| Compound | Compound 23 | | | | | | | | | |
|---|---|---|---|---|---|---|---|---|---|---|
| Rank | 146 | 147 | 149 | 150 | 152 | 153 | 154 | 155 | 167 | Log2FC |
| 1 | F | W | E | I | G | A | S | F | L | 4.4149 |
| 2 | F | W | D | I | G | A | N | F | L | 4.2762 |
| 3 | F | W | N | I | G | A | Q | F | L | 4.2715 |
| 4 | F | W | D | I | G | A | A | F | L | 4.2601 |
| 5 | F | W | E | V | G | A | S | F | L | 4.2567 |
| 6 | F | W | D | V | G | A | Q | F | L | 4.2369 |
| 7 | F | Y | E | V | G | A | S | F | L | 4.2356 |
| 8 | F | W | D | I | G | A | S | F | L | 4.2199 |
| 9 | F | W | D | V | G | A | S | F | L | 4.2017 |
| 10 | F | W | P | V | G | A | Q | F | L | 4.1974 |
| 11 | F | W | E | Q | G | A | S | F | L | 4.1915 |
| 12 | F | W | E | I | G | A | H | F | L | 4.1861 |
| 13 | F | W | E | I | G | A | C | F | L | 4.1830 |
| 14 | F | Y | E | I | G | A | T | F | L | 4.1715 |
| 15 | F | W | D | V | G | A | A | F | L | 4.1742 |
| 16 | F | W | E | I | G | A | K | F | L | 4.1692 |
| 17 | F | Y | E | V | G | A | E | F | L | 4.1417 |
| 18 | F | Y | E | I | G | A | S | F | L | 4.1371 |
| 19 | F | W | N | V | G | A | S | F | L | 4.1357 |
| 20 | F | W | E | V | G | A | Q | F | L | 4.1344 |
| WT | F | Q | N | Q | G | A | S | F | L | -1.4561 |

| Compound | Compound 33 | | | | | | | | | |
|---|---|---|---|---|---|---|---|---|---|---|
| Rank | 146 | 147 | 149 | 150 | 152 | 153 | 154 | 155 | 167 | Log2FC |
| 1 | F | R | E | V | G | A | P | F | L | 3.3928 |
| 2 | F | R | D | I | G | A | P | F | L | 3.3226 |
| 3 | F | R | E | I | G | A | P | F | L | 3.3123 |
| 4 | F | R | P | V | G | A | P | F | L | 3.2684 |
| 5 | F | M | N | I | G | A | P | F | L | 3.2358 |
| 6 | F | R | D | V | G | A | P | F | L | 3.2095 |
| 7 | F | M | E | I | G | A | A | F | L | 3.1949 |
| 8 | L | Y | E | I | G | F | T | C | L | 3.1905 |
| 9 | L | C | P | I | G | F | Y | C | L | 3.1590 |
| 10 | L | C | P | I | G | F | T | C | L | 3.1410 |
| 11 | F | Y | E | I | G | A | Q | F | L | 3.1346 |
| 12 | F | M | E | I | G | A | P | F | L | 3.1328 |
| 13 | F | M | E | V | G | A | P | F | L | 3.1157 |
| 14 | F | G | P | V | G | A | F | F | L | 3.1139 |
| 15 | F | Y | E | V | G | A | C | F | L | 3.1049 |
| 16 | L | Y | E | I | G | F | S | C | L | 3.0997 |
| 17 | L | Y | P | I | G | F | S | C | L | 3.0902 |
| 18 | F | M | P | I | G | A | P | F | L | 3.0896 |
| 19 | F | M | P | V | G | A | P | F | L | 3.0869 |
| 20 | F | M | P | V | G | A | A | F | L | 3.0783 |
| WT | F | Q | N | Q | G | A | S | F | L | -1.2556 |

| Compound | DMSO Control | | | | | | | | | |
|---|---|---|---|---|---|---|---|---|---|---|
| Rank | 146 | 147 | 149 | 150 | 152 | 153 | 154 | 155 | 167 | Log2FC |
| 1 | L | W | P | I | G | F | F | C | L | 3.7923 |
| 2 | L | W | P | I | G | F | L | C | L | 3.6101 |
| 3 | L | W | P | I | G | F | P | C | L | 3.5229 |
| 4 | L | W | P | I | G | F | Y | C | L | 3.4893 |
| 5 | L | W | P | I | G | F | I | C | L | 3.3061 |
| 6 | L | W | P | I | G | F | D | C | L | 3.1185 |
| 7 | L | W | P | I | G | F | M | C | L | 3.0653 |
| 8 | L | W | P | I | G | F | W | C | L | 3.0491 |
| 9 | L | W | E | I | G | F | F | C | L | 3.0278 |
| 10 | L | W | P | I | G | F | C | C | L | 2.9859 |
| 11 | L | W | N | Q | G | F | I | C | L | 2.9385 |
| 12 | L | W | N | Q | G | F | F | C | L | 2.9343 |
| 13 | L | W | E | I | G | F | I | C | L | 2.8293 |
| 14 | L | W | P | I | G | F | P | C | L | 2.8090 |
| 15 | L | W | N | Q | G | F | W | C | L | 2.8031 |
| 16 | F | T | P | I | G | H | Q | F | L | 2.7099 |
| 17 | F | W | P | I | G | H | G | F | L | 2.6906 |
| 18 | L | W | N | Q | G | F | L | C | L | 2.6802 |
| 19 | F | W | P | I | G | A | F | F | L | 2.6417 |
| 20 | L | W | P | I | G | F | V | C | L | 2.5893 |
| WT | F | Q | N | Q | G | A | S | F | L | -1.325 |

| |
|---|
| A |
| C |
| D |
| E |
| F |
| G |
| H |
| I |
| K |
| L |
| M |
| N |
| P |
| Q |
| R |
| S |
| T |
| V |
| W |
| Y |

**Figure S10.**

Table showing the 20 highest ranked mutant ZF sequences by $log_2FC$ for compounds **1** (thalidomide), **19**, **22**, **23**, **33**, lenalidomide and a DMSO control; the WT (IKZF1/3 ZF2) sequence and $log_2FC$ for each of the 3 compound screens are also shown for comparison.

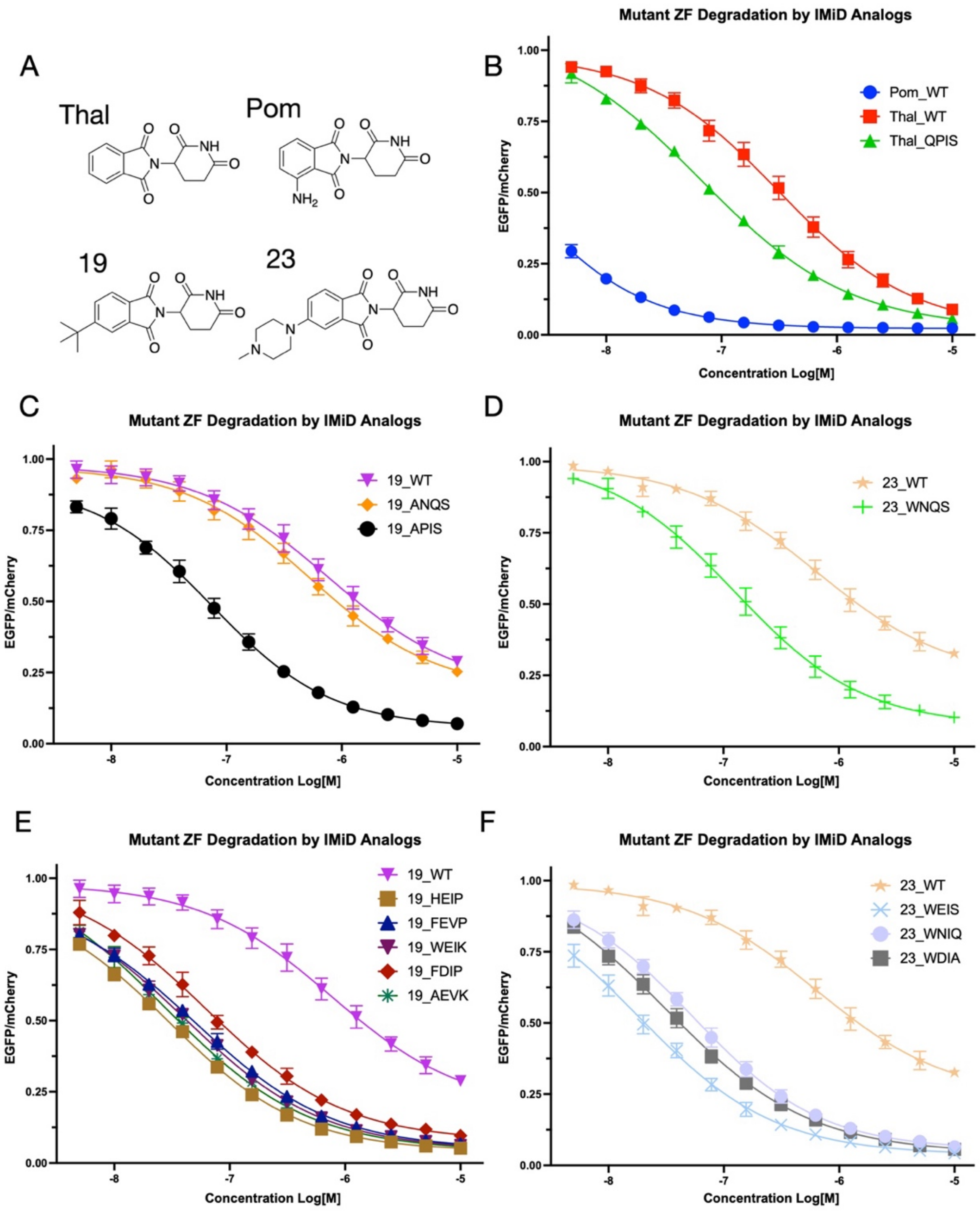

**Figure S11.**
**A.** Chemical structure of four IMiD analogs. Dose-response curves for: **B.** pomalidomide and thalidomide degradation of the WT degron; thalidomide degradation of the QPIS degron. **C.** Compound **19** degradation of the WT, ANQS and APIS degrons. **D.** Compound **23** degradation of the WT and WNQS degrons. **E.** Compound **19** degradation of the WT degron and five top ranking mutant degrons from the compound **19** mutant library screen. **F.** Compound **23** degradation of the WT degron and three (of five) top ranking mutant degrons from the compound **23** mutant library screen.

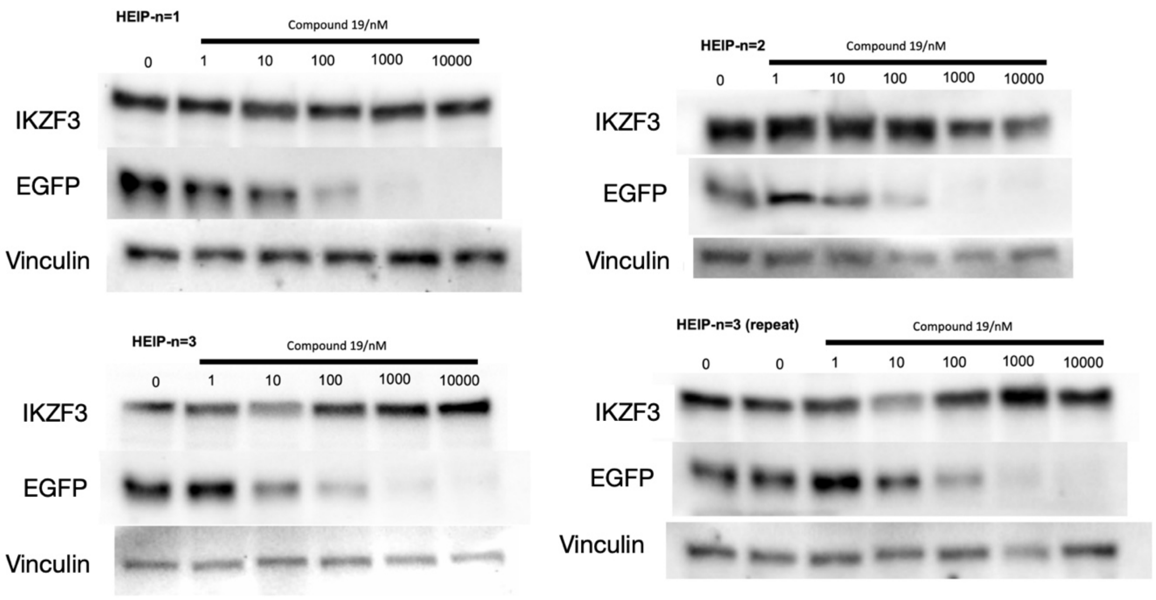

**Figure S12.**
Western blot performed on transduced Jurkat cells expressing a ZF-EGFP fusion protein with the HEIP degron at five concentrations of compound **19** and an untreated DMSO control. Bands show levels of IKZF3, EGFP, or vinculin.

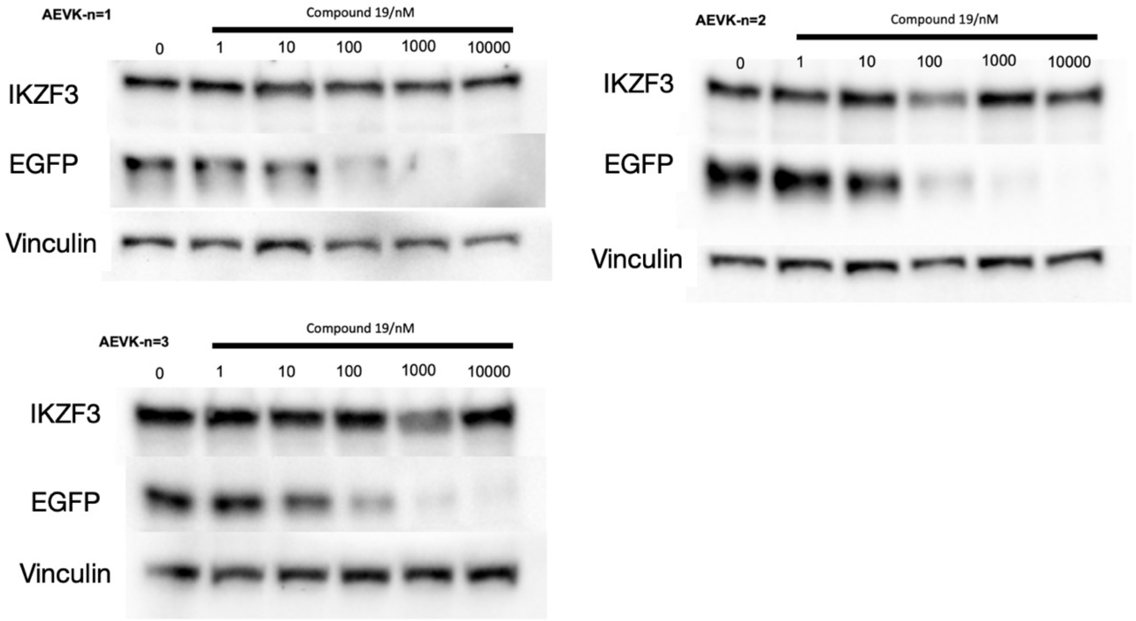

**Figure S13.**
Western blot performed on transduced Jurkat cells expressing a ZF-EGFP fusion protein with the AEVK degron at five concentrations of compound **19** and an untreated DMSO control. Bands show levels of IKZF3, EGFP, or vinculin.

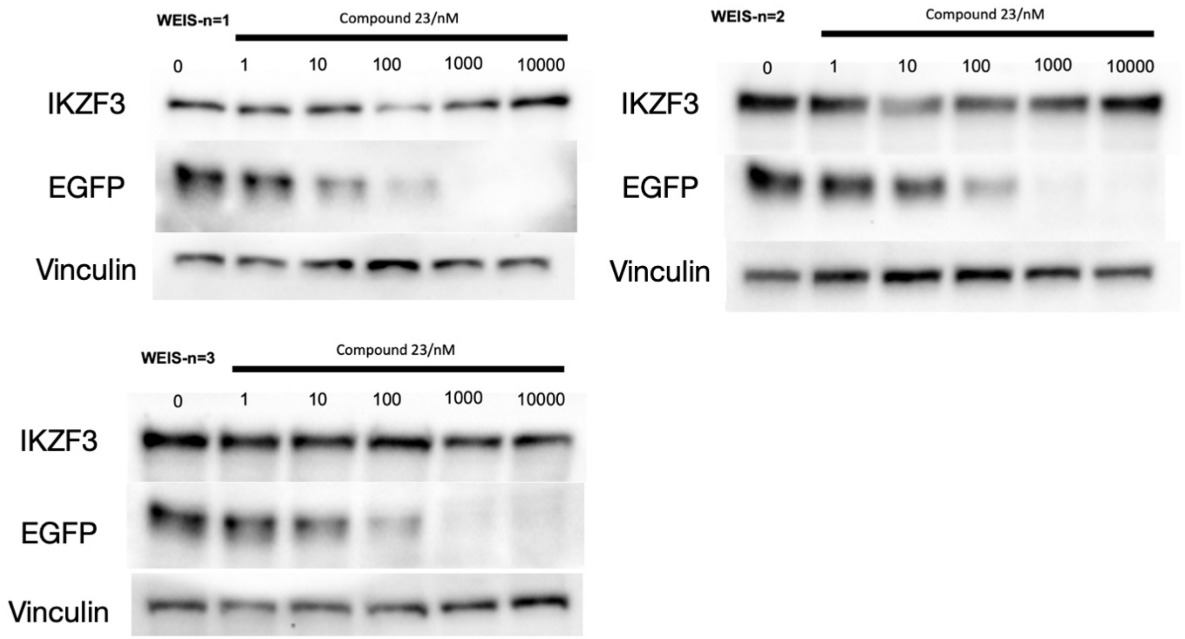

**Figure S14.**
Western blot performed on transduced Jurkat cells expressing a ZF-EGFP fusion protein with the WEIS degron at five concentrations of compound **23** and an untreated DMSO control. Bands show levels of IKZF3, EGFP, or vinculin.

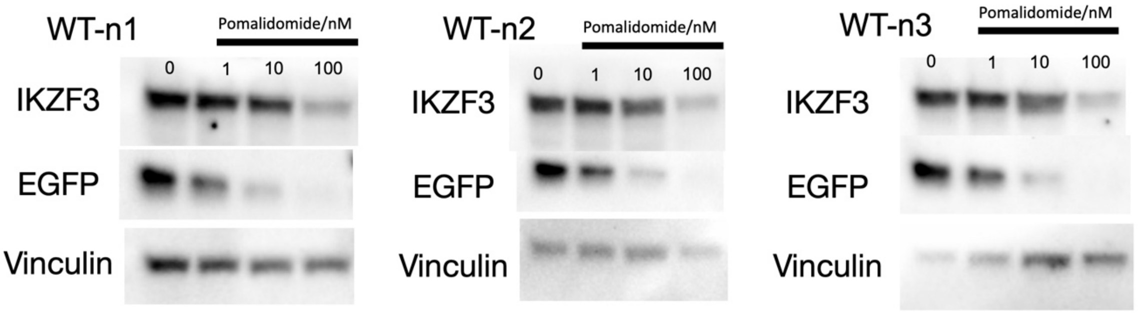

**Figure S15.**
Western blot performed on transduced Jurkat cells expressing a ZF-EGFP fusion protein with the WT degron at three concentrations of compound **2** (pomalidomide) and an untreated DMSO control. Bands show levels of IKZF3, EGFP, or vinculin.

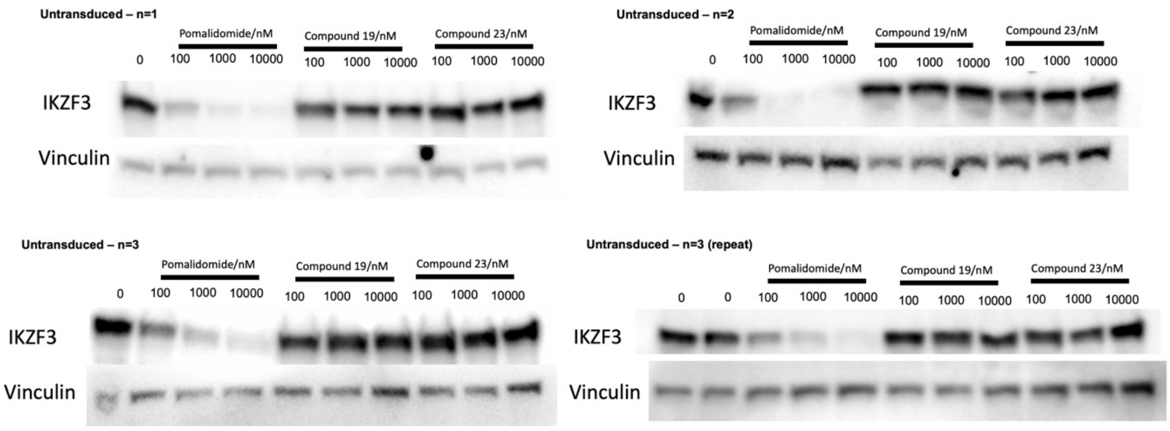

**Figure S16.**
Western blot performed on untransduced Jurkat cells at three concentrations of compound **2** (pomalidomide), compound **19**, or compound **23**, and an untreated DMSO control. Bands show levels of IKZF3 or vinculin.

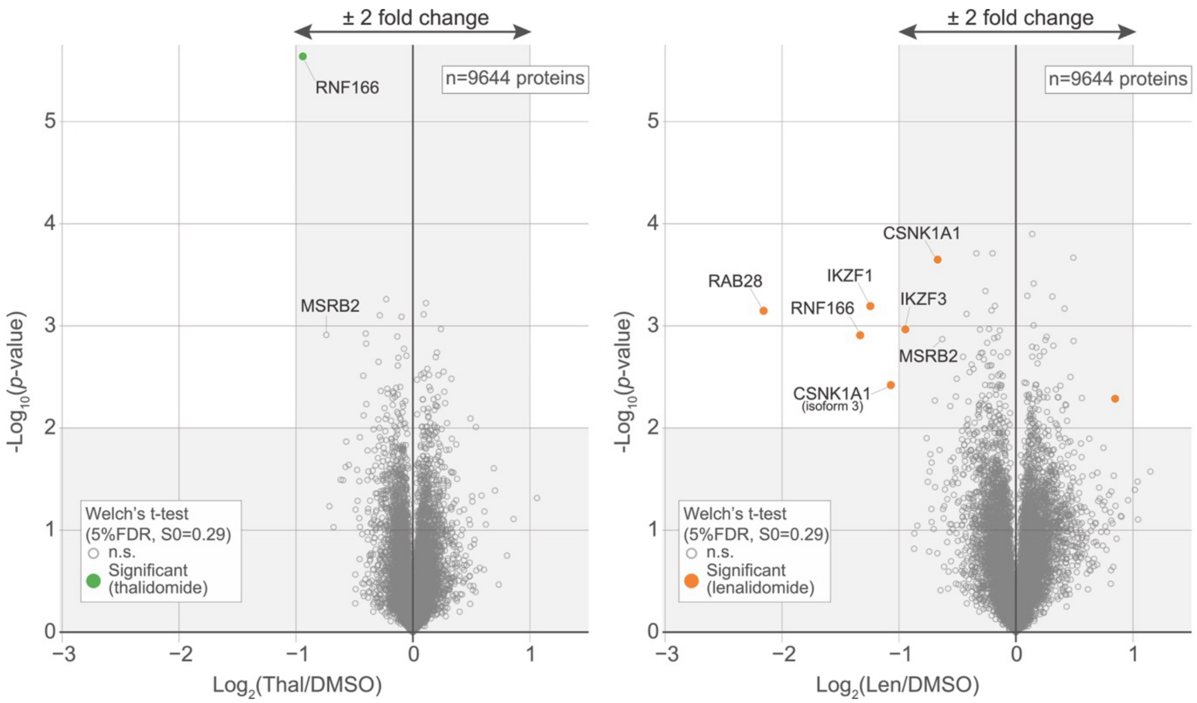

**Figure S17.**
Quantitative proteomics profiling following cell treatment with thalidomide (10 μM) or lenalidomide (1 μM). Jurkats cells were treated for 16 h with compound or DMSO control, and protein abundance was analysed using TMT quantification mass spectrometry. $\log_2$FC is shown on the x-axis, and $-\log_{10}$($p$-value) is shown on the y-axis. Values shown are an average of three biological replicates.

**Figure S18 (separate file).**
$log_2FC$ data for six compounds and a DMSO control against 8380 library mutant ZF degrons. $log_2FC$ values are arranged according to residue 154 on the x-axis, and values are arranged according to residues 146/153/155, 147, 149 and 150 on the y-axis. Color scale shows high $log_2FC$ as orange, representing high sequence enrichment in the EGFP-low population, and low $log_2FC$ as blue, representing low sequence occurrence. A $log_2FC$ score of zero, colored as tan, represents equal representation of a sequence in the 'degraded' cell population and the remaining cell population after FACS.

#### Materials and Methods

##### Chemistry Experimental Section

Reagents and solvents used were of commercially available reagent grade quality from Alfa Aesar, Fluorochem, Merck (formally known as Sigma-Aldrich), or Tokyo Chemical Industry, and were used without further purification unless stated. Anhydrous solvents were obtained from an MBRAUN Solvent Purification Systems 5 and stored under an argon atmosphere over 3 Å molecular sieves. Concentration *in vacuo* was performed at 40 °C for organic solvents using a Buchi™ rotary evaporator. Brine refers to a saturated aqueous solution of sodium chloride. Petroleum ether refers to the fraction of light petroleum ether boiling in the range 40-60 °C. Celite® refers to Celite® 545 filter aid, treated with sodium carbonate, flux-calcined (Merck).

$^{1}$H NMR spectra were measured on a Bruker AV400 (400 MHz), a Bruker AVII 500 (500 MHz) or a Bruker AV600 (600 MHz) spectrometer in the stated solvents as a reference for the internal deuterium lock. The chemical shift data for each signal are given as δ in units of parts per million (ppm) relative to tetramethylsilane (TMS) where δ(TMS) = 0.00. The spectra are calibrated using the solvent peak with the data provided by Fulmer *et al.* (*58*). The multiplicity of each signal is indicated by: s (singlet); bs (broad singlet); d (doublet); t (triplet); q (quartet); p (pentet); sept (septet); m (multiplet) or combinations thereof. The number of protons (n) for a given resonance signal is indicated by nH. Where appropriate, coupling constants (*J*) are quoted in Hz, recorded to the nearest 0.1 Hz, and were determined by analysis using Bruker TopSpin v3.2 software or MestreNova software. The mean value of identical coupling constants is reported. Spectra were assigned using COSY, HSQC and HMBC experiments as necessary. Spectra acquired at high temperature (353 K or 363 K) are indicated.

$^{13}$C NMR spectra were measured on a Bruker AV400 (101 MHz), a Bruker AVII 500 (126 MHz) or a Bruker AV600 (151 MHz) spectrometer in the stated solvents as a reference for the internal deuterium lock using the standard $^{13}$C experiment. The chemical shift data for each signal are given as δ in units of parts per million (ppm) relative to tetramethylsilane (TMS) where δ(TMS) = 0.00. The spectra are calibrated using the solvent peak with the data provided by Fulmer *et al.* (*58*). Signals are quoted to one decimal place unless peaks are indistinguishable, in which case two decimal places are used. Where appropriate, coupling constants (*J*) are quoted in Hz, recorded to the nearest 0.1 Hz, and were determined by analysis using Bruker TopSpin v3.2 software or MestreNova software. Spectra were assigned using HSQC and HMBC experiments as necessary. Spectra acquired at high temperature (353 K or 363 K) are indicated.

$^{19}$F NMR spectra were measured on a Bruker AVIII HD 400 (376 MHz) spectrometer in the solvent stated, and are $^{13}$C decoupled. Chemical shifts are given as δ in units of parts per million (ppm), to the nearest 0.1 ppm. The multiplicity of each signal is a singlet unless stated otherwise.

Mass spectra were acquired on either an Agilent 6120 (low resolution) or Bruker microToF spectrometer (high resolution) using electrospray ionisation (ESI) from solutions of either methanol or water. *m/z* values are reported in Daltons and are followed by their percentage abundance in parentheses.

Melting points were determined using a Griffin capillary tube melting point apparatus (Registered Design No. 889339) and are uncorrected. The solvent(s) from which the sample was crystallised is given in parentheses. Dec. indicates that the sample decomposed at the stated temperature.

Infrared spectra were obtained from thin films, using a diamond ATR module on a Bruker Tensor 27 spectrometer. Absorption maxima are reported in wavenumbers ($cm^{-1}$) and reported as s (strong), m (medium), w (weak) or br (broad).

Analytical high performance liquid chromatography (HPLC) was performed on a PerkinElmer Flexar system with a binary LC Pump and UV/vis LC detector set at 254 nm or Agilent 1260 Infinity II® system equipped with a Poroshell 120 EC-C18 column [4 µm, 4.6 × 100 mm] with a diode array UV/vis detector. To determine compound purity, a Dionex Acclaim® 120 C18 [5 µm, 12 Å, 150 mm × 4.6 mm] reverse phase column was used with a constant flow rate of 1.5 mL $min^{-1}$ and gradient method of 10 min from 95:5 $H_2O$:acetonitrile (0.1% TFA) to 5:95 $H_2O$:acetonitrile (0.1% TFA) with a 5 min hold. Samples injected were prepared by dissolving in methanol, water, or acetonitrile, and filtered. All samples are run with 0.1% trifluoroacetic acid or 0.1% formic acid added unless indicated.

All biologically assessed compounds had a purity of ≥95% determined using analytical HPLC.

Normal Phase silica gel analytical Thin Layer Chromatography (NP TLC) to monitor reaction progress was carried out on normal phase Merck silica gel 60 F254 aluminium-supported thin layer chromatography sheets. Visualisation was carried out using absorption of UV light ($\lambda_{max}$ = 254 nm and 365 nm) or thermal development after staining in an ethanolic solution of ninhydrin or an aqueous solution of potassium permanganate.

Normal phase silica gel flash column chromatography was carried out manually on Merck Geduran® silica gel 60 (40–63 µm), eluting with solvents as supplied, under a positive pressure of compressed, gaseous, nitrogen.

<u>Compound synthesis</u>

**2-(2,6-Dioxopiperidin-3-yl)isoindoline-1,3-dione (1)**

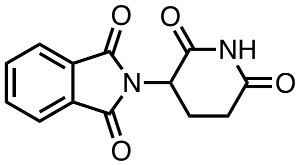

Phthalic anhydride (200 mg, 1.35 mmol, 1.0 eq), 3-aminopiperidine-2,6-dione hydrochloride (222 mg, 1.35 mmol, 1.0 eq), and NaOAc (166 mg, 2.03 mmol, 1.5 eq) were dissolved in AcOH (5 mL) and stirred at 120 °C for 16.5 h. After this time the reaction solution was cooled to rt, and the solvent removed *in vacuo* (azeotrope with cyclohexane). The product was purified using flash column chromatography (2/98–10/90 MeOH/$CH_2Cl_2$) to afford the title compound (270 mg, 77%) as a colorless solid: $R_f$ 0.60 (10/90 MeOH/$CH_2Cl_2$); m.p. 263–266 °C (from $CH_2Cl_2$) [lit. (*59*) 269–271 °C, lit. (*60*) 258–260 °C, lit. (*61*) 270–272 °C]; $^1H$ NMR (400 MHz, $D_6$-DMSO) δ 11.12 (1H, s), 7.98 – 7.85 (4H, m), 5.16 (1H, dd, *J* 12.9, 5.4 Hz), 2.96 – 2.84 (1H, m), 2.66 – 2.52 (2H, m), 2.13 – 1.97 (1H, m); LRMS *m/z* ($ESI^-$) 257 ($[M-H]^-$, 100%); HPLC Retention time 220 nm: 7.0 min, 98.7%; 254 nm: 7.0 min, 100.0%. These data are in good agreement with the literature values (*59*).

**4-Amino-2-(2,6-dioxopiperidin-3-yl)isoindoline-1,3-dione (2)**

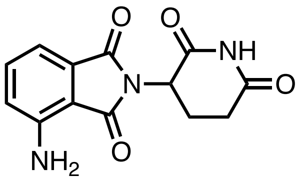

2-(2,6-Dioxopiperidin-3-yl)-4-nitroisoindoline-1,3-dione (497 mg, 1.64 mmol, 3.0) and palladium on carbon (10% *w/w*; 50 mg, 0.47 mmol, 1.0 eq) were dissolved in DMF (20 mL) and heated to 35 °C under an atmosphere of $H_2$ for 16 h. After this time the mixture was cooled to rt and filtered through Celite®. The DMF was removed from the filtrate *in vacuo* (azeotrope with toluene). The residual light green powder was dissolved in 20 mL of EtOAc and heated under reflux at 77 °C for 30 min. After this time, the solution was cooled, filtered, and the solvent was removed *in vacuo* to afford the title compound (363 mg, 81%) as a yellow, fluorescent powder: $R_f$ 0.22 (50/50 EtOAc/petroleum ether); m.p. 290 °C – Dec. (from $CH_2Cl_2$) [lit. (*62*) 319–322 °C, lit. (*63*) 252 °C]; $^1$H NMR (400 MHz, $D_6$-DMSO) δ 11.07 (1H, s), 7.47 (1H, dd, *J* 8.5, 7.0 Hz), 7.05 – 6.97 (2H, m), 6.51 (2H, s), 5.04 (1H, dd, *J* 12.9, 5.4 Hz), 2.94 – 2.82 (1H, m), 2.64 – 2.51 (2H, m), 2.07 – 1.96 (1H, m); LRMS *m/z* ($ESI^-$) 272 ($[M-H]^-$, 100%); HPLC Retention time 220 nm: 6.4 min, 99.9%; 254 nm: 6.4 min, 100.0%. These data are in good agreement with the literature values (*64*).

**5-Amino-2-(2,6-dioxopiperidin-3-yl)isoindoline-1,3-dione (3)**

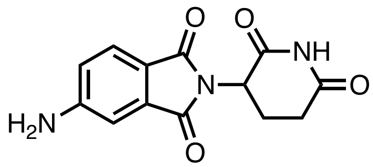

2-(2,6-Dioxopiperidin-3-yl)-5-nitroisoindoline-1,3-dione (691 mg, 2.28 mmol, 3.0 eq) and palladium on carbon (10% *w/w*; 70 mg, 0.74 mmol, 1.0 eq) were dissolved in DMF and heated to 35 °C under an atmosphere of argon. $H_2$ (g) was then bubbled through the solution for 10 min and the reaction was stirred at rt for 16 h under an atmosphere of $H_2$ (g). The mixture was then filtered through Celite® and the residue washed with toluene. The DMF was removed from the filtrate *in vacuo* (azeotrope with toluene). The residual dark green powder was dissolved in 25 mL of EtOAc and heated under reflux at 77 °C for 30 min. The solution was cooled, filtered, and the solvent was removed *in vacuo* to afford a yellow, fluorescent powder (369 mg, 52%): $R_f$ 0.40 (10% MeOH: $CH_2Cl_2$); m.p. >300 °C – Dec. (from $CH_2Cl_2$) [lit. (*65*) 318–320 °C, lit. (*66*) 320–322 °C]; $^1$H NMR (400 MHz, $D_6$-DMSO) δ 11.06 (1H, s), 7.52 (1H, d, *J* 8.3 Hz), 6.94 (1H, d, *J* 2.1 Hz), 6.83 (1H, dd, *J* 8.3, 2.1 Hz), 6.55 (2H, s), 5.01 (1H, dd, *J* 12.8, 5.5 Hz), 2.94 – 2.80 (1H, m), 2.62 – 2.52 (2H, m), 2.04 – 1.93 (1H, m); LRMS *m/z* ($ESI^-$) 272 ($[M-H]^-$, 100%); HPLC Retention time

220 nm: 6.2 min, 98.7%; 254 nm: 6.2 min, 100.0%. These data are in good agreement with the literature values (*67*).

**2-(2,6-Dioxopiperidin-3-yl)-4-hydroxyisoindoline-1,3-dione (4)**

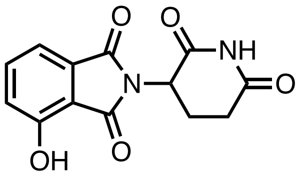

3-Hydroxyphthalic anhydride (2.00 g, 12.2 mmol, 1.0 eq), 3-aminopiperidine-2,6-dione hydrochloride (2.01 g, 12.2 mmol, 1.0 eq) and NaOAc (1.50 g, 18.3 mmol, 1.5 eq) were dissolved in AcOH (50 mL) and heated at 140 °C under reflux for 17.5 h. The AcOH was removed *in vacuo* (azeotrope with cyclohexane). The residue was dissolved in water and extracted with EtOAc, washed with water and brine, then dried with sodium sulfate, and filtered. The EtOAc was removed *in vacuo* to afford a salmon-pink solid (1.14 g, 34%). No further purification was required: $R_f$ 0.31 (5/95 MeOH/$CH_2Cl_2$); m.p. 257–259 °C – Dec. (from $CH_2Cl_2$) [lit. (*68*) 243–244 °C, lit. (*66*) 275–276 °C, lit. (*69*) 281–282 °C]; $^1$H NMR (400 MHz, $D_6$-DMSO) δ 11.19 (1H, s), 11.08 (1H, s), 7.65 (1H, dd, *J* 8.4, 7.2 Hz), 7.32 (1H, dd, *J* 7.2, 0.8 Hz), 7.25 (1H, dd, *J* 8.4, 0.8 Hz), 5.07 (1H, dd, *J* 12.8, 5.4 Hz), 2.95 – 2.81 (1H, m), 2.64 – 2.50 (2H, m), 2.08 – 1.96 (1H, m); LRMS *m/z* (ESI$^-$) 273 ([M−H]$^-$, 100%); HPLC Retention time 220 nm: 6.2 min, 97.0%; 254 nm: 6.2 min, 100%. These data are in good agreement with the literature values (*70*).

**2-(2,6-Dioxopiperidin-3-yl)-5-hydroxyisoindoline-1,3-dione (5)**

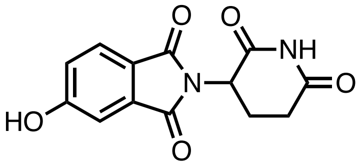

4-Hydroxyphthalic acid (246 mg, 1.35 mmol, 1.0 eq), 3-aminopiperidine-2,6-dione hydrochloride (222 mg, 1.35 mmol, 1.0 eq) and NaOAc (166 mg, 2.03 mmol, 1.5 eq) were dissolved in AcOH (5 mL) and heated at 120 °C under reflux for 3 h. The AcOH was removed *in vacuo* and the residue purified using flash column chromatography (5% MeOH/$CH_2Cl_2$) to afford a colorless solid (274 mg, 74%) which fluoresces yellow when dissolved: $R_f$ 0.43 (10% MeOH/$CH_2Cl_2$); m.p. >300 °C (from $H_2O$/MeCN) [lit. (*71*) 317–318 °C]; $^1$H NMR (400 MHz, $D_6$-DMSO) δ 11.14 (1H, br s), 11.11 (1H, s), 7.74 (1H, dd, *J* 7.9, 0.8 Hz), 7.24 – 7.08 (2H, m), 5.08 (1H, dd, *J* 12.9, 5.3 Hz), 2.94 – 2.82 (1H, m), 2.64 – 2.51 (2H, m), 2.11 – 1.97 (1H, m). LRMS *m/z* (ESI$^-$) 273 ([M−H]$^-$, 100%); HPLC Retention time 220 nm: 6.3 min, 95.3%; 254 nm: 6.4 min, 99.7%. These data are in good agreement with the literature values (*72*).

**2-(2,6-Dioxopiperidin-3-yl)-4-methoxyisoindoline-1,3-dione (6)**

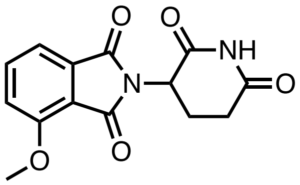

3-Methoxyphthalic acid (125 mg, 0.637 mmol, 1.0 eq), 3-aminopiperidine-2,6-dione hydrochloride (115 mg, 0.699 mmol, 1.0 eq) and NaOAc (86.0 mg, 1.05 mmol, 1.5 eq) were dissolved in AcOH (5 mL) and heated at 120 °C under reflux for 16.5 h. The AcOH was removed *in vacuo* and the residue purified using flash column chromatography (2/98 MeOH/$CH_2Cl_2$) to afford a colorless solid (96 mg, 47%): $R_f$ 0.50 (2/98 MeOH/$CH_2Cl_2$); m.p. 276–278 °C (from $CH_2Cl_2$) [lit.[17] 281–282 °C]; $^1$H NMR (400 MHz, $D_6$-DMSO) δ 11.10 (1H, s), 7.84 (1H, dd, *J* 8.6, 7.3 Hz), 7.53 (1H, dd, *J* 8.6, 0.7 Hz), 7.46 (1H, dd, *J* 7.3, 0.7 Hz), 5.09 (1H, dd, *J* 12.7, 5.4 Hz), 3.97 (3H, s), 2.94 – 2.83 (1H, m), 2.63 – 2.51 (2H, m), 2.11 – 1.97 (1H, m); LRMS *m/z* (ESI$^-$) 287 ([M−H]$^-$, 100%); HPLC Retention time 220 nm: 6.9 min, 96.7%; 254 nm: 6.9 min, 100.0%. These data are in good agreement with the literature value (*73*).

**2-(2,6-Dioxopiperidin-3-yl)-5-methoxyisoindoline-1,3-dione (7)**

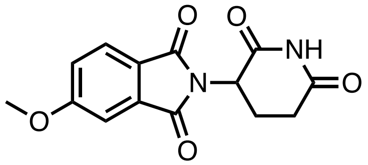

4-Methoxyphthalic acid (167 mg, 0.851 mmol, 1.0 eq), 3-aminopiperidine-2,6-dione hydrochloride (140 mg, 0.851 mmol, 1.0 eq) and NaOAc (105 mg, 1.28 mmol, 1.5 eq) were dissolved in AcOH (5 mL) and heated at 120 °C under reflux for 20 h. The AcOH was removed *in vacuo* and the residue purified using flash column chromatography (2/98 MeOH/$CH_2Cl_2$) to afford a colorless solid (122 mg, 50%): $R_f$ 0.61 (2% MeOH/$CH_2Cl_2$); m.p. 218–219 °C (from $CH_2Cl_2$) [lit. (*74*) 218–220 °C]; $\nu_{max}$ (thin film)/cm$^{-1}$ 1717 (C=O, s); $^1$H NMR (400 MHz, $D_6$-DMSO) δ 11.11 (1H, s), 7.85 (1H, d, *J* 8.3 Hz), 7.44 (1H, d, *J* 2.3 Hz), 7.36 (1H, dd, *J* 8.3, 2.3 Hz), 5.12 (1H, dd, *J* 12.9, 5.5 Hz), 3.94 (3H, s), 2.94 – 2.83 (1H, m), 2.65 – 2.51 (2H, m), 2.09 – 1.99 (1H, m); $^{13}$C NMR (151 MHz, $D_6$-DMSO) δ 172.7, 169.9, 166.9, 166.8, 164.7, 133.9, 125.3, 123.0, 120.3, 108.5, 56.4, 49.0, 30.9, 22.0; HRMS *m/z* (ESI$^-$) [Found: 287.0676, $C_{14}H_{11}N_2O_5$ requires [M−H$^-$] 287.0673)]; LRMS *m/z* (ESI$^-$) 287 ([M−H$^-$, 100%); HPLC Retention time 220 nm: 7.6 min, 95.5%; 254 nm: 7.6 min, 100.0%.

**2-(2,6-Dioxopiperidin-3-yl)-4-fluoroisoindoline-1,3-dione (8)**

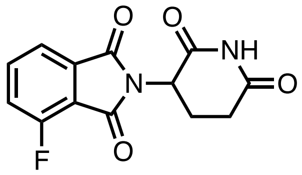

3-Fluorophthalic anhydride (2.00 g, 12.0 mmol, 1.0 eq), 3-aminopiperidine-2,6-dione hydrochloride (1.98 g, 12.0 mmol, 1.0 eq) and NaOAc (1.27 g, 18.1 mmol, 1.5 eq) were dissolved in AcOH (50 mL) and heated at 140 °C under reflux for 17 h. The mixture was cooled to rt and the AcOH removed *in vacuo* (azeotrope with cyclohexane). The residue was extracted from an aqueous solution of LiCl (0.5 M) with EtOAc. The EtOAc was removed *in vacuo* and the residue purified using flash column chromatography (60/40 EtOAc/petroleum ether) to afford the title compound (3.18 g, 67%) as a colorless solid: $R_f$ 0.50 (10% MeOH/$CH_2Cl_2$); m.p. 243–247 °C (from $CH_2Cl_2$) [lit. (*75*) 255–257 °C, lit. (*76*) 289 °C]; $^1H$ NMR (400 MHz, $D_6$-DMSO) δ 11.16 (1H, s), 7.99 – 7.88 (1H, m), 7.82 – 7.68 (2H, m), 5.16 (1H, dd, *J* 12.7, 5.5 Hz), 2.95 – 2.82 (1H, m), 2.70 – 2.50 (2H, m), 2.10 – 2.02 (1H, m); $^{19}F$ NMR (376 MHz, $D_6$-DMSO) δ −114.69; LRMS *m/z* ($ESI^-$) 275 ($[M-H]^-$, 100%); HPLC Retention time 220 nm: 6.8 min, 97.5%; 254 nm: 6.8 min, 98.5%. These data are in good agreement with the literature values (*77*).

**2-(2,6-Dioxopiperidin-3-yl)-5-fluoroisoindoline-1,3-dione (9)**

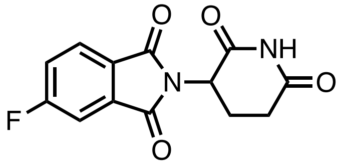

4-Fluorophthalic anhydride (2.00 g, 12.0 mmol, 1.0 eq), 3-aminopiperidine-2,6-dione hydrochloride (1.98 g, 12.0 mmol, 1.0 eq) and NaOAc (1.27 g, 18.1 mmol, 1.5 eq) were dissolved in AcOH (50 mL) and heated at 120 °C under reflux for 1.5 h. The AcOH was removed *in vacuo*, and the residue purified using flash column chromatography (5/95 MeOH/$CH_2Cl_2$) to afford a light lilac solid (1.92 g, 58%): $R_f$ 0.50 (5/95 MeOH/$CH_2Cl_2$); m.p. 245–250 °C (from $CH_2Cl_2$); $^1H$ NMR (400 MHz, $D_6$-DMSO) δ 11.15 (1H, s), 8.02 (1H, dd, *J* 8.3, 4.5 Hz), 7.86 (1H, dd, *J* 7.5, 2.4 Hz), 7.73 (1H, ddd, *J* 9.5, 8.3, 2.4 Hz), 5.17 (1H, dd, *J* 12.7, 5.4 Hz), 2.95 – 2.83 (1H, m), 2.65 – 2.51 (2H, m), 2.12 – 2.02 (1H, m); $^{19}F$ NMR (376 MHz, $D_6$-DMSO) δ −102.35; LRMS *m/z* ($ESI^-$) 275 ($[M-H]^-$, 100%); HPLC Retention time 220 nm: 7.1 min, 99.5%; 254 nm: 7.1 min, 100.0%. These data are in good agreement with the literature values (*78*).

**4-Chloro-2-(2,6-dioxopiperidin-3-yl)isoindoline-1,3-dione (10)**

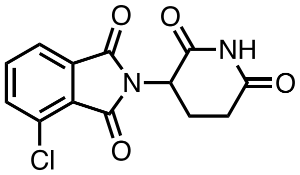

3-Chlorophthalic anhydride (155 mg, 0.849 mmol, 1.0 eq), 3-aminopiperidine-2,6-dione hydrochloride (140 mg, 0.851 mmol, 1.0 eq) and NaOAc (105 mg, 1.28 mmol, 1.5 eq) were dissolved in AcOH (5 mL) and heated at 120 °C under reflux for 20 h. After this time, the AcOH was removed *in vacuo* and the residue purified using flash column chromatography (2/98 MeOH/$CH_2Cl_2$) to afford a colorless solid (19 mg, 8%): $R_f$ 0.64 (5/95 MeOH/$CH_2Cl_2$); m.p. 273–277 °C (from $CH_2Cl_2$) [lit. (*73*) 290–291 °C]; $^1$H NMR (400 MHz, $D_6$-DMSO) δ 11.15 (1H, s), 7.95 – 7.84 (3H, m), 5.17 (1H, dd, *J* 12.8, 5.5 Hz), 2.94 – 2.85 (1H, m), 2.66 – 2.50 (2H, m), 2.12 – 2.02 (1H, m); LRMS *m/z* ($ESI^-$) 291 ($[M-H]^-$, 100%), 293 ($[M-H]^-$, 33%); HPLC Retention time 220 nm: 7.7 min, 100.0 %; 254 nm: 7.7 min, 96.4%. These data are in good agreement with the literature values (*73*).

**5-Chloro-2-(2,6-dioxopiperidin-3-yl)isoindoline-1,3-dione (11)**

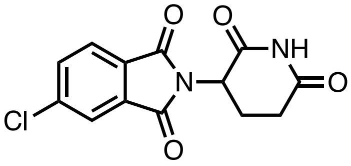

4-Chlorophthalic anhydride (155 mg, 0.849 mmol, 1.0 eq), 3-aminopiperidine-2,6-dione hydrochloride (140 mg, 0.851 mmol, 1.0 eq) and NaOAc (105 mg, 1.28 mmol, 1.5 eq) were dissolved in AcOH (5 mL) and heated at 120 °C under reflux for 20 h. The AcOH was removed *in vacuo* and the residue purified using flash column chromatography (2/98 MeOH/$CH_2Cl_2$) to afford a pale pink solid (68 mg, 28%): $R_f$ 0.50 (2/98 MeOH/$CH_2Cl_2$); m.p. >300 °C (from $CH_2Cl_2$) [lit. (*66*) 312–313 °C]; $^1$H NMR (400 MHz, $D_6$-DMSO) δ 11.15 (1H, s), 8.04 (1H, dd, *J* 1.2, 1.2 Hz), 7.97 – 7.93 (2H, m), 5.17 (1H, dd, *J* 12.8, 5.5 Hz), 2.95 – 2.85 (1H, m), 2.65 – 2.52 (2H, m), 2.11 – 2.01 (1H, m); LRMS *m/z* ($ESI^-$) 291 ($[M-H]^-$, 100%), 293 ($[M-H]^-$, 33%); HPLC Retention time 220 nm: 7.9 min, 95.2%; 254 nm: 7.9 min, 100.0%. These data are in good agreement with the literature values (*66*).

**4-Bromo-2-(2,6-dioxopiperidin-3-yl)isoindoline-1,3-dione (12)**

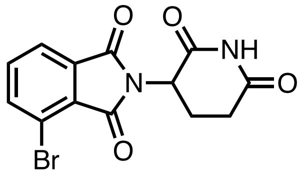

3-Bromophthalic anhydride (189 mg, 0.832 mmol, 1.0 eq), 3-aminopiperidine-2,6-dione hydrochloride (137 mg, 0.832 mmol, 1.0 eq) and NaOAc (103 mg, 1.25 mmol, 1.5 eq) were dissolved in AcOH (5 mL) and heated at 120 °C under reflux for 15 h. The AcOH was removed *in vacuo* and the residue purified using flash column chromatography (2/98 MeOH/$CH_2Cl_2$) to afford a colorless solid (115 mg, 41%): $R_f$ 0.68 (5.95 MeOH/$CH_2Cl_2$); m.p. 283–286 °C (from $CH_2Cl_2$); $^1$H NMR (400 MHz, $D_6$-DMSO) δ 11.15 (1H, s), 8.07 (1H, dd, *J* 8.1, 0.9 Hz), 7.94 (1H, dd, *J* 7.4, 0.9 Hz), 7.78 (1H, dd, *J* 8.1, 7.4 Hz), 5.17 (1H, dd, *J* 12.8, 5.4 Hz), 2.94 – 2.85 (1H, m), 2.66 – 2.50 (2H, m), 2.12 – 2.01 (1H, m); LRMS *m/z* ($ESI^-$) 335 ([M−H]$^-$, 97%), 337 ([M−H]$^-$, 100%); HPLC Retention time 220 nm: 7.9 min, 100.0%; 254 nm: 7.9 min, 98.8%. These data are in good agreement with the literature values (*79*).

**5-Bromo-2-(2,6-dioxopiperidin-3-yl)isoindoline-1,3-dione (13)**

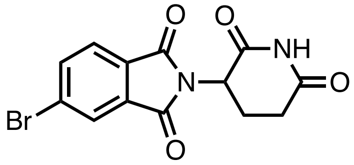

4-Bromophthalic anhydride (1.00 g, 4.18 mmol, 1.0 eq), 3-aminopiperidine-2,6-dione hydrochloride (688 mg, 4.18 mmol, 1.0 eq) and NaOAc (514 mg, 6.27 mmol, 1.5 eq) were dissolved in AcOH (30 mL) and heated at 120 °C under reflux for 1 h. The AcOH was removed *in vacuo*, and the residue purified using flash column chromatography (5/95 MeOH/$CH_2Cl_2$) to afford a colorless solid (585 mg, 42%): $R_f$ 0.53 (5/95 MeOH/$CH_2Cl_2$); m.p. 292–297 °C – Dec. (from $CH_2Cl_2$) [lit. (*80*) 230 °C, lit. (*68*) 240–241 °C, lit. (*81*) 303 °C]; $^1$H NMR (400 MHz, $D_6$-DMSO) δ 11.15 (1H, s), 8.15 (1H, dd, *J* 1.7, 0.6 Hz), 8.10 (1H, dd, *J* 7.9, 1.7 Hz), 7.87 (1H, dd, *J* 7.9, 0.6 Hz), 5.17 (1H, dd, *J* 12.8, 5.4 Hz), 2.93 – 2.84 (1H, m), 2.65 – 2.52 (2H, m), 2.11 – 2.01 (1H, m); LRMS *m/z* ($ESI^-$) 335 ([M−H]$^-$, 97%), 337 ([M−H]$^-$, 100%); HPLC Retention time 220 nm: 8.3 min, 96.2%; 254 nm: 8.3 min, 97.6%. These data are in good agreement with the literature values (*72*).

**2-(2,6-Dioxopiperidin-3-yl)-4-iodoisoindoline-1,3-dione (14)**

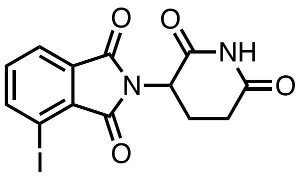

3-Iodophthalic acid (300 mg, 1.03 mmol, 1.0 eq), 3-aminopiperidine-2,6-dione hydrochloride (169 mg, 1.03 mmol, 1.0 eq) and NaOAc (126 mg, 1.54 mmol, 1.5 eq) were dissolved in AcOH (5 mL) and heated at 120 °C under reflux for 1 h. After this time the AcOH was removed *in vacuo* and the residue purified using flash column chromatography (2/98 MeOH/$CH_2Cl_2$) to afford a colorless solid (156 mg, 40%): $R_f$ 0.60 (5/95 MeOH/$CH_2Cl_2$); m.p. 286–290 °C (from $CH_2Cl_2$) [lit. (*80*) 304–305 °C]; $^1$H NMR (400 MHz, $D_6$-DMSO) δ 11.15 (1H, s), 8.28 (1H, dd, *J* 7.9, 0.9 Hz), 7.93 (1H, dd, *J* 7.4, 0.9 Hz), 7.58 (1H, dd, *J* 7.9, 7.4 Hz), 5.16 (1H, dd, *J* 12.8, 5.4 Hz), 2.94 – 2.83 (1H, m), 2.65 – 2.52 (2H, m), 2.11 – 2.01 (1H, m); LRMS *m/z* ($ESI^-$) 382.9 ($[M-H]^-$, 100%); HPLC Retention time 220 nm: 8.2 min, 95.9%; 254 nm: 8.2 min, 100.0%. These data are in good agreement with the literature values (*76*).

**2-(2,6-Dioxopiperidin-3-yl)-5-iodoisoindoline-1,3-dione (15)**

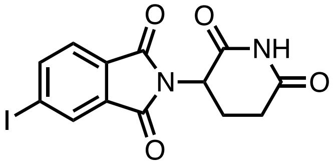

4-Iodophthalic acid (124 mg, 0.425 mmol, 1.0 eq), 3-aminopiperidine-2,6-dione hydrochloride (70 mg, 0.425 mmol, 1.0 eq) and NaOAc (53 mg, 0.646 mmol, 1.5 eq) were dissolved in AcOH (5 mL) and heated at 120 °C under reflux for 2 h. The AcOH was removed *in vacuo* and the residue purified using flash column chromatography (2/98 MeOH/$CH_2Cl_2$) to afford a colorless solid (93 mg, 57%): $R_f$ 0.48 (2/98 MeOH/$CH_2Cl_2$); m.p. 243–245 °C (from $CH_2Cl_2$); $\nu_{max}$ (thin film)/cm$^{-1}$ 1726 (C=O, s); $^1$H NMR (400 MHz, $D_6$-DMSO) δ 11.14 (1H, s), 8.31 – 8.24 (2H, m), 7.69 (1H, d, *J* 8.1 Hz), 5.15 (1H, dd, *J* 12.9, 5.4 Hz), 2.93 – 2.82 (1H, m), 2.65 – 2.51 (2H, m), 2.10 – 2.00 (1H, m); $^{13}$C NMR (151 MHz, $D_6$-DMSO) δ 172.7, 169.7, 166.8, 165.9, 143.5, 132.7, 131.8, 130.4, 124.9, 102.8, 49.1, 30.9, 21.9; HRMS *m/z* ($ESI^-$) [Found: 382.9538, $C_{13}H_8IN_2O_4$ requires $[M-H^-]$ 382.9534)]; LRMS *m/z* ($ESI^-$) 382.9 ($[M-H]^-$, 100%); HPLC Retention time 220 nm: 8.6 min, 96.6%; 254 nm: 8.6 min, 98.7%.

**2-(2,6-Dioxopiperidin-3-yl)-4-methylisoindoline-1,3-dione (16)**

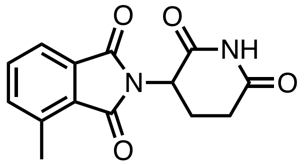

3-Methylphthalic anhydride (200 mg, 1.20 mmol, 1.0 eq), 3-aminopiperidine-2,6-dione hydrochloride (203 mg, 1.20 mmol, 1.0 eq) and NaOAc (152 mg, 1.81 mmol, 1.5 eq) were dissolved in AcOH (5 mL) and heated at 140 °C under reflux for 24 h. The residue was extracted from an aqueous solution of LiCl (0.5 M) with EtOAc. The EtOAc was removed *in vacuo*, and the residue purified using flash column chromatography (2/98 MeOH/$CH_2Cl_2$) to afford an off-white solid (135 mg, 40%): $R_f$ 0.64 (10/90 MeOH/$CH_2Cl_2$); m.p. 284–288 °C (from $CH_2Cl_2$) [lit. (*73*) 290–292 °C]; $^{1}$H NMR (400 MHz, $D_6$-DMSO) δ 11.11 (1H, s), 7.72 – 7.64 (3H, m), 5.13 (1H, dd, *J* 12.9, 5.4 Hz), 2.95 – 2.83 (1H, m), 2.63 (3H, s), 2.61 – 2.50 (2H, m), 2.11 – 1.99 (1H, m); LRMS *m/z* ($ESI^-$) 271 ($[M-H]^-$, 100%); HPLC Retention time 220 nm: 7.9 min, 96.7%; 254 nm: 7.9 min, 100.0%. These data are in good agreement with the literature values (*72*).

**2-(2,6-Dioxopiperidin-3-yl)-5-methylisoindoline-1,3-dione (17)**

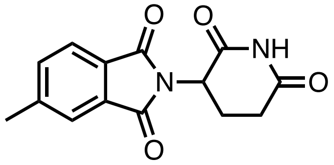

4-Methylphthalic anhydride (200 mg, 1.20 mmol, 1.0 eq), 3-aminopiperidine-2,6-dione hydrochloride (203 mg, 1.20 mmol, 1.0eq) and NaOAc (152 mg, 1.81 mmol, 1.5 eq) were dissolved in AcOH (5 mL) and heated at 120 °C under reflux for 2 h. The AcOH was removed *in vacuo* and the residue purified using flash column chromatography (5/95 MeOH/$CH_2Cl_2$) to afford a colorless solid (277 mg, 84%): $R_f$ 0.60 (5/95 MeOH/$CH_2Cl_2$); m.p. 260–266 °C (from $CH_2Cl_2$) [lit. (*82*) 265–267 °C]; $^{1}$H NMR (400 MHz, $D_6$-DMSO) δ 11.12 (1H, s), 7.88 (1H, d, J 7.7), 7.83 (1H, q, J 0.8), 7.76 (1H, dq, J 7.7, 0.8), 5.18 – 5.09 (1H, m), 2.96 – 2.82 (1H, m), 2.65 – 2.52 (5H, m), 2.11 – 2.00 (1H, m); LRMS *m/z* ($ESI^-$) 271 ($[M-H]^-$, 100%); HPLC Retention time 220 nm: 7.8 min, 98.7%; 254 nm: 7.8 min, 98.4%. These data are in good agreement with the literature values (*72*).

**2-(2,6-Dioxopiperidin-3-yl)-5-(trifluoromethyl)isoindoline-1,3-dione (18)**

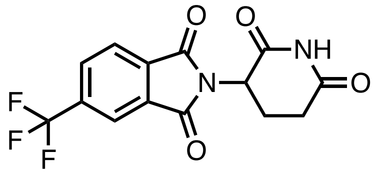

4-(Trifluoromethyl)phthalic acid (200 mg, 0.854 mmol, 1.0 eq), 3-aminopiperidine-2,6-dione hydrochloride (140 mg, 0.851 mmol, 1.0 eq) and NaOAc (105 mg, 1.28 mmol, 1.5 eq) were dissolved in AcOH (5 mL) and heated at 120 °C under reflux for 20 h. After this time the AcOH was removed *in vacuo* and the residue purified using flash column chromatography (5/95 MeOH/$CH_2Cl_2$) to afford a pale pink solid (192 mg, 69%): $R_f$ 0.75 (10/90 MeOH/$CH_2Cl_2$); m.p. 190–192 °C (from $CH_2Cl_2$); $\nu_{max}$ (thin film)/cm$^{-1}$ 1730 (C=O, s); $^1$H NMR (400 MHz, $D_6$-DMSO) δ 11.18 (1H, s), 8.30 (1H, s), 8.29 (1H, d, J 7.7), 8.15 (1H, d, J 7.7), 5.22 (1H, dd, *J* 12.8, 5.4 Hz), 2.96 – 2.84 (1H, m), 2.67 – 2.52 (2H, m), 2.13 – 2.03 (1H, m); $^{13}$C NMR (151 MHz, $D_6$-DMSO) δ 172.7, 169.6, 166.0, 165.8, 134.7, 134.5 (q, $J_{C\text{-}F}$ = 32.8 Hz), 132.2, 132.0 (d, $J_{C\text{-}F}$ = 21.2 Hz), 124.5 (d, $J_{C\text{-}F}$ = 22.4 Hz), 123.2 (q, $J_{C\text{-}F}$ = 273.1 Hz), 120.4 (d, $J_{C\text{-}F}$ = 21.2 Hz), 49.3, 30.87, 21.8; $^{19}$F NMR (565 MHz, $CDCl_3$) δ −56.61; HRMS *m/z* (ESI$^-$) [Found: 325.0439, $C_{14}H_8F_3N_2O_4$ requires [M−H$^-$] 325.0442)]; LRMS *m/z* (ESI$^-$) 325 ([M−H]$^-$, 100%); HPLC Retention time 220 nm: 8.5 min, 100.0%; 254 nm: 9.5 min, 100.0%.

**5-(*tert*-Butyl)-2-(2,6-dioxopiperidin-3-yl)isoindoline-1,3-dione (19)**

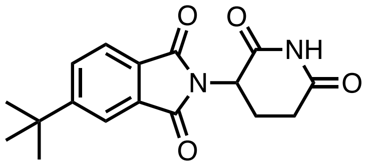

4-*tert*-Butylphthalic anhydride (200 mg, 0.979 mmol, 1.0 eq), 3-aminopiperidine-2,6-dione hydrochloride (161 mg, 0.978 mmol, 1.0 eq) and NaOAc (121 mg, 1.47 mmol, 1.5 eq) were dissolved in AcOH (5 mL) and heated at 120 °C under reflux for 21 h. The AcOH was removed *in vacuo* and the residue purified using flash column chromatography (10% MeOH/$CH_2Cl_2$) to afford a colorless solid (139 mg, 45%): $R_f$ 0.69 (10/90 MeOH/$CH_2Cl_2$); m.p. 145–155 °C (from $CHCl_3$); $^1$H NMR (400 MHz, $D_6$-DMSO) δ 11.12 (1H, s), 7.96 – 7.82 (3H, m), 5.14 (1H, dd, *J* 12.9, 5.4 Hz), 2.96 – 2.82 (1H, m), 2.65 – 2.52 (2H, m), 2.11 – 1.99 (1H, m), 1.36 (9H, s); LRMS *m/z* (ESI$^-$) 313 ([M−H]$^-$, 100%); HPLC Retention time 220 nm: 9.7 min, 99.8%; 254 nm: 9.7 min, 100.0%. These data are in good agreement with the literature values (*72*).

**2-(2,6-Dioxopiperidin-3-yl)-5-phenylisoindoline-1,3-dione (20)**

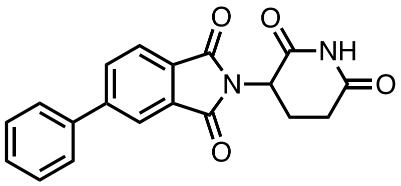

5-Phenylisobenzofuran-1,3-dione (125 mg, 0.557 mmol, 1.0 eq), 3-aminopiperidine-2,6-dione hydrochloride (92.0 mg, 0.559 mmol, 1.0 eq) and NaOAc (69.0 mg, 0.841 mmol, 1.5 eq) were dissolved in AcOH (5 mL) and heated at 120 °C under reflux for 2 h. The AcOH was removed *in vacuo* and the residue purified using flash column chromatography (2/98 MeOH/$CH_2Cl_2$) to afford a colorless solid (119 mg, 64%): $R_f$ 0.65 (5/95 MeOH/$CH_2Cl_2$); m.p. 216–220 °C (from $CH_2Cl_2$) [lit. (*80*) 230 °C]; $^1$H NMR (400 MHz, $D_6$-DMSO) δ 11.15 (1H, s), 8.23 – 8.15 (2H, m), 8.05 – 7.96 (1H, m), 7.88 – 7.81 (2H, m), 7.59 – 7.44 (3H, m), 5.19 (1H, dd, *J* 12.9, 5.5 Hz), 2.97 – 2.85 (1H, m), 2.66 – 2.52 (2H, m), 2.14 – 2.03 (1H, m); LRMS *m/z* (ESI$^-$) 333 ($[M-H]^-$, 100%); HPLC Retention time 220 nm: 9.5 min, 98.2%; 254 nm: 9.5 min, 99.7%. These data are in good agreement with the literature values (*80*).

**5-(Dimethylamino)-2-(2,6-dioxopiperidin-3-yl)isoindoline-1,3-dione (21)**

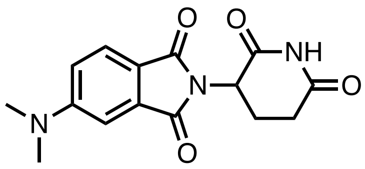

2-(2,6-Dioxopiperidin-3-yl)-5-fluoroisoindoline-1,3-dione (50.0 mg, 0.181 mmol, 1.0 eq), dimethylamine (2.0 M in THF, 2.90 mmol, 16.0 eq) and DIPEA (0.50 mL, 2.90 mmol, 16.0 eq) were dissolved in DMF (4 mL) and heated at 110 °C for 21 h. The DMF was removed by vigorously blowing nitrogen over the reaction vessel. The residual dark brown oil was purified using flash chromatography (45/55–50/50 EtOAc/petroleum ether) to afford a bright yellow powder (48 mg, 88%): $R_f$ 0.07 (45/55 EtOAc/petroleum ether); m.p. 241–243 °C (from EtOAc) [lit. (*83*) 240–242 °C]; $^1$H NMR (400 MHz, $CDCl_3$) δ 7.99 (1H, s), 7.67 (1H, d, *J* 8.6 Hz), 7.09 (1H, d, *J* 2.4 Hz), 6.82 (1H, dd, *J* 8.6, 2.4 Hz), 4.98 – 4.89 (1H, m), 3.13 (6H, s), 2.94 – 2.64 (3H, m), 2.21 – 2.08 (1H, m); LRMS *m/z* (ESI$^-$) 300 ($[M-H]^-$, 100%); HPLC Retention time 220 nm: 7.5 min, 98.9%; 254 nm: 7.5 min, 98.6%. These data are in good agreement with the literature values (*83*).

**2-(2,6-Dioxopiperidin-3-yl)-5-morpholinoisoindoline-1,3-dione (22)**

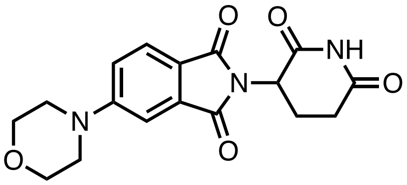

In a microwave vial, 2-(2,6-dioxopiperidin-3-yl)-5-fluoroisoindoline-1,3-dione (50.0mg, 0.181 mmol, 1.0 eq), morpholine (32 µL, 0.36 mmol, 2.0 eq) and DIPEA (0.13 mL, 0.72 mmol, 4.0 eq) were dissolved in NMP (2 mL) and heated at 110 °C for 2 h in a microwave. The NMP was removed by vigorously blowing nitrogen over the reaction vessel while heating to 60 °C. The residual dark brown oil was purified using flash chromatography (40/60 EtOAc/petroleum ether) to afford a bright yellow solid (23 mg, 37%): $R_f$ 0.09 (50/50 EtOAc/petroleum ether); m.p. 207–208 °C (from EtOAc); $\nu_{max}$ (thin film)/cm$^{-1}$ 1707 (C=O, s); $^1$H NMR (400 MHz, $CDCl_3$) δ 7.95 (1H, s), 7.73 (1H, d, *J* 8.5 Hz), 7.30 (1H, d, *J* 2.4 Hz), 7.08 (1H, dd, *J* 8.5, 2.4 Hz), 4.95 (1H, dd, *J* 12.3, 5.3 Hz), 3.88 (4H, dd, *J* 6.0, 3.8 Hz), 3.41 – 3.34 (4H, m), 2.95 – 2.68 (3H, m), 2.19 – 2.09 (1H, m); $^{13}$C NMR (151 MHz, $CDCl_3$) δ 170.9, 168.2, 167.9, 167.3, 155.8, 134.4, 125.5, 120.5, 118.1, 108.8, 66.5, 49.4, 47.8, 31.6, 22.9; HRMS *m/z* (ESI$^-$) [Found: 342.1098, $C_{17}H_{16}N_3O_5$ requires [M−H$^-$] 342.1095)]; LRMS *m/z* (ESI$^-$) 342 ([M−H]$^-$, 100%); HPLC Retention time 220 nm: 7.4 min, 96.5%; 254 nm: 7.4 min, 97.5%.

**2-(2,6-Dioxopiperidin-3-yl)-5-(4-methylpiperazin-1-yl)isoindoline-1,3-dione (23)**

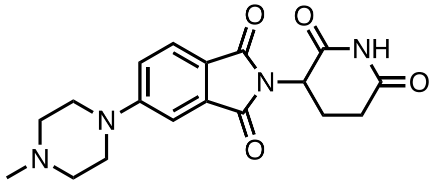

In a microwave vial, 2-(2,6-dioxopiperidin-3-yl)-5-fluoroisoindoline-1,3-dione (50.0 mg, 0.181 mmol, 1.0 eq), 1-methylpiperazine (40 µL, 0.36 mmol, 2.0 eq) and DIPEA (0.13 mL, 0.72 mmol, 4.0 eq) were dissolved in NMP (2 mL) and heated at 110 °C for 2 h in a microwave. The NMP was removed by vigorously blowing nitrogen over the reaction vessel while heating to 60 °C. The residual dark brown oil was purified using flash chromatography (2/98–10/90 MeOH/$CH_2Cl_2$) to afford a bright yellow solid (40 mg, 62%): $R_f$ 0.44 (10/90 MeOH/$CH_2Cl_2$); m.p. 192–194 °C (from EtOAc); $\nu_{max}$ (thin film)/cm$^{-1}$ 1709 (C=O, s); $^1$H NMR (400 MHz, $CDCl_3$) δ 8.40 (1H, s), 7.69 (1H, d, *J* 8.6 Hz), 7.28 (1H, d, *J* 2.4 Hz), 7.06 (1H, dd, *J* 8.6, 2.4 Hz), 4.98 – 4.88 (1H, m), 3.47 – 3.40 (4H, m), 2.93 – 2.64 (3H, m), 2.62 – 2.53 (4H, m), 2.35 (3H, s), 2.19 – 2.02 (1H, m); $^{13}$C NMR (151 MHz, $CDCl_3$) δ 171.0, 168.3, 168.0, 167.3, 155.6, 134.4, 125.5, 119.7, 118.2, 108.9, 54.6, 49.3, 47.6, 46.1, 31.6, 22.9; HRMS *m/z* (ESI$^-$) [Found: 355.1417, $C_{18}H_{19}N_4O_4$ requires [M−H$^-$] 355.1412)]; LRMS *m/z* (ESI$^-$) 355 ([M−H]$^-$, 100%); HPLC Retention time 220 nm: 5.1 min, 100.0%; 254 nm: 5.1 min, 100.0%.

**2-(2,6-Dioxopiperidin-3-yl)-4-(ethylamino)isoindoline-1,3-dione (24)**

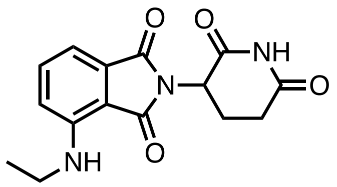

In a microwave vial, 2-(2,6-dioxopiperidin-3-yl)-4-fluoroisoindoline-1,3-dione (200 mg, 0.724 mmol, 1.0 eq), ethylamine (2.0 M in THF; 1.44 mL, 2.90 mmol, 4.0 eq) and DIPEA (0.50 mL, 2.9 mmol, 4.0 eq) were dissolved in NMP (4 mL) and heated at 110 °C for 2 h in a microwave. The NMP was removed by vigorously blowing nitrogen over the reaction vessel while heating to 60 °C. The residual oil was purified using flash column chromatography (40/60 EtOAc/petroleum ether) to afford a yellow powder (6 mg, 3%): $R_f$ 0.30 (30/70 EtOAc/petroleum ether); m.p. 209–211 °C (from EtOAc) [lit. (*83*) 221–224 °C]; $^1$H NMR (400 MHz, $CDCl_3$) δ 7.96 (1H, s), 7.50 (1H, ddd, *J* 8.5, 7.1, 0.6 Hz), 7.10 (1H, dd, *J* 7.1, 0.6 Hz), 6.89 (1H, d, *J* 8.5 Hz), 6.17 (1H, s), 4.96 – 4.87 (1H, m), 3.32 (2H, qd, *J* 7.2, 5.4 Hz), 2.98 – 2.64 (3H, m), 2.19 – 2.08 (1H, m), 1.31 (3H, t, *J* 7.2 Hz); LRMS *m/z* ($ESI^-$) 300 ($[M-H]^-$, 100%); HPLC Retention time 220 nm: 8.5 min, 97.9%; 254 nm: 8.5 min, 98.6%. These data are in good agreement with the literature values (*83*).

**2-(2,6-Dioxopiperidin-3-yl)-4-(prop-2-yn-1-ylamino)isoindoline-1,3-dione (25)**

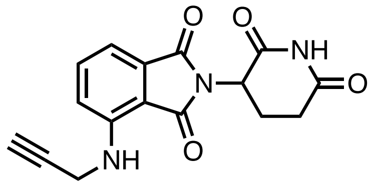

In a microwave vial, 2-(2,6-dioxopiperidin-3-yl)-4-fluoroisoindoline-1,3-dione (200 mg, 0.724 mmol, 1.0 eq), propargylamine (0.19 mL, 2.9 mmol, 4.0 eq) and DIPEA (0.50 mL, 2.9 mmol, 4.0 eq) were dissolved in NMP (4 mL) and heated at 110 °C for 2 h in a microwave. The NMP was removed by vigorously blowing nitrogen over the reaction vessel while heating to 60 °C. The residual oil was purified using flash column chromatography (30/70–50/50 EtOAc/petroleum ether) to afford a bright yellow solid (123 mg, 54%): $R_f$ 0.55 (60/40 EtOAc/petroleum ether); m.p. 174–176 °C (from EtOAc); $^1$H NMR (400 MHz, $CDCl_3$) δ 8.03 (1H, s), 7.57 (1H, dd, *J* 8.5, 7.2 Hz), 7.20 (1H, d, *J* 7.2 Hz), 7.03 (1H, d, *J* 8.5 Hz), 6.45 (1H, t, *J* 6.2 Hz), 4.92 (1H, dd, *J* 12.2, 5.3 Hz), 4.09 (2H, dd, *J* 6.2, 2.5 Hz), 2.94 – 2.67 (3H, m), 2.27 (1H, t, *J* 2.5 Hz), 2.18 – 2.08 (1H, m); LRMS *m/z* ($ESI^-$) 310 ($[M-H]^-$, 100%); HPLC Retention time 220 nm: 8.0 min, 96.7%; 254 nm: 8.1 min, 97.7%. These data are in good agreement with the literature values (*84*).

**4-(Benzylamino)-2-(2,6-dioxopiperidin-3-yl)isoindoline-1,3-dione (26)**

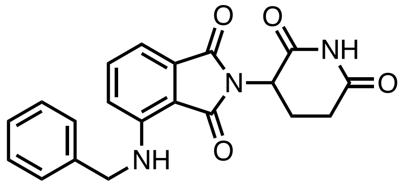

In a microwave vial, 2-(2,6-dioxopiperidin-3-yl)-4-fluoroisoindoline-1,3-dione (200 mg, 0.724 mmol, 1.0 eq), benzylamine (0.16 mL, 1.5 mmol, 2.0 eq) and DIPEA (0.50 mL, 2.9 mmol, 4.0 eq) were dissolved in NMP (4 mL) and heated at 110 °C for 2 h in a microwave. The NMP was removed by blowing nitrogen over it overnight while heating the vessel to 60 °C. The resultant oil was purified using column chromatography (50/50 EtOAc/petroleum ether) to afford a yellow solid (125 mg, 48%): $R_f$ 0.40 (50/50 EtOAc/petroleum ether); m.p. 199–201 °C (from EtOAc) [lit. (*85*) 209–211 °C]; $^{1}$H NMR (400 MHz, $CDCl_3$) δ 8.00 (1H, s), 7.45 (1H, dd, *J* 8.5, 7.1 Hz), 7.40 – 7.27 (5H, m), 7.12 (1H, d, *J* 7.1 Hz), 6.84 (1H, d, *J* 8.5 Hz), 6.70 (1H, t, *J* 6.0 Hz), 4.97 – 4.88 (1H, m), 4.52 (2H, d, *J* 6.0 Hz), 2.94 – 2.68 (3H, m), 2.20 – 2.10 (1H, m); LRMS *m/z* ($ESI^-$) 362 ($[M-H]^-$, 100%); HPLC Retention time 220 nm: 9.8 min, 99.5%; 254 nm: 9.8 min, 97.7%. These data are in good agreement with the literature values (*86*).

**2-(2,6-Dioxopiperidin-3-yl)-4-((pyridin-3-ylmethyl)amino)isoindoline-1,3-dione (27)**

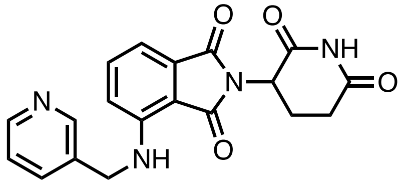

In a microwave vial, 2-(2,6-dioxopiperidin-3-yl)-4-fluoroisoindoline-1,3-dione (200 mg, 0.724 mmol, 1.0 eq), 3-picolylamine (0.15 mL, 1.5 mmol, 2.0 eq) and DIPEA (0.50 mL, 2.9 mmol, 4.0 eq) were dissolved in NMP (4 mL) and heated at 110 °C for 2 h in a microwave. The NMP was removed by vigorously blowing nitrogen over the reaction vessel while heating to 60 °C. The residual orange oil was purified using flash column chromatography (EtOAc) to afford a bright, light yellow solid (149 mg, 57%): $R_f$ 0.31 (EtOAc); m.p. 217–220 °C (from EtOAc); $\nu_{max}$ (thin film)/cm$^{-1}$ 1696 (C=O, s); $^{1}$H NMR (400 MHz, $CDCl_3$) δ 8.63 (1H, d, *J* 2.2 Hz), 8.56 (1H, dd, *J* 4.8, 1.6 Hz), 8.05 (1H, s), 7.68 (1H, ddd, *J* 7.9, 2.2, 1.6 Hz), 7.47 (1H, dd, *J* 8.5, 7.2 Hz), 7.30 (1H, dd, *J* 7.9, 4.8 Hz), 7.16 (1H, d, *J* 7.2 Hz), 6.82 (1H, d, *J* 8.5 Hz), 6.71 (1H, t, *J* 6.0 Hz), 4.93 (1H, dd, *J* 12.2, 5.3 Hz), 4.55 (2H, d, *J* 6.0 Hz), 2.95 – 2.68 (3H, m), 2.20 – 2.12 (1H, m); $^{13}$C NMR (151 MHz, $CDCl_3$) δ 170.9, 169.6, 168.3, 167.5, 149.4, 149.0, 146.4, 136.5, 134.8, 133.4, 132.7, 123.9, 117.0, 112.7, 111.1, 49.1, 44.6, 31.6, 22.9; HRMS *m/z* ($ESI^-$) [Found: 363.1100, $C_{19}H_{15}N_4O_4$ requires $[M-H^-]$ 363.1099)]; LRMS *m/z* ($ESI^-$) 363 ($[M-H]^-$, 100%); HPLC Retention time 220 nm: 5.8 min, 97.7%; 254 nm: 5.8 min, 100.0%.

**2-(2,6-Dioxopiperidin-3-yl)-4-((2-methylbenzyl)amino)isoindoline-1,3-dione (28)**

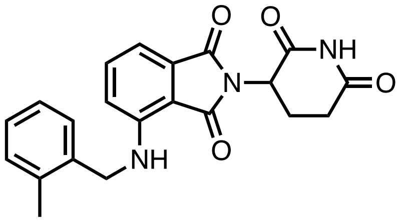

In a microwave vial, 2-(2,6-dioxopiperidin-3-yl)-4-fluoroisoindoline-1,3-dione (200 mg, 0.724 mmol, 1.0 eq), 2-methylbenzylamine (0.18 mL, 1.5 mmol, 2.0 eq) and DIPEA (0.50 mL, 2.9 mmol, 4.0 eq) were dissolved in NMP (4 mL) and heated at 110 °C for 2 h in a microwave. The NMP was removed by vigorously blowing nitrogen over the reaction vessel while heating to 60 °C. The residual orange oil was purified using flash column chromatography (40/60–60/40 EtOAc/petroleum ether) to afford a bright, light yellow solid (107 mg, 39%): $R_f$ 0.50 (50/50 EtOAc/petroleum ether); m.p. 218–221 °C (from EtOAc); $\nu_{max}$ (thin film)/cm$^{-1}$ 1690 (C=O, s); $^1$H NMR (400 MHz, $CDCl_3$) δ 7.97 (1H, s), 7.48 (1H dd, *J* 8.5, 7.1 Hz), 7.24 – 7.16 (4H, m), 7.12 (1H, d, *J* 7.1 Hz), 6.85 (1H, d, *J* 8.5 Hz), 6.52 (1H, t, *J* 5.6 Hz), 4.96 – 4.86 (1H, m), 4.45 (2H, d, *J* 5.6 Hz), 2.95 – 2.66 (3H, m), 2.38 (3H, s), 2.20 – 2.09 (1H, m); $^{13}$C NMR (151 MHz, $CDCl_3$) δ 171.0, 169.6, 168.3, 167.7, 146.88, 136.4, 136.1, 135.3, 132.6, 130.8, 127.9, 127.6, 126.5, 117.1, 112.1, 110.6, 49.1, 45.1, 31.6, 23.0, 19.2; HRMS *m/z* (ESI$^-$) [Found: 376.1306, $C_{21}H_{18}N_3O_4$ requires [M−H$^-$] 376.1303)]; LRMS *m/z* (ESI$^-$) 376 ([M−H]$^-$, 100%); HPLC Retention time 220 nm: 10.3 min, 95.4%; 254 nm: 10.3 min, 98.8%.

**2-(2,6-Dioxopiperidin-3-yl)-4-(((4-methylpyridin-3-yl)methyl)amino)isoindoline-1,3-dione (29)**

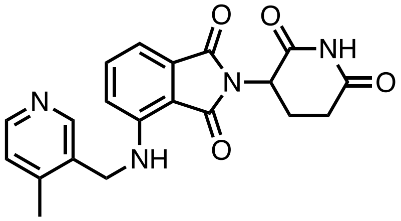

In a microwave vial, 2-(2,6-dioxopiperidin-3-yl)-4-fluoroisoindoline-1,3-dione (282 mg, 1.02 mmol, 1.0 eq), (4-methylpyridin-3-yl)methylamine (250 mg, 2.05 mmol, 2.0 eq) and DIPEA (0.71 mL, 4.1 mmol, 4.0 eq) were dissolved in NMP (4 mL) and heated at 110 °C for 2 h in a microwave. The NMP was removed by vigorously blowing nitrogen over the reaction vessel while heating to 60 °C. The residual orange oil was purified using flash column chromatography (EtOAc) to afford a bright yellow oil. TLC showed that the product was still impure; further flash column chromatography (2/98 MeOH/$CH_2Cl_2$) yielded a bright yellow/green glass-like oil, which was placed under vacuum for 72 h. Redissolving this in $CHCl_3$ and removal of solvent *via* nitrogen stream yielded a bright yellow powder (338 mg, 87%): $R_f$ 0.21 (2/98 MeOH/$CH_2Cl_2$); m.p. 199–202 °C (from EtOAc); $\nu_{max}$ (thin film)/cm$^{-1}$ 1694 (C=O, s); $^1$H NMR (400 MHz, $CDCl_3$) δ 8.51 (1H, s), 8.50 (1H, s), 8.45 (1H, d, *J* 4.9 Hz,), 7.50 (1H, dd, *J* 8.5, 7.1 Hz), 7.19 – 7.10 (2H, m), 6.89 (1H, d, *J* 8.5 Hz), 6.45 (1H, t, *J* 5.6 Hz), 4.90 (1H, dd, *J* 12.1, 5.3 Hz), 4.47 (2H, d, *J* 5.6 Hz),

2.92 – 2.66 (3H, m), 2.39 (3H, s), 2.18 – 2.08 (1H, m); $^{13}C$ NMR (151 MHz, $CDCl_3$) δ 170.9, 169.6, 168.3, 167.5, 149.3, 148.9, 146.4, 146.3, 136.5, 132.7, 131.4, 125.8, 116.8, 112.6, 111.1, 49.1, 43.1, 31.6, 22.9, 18.8; HRMS *m/z* ($ESI^-$) [Found: 377.1253, $C_{20}H_{17}N_4O_4$ requires $[M-H]^-$ 377.1255)]; LRMS *m/z* ($ESI^-$) 377 ($[M-H]^-$, 100%); HPLC Retention time 220 nm: 5.9 min, 99.5%; 254 nm: 5.9 min, 100.0%.

**4-(Benzyloxy)-2-(2,6-dioxopiperidin-3-yl)isoindoline-1,3-dione (30)**

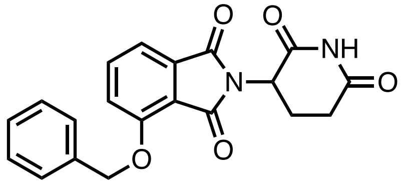

2-(2,6-Dioxopiperidin-3-yl)-4-hydroxyisoindoline-1,3-dione (100 mg, 0.364 mmol, 1.0 eq) was dissolved in dry DMF (5 mL) under an argon atmosphere. Benzyl bromide (43 µL, 0.36 mmol, 1.0 eq) and potassium carbonate (76.0 mg, 0.550 mmol, 1.5 eq) were added and the mixture stirred at room temperature for 1.5 h. The reaction mixture was dissolved in an aqueous solution of LiCl (0.5 M) and extracted with EtOAc. The organic layer was washed with water, brine, then dried with sodium sulfate, filtered, and the solvent removed *in vacuo*. The residue was purified using flash chromatography (5/95 MeOH/$CH_2Cl_2$). Further purification was achieved using a second round of flash chromatography (3/97 MeOH/$CH_2Cl_2$), as residual benzyl bromide was still present after the first column. This afforded a colorless solid (71 mg, 53%): $R_f$ 0.65 (10/90 MeOH/$CH_2Cl_2$); m.p. 223–226 °C (from $CH_2Cl_2$) [lit. (*71*) 230–234 °C, lit. (*87*) 238–240 °C]; $^1H$ NMR (400 MHz, $D_6$-DMSO) δ 11.10 (1H, s), 7.83 (1H, dd, *J* 8.5, 7.3 Hz), 7.60 (1H, d, *J* 8.5 Hz), 7.55 – 7.30 (6H, m), 5.38 (2H, s), 5.09 (1H, dd, *J* 12.8, 5.4 Hz), 2.95 – 2.83 (1H, m), 2.64 – 2.51 (1H, m), 2.09 – 1.99 (1H, m); LRMS *m/z* ($ESI^-$) 363 ($[M-H]^-$, 100%); HPLC Retention time 220 nm: 9.2 min, 96.7%; 254 nm: 9.2 min, 96.8%. These data are in good agreement with the literature values (*69*).

**2-(2,6-Dioxopiperidin-3-yl)-4-(pyridin-3-ylmethoxy)isoindoline-1,3-dione (31)**

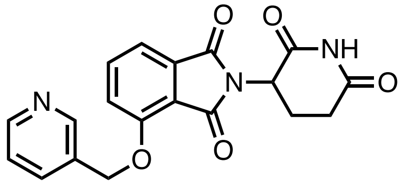

2-(2,6-Dioxopiperidin-3-yl)-4-hydroxy-1*H*-isoindole-1,3(2*H*)-dione (100 mg, 0.364 mmol, 1.0 eq) was dissolved in DMF (5 mL). Potassium carbonate (76.0 mg, 0.550 mmol, 1.5 eq) was then added and mixed. 3-(Bromomethyl)pyridine hydrobromide (82.0 mg, 0.324 mmol, 0.9 eq) was added and the mixture stirred at room temperature for 1.5 h. The DMF was removed by vigorously blowing nitrogen over the reaction vessel, and the residue was purified using flash chromatography (2/98 MeOH/$CH_2Cl_2$) to afford a colorless solid (69 mg, 59%): $R_f$ 0.38 (10/90 MeOH/$CH_2Cl_2$); m.p. 217–220 °C (from $CH_2Cl_2$); $\nu_{max}$ (thin film)/$cm^{-1}$ 1709 (C=O, s); $^1H$ NMR (400 MHz, $D_6$-

DMSO) δ 11.12 (1H, s), 8.73 (1H, s), 8.57 (1H, d, *J* 4.8 Hz), 7.93 (1H, d, *J* 6.8 Hz), 7.86 (1H, dd, *J* 7.7, 7.7 Hz), 7.64 (1H, d, *J* 8.6 Hz), 7.50 (1H, d, *J* 8.6 Hz), 7.46 (1H, dd, *J* 6.8, 4.8 Hz), 5.42 (1H, s), 5.09 (1H, dd, *J* 12.9, 5.2 Hz), 2.93 – 2.83 (1H, m), 2.65 – 2.52 (2H, m), 2.07 – 1.99 (1H, m); $^{13}$C NMR (151 MHz, $D_6$-DMSO) δ 172.7, 169.9, 166.7, 165.3, 155.3, 149.3, 148.7, 137.1, 135.3, 133.3, 131.8, 123.6, 120.2, 116.8, 115.8, 67.9, 48.8, 30.9, 22.0; HRMS *m/z* ($ESI^-$) [Found: 364.0940, $C_{19}H_{14}N_3O_5$ requires $[M{-}H^-]$ 364.0939)]; LRMS *m/z* ($ESI^-$) 364 ($[M{-}H]^-$, 100%); HPLC Retention time 220 nm: 5.7 min, 98.3%; 254 nm: 5.7 min, 100.0%.

**2-(2,6-Dioxopiperidin-3-yl)-4-((2-methylbenzyl)oxy)isoindoline-1,3-dione (32)**

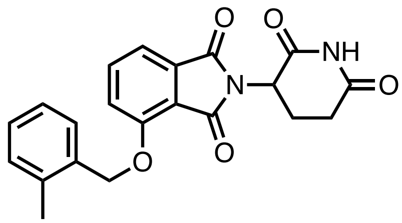

2-(2,6-Dioxopiperidin-3-yl)-4-hydroxy-1*H*-isoindole-1,3(2*H*)-dione (100 mg, 0.364 mmol, 1.0 eq) was dissolved in DMF (5 mL). Potassium carbonate (76.0 mg, 0.550 mmol, 1.5 eq) was then added and mixed. 2-Methylbenzyl bromide (44 μL, 0.32 mmol, 0.9 eq) was then added and the mixture stirred at room temperature for 1.5 h. The DMF was removed by vigorously blowing nitrogen over the reaction vessel, and the residue was purified using flash chromatography (2/98–5/95 MeOH/$CH_2Cl_2$) to afford a colorless solid (117 mg, 97%): $R_f$ 0.56 (10% MeOH/$CH_2Cl_2$); m.p. 213–215 °C (from $CH_2Cl_2$); $\nu_{max}$ (thin film)/cm$^{-1}$ 1704 (C=O, s); $^1$H NMR (400 MHz, $D_6$-DMSO) δ 11.12 (1H, s), 7.85 (1H, dd, *J* 8.6, 7.2 Hz), 7.68 (1H, d, *J* 8.6 Hz), 7.55 (1H, d, *J* 7.0 Hz), 7.48 (1H, d, *J* 7.2 Hz), 7.31 – 7.18 (3H, m), 5.35 (2H, s), 5.09 (1H, dd, *J* 12.8, 5.4 Hz), 2.93 – 2.82 (1H, m), 2.63 – 2.51 (2H, m), 2.36 (3H, s), 2.08 – 1.99 (1H, m); $^{13}$C NMR (151 MHz, $D_6$-DMSO) δ 172.7, 169.9, 166.8, 165.3, 155.5, 137.0, 136.5, 134.1, 133.3, 130.1, 128.2, 128.0, 125.8, 120.2, 116.6, 115.5, 68.8, 48.8, 30.9, 22.0, 18.5; HRMS *m/z* ($ESI^-$) [Found: 377.1144, $C_{21}H_{17}N_2O_5$ requires $[M{-}H^-]$ 377.1143)]; LRMS *m/z* ($ESI^-$) 377 ($[M{-}H]^-$, 100%); HPLC Retention time 220 nm: 9.7 min, 96.2%; 254 nm: 9.7 min, 97.4%.

**4-(Benzhydrylamino)-2-(2,6-dioxopiperidin-3-yl)isoindoline-1,3-dione (33)**

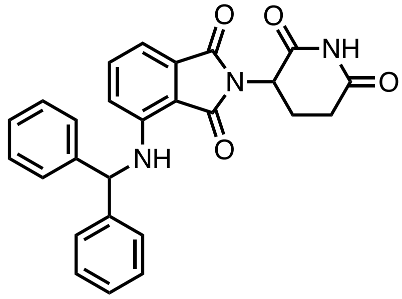

2-(2,6-Dioxopiperidin-3-yl)-4-fluoroisoindoline-1,3-dione (100 mg, 0.362 mmol, 1.0 eq), benzhydrylamine (0.25 mL, 1.5 mmol, 4.0 eq) and DIPEA (0.25 mL, 1.5 mmol, 4.0 eq) were dissolved in NMP (4 mL) and heated at 110 °C for 20 h. The NMP was removed by vigorously blowing nitrogen over the reaction vessel while heating to 60 °C. The residual dark brown oil was purified using flash column chromatography (50/50 EtOAc/petroleum ether) from which an impure product was gained; further flash column chromatography (30/70–40/60 EtOAc/petroleum ether) yielded a bright yellow powder (22 mg, 14%): $R_f$ 0.36 (50/50 EtOAc/petroleum ether); m.p. 86–88 °C (from EtOAc); $\nu_{max}$ (thin film)/cm$^{-1}$ 1701 (C=O, s); $^1$H NMR (600 MHz, $CDCl_3$) δ 7.92 (1H, s), 7.41 – 7.27 (11H, m), 7.12 (1H, d, *J* 8.5 Hz), 6.86 (1H, d, *J* 5.5 Hz), 6.71 (1H, d, *J* 8.5 Hz), 5.69 (1H, d, *J* 5.5 Hz), 4.92 (1H, dd, *J* 12.5, 5.4 Hz), 2.92 – 2.68 (3H, m), 2.18 – 2.08 (1H, m); $^{13}$C NMR (151 MHz, $CDCl_3$) δ 170.8, 169.6, 168.3, 167.6, 145.9, 141.44, 141.41, 136.2, 132.5, 129.22, 129.21, 128.06, 128.03, 127.35, 127.31, 118.3, 112.4, 111.0, 61.9, 49.1, 31.6, 23.0; HRMS *m/z* ($ESI^-$) [Found: 438.1460, $C_{26}H_{20}N_3O_4$ requires [M−H$^-$] 438.1459)]; LRMS *m/z* ($ESI^-$) 438 ([M−H]$^-$, 100%); HPLC Retention time 220 nm: 11.0 min, 96.0%; 254 nm: 11.1 min, 99.1%.

**4-(Benzhydryloxy)-2-(2,6-dioxopiperidin-3-yl)isoindoline-1,3-dione (34)**

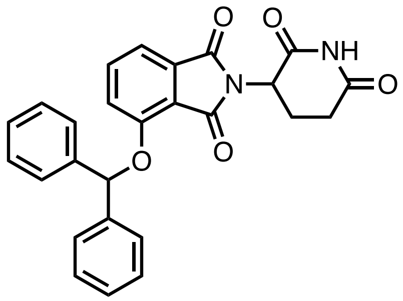

2-(2,6-Dioxopiperidin-3-yl)-4-hydroxyisoindoline-1,3-dione (100 mg, 0.364 mmol, 1.0 eq) was dissolved in dry DMF (5 mL) under an argon atmosphere. Benzhydryl bromide (95.0 mg, 0.364 mmol, 1.0 eq) and potassium carbonate (76.0 mg, 0.550 mmol, 1.5 eq) were added and the mixture stirred at room temperature for 1.5 h. The DMF was then removed by vigorously blowing nitrogen over the reaction vessel. The crude product was purified using flash chromatography (2/98 MeOH/$CH_2Cl_2$) to afford a colorless solid (100 mg, 62%): $R_f$ 0.53 (10/90 MeOH/$CH_2Cl_2$); m.p. 201–204 °C (from $CH_2Cl_2$); $\nu_{max}$ (thin film)/cm$^{-1}$ 1713 (C=O, s); $^1$H NMR (400 MHz, $D_6$-DMSO) δ 11.15 (1H, s), 7.72 (1H, dd, *J* 8.6, 7.3 Hz), 7.63 – 7.56 (4H, m), 7.50 (1H, d, *J* 8.6 Hz), 7.42 (1H, d, *J* 7.3 Hz), 7.40 – 7.34 (4H, m), 7.30 – 7.23 (2H, m), 6.90 (1H, s), 5.14 (1H, dd, *J* 12.9, 5.4 Hz), 2.97 – 2.85 (1H, m), 2.67 – 2.51 (2H, m), 2.12 – 2.02 (1H, m); $^{13}$C NMR (151 MHz, $D_6$-DMSO) δ 173.3, 170.4, 167.2, 165.9, 154.8, 141.37, 141.35, 137.2, 133.8, 129.20, 129.18, 128.30, 128.28, 126.67, 126.64, 121.7, 117.9, 116.2, 80.8, 49.3, 31.5, 22.5; HRMS *m/z* ($ESI^-$) [Found: 439.1300, $C_{26}H_{19}N_2O_5$ requires [M−H$^-$] 439.1299)]; LRMS *m/z* ($ESI^-$) 439 ([M−H]$^-$, 100%); HPLC Retention time 220 nm: 10.5 min, 95.9%; 254 nm: 10.5 min, 96.3%.

**6-(2,6-Dioxopiperidin-3-yl)-5*H*-pyrrolo[3,4-*b*]pyridine-5,7(6*H*)-dione (35)**

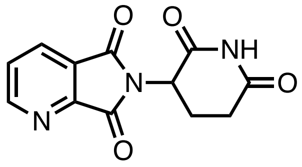

2,3-Pyridinedicarboxylic anhydride (201 mg, 1.35 mmol, 1.0 eq), 3-aminopiperidine-2,6-dione hydrochloride (222 mg, 1.35 mmol, 1.0 eq) and NaOAc (166 mg, 2.03 mmol, 1.5 eq) were dissolved in AcOH (5 mL) and heated at 120 °C under reflux for 19 h. The AcOH was removed *in vacuo* and the residue purified using flash column chromatography (5/95 MeOH/$CH_2Cl_2$) to afford a colorless solid (200 mg, 57%): $R_f$ 0.50 (10/90 MeOH/$CH_2Cl_2$); m.p. 245–247 °C (from $H_2O$/MeCN) [lit. (*88*) 258–259 °C, lit. (*89*) 266–267 °C]; $^1$H NMR (400 MHz, $D_6$-DMSO) δ 11.17 (1H, s), 9.04 (1H, dd, J 4.9, 1.4), 8.38 (1H, dd, J 7.7, 1.4), 7.85 (1H, dd, J 7.7, 4.9), 5.24 (1H, dd, *J* 12.9, 5.4 Hz), 2.96 – 2.84 (1H, m), 2.67 – 2.52 (2H, m), 2.13 – 2.03 (1H, m); LRMS *m/z* (ESI$^-$) 258 ([M−H]$^-$, 100%); HPLC Retention time 220 nm: 5.1 min, 99.7%; 254 nm: 5.2 min, 100.0%. These data are in good agreement with the literature values (*72*).

**2-(2,6-Dioxopiperidin-3-yl)-1*H*-pyrrolo[3,4-*c*]pyridine-1,3(2*H*)-dione (36)**

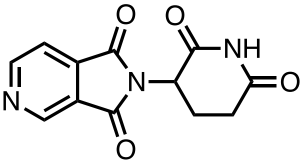

3,4-Pyridinedicarboxylic anhydride (201 mg, 1.35 mmol, 1.0 eq), 3-aminopiperidine-2,6-dione hydrochloride (222 mg, 1.35 mmol, 1.0 eq) and NaOAc (166 mg, 2.03 mmol, 1.5 eq) were dissolved in AcOH (5 mL) and heated at 120 °C under reflux for 19.5 h. The AcOH was removed *in vacuo* and the residue purified using flash column chromatography (5/95 MeOH/$CH_2Cl_2$) to afford a colorless solid (314 mg, 90%): $R_f$ 0.61 (10/90 MeOH/$CH_2Cl_2$); m.p. 219–221 °C (from $H_2O$/MeCN) [lit. (*90*) 233–235 °C, lit. (*89*) 234–235 °C]; $^1$H NMR (400 MHz, $D_6$-DMSO) δ 11.18 (1H, s), 9.22 – 9.11 (2H, m), 7.97 (1H, dd, *J* 4.8, 1.1 Hz), 5.22 (1H, dd, *J* 12.9, 5.4 Hz), 2.97 – 2.85 (1H, m), 2.67 – 2.50 (2H, m), 2.12 – 2.02 (1H, m); LRMS *m/z* (ESI$^-$) 258 ([M−H]$^-$, 100%); HPLC Retention time 220 nm: 5.2 min, 99.4%; 254 nm: 5.2 min, 100.0%. These data are in good agreement with the literature values (*72*).

Biological Methods

**Cell Culture**

Jurkat cells were cultured in a media of RPMI (Gibco), 10% heat-inactivated FBS (Gibco), 1% Glutamax (Gibco), 1% MEM non-essential amino acids (Gibco), 1% sodium pyruvate (Gibco) and 1% PenStrep (Sigma). Lenti-X 293-T cells (Takara) were cultured in a media of DMEM high glucose (Gibco), 10% Tet-free FBS (Takara), 1% sodium pyruvate and 1% PenStrep. HEK293T cells were cultured in a media of DMEM, 10% heat-inactivated FBS, 1% MEM non-essential amino acids and 1% PenStrep. Cells were incubated at 37 °C and 5% $CO_2$.

**Cloning**

The degradation reporter plasmid, pLVX-Sprout, was generated by inserting a gene fragment encoding for IKZF3 130aa-142aa 170-189aa fused to EGFP *via* a linker into pLVX-EF1a-IRES-mCherry (Takara) using XbaI/BamHI restriction sites. Gene fragments encoding for IKZF3 143aa-169aa sequences were cloned into pLVX-Sprout using the Golden Gate Assembly Kit BsmBI-v2 (New England Biolabs). Gene fragments encoding for either HiBit-TRIM28, HiBit-TRIM28 N-terminal degron or HiBit-TRIM28 C-terminal degron were inserted into pLVX-EF1a-IRES-mCherry (Takara) using XbaI/BamHI restriction sites.

**Lentivirus Vector Production and Transduction**

The below method was used to generate lentiviral vectors for plasmids used in initial compound screening (WT and Q147A) and TRIM28 degradation (TRIM28, TRIM28 N-degron and TRIM28 C-degron). The following method was also used to generate the library lentiviral vector pool. Lenti-X 293-T cells were seeded into 100 mm TC-treated cell culture dishes (Falcon). When cells were 90% confluent, 7 µg transgene plasmid was mixed with Lenti-X™ Packaging Single Shot (VSV-G) (Takara) and incubated for 10 min at room temperature before being added to cells. After 6 h, Viralboost (Alstem) was added at a 500X dilution. After 3 days, the supernatant was harvested and centrifuged at 1000 x g for 10 minutes, then supernatant was passed through a 0.22 µm PES filter. Lenti-X concentrator (Takara) was added at a 4X dilution, and the supernatant was incubated at 4 °C for 6 h. Supernatant was centrifuged at 4 °C 1500 x g for 45 min, supernatant was removed, and virus pellet was resuspended at a 50X concentration in cell culture media used for Jurkat cells. Lentivirus vector aliquots were frozen at −80 °C. For transductions, lentivirus vectors were added to Jurkat cells in 6 well, TC-treated, cell culture dishes (Falcon) with $3 \times 10^6$ cells per well and incubated for 2 days at 37 °C and 5% $CO_2$. After 2 days, the supernatant was removed and replaced with fresh media.

The following method was used to generate lentiviral vectors for plasmids tested in library hit confirmation experiments: WT, QPIS, ANQS, APIS, HEIP, FEVP, WEIK, FDIP, AEVK, WNQS, WEIS, WNIQ, and WDIA. HEK293T cells were seeded into 6 well, TC-treated, cell culture dishes (Falcon) at $7 \times 10^5$ cells per well and incubated overnight at 37 °C and 5% $CO_2$. 888 ng of transgene plasmid, 444 ng of psPAX2 (Addgene Cat# 12260), and 222 ng of pMD2.G (Addgene Cat#12259) were combined in 100 µL of OptiMEM (Thermo). In a separate tube, of PEIPro (Polyplus, 2 µL) was added to OptiMEM (100 µL). The solution containing PEIPro and DNA was mixed and incubated at room temperature for 15 min to create the transfection mixture. The transfection mixture was added to cells and incubated for 5–6 h at 37 °C and 5% $CO_2$. After 2 days, the supernatant was harvested, and fresh media was added to cells. After 24 h, the supernatant was harvested and combined with the first supernatant harvested. Supernatant was centrifuged at 2000 x g room temperature for 5 min, then the supernatant was filtered through a 0.45 µm cellulose acetate syringe filter and frozen at −80 °C. For transductions, lentivirus vectors were added to Jurkat cells in 6 well, TC-treated, cell culture dishes (Falcon) with $2.5 \times 10^6$ cells per well in media containing 4 ug/mL polybrene (Sigma). The cells were centrifuged at 500 x g for 60 min at 30 °C. Cells were incubated for 3 days at 37 °C and 5% $CO_2$ prior to replacing supernatant with fresh media. (psPAX2 and pMD2.G were a gift from Didier Trono's lab).

**GFP Degradation Flow Cytometry Assay**

Test compounds (10 mM stock in DMSO) were either serially diluted 10-fold to create a 6 point dilution spanning 1 nM–100 µM (final assay concentration) or serially diluted 2-fold to create a 12 point dilution spanning 5 nM–10 µM. Jurkat cells were added to 96 well U-bottom assay plates (Falcon) at $1 \times 10^5$ cells per well and treated with compounds for 18 hours at 37 °C and 5% $CO_2$. Cells were stained with 1µg/mL DAPI (ThermoFisher) and acquired on a Cytoflex S (Beckman Coulter). Data were analyzed using FlowJo 10.8.1 (BD).

**Compound Solubility**

A DMSO stock solution of IMiD analog (5 µL, 10 mM) was diluted to a volume of 100 µL with pH 7.4 PBS, equilibrated for 1 h at room temperature and filtered through Millipore Multiscreen HTS-PCF filter plates (MSSL BPC). The filtrate was quantified by suitably calibrated Charged Aerosol Detector (*91*).

**Protein Expression and Purification**

Wild-type human 6×His-TEV-Cereblon and 6×His-TEV-DDB1ΔBPB were cloned in pFastbac-HTb (Genscript), and recombinant proteins were expressed in Super 9 SFX insect cells (SH3A3187.02, Cytiva) using the BacMam baculovirus expression system (ThermoFisher). For protein purification cells were resuspended in buffer containing 500 mM NaCl (Sigma), 10% glycerol (Sigma), 50 mM Tris pH 7.7 (Sigma), 15 mM imidazole (Sigma), 90.5 mM TCEP (Sigma), 5 µL/mL protease inhibitor cocktail (Sigma) and 0.66 µL/mL benzonase (Novagen). Cells were lysed by sonication and pelleted by ultra-centrifugation. The supernatant was passed over Ni-NTA affinity resin (Qiagen) and eluted in buffer containing 500 mM NaCl, 10% glycerol, 50 mM Tris pH 7.7 and 90.5 mM TCEP supplemented with 350 mM imidazole. Proteins were further purified using size exclusion chromatography in 50 mM Tris pH 7.5, 5% glycerol, 2 mM DTT (Sigma), and 150 mM NaCl. Protein fractions were concentrated by ultrafiltration using an Ultra 10 K MWCO device (Amicon), then flash frozen in liquid nitrogen and stored at −80 °C.

**CRBN Binding**

Test compounds (10 mM stock in DMSO) were serially diluted 4-fold to create an 11-point dilution spanning 0.1 nM–100 µM. 100 nL of compounds were added to black 384 well low volume assay plates (Greiner) using an Echo 555 acoustic dispenser (Labcyte). 5 µL 6×His-TEV-Cereblon/6×His-TEV-DDB1ΔBPB (5 nM final assay concentration) containing Alexa647 labelled lenalidomide (50 nM final assay concentration) were added to the assay plates using a Multidrop Combi dispenser (ThermoFisher) in assay buffer (50 mM HEPES (Sigma), 150 mM NaCl (Sigma), 5% Glycerol (Sigma), 1 mM CHAPS (Sigma) and 1 mM DTT (Sigma) pH 7.4). Assay plates were centrifuged at 1000 rpm for 1 min and then incubated for 15 min at rt. 5 µL Eu-W1024-labeled Anti-6×His antibody (1 nM final assay concentration) (Perkin Elmer) in assay buffer was added to all wells, plates were centrifuged at 1000 rpm for 1 min and then incubated for 30 min at room temperature. After excitation of europium fluorescence at 337 nm, emission at 615 nm (donor, europium) and 665 nm (acceptor, Alexa647) were recorded with a 20 µs delay to reduce background fluorescence on an Envision 2104 microplate reader (Perkin Elmer). The data were reported as a 665/615 ratio and normalized between high (no test compound) and low (no protein) controls.

**Endogenous IKZF1 Immunostaining**

Test compounds (10 mM stock in DMSO) were serially diluted 10-fold to create a 6-point dilution spanning 1 nM–100 µM (final assay concentration). Jurkat cells were added to 96 well U-bottom assay plates (Falcon) at 1 x $10^5$ cells per well and treated with compounds for 18 hours at 37 °C and 5% $CO_2$. Following incubation, cells were stained with LIVE/DEAD™ Fixable Aqua Dead Cell Stain (Thermo) diluted 1000X in PBS for 30 min at room temperature and then incubated with Human TruStain FcX block (Biolegend) diluted 40X in PBS + 1% FBS (Gibco) for 10 min at room temperature. Intranuclear staining of IKZF1 was carried out using the Transcription Factor Buffer Set (BD Bioscience) according to the manufacturer's protocol and staining with IKZF1 AF488 antibody (clone R32-1149, BD Bioscience) diluted 40X for 30 minutes at room temperature. Cells were acquired on a Cytoflex S (Beckman Coulter) and data analysed using FlowJo 10.8.1 9 (BD).

**Pooled Library Construction**

A library of 8380 IKZF3 degrons was cloned into pLVX-Sprout to create the EGFP/mCherry protein degradation reporter plasmid pool. A lentivirus pool was then generated using the Lenti-X lentiviral vector production method.

**Pooled Library Screen**

Jurkat cells were transduced with the lentivirus vector library at a low level of <30% to maximise the number of cells with a single integration. Transductions were carried out in 6 well, TC-treated, cell culture dishes (Falcon) with 3 × $10^6$ cells per well. Cells were incubated for 2 days at 37 °C and 5% $CO_2$ before the supernatant was removed and replaced with fresh media. Six days post-transduction, mCherry+ cells were isolated *via* FACS using the FACS Aria Fusion (BD) in a buffer containing PBS (Gibco), 2 mM EDTA (Gibco), 25 mM HEPES (Sigma) and 1% BSA (Sigma). After 8 days, cells were treated with DMSO, or a thalidomide analog, for 18 h at 37 °C and 5% $CO_2$. $EGFP_{low}$ and EGFP cells were sorted by FACS using the FACS Aria Fusion (BD). DMSO treated cells were used to set the $EGFP_{low}$ gate and dead cells were excluded from sorting by staining cells with 1 µg/mL DAPI (ThermoFisher). gDNA was isolated using the Maxwell RSC 48 (Promega). gDNA from cells that were unsorted and untreated were also isolated to determine the library representation on the day of compound treatment. Throughout the screen >600X coverage of the library was maintained. Three biological replicates were carried out with a new transduction taking place each time.

**Next-Generation Sequencing (NGS)**

A two-step PCR was used where the first PCR (PCR1) amplified the integrated degron sequence from gDNA and the second PCR (PCR2) added Illumina adaptors and barcodes to permit multiplexing. See **Sequences for NGS** for PCR1 and PCR2 primers. Q5 Hot Start High-Fidelity 2X Master Mix (New England Biolabs) was used for both PCRs. Sufficient gDNA was amplified in PCR1 in order to maintain >600X coverage and amplicons were purified using solid phase reversible immobilization (SPRI) beads (Beckman Coulter). Purified PCR1 products were analysed on the Tapestation 4200 (Agilent) to confirm purity prior to proceeding to PCR2. Sufficient PCR1 product was amplified in PCR2 so as to maintain >600X coverage and amplicons were run on a 2% agarose TAE gel. Target bands were excised and purified using the NucleoSpin Gel and PCR Clean up Kit (Machery Nagel). Amplicons were analysed on the Tapestation 4200 and quantified using Qubit dsDNA HS kit (ThermoFisher). PCR products were pooled and sequenced as paired-reads on the NovaSeq (Illumina) using a SP2 200 cycles kit (Illumina).

**NGS Data Analysis**

Sample fastq files were processed to align reads to reference degron sequences using the alignment tool Vsearch (*92*) to count the number of times each degron was present within each sample. Count distributions, percentage of read mapping, library coverage and replicate correlation per sample were assessed for QC purposes. Counts for each degron were then subject to Limma Voom linear regression analysis (*93*) to generate an estimated log fold change between replicate samples in the positively ($EGFP_{low}$) sorted fraction *vs* replicate samples in the negatively ($EGFP_{+}$) sorted fraction for each degron sequence.

**Immunoblotting**

Test compounds (10 mM stock in DMSO) were serially diluted 10-fold to create a 3 or 6 point dilution spanning 100 nM–10 µM or 1 nM–10 µM, respectively (final assay concentration). Jurkat cells were added to T25 flasks (Falcon) at $2 \times 10^6$ cells per flask and were incubated with compounds for 18 hours at 37 °C and 5% $CO_2$. Cells were lysed in buffer containing 9 M urea (Sigma), 75 mM Tris-HCl (Sigma), 0.15 M β-mercaptoethanol (Merck) and 10X diluted protease inhibitor cocktail (Merck). Cells lysates were sonicated for 10 secs 3 times, centrifuged at 13000 rpm for 10 min at 4 °C and the supernatants were transferred into fresh tubes. Protein concentrations were measured using a Pierce BCA Protein Assay Kit (Thermo). Proteins were boiled in Laemmli sample buffer (Bio-Rad) for 5 min at 100 °C and separated by SDS-PAGE (7.5% Mini-PROTEAN TGX gels, Bio-Rad) and compared to the PageRuler Prestained Protein Ladder (Thermo). Semi-dry blotting was done on a Trans-Blot Turbo Transfer System (Bio-Rad) with the Trans-Blot Turbo RTA Mini 0.2 µm PVDF Transfer Kit (Bio-Rad) using the default mixed molecular weight mode. Membranes were blocked in 5% non-fat dry milk (Bio Rad) in TBS-T. Primary IKZF3 (D1C1E, Cell Signaling Technology Europe), EGFP (F56-6A1.2.3, Thermo) and vinculin (E1E9V, Cell Signaling Technology Europe and sc-25336, Santa Cruz Biotechnologies) antibodies were diluted 1000X, 500X and 50X respectively in 5% non-fat dry milk in TBS-T. Membranes were incubated with primary antibodies overnight at 4 °C, washed with TBS-T three times, incubated with HRP conjugated secondary antibodies (Promega) for 1 h at room temperature, washed with TBS-T three times and imaged with Clarity Western ECL substrate (Bio-Rad). Membranes were scanned with a Bio-Rad ChemiDoc XRS+ Imaging System (Bio-Rad) and signal intensities were quantified using Image Lab software (Bio Rad). Band densitometry was assessed, normalized to vinculin bands, and reported as percentage of the DMSO control lane.

**CRISPR Editing**

Jurkat cells were edited eight days after lentiviral transduction. For each gene, three sgRNAs targeting a single exon were designed using the Synthego CRISPR Design tool. For sequences see ‘sgRNA sequences’ below. sgRNA were resuspended to 100 µM in IDT duplex buffer and then combined to create pools of sgRNA. Cas9 RNPs were prepared by combining 2.5 µL 36.6 µM Alt-R® S.p. Cas9 Nuclease V3 (IDT) and 2 µL 100 µM sgRNA pool in IDT duplex buffer and incubating for 10 minutes at room temperature. $1.5 \times 10^6$ cells in 20 µL SE buffer (Lonza) were mixed with the Cas9 RNPs and then electroporated using a 4D-Nucleofector (Lonza) on pulse code CL-120. Estimation of knockout efficiency was performed by PCR amplification of the sgRNA target site using primers described in 'PCR primers for amplifying sgRNA target site' below. PCR products were Sanger sequenced and editing efficiency was assessed using the online ICE webtool (*94*).

**HiBit and Cell Titre Glo (CTG)**

Compound (10 mM stock in DMSO) was serially diluted 3-fold to create an 11-point dilution spanning 169 pM–10 µM (final assay concentration). Jurkat cells transduced with Hibit-TRIM28 lentiviral vectors were added to white opaque-bottom 384-well plates (ThermoFisher) at 1 × $10^4$ cells per well in 25 µL volume, and treated with compound for 24 h at 37 °C and 5% $CO_2$. The HiBit signal was measured by adding 25 µL of the lytic detection reagent from the Nano-Glo® HiBiT Extracellular Detection System (Promega), prepared as per manufacturer protocol. The CTG signal was developed using 25 µL of CTG reagent (Promega). Both assays were read using PHERAstar microplate luminescence reader (BMG Labtech). The data have been expressed as % of DMSO control and represent three independent biological replicates.

**Proteomics - cell lysis and protein digestion**

The viability of Jurkat cells was tested after 16-h treatment with corresponding chemicals followed by trypan blue staining (CytoSmart cell counter, Corning) and confirmed to be >95 % viable cells in each condition. Next, ~107 cells per condition were collected by centrifugation (1000 rpm), washed twice with ice-cold PBS, and lysed in RIPA buffer [50 mM HEPES pH 7.4, 150 mM NaCl, 1% sodium deoxycholate, 1% NP-40, 0.1% SDS, 10 mM sodium pyrophosphate, 10 mM β-glycerophosphate, 2.5 mM $MgCl_2$, 200 µM TCEP, phosphatase and protease inhibitor cocktail (in-house)], to produce whole cell extracts. Whole-cell extracts were sonicated, and protein concentrations were determined using the Bradford assay. Protein extracts (100 µg) were subjected to disulfide bond reduction with 5 mM TCEP (10 min) and alkylation with 25 mM chloroacetamide (20 min). Methanol–chloroform precipitation was performed prior to protease digestion. In brief, four parts of neat methanol were added to each sample and vortexed, one part chloroform was added to the sample and vortexed, and finally, three parts water was added to the sample and vortexed. The sample was centrifuged at 8000 rpm for 5 min at room temperature and subsequently washed twice with 100% methanol. Samples were resuspended in 100 mM EPPS pH8.5 containing 0.1% RapiGest and digested at 37°C for 6 h with trypsin at a 100:1 protein-to-protease ratio. The digestion efficiency of a small aliquot was tested. Samples were then subjected to TMTpro labeling.

**Proteomics - Tandem mass tag labeling**

Proline-based reporter isobaric Tandem Mass Tag (TMTpro) labeling of dried peptide samples resuspended in 100 mM EPPS pH 8.5, was carried out as follows. TMTpro reagent was added to samples (100 µg peptide), along with acetonitrile, to achieve a final acetonitrile concentration of approximately 30% (*v/v*). Following incubation at room temperature for 1 h, the labeling efficiency of a small aliquot was tested. The reaction was then quenched with hydroxylamine to a final concentration of 0.5% (*v/v*) for 15 min. The TMTpro-labeled samples were pooled together at a 1:1 ratio. The sample was vacuum centrifuged to near dryness and subjected to C18 solid-phase extraction (SPE).

**Proteomics - off-line basic pH reversed-phase (BPRP) fractionation**

The dried TMTpro-labeled sample was resuspended in 100 µL of 10 mM $NH_4HCO_3$ pH 8.0 and fractionated using basic pH reverse phase HPLC (*95*). Briefly, samples were offline fractionated over a 90 min run, into 96 fractions by high pH reverse-phase HPLC (Agilent LC1260) through an aeris peptide xb-c18 column (Phenomenex; 250 mm × 3.6 mm) for total proteome with mobile phase A containing 5% acetonitrile and 10 mM $NH_4HCO_3$, and mobile phase B containing 90% acetonitrile and 10 mM $NH_4HCO_3$ (both pH 8.0). The 96 resulting fractions were then pooled non-continuously into 24 fractions (as outlined in Supplemental Figure 5 of Paulo et al.) (*96*) used for subsequent mass spectrometry analysis. Fractions were vacuum centrifuged to near dryness. Each consolidated fraction was desalted via StageTip, dried again *via* vacuum centrifugation, and reconstituted in 5% acetonitrile, 1% formic acid for LC-MS/MS processing.

**Proteomics – total proteomics analysis using TMTpro**

Mass spectrometry data were collected using an Orbitrap Eclipse Tribrid mass spectrometer (Thermo Fisher Scientific, San Jose, CA) coupled to an UltiMate 3000 RSLCnano system liquid chromatography (LC) pump (Thermo Fisher Scientific). Peptides were separated on a 100 µm inner diameter microcapillary column packed in-house with ~40 cm of HALO Peptide ES-C18 resin (2.7 µm, 160 Å, Advanced Materials Technology, Wilmington, DE) with a gradient consisting of 5%–24% (0-85 min), 24-35% (85-110min) (ACN, 0.1% FA) over a total 125 min run at ~500 nL/min. For analysis, we loaded 1/10 of each fraction onto the column. Each analysis used the Multi-Notch MS3-based TMT method, (*97*) to reduce ion interference compared to MS2 quantification, (*98*) combined with the FAIMS Pro Interface (using previously optimized 3 CV parameters for TMT multiplexed samples) (*99*) and combined with Real-Time Search analysis software (*100*, *101*). The scan sequence began with an MS1 spectrum (Orbitrap analysis; resolution 120,000 at 200 Th; mass range 400−1500 *m/z*; automatic gain control (AGC) target 4×105; maximum injection time 50 ms). Precursors for MS2 analysis were selected using a cycle type of 1.25 sec/CV method (FAIMS CV=-40/-60/-80). MS2 analysis consisted of collision-induced dissociation (quadrupole ion trap analysis; Rapid scan rate; AGC 1.0×104; isolation window 0.5 Th; normalized collision energy (NCE) 35; maximum injection time 35 ms). Monoisotopic peak assignment was used, and previously interrogated precursors were excluded using a dynamic window (120 s ±10 ppm). Following the acquisition of each MS2 spectrum, a synchronous-precursor-selection (SPS) API-MS3 scan was collected on the top 10 most intense ions b or y-ions matched by the online search algorithm in the associated MS2 spectrum (*100*, *101*). MS3 precursors were fragmented by high energy collision-induced dissociation (HCD) and analyzed using the Orbitrap (NCE 45; AGC 2.5×105; maximum injection time 200 ms, resolution was 50000 at 200 Th). The closeout was set at two peptides per protein per fraction so that MS3s were no longer collected for proteins having two peptide-spectrum matches (PSMs) that passed the quality filters (*101*).

**Proteomics – data analysis**

Mass spectra were converted to mzXML (*102*) and processed using the open-source Comet search engine (2020.01 rev. 4) software pipeline (Eng et al., 2013) with the Human Reference Proteome (2020-03 - SwissProt (w isoforms) entries only) UniProt database with contaminants and reverse decoy sequences appended. For analysis, searches were performed with a 50 ppm precursor ion tolerance, and product ion parameters for ion trap MS/MS were used. The theoretical fragment ions were set to 1 with a tolerance of 1.0005 and a 0.4 offset (mono masses). TMTpro tags on lysine residues and peptide N termini (+304.207 Da) and carbamidomethylation of cysteine residues (+57.021 Da) were set as static modifications, while oxidation of methionine residues (+15.995 Da) was set as a variable modification. Search results were first filtered to a 1% peptide FDR using linear discriminant analysis employing a target-decoy strategy and further filtered to obtain a protein level FDR of 1% (*103–105*). Moreover, protein assembly was guided by principles of parsimony to produce the smallest set of proteins necessary to account for all observed peptides.

For TMTpro-based reporter ion quantitation, we extracted the summed signal-to-noise (S:N) ratio for each TMTpro channel and found the closest matching centroid to the expected mass of the TMT reporter ion (integration tolerance of 0.003 Da). Reporter ion intensities were adjusted to correct for the isotopic impurities of the different TMTpro reagents according to manufacturer specifications. Proteins were quantified by summing reporter ion signal-to-noise measurements across all matching PSMs, yielding a ‘‘summed signal-to-noise’’ measurement. PSMs with poor quality, MS3 spectra with 8 or more TMTpro reporter ion channels missing, or isolation specificity less than 0.5, or with TMT reporter summed signal-to-noise ratio that was less than 160 or had no MS3 spectra were excluded from quantification.

Protein quantification values were exported for further analysis in Microsoft Excel, GraphPad Prism, and Perseus (*106*). Each reporter ion channel was summed across all quantified proteins and normalized, assuming equal protein loading of all samples. A two-sided Welch's t-test was conducted on the specified sample comparisons with FDR correction applied for multiple comparisons (Figures 6A, 6B and S18).

Supplemental Data Tables list all quantified proteins and the associated TMTpro reporter ratio to control channels used for quantitative analysis.

**Plasmid Information**

Plasmid: pLVX-Sprout
Backbone: pLVX-EF1a-IRES-mCherry (Takara)
Insert sequence cloned into MCS:
cagtgttctagaggaatagccaccatgtccggattcaatgtcttaatggttcataagcgaagccatactggtgaagagacggacgtctcagaa
aaaccttttaagtgtcacctctgcaactatgcatgccaaagaagagatgcgctcacgcgtgctgaagctgctgcaaaggaagctgcagctaa
ggaggctgcagctaaggctgtgagcaagggcgaggagctgttcaccggggtggtgcccatcctggtcgagctggacggcgacgtaaac
ggccacaagttcagcgtgtccggcgagggcgagggcgatgccacctatggcaaactgaccctgaaattcatctgcaccaccggcaaact
gcccgtgccctggcccaccctcgtgaccaccctgacctatggcgtgcagtgcttcagccgctatcccgaccacatgaaacagcacgacttc
ttcaagtccgccatgcccgaaggctacgtccaggagcgcaccatcttcttcaaggacgacggcaactacaagacccgcgccgaggtgaa
gttcgagggcgacaccctggtgaaccgcatcgagctgaagggcatcgacttcaaggaggacggcaacatcctggggcacaagctggag
tacaactacaacagccacaacgtctatatcatggccgacaagcagaagaacggcatcaaggtgaacttcaagatccgccacaacatcgag
gacggcagcgtgcagctcgccgaccactatcagcagaacacccccatcggcgacggccccgtgctgctgcccgacaaccactatctgag
cacccagtccgccctgagcaaagaccccaacgagaaacgcgatcacatggtcctgctggagttcgtgaccgccgccgggatcactctcg
gcatggatgaactgtataaataaggatccccacat

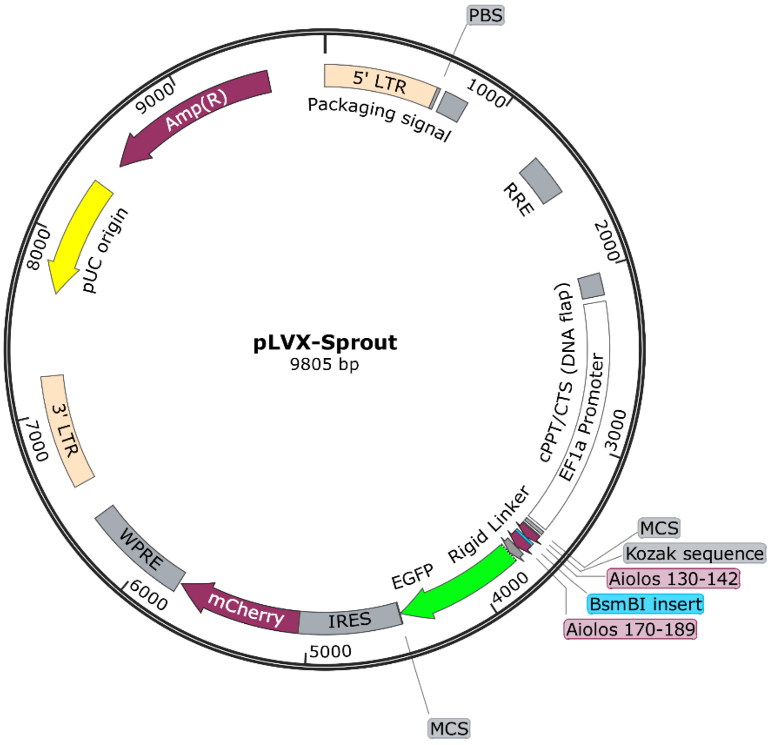

Plasmid: pLVX-Sprout-IKZF3 130-189aa (WT)
Backbone: pLVX-Sprout
Insert sequence cloned into pLVX-Sprout:
cagtgtgggctaccgtctcagtgaacgcccattccagtgtaatcagtgtggggcatcttttactcagaaaggtaacctcctccgccacattaaa
ctgcacacaggggaaacgagacggtagccgccacat

Plasmid map:

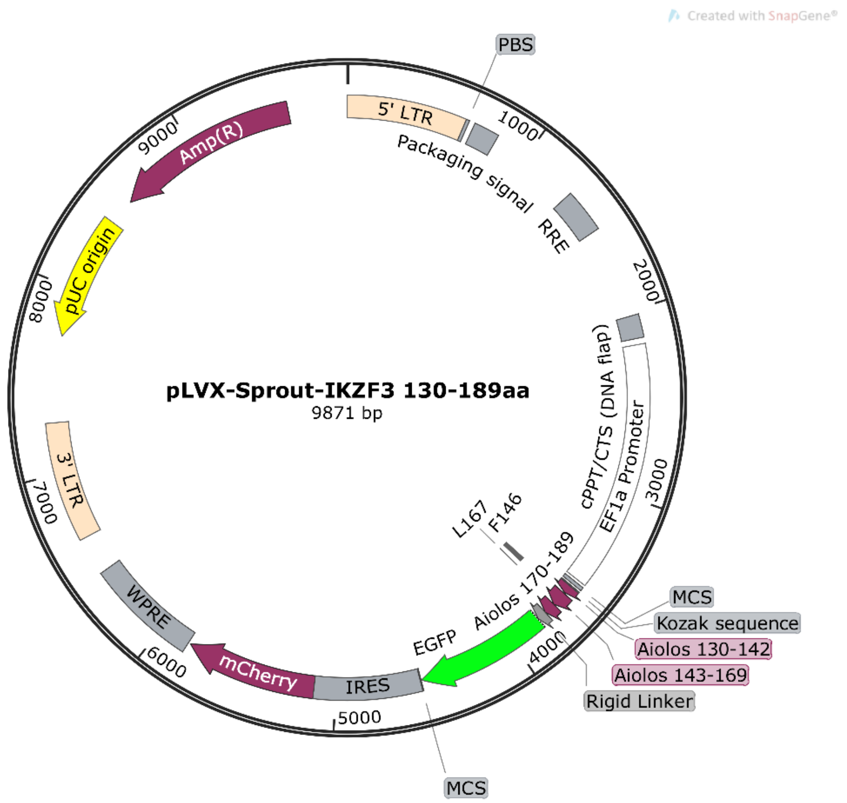

Plasmid: pLVX-Sprout-IKZF3 130-189aa Q147A (ANQS)
Backbone: pLVX-Sprout
Insert sequence cloned into pLVX-Sprout:
cagtgtgggctaccgtctcagtgaacgcccattcgcctgtaatcagtgtggggcatcttttactcagaaaggtaacctcctccgccacattaaa
ctgcacacaggggaaacgagacggtagccgccacat

Plasmid: pLVX-Sprout-IKZF3 130-189aa Q147A N149P Q150I (APIS)
Backbone: pLVX-Sprout
Insert sequence cloned into pLVX-Sprout:
cagtgtgggctaccgtctcagtgaacgcccattcgcctgtccaatctgtggggcatcttttactcagaaaggtaacctcctccgccacattaaa
ctgcacacaggggaaacgagacggtagccgccacat

Plasmid: pLVX-Sprout-IKZF3 130-189aa Q147W (WNQS)
Backbone: pLVX-Sprout
Insert sequence cloned into pLVX-Sprout:
cagtgtgggctaccgtctcagtgaacgcccattctggtgtaatcagtgtggggcatcttttactcagaaaggtaacctcctccgccacattaaa
ctgcacacaggggaaacgagacggtagccgccacat

Plasmid: pLVX-Sprout-IKZF3 130-189aa Q147H N149E Q150I S154P (HEIP)
Backbone: pLVX-Sprout
Insert sequence cloned into pLVX-Sprout:
cagtgtgggctaccgtctcagtgaacgcccattccattgtgagatctgtggggcaccatttactcagaaaggtaacctcctccgccacattaa
actgcacacaggggaaacgagacggtagccgccacat

Plasmid: pLVX-Sprout-IKZF3 130-189aa Q147F N149E Q150V S154P (FEVP)
Backbone: pLVX-Sprout
Insert sequence cloned into pLVX-Sprout:
cagtgtgggctaccgtctcagtgaacgcccattcttctgtgaggtgtgtggggcaccctttactcagaaaggtaacctcctccgccacattaa
actgcacacaggggaaacgagacggtagccgccacat

Plasmid: pLVX-Sprout-IKZF3 130-189aa Q147W N149E Q150I S154K (WEIK)
Backbone: pLVX-Sprout
Insert sequence cloned into pLVX-Sprout:
cagtgtgggctaccgtctcagtgaacgcccattctggtgtgaaatctgtggggcaaagtttactcagaaaggtaacctcctccgccacattaa
actgcacacaggggaaacgagacggtagccgccacat

Plasmid: pLVX-Sprout-IKZF3 130-189aa Q147F N149D Q150I S154P (FDIP)
Backbone: pLVX-Sprout
Insert sequence cloned into pLVX-Sprout:
cagtgtgggctaccgtctcagtgaacgcccattcttctgtgacatatgtggggcaccctttactcagaaaggtaacctcctccgccacattaaa
ctgcacacaggggaaacgagacggtagccgccacat

Plasmid: pLVX-Sprout-IKZF3 130-189aa Q147A N149E Q150V S154K (AEVK)
Backbone: pLVX-Sprout
Insert sequence cloned into pLVX-Sprout:
cagtgtgggctaccgtctcagtgaacgcccattcgcttgtgaagtgtgtggggcaaaatttactcagaaaggtaacctcctccgccacattaa
actgcacacaggggaaacgagacggtagccgccacat

Plasmid: pLVX-Sprout-IKZF3 130-189aa Q147W N149E Q150I (WEIS)
Backbone: pLVX-Sprout
Insert sequence cloned into pLVX-Sprout:
cagtgtgggctaccgtctcagtgaacgcccattctggtgtgagatttgtggggcatcttttactcagaaaggtaacctcctccgccacattaaa
ctgcacacaggggaaacgagacggtagccgccacat

Plasmid: pLVX-Sprout-IKZF3 130-189aa Q147W Q150I S154Q (WNIQ)
Backbone: pLVX-Sprout
Insert sequence cloned into pLVX-Sprout:
cagtgtgggctaccgtctcagtgaacgcccattctggtgtaacatctgtggggcacagtttactcagaaaggtaacctcctccgccacattaa
actgcacacaggggaaacgagacggtagccgccacat

Plasmid: pLVX-Sprout-IKZF3 130-189aa Q147W N149D Q150I S154A (WDIA)
Backbone: pLVX-Sprout
Insert sequence cloned into pLVX-Sprout:
cagtgtgggctaccgtctcagtgaacgcccattctggtgtgatatctgtggggcagcctttactcagaaaggtaacctcctccgccacattaa
actgcacacaggggaaacgagacggtagccgccacat

Plasmid: pLVX-Sprout-IKZF3 130-189aa N149P Q150I (QPIS)
Backbone: pLVX-Sprout
Insert sequence cloned into pLVX-Sprout:
cagtgtgggctaccgtctcagtgaacgcccattccagtgtcctatctgtggggcaagctttactcagaaaggtaacctcctccgccacattaa
actgcacacaggggaaacgagacggtagccgccacat

Plasmid: pLVX-HiBit-TRIM28
Backbone: pLVX-EF1a-IRES-mCherry (Takara)
Insert sequence cloned into MCS:
cagtgttctagaggaatagccaccatggtgagcggctggcggctgttcaagaagattagcggcggaggcggaggtggcgcagcaagtgcagctgctgcaagcgccgctgctgcttccgcagctagtggttcaccgggtccgggagagggatctgctggaggtgagaagcgcagcacagccccaagtgctgctgcgtccgcttcagcttcagctgcagcaagtagcccagcgggggggtggcgcagaggctctcgaactgttggaacattgtggcgtctgtagagagaggcttcgcccggagagggagcctaggctccttccgtgtcttcattctgcctgcagcgcatgtttgggcccagcggcaccggcggctgcaaattcatccggggatgggggcgcagcaggtgacggtaccgtggtagactgtccagtatgcaagcagcaatgtttttcaaaagacatagttgagaactattttatgagagatagcggtagcaaggctgcaacagacgcgcaagatgcgaatcaatgctgtacctcatgtgaagataatgctccggccacgagttactgtgtggagtgttcagaaccgctttgtgaaacttgcgtcgaggcgcaccaaagggtcaaatacactaaagatcataccgtaaggtcaacaggaccagcgaaatcccgcgacggagaacgaacggtctactgcaacgtccataagcatgagccgttggtccttttctgcgagtcttgcgatacgctcacctgtagggattgtcaactgaacgcgcataaggatcatcaatatcagtttctggaagatgctgtccggaatcagcgaaaacttctcgcaagcctcgtgaaaaggctcggcgacaaacatgcgactctccagaagtcaaccaaggaagtccggagcagcatacgacaagtgtctgatgttcaaaagcgagtgcaagtagatgtcaagatggccatactccaaataatgaaggagctcaataaacggggtcgagtccttgtcaacgatgcacagaaggtcacggagggggcaacaggaaaggctggaacgacaacactggactatgaccaagattcaaaagcaccaagagcacatacttcgctttgcatcatgggcgttggagtctgataacaatacggcactgctcctgtctaagaaattgatctattttcagctgcacagggctctgaaaatgatagtcgatccagttgaaccccacggggagatgaaatttcagtgggacctgaatgcatggacaaagtccgctgaagcattcggaaagattgtagctgaaaggcccgggactaactcaacaggacccgctcctatggctcctccacgcgcacctggtcctctgtcaaaacaggggagtggctcatcacaacccatggaagtgcaagagggttatggttttggaagcggagatgatccctattctagcgcggaacctcacgtatcaggggtcaaacgatcccgctcaggtgagggcgaggtcagtgggctgatgcgaaaggtaccccgcgtcagtctggagcggttggatcttgatcttaccgcggatagtcaacctcccgttttcaaagtttttccgggttcaactacggaagattataacctcatagtaatcgagagaggagctgcagccgcggctacaggccaacctgggacagcaccagcaggtacacccggggcccctccacttgcggggatggccattgttaaagaagaagagacggaagcagcgattggggctcctccaaccgccaccgaaggacccgaaacaaaaccagttctcatggcactggctgaaggccctggggctgaagggccgagattggccagtccgtcagggagcacttcatctgggttggaagtcgtggcccctgagggcacaagtgcacccgggggggggcccaggcactctcgatgactctgctacgatctgtcgggtatgtcaaaaaccaggcgaccttgttatgtgcaatcagtgcgaattctgcttccacctggactgtcaccttccggccctgcaggacgttccaggtgaggaatggagctgtagcctgtgtcacgtactcccggatctgaaggaggaagatggctcactgtcactggacggtgcagattccaccggagttgtggccaagttgtccccagcaaatcagagaaagtgcgaaagggtcctgcttgcgttgttctgccacgaaccgtgtcggcctctgcaccagttggcgacggactccacgttctctctggatcagcccggtgggaccttggatcttactctcatacgggctcgactccaagagaagctttcacctccttatagttccccccaggagttcgctcaagacgtaggcaggatgttcaagcaatttaataagttgactgaggacaaagcggatgttcaaagcattatcggattgcaaagattctttgagacacgaatgaatgaagcattcggggacaccaaatttagcgcggtacttgtagagcccccctcctatgtccttgccaggtgcaggtttgtcctcccaggaattgtctgggggaccgggtgatggaccctgaggatccccacat

Plasmid map:

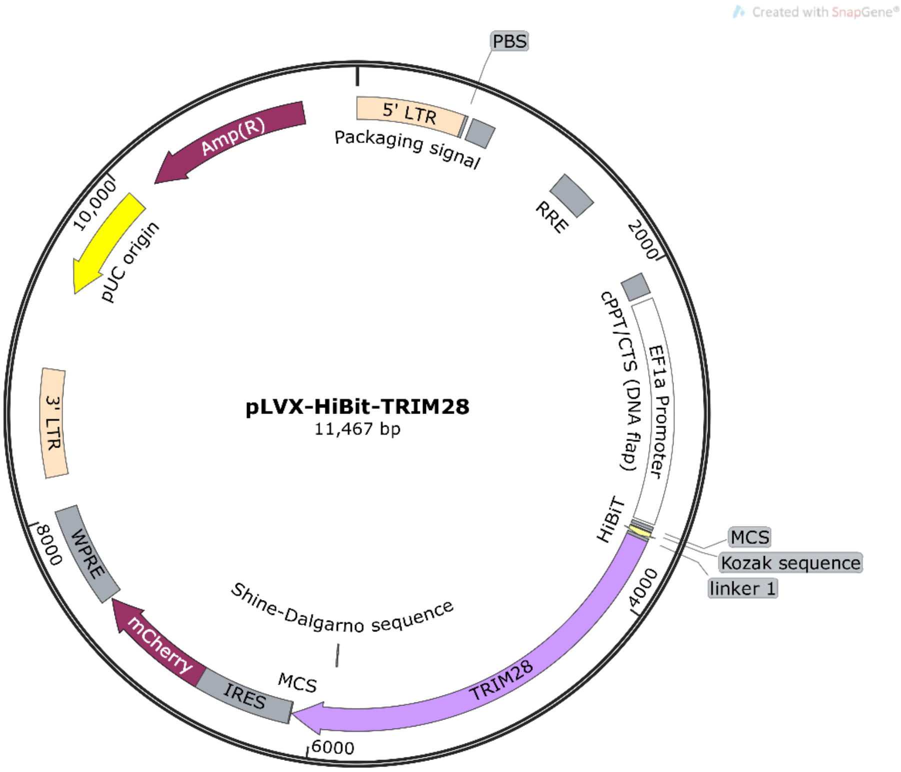

Plasmid: pLVX-HiBit-TRIM28 N-degron
Backbone: pLVX-EF1a-IRES-mCherry (Takara)
Insert sequence cloned into MCS:
cagtgttctagaggaatagccaccatggtgagcggctggcggctgttcaagaagattagcggcggaggcggaggtggctccggattcaatgtcttaatggttcataagcgaagccatactggtgaacgcccattctggtgtgagatttgtggggcatcttttactcagaaaggtaacctcctccgccacattaaactgcacacaggggaaaaaccttttaagtgtcacctctgcaactatgcatgccaaagaagagatgcgctcacgcgtgctgaagccgctgcaaaggaagccgcagctaaggaggcagctgccaaggccgcagcaagtgcagctgctgcaagcgccgctgctgcttccgcagctagtggttcaccgggtccgggagagggatctgctggaggtgagaagcgcagcacagccccaagtgctgctgcgtccgcttcagcttcagctgcagcaagtagcccagcgggggggtggcgcagaggctctcgaactgttggaacattgtggcgtctgtagagagaggcttcgcccggagagggagcctaggctccttccgtgtcttcattctgcctgcagcgcatgtttgggcccagcggcaccggcggctgcaaattcatccggggatgggggcgcagcaggtgacggtaccgtggtagactgtccagtatgcaagcagcaatgtttttcaaaagacatagttgagaactattttatgagagatagcggtagcaaggctgcaacagacgcgcaagatgcgaatcaatgctgtacctcatgtgaagataatgctccggccacgagttactgtgtggagtgttcagaaccgctttgtgaaacttgcgtcgaggcgcaccaaagggtcaaatacactaaagatcataccgtaaggtcaacaggaccagcgaaatcccgcgacggagaacgaacggtctactgcaacgtccataagcatgagccgttggtccttttctgcgagtcttgcgatacgctcacctgtagggattgtcaactgaacgcgcataaggatcatcaatatcagtttctggaagatgctgtccggaatcagcgaaaacttctcgcaagcctcgtgaaaaggctcggcgacaaacatgcgactctccagaagtcaaccaaggaagtccggagcagcatacgacaagtgtctgatgttcaaaagcgagtgcaagtagatgtcaagatggccatactccaaataatgaaggagctcaataaacggggtcgagtccttgtcaacgatgcacagaaggtcacggaggggcaacaggaaaggctggaacgacaacactggactatgaccaagattcaaaagcaccaagagcacatacttcgctttgcatcatgggcgttggagtctgataacaatacggcactgctcctgtctaagaaattgatctattttcagctgcacagggctctgaaaatgatagtcgatccagttgaaccccacggggagatgaaatttcagtgggacctgaatgcatggacaaagtccgctgaagcattcggaaagattgtagctgaaaggcccgggactaactcaacaggacccgctcctatggctcctccacgcgcacctggtcctctgtcaaaacaggggagtggctcatcacaacccatggaagtgcaagagggttatggttttggaagcggagatgatccctattctagcgcggaacctcacgtatcaggggtcaaacgatcccgctcaggtgagggcgaggtcagtgggctgatgcgaaaggtaccccgcgtcagtctggagcggttggatcttgatcttaccgcggatagt

caacctcccgttttcaaagtttttccgggttcaactacggaagattataacctcatagtaatcgagagaggagctgcagccgcggctacaggc
caacctgggacagcaccagcaggtacacccggggcccctccacttgcggggatggccattgttaaagaagaagagacggaagcagcga
ttggggctcctccaaccgccaccgaaggacccgaaacaaaaccagttctcatggcactggctgaaggccctggggctgaagggccgag
attggccagtccgtcagggagcacttcatctgggttggaagtcgtggcccctgagggcacaagtgcacccgggggggcccaggcactc
tcgatgactctgctacgatctgtcgggtatgtcaaaaaccaggcgaccttgttatgtgcaatcagtgcgaattctgcttccacctggactgtcac
cttccggccctgcaggacgttccaggtgaggaatggagctgtagcctgtgtcacgtactcccggatctgaaggaggaagatggctcactgt
cactggacggtgcagattccaccggagttgtggccaagttgtccccagcaaatcagagaaagtgcgaaagggtcctgcttgcgttgttctgc
cacgaaccgtgtcggcctctgcaccagttggcgacggactccacgttctctctggatcagcccggtgggaccttggatcttactctcatacg
ggctcgactccaagagaagctttcacctccttatagttccccccaggagttcgctcaagacgtaggcaggatgttcaagcaatttaataagttg
actgaggacaaagcggatgttcaaagcattatcggattgcaaagattctttgagacacgaatgaatgaagcattcggggacaccaaatttag
cgcggtacttgtagagccccctcctatgtccttgccaggtgcaggtttgtcctcccaggaattgtctgggggaccgggtgatggaccctgag
gatccccacat

Plasmid map:

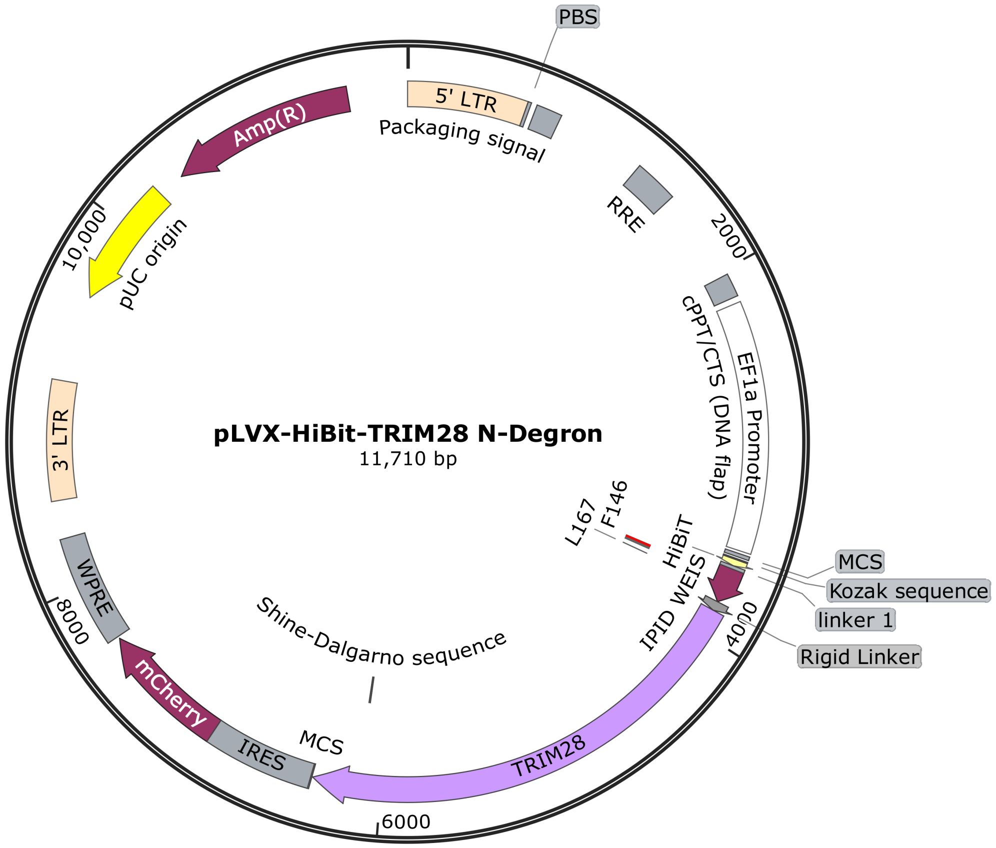

Plasmid: pLVX-HiBit-TRIM28 C-degron
Backbone: pLVX-EF1a-IRES-mCherry (Takara)
Insert sequence cloned into MCS:
cagtgttctagaggaatagccaccatggtgagcggctggcggctgttcaagaagattagcggcggaggcggaggtggagcagcaagtg
cagctgctgcaagcgccgctgctgcttccgcagctagtggttcaccgggtccgggagagggatctgctggaggtgagaagcgcagcaca
gcccccaagtgctgctgcgtccgcttcagcttcagctgcagcaagtagcccagcggggggtggcgcagaggctctcgaactgttggaacat
tgtggcgtctgtagagagaggcttcgcccggagagggagcctaggctccttccgtgtcttcattctgcctgcagcgcatgtttgggcccagc

ggcaccggcggctgcaaattcatccggggatgggggcgcagcaggtgacggtaccgtggtagactgtccagtatgcaagcagcaatgttt
ttcaaaagacatagttgagaactattttatgagagatagcggtagcaaggctgcaacagacgcgcaagatgcgaatcaatgctgtacctcat
gtgaagataatgctccggccacgagttactgtgtggagtgttcagaaccgctttgtgaaacttgcgtcgaggcgcaccaaagggtcaaatac
actaaagatcataccgtaaggtcaacaggaccagcgaaatcccgcgacggagaacgaacggtctactgcaacgtccataagcatgagcc
gttggtccttttctgcgagtcttgcgatacgctcacctgtagggattgtcaactgaacgcgcataaggatcatcaatatcagtttctggaagatg
ctgtccggaatcagcgaaaacttctcgcaagcctcgtgaaaaggctcggcgacaaacatgcgactctccagaagtcaaccaaggaagtcc
ggagcagcatacgacaagtgtctgatgttcaaaagcgagtgcaagtagatgtcaagatggccatactccaaataatgaaggagctcaataa
acggggtcgagtccttgtcaacgatgcacagaaggtcacggaggggcaacaggaaaggctggaacgacaacactggactatgaccaag
attcaaaagcaccaagagcacatacttcgctttgcatcatgggcgttggagtctgataacaatacggcactgctcctgtctaagaaattgatct
attttcagctgcacagggctctgaaaatgatagtcgatccagttgaaccccacggggagatgaaatttcagtgggacctgaatgcatggaca
aagtccgctgaagcattcggaaagattgtagctgaaaggcccgggactaactcaacaggacccgctcctatggctcctccacgcgcacct
ggtcctctgtcaaaacaggggagtggctcatcacaacccatggaagtgcaagagggttatggttttggaagcggagatgatccctattctag
cgcggaacctcacgtatcaggggtcaaacgatcccgctcaggtgagggcgaggtcagtgggctgatgcgaaaggtaccccgcgtcagtc
tggagcggttggatcttgatcttaccgcggatagtcaacctcccgttttcaaagtttttccgggttcaactacggaagattataacctcatagtaa
tcgagagaggagctgcagccgcggctacaggccaacctgggacagcaccagcaggtacacccggggcccctccacttgcggggatgg
ccattgttaaagaagaagagacggaagcagcgattggggctcctccaaccgccaccgaaggacccgaaacaaaaccagttctcatggca
ctggctgaaggccctggggctgaagggccgagattggccagtccgtcagggagcacttcatctgggttggaagtcgtggcccctgaggg
cacaagtgcacccgggggggggcccaggcactctcgatgactctgctacgatctgtcgggtatgtcaaaaaccaggcgaccttgttatgtgc
aatcagtgcgaattctgcttccacctggactgtcaccttccggccctgcaggacgttccaggtgaggaatggagctgtagcctgtgtcacgta
ctcccggatctgaaggaggaagatggctcactgtcactggacggtgcagattccaccggagttgtggccaagttgtccccagcaaatcaga
gaaagtgcgaaagggtcctgcttgcgttgttctgccacgaaccgtgtcggcctctgcaccagttggcgacggactccacgttctctctggatc
agcccggtgggaccttggatcttactctcatacgggctcgactccaagagaagctttcacctccttatagttccccccaggagttcgctcaag
acgtaggcaggatgttcaagcaatttaataagttgactgaggacaaagcggatgttcaaagcattatcggattgcaaagattctttgagacac
gaatgaatgaagcattcggggacaccaaatttagcgcggtacttgtagagccccctcctatgtccttgccaggtgcaggtttgtcctcccagg
aattgtctgggggaccgggtgatggacccgctgaagccgctgcaaaggaagccgcagctaaggaggcagctgccaaggcctccggatt
caatgtcttaatggttcataagcgaagccatactggtgaacgcccattctggtgtgagatttgtggggcatcttttactcagaaaggtaacctcc
tccgccacattaaactgcacacaggggaaaaaccttttaagtgtcacctctgcaactatgcatgccaaagaagagatgcgctcacgcgttga
ggatccccacat

Plasmid map:

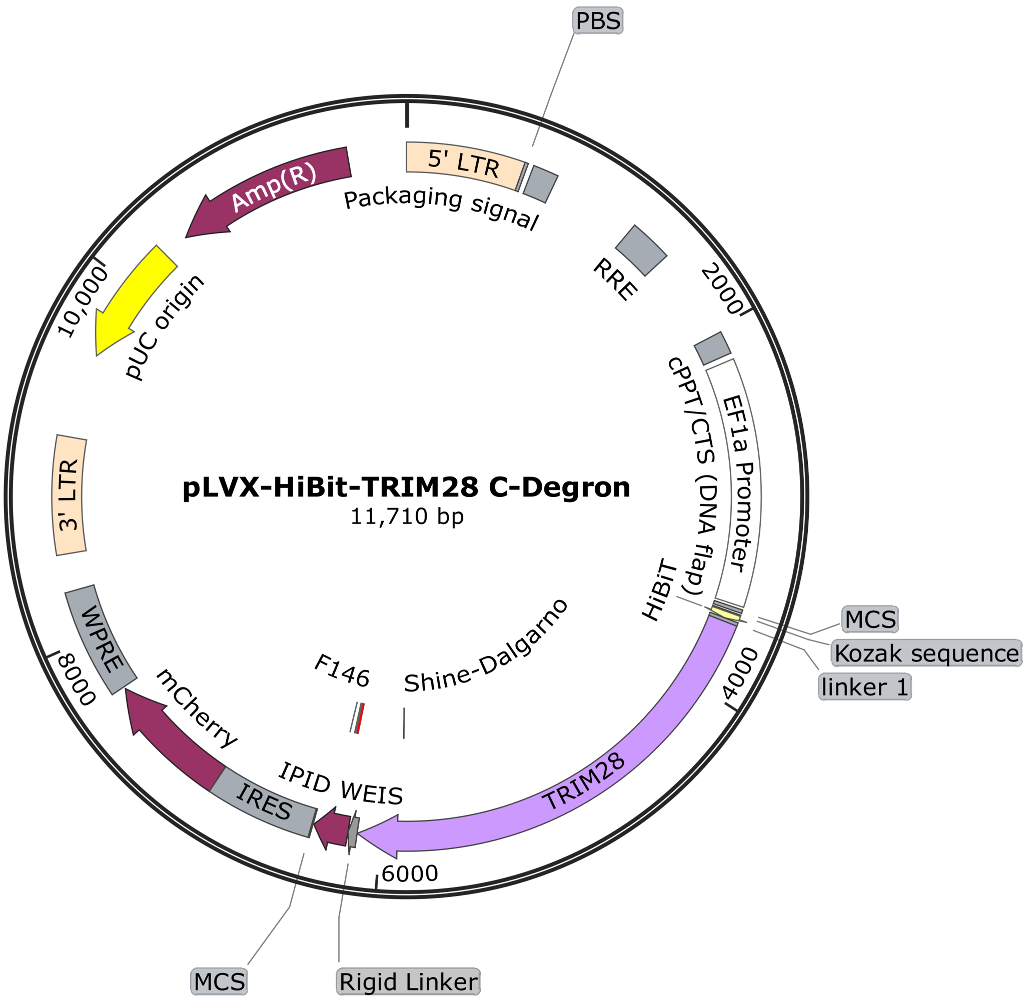

5

**Sequences for NGS**

NGS PCR1 primers

| Name | Sequence |
|---|---|
| PCR1 F | CAGGTGTCGTGAGGATCTATTTCCG |
| PCR1 R | GTGCAGATGAATTTCAGGGTCAGG |

### NGS PCR2 P5 primers

| Sample | Name | Sequence |
|---|---|---|
| Replicate 1 | N502 index P5 0 nt stagger | AATGATACGGCGACCACCGAGATCTACACCTCTCTATACACTCTTTCCCTACACGACGCTCTTCCGATCTAGCGAAGCCATACTGGTGAA*C |
| Replicate 1 | N502 index P5 1 nt stagger | AATGATACGGCGACCACCGAGATCTACACCTCTCTATACACTCTTTCCCTACACGACGCTCTTCCGATCTCAGCGAAGCCATACTGGTGAA*C |
| Replicate 1 | N502 index P5 2 nt stagger | AATGATACGGCGACCACCGAGATCTACACCTCTCTATACACTCTTTCCCTACACGACGCTCTTCCGATCTGCAGCGAAGCCATACTGGTGAA*C |
| Replicate 1 | N502 index P5 3 nt stagger | AATGATACGGCGACCACCGAGATCTACACCTCTCTATACACTCTTTCCCTACACGACGCTCTTCCGATCTAGCAGCGAAGCCATACTGGTGAA*C |
| Replicate 1 | N502 index P5 4 nt stagger | AATGATACGGCGACCACCGAGATCTACACCTCTCTATACACTCTTTCCCTACACGACGCTCTTCCGATCTCAACAGCGAAGCCATACTGGTGAA*C |
| Replicate 1 | N502 index P5 6 nt stagger | AATGATACGGCGACCACCGAGATCTACACCTCTCTATACACTCTTTCCCTACACGACGCTCTTCCGATCTTGCACCAGCGAAGCCATACTGGTGAA*C |
| Replicate 1 | N502 index P5 7 nt stagger | AATGATACGGCGACCACCGAGATCTACACCTCTCTATACACTCTTTCCCTACACGACGCTCTTCCGATCTACGCAACAGCGAAGCCATACTGGTGAA*C |
| Replicate 1 | N502 index P5 8 nt stagger | AATGATACGGCGACCACCGAGATCTACACCTCTCTATACACTCTTTCCCTACACGACGCTCTTCCGATCTGAAGACCCAGCGAAGCCATACTGGTGAA*C |
| Replicate 2 | N505 index P5 0 nt stagger | AATGATACGGCGACCACCGAGATCTACACGTAAGGAGACACTCTTTCCCTACACGACGCTCTTCCGATCTAGCGAAGCCATACTGGTGAA*C |
| Replicate 2 | N505 index P5 1 nt stagger | AATGATACGGCGACCACCGAGATCTACACGTAAGGAGACACTCTTTCCCTACACGACGCTCTTCCGATCTCAGCGAAGCCATACTGGTGAA*C |
| Replicate 2 | N505 index P5 2 nt stagger | AATGATACGGCGACCACCGAGATCTACACGTAAGGAGACACTCTTTCCCTACACGACGCTCTTCCGATCTGCAGCGAAGCCATACTGGTGAA*C |
| Replicate 2 | N505 index P5 3 nt stagger | AATGATACGGCGACCACCGAGATCTACACGTAAGGAGACACTCTTTCCCTACACGACGCTCTTCCGATCTAGCAGCGAAGCCATACTGGTGAA*C |
| Replicate 2 | N505 index P5 4 nt stagger | AATGATACGGCGACCACCGAGATCTACACGTAAGGAGACACTCTTTCCCTACACGACGCTCTTCCGATCTCAACAGCGAAGCCATACTGGTGAA*C |
| Replicate 2 | N505 index P5 6 nt stagger | AATGATACGGCGACCACCGAGATCTACACGTAAGGAGACACTCTTTCCCTACACGACGCTCTTCCGATCTTGCACCAGCGAAGCCATACTGGTGAA*C |
| Replicate 2 | N505 index P5 7 nt stagger | AATGATACGGCGACCACCGAGATCTACACGTAAGGAGACACTCTTTCCCTACACGACGCTCTTCCGATCTACGCAACAGCGAAGCCATACTGGTGAA*C |
| Replicate 2 | N505 index P5 8 nt stagger | AATGATACGGCGACCACCGAGATCTACACGTAAGGAGACACTCTTTCCCTACACGACGCTCTTCCGATCTGAAGACCCAGCGAAGCCATACTGGTGAA*C |
| Replicate 3 | N506 index P5 0 nt stagger | AATGATACGGCGACCACCGAGATCTACACACTGCATAACACTCTTTCCCTACACGACGCTCTTCCGATCTAGCGAAGCCATACTGGTGAA*C |
| Replicate 3 | N506 index P5 1 nt stagger | AATGATACGGCGACCACCGAGATCTACACACTGCATAACACTCTTTCCCTACACGACGCTCTTCCGATCTCAGCGAAGCCATACTGGTGAA*C |
| Replicate 3 | N506 index P5 2 nt stagger | AATGATACGGCGACCACCGAGATCTACACACTGCATAACACTCTTTCCCTACACGACGCTCTTCCGATCTGCAGCGAAGCCATACTGGTGAA*C |
| Replicate 3 | N506 index P5 3 nt stagger | AATGATACGGCGACCACCGAGATCTACACACTGCATAACACTCTTTCCCTACACGACGCTCTTCCGATCTAGCAGCGAAGCCATACTGGTGAA*C |
| Replicate 3 | N506 index P5 4 nt stagger | AATGATACGGCGACCACCGAGATCTACACACTGCATAACACTCTTTCCCTACACGACGCTCTTCCGATCTCAACAGCGAAGCCATACTGGTGAA*C |
| Replicate 3 | N506 index P5 6 nt stagger | AATGATACGGCGACCACCGAGATCTACACACTGCATAACACTCTTTCCCTACACGACGCTCTTCCGATCTTGCACCAGCGAAGCCATACTGGTGAA*C |
| Replicate 3 | N506 index P5 7 nt stagger | AATGATACGGCGACCACCGAGATCTACACACTGCATAACACTCTTTCCCTACACGACGCTCTTCCGATCTACGCAACAGCGAAGCCATACTGGTGAA*C |
| Replicate 3 | N506 index P5 8 nt stagger | AATGATACGGCGACCACCGAGATCTACACACTGCATAACACTCTTTCCCTACACGACGCTCTTCCGATCTGAAGACCCAGCGAAGCCATACTGGTGAA*C |

| |
|---|
| P5 flowcell attachment sequence |
| Illumina sequencing primer |
| Vector primer binding sequence |
| Barcode region |
| Stagger region |
| * = Phosphorothioate bond |

NGS PCR2 P7 primers

| Compound | Name | Sequence |
|---|---|---|
| DMSO -ve | N703 index P7.3 | CAAGCAGAAGACGGCATACGAGATTTCTGCCTGTGACTGGAGTTCAGACGTGTGCTCTTCCGATCTGTGACACTTAAAAGGTTTTTCCCCT*G |
| Thal -ve | N704 index P7.3 | CAAGCAGAAGACGGCATACGAGATGCTCAGGAGTGACTGGAGTTCAGACGTGTGCTCTTCCGATCTGTGACACTTAAAAGGTTTTTCCCCT*G |
| 19 -ve | N705 index P7.3 | CAAGCAGAAGACGGCATACGAGATAGGAGTCCGTGACTGGAGTTCAGACGTGTGCTCTTCCGATCTGTGACACTTAAAAGGTTTTTCCCCT*G |
| 23 -ve | N706 index P7.3 | CAAGCAGAAGACGGCATACGAGATCATGCCTAGTGACTGGAGTTCAGACGTGTGCTCTTCCGATCTGTGACACTTAAAAGGTTTTTCCCCT*G |
| Len -ve | N707 index P7.3 | CAAGCAGAAGACGGCATACGAGATGTAGAGAGGTGACTGGAGTTCAGACGTGTGCTCTTCCGATCTGTGACACTTAAAAGGTTTTTCCCCT*G |
| 33 -ve | N710 index P7.3 | CAAGCAGAAGACGGCATACGAGATCAGCCTCGGTGACTGGAGTTCAGACGTGTGCTCTTCCGATCTGTGACACTTAAAAGGTTTTTCCCCT*G |
| 22 -ve | N711 index P7.3 | CAAGCAGAAGACGGCATACGAGATTGCCTCTTGTGACTGGAGTTCAGACGTGTGCTCTTCCGATCTGTGACACTTAAAAGGTTTTTCCCCT*G |
| DMSO +ve | N714 index P7.3 | CAAGCAGAAGACGGCATACGAGATTCATGAGCGTGACTGGAGTTCAGACGTGTGCTCTTCCGATCTGTGACACTTAAAAGGTTTTTCCCCT*G |
| Thal +ve | N715 index P7.3 | CAAGCAGAAGACGGCATACGAGATCCTGAGATGTGACTGGAGTTCAGACGTGTGCTCTTCCGATCTGTGACACTTAAAAGGTTTTTCCCCT*G |
| 19 +ve | N716 index P7.3 | CAAGCAGAAGACGGCATACGAGATTAGCGAGTGTGACTGGAGTTCAGACGTGTGCTCTTCCGATCTGTGACACTTAAAAGGTTTTTCCCCT*G |
| 23 +ve | N718 index P7.3 | CAAGCAGAAGACGGCATACGAGATGTAGCTCCGTGACTGGAGTTCAGACGTGTGCTCTTCCGATCTGTGACACTTAAAAGGTTTTTCCCCT*G |
| Len +ve | N719 index P7.3 | CAAGCAGAAGACGGCATACGAGATTACTACGCGTGACTGGAGTTCAGACGTGTGCTCTTCCGATCTGTGACACTTAAAAGGTTTTTCCCCT*G |
| 33 +ve | N720 index P7.3 | CAAGCAGAAGACGGCATACGAGATAGGCTCCGGTGACTGGAGTTCAGACGTGTGCTCTTCCGATCTGTGACACTTAAAAGGTTTTTCCCCT*G |
| 22 +ve | N721 index P7.3 | CAAGCAGAAGACGGCATACGAGATGCAGCGTAGTGACTGGAGTTCAGACGTGTGCTCTTCCGATCTGTGACACTTAAAAGGTTTTTCCCCT*G |

| |
|---|
| P7 flowcell attachment sequence |
| Illumina sequencing primer |
| Vector primer binding sequence |
| Barcode region |
| * = Phosphorothioate bond |

#### sgRNA

| sgRNA | Sequence |
|---|---|
| TRIM28 sgRNA 1 | AUCCCUGCUUCUCGAAGUGG |
| TRIM28 sgRNA 2 | GUGCUUCUCCAAAGACAUCG |
| TRIM28 sgRNA 3 | CGACGCCCAGGAUGCGAACC |

#### PCR primers for amplifying sgRNA target site

| Primer | Sequence |
|---|---|
| TRIM28 F | CGAAGTGATCGGTGCCAC |
| TRIM28 R | GCATTATCCTCACAGCTAGTGC |

Computational Methods

Docking was carried out using the crystal structure DDB1-CRBN-pomalidomide complex bound to IKZF1(ZF2) from the Protein Data Bank (PDB ID: 6H0F) as a template (*6*).

Protein energetics and mutation evaluation were carried out using the PositionScan function in FoldX 5. The 3D coordinate files generated for the mutant proteins were then used for subsequent docking.

Docking was carried out using GOLD version 2020.1 from the Cambridge Crystallographic Data Centre. The position of the original pomalidomide ligand in the template crystal structure was used to define the binding pocket for new ligands using GOLD’s ‘cavity’ functionality. The imide ring region of the original pomalidomide ligand was used to specify the position onto which the imide region of screened ligands should be superimposed using GOLD’s ‘scaffold’ functionality. All parameters were kept as default during docking screens, except constraint weight, which was increased from the default value of 5 to 10.

Mutational analysis for use in the lentiviral library screen was performed using the PositionScan functionality in FoldX and the PositionMutation functionality in Rosetta. The same crystal structure (PDB ID: 6H0F) was used with the pomalidomide ligand removed (*6*).

NMR Spectra for Compounds

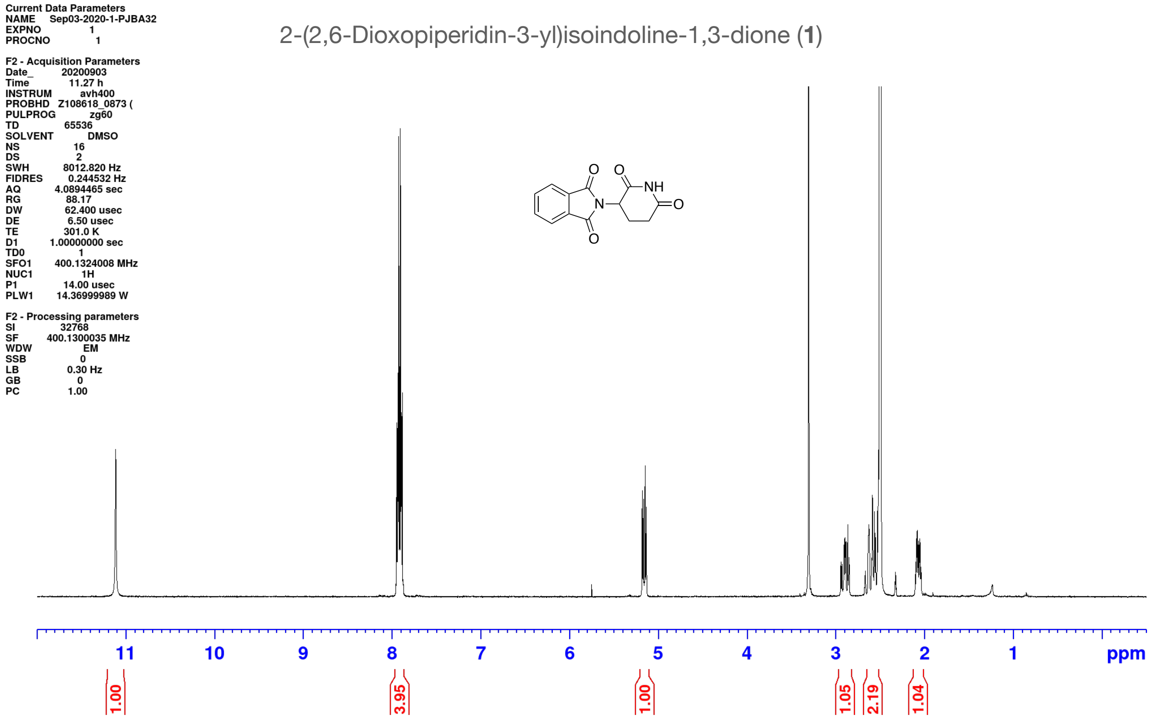

2-(2,6-Dioxopiperidin-3-yl)isoindoline-1,3-dione (**1**)

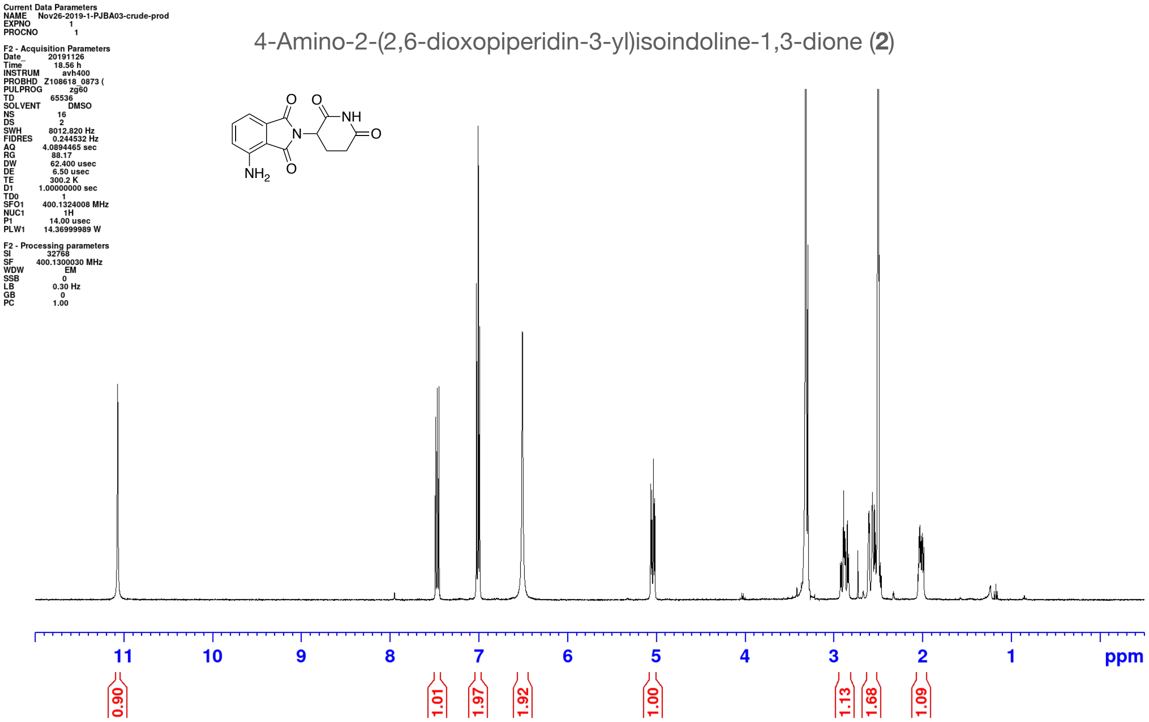
Current Data Parameters
NAME Nov26-2019-1-PJBA03-crude-prod
EXPNO 1
PROCNO 1
F2 - Acquisition Parameters
Date_ 20191126
Time 18.56 h
INSTRUM avh400
PROBHD Z108618_0873 (
PULPROG zg60
TD 65536
SOLVENT DMSO
NS 16
DS 2
SWH 8012.820 Hz
FIDRES 0.244532 Hz
AQ 4.0894465 sec
RG 88.17
DW 62.400 usec
DE 6.50 usec
TE 300.2 K
D1 1.00000000 sec
TD0 1
SFO1 400.1324008 MHz
NUC1 1H
P1 14.00 usec
PLW1 14.36999989 W
F2 - Processing parameters
SI 32768
SF 400.1300030 MHz
WDW EM
SSB 0
LB 0.30 Hz
GB 0
PC 1.00
4-Amino-2-(2,6-dioxopiperidin-3-yl)isoindoline-1,3-dione (2)
O
O
NH
N
O
$NH_2$
O
11
10
9
8
7
6
5
4
3
2
1
ppm
0.90
1.01
1.97
1.92
1.00
1.13
1.68
1.09

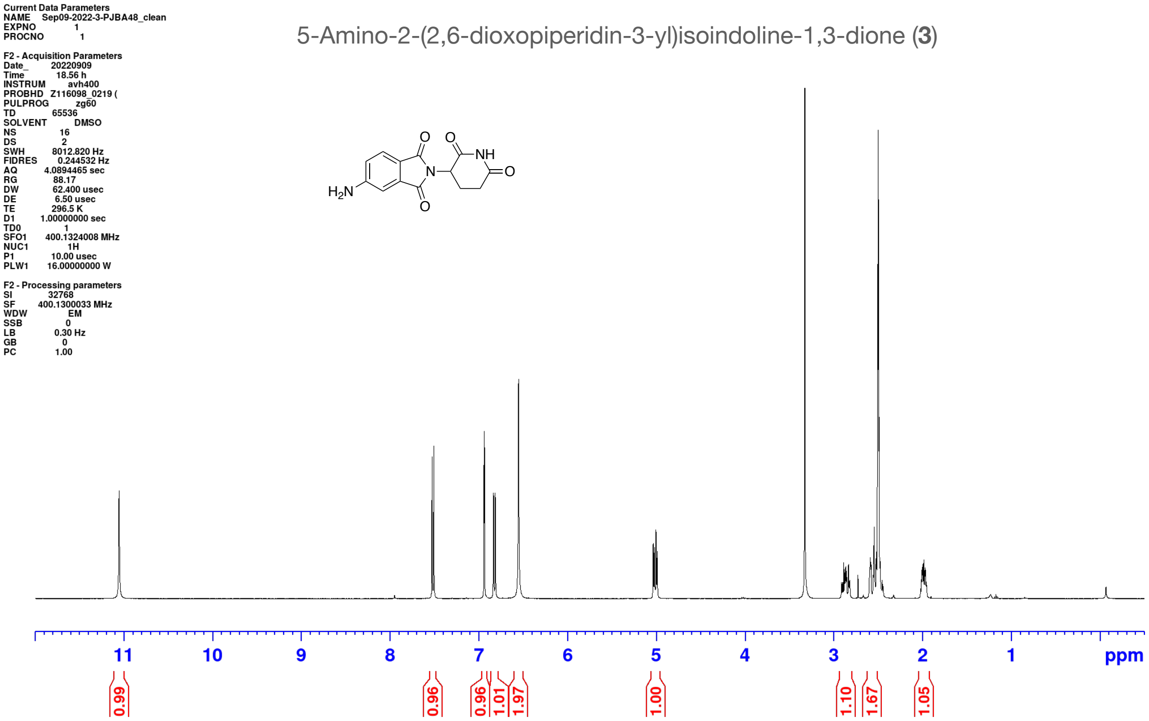
Current Data Parameters
NAME Sep09-2022-3-PJBA48_clean
EXPNO 1
PROCNO 1
F2 - Acquisition Parameters
Date_ 20220909
Time 18.56 h
INSTRUM avh400
PROBHD Z116098_0219 (
PULPROG zg60
TD 65536
SOLVENT DMSO
NS 16
DS 2
SWH 8012.820 Hz
FIDRES 0.244532 Hz
AQ 4.0894465 sec
RG 88.17
DW 62.400 usec
DE 6.50 usec
TE 296.5 K
D1 1.00000000 sec
TD0 1
SFO1 400.1324008 MHz
NUC1 1H
P1 10.00 usec
PLW1 16.00000000 W
F2 - Processing parameters
SI 32768
SF 400.1300033 MHz
WDW EM
SSB 0
LB 0.30 Hz
GB 0
PC 1.00
5-Amino-2-(2,6-dioxopiperidin-3-yl)isoindoline-1,3-dione (3)
O
O
NH
N
O
$H_2N$
O
11
10
9
8
7
6
5
4
3
2
1
ppm
0.99
0.96
0.96
1.01
1.97
1.00
1.10
1.67
1.05

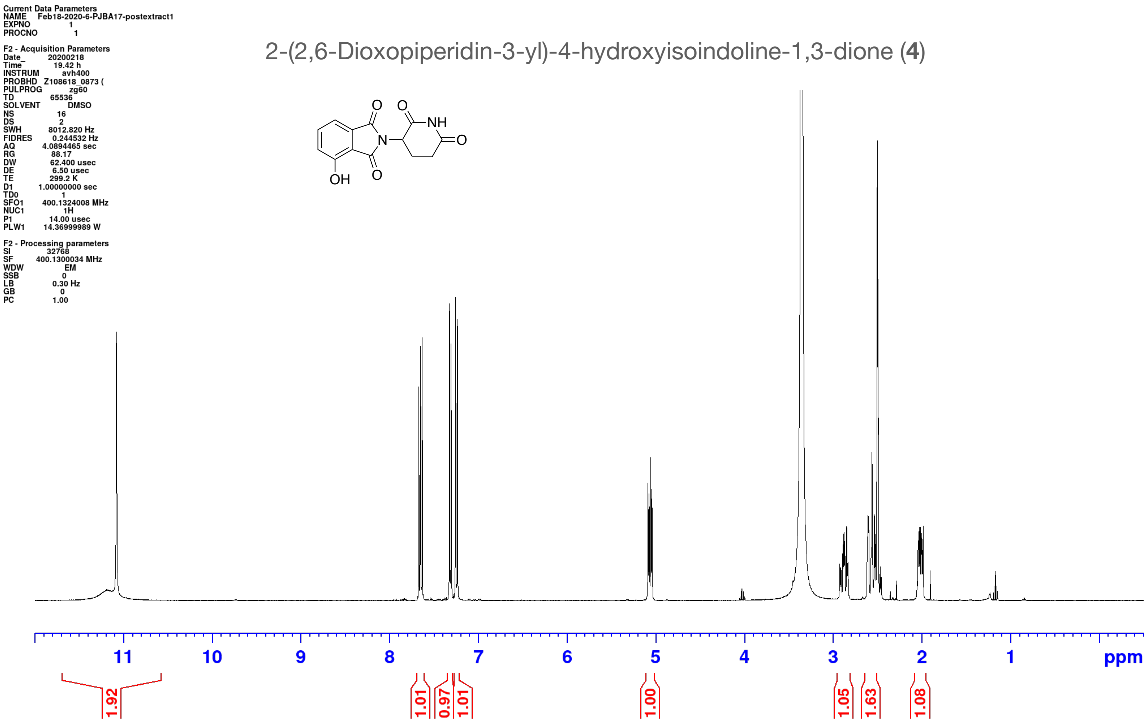
Current Data Parameters
NAME Feb18-2020-6-PJBA17-postextract1
EXPNO 1
PROCNO 1
F2 - Acquisition Parameters
Date_ 20200218
Time 19.42 h
INSTRUM avh400
PROBHD Z108618_0873 (
PULPROG zg60
TD 65536
SOLVENT DMSO
NS 16
DS 2
SWH 8012.820 Hz
FIDRES 0.244532 Hz
AQ 4.0894465 sec
RG 88.17
DW 62.400 usec
DE 6.50 usec
TE 299.2 K
D1 1.00000000 sec
TD0 1
SFO1 400.1324008 MHz
NUC1 1H
P1 14.00 usec
PLW1 14.36999989 W
F2 - Processing parameters
SI 32768
SF 400.1300034 MHz
WDW EM
SSB 0
LB 0.30 Hz
GB 0
PC 1.00
2-(2,6-Dioxopiperidin-3-yl)-4-hydroxyisoindoline-1,3-dione (4)
O
O
NH
N
O
O
OH
11
10
9
8
7
6
5
4
3
2
1
ppm
1.92
1.01
0.97
1.01
1.00
1.05
1.63
1.08

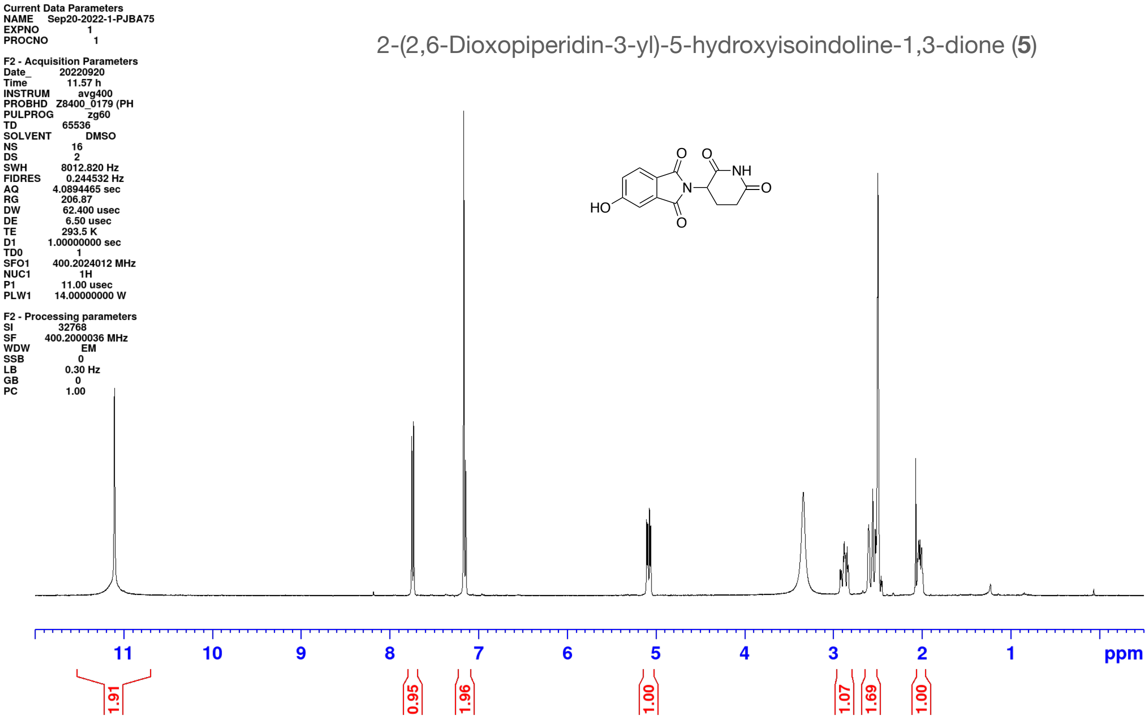
Current Data Parameters
NAME Sep20-2022-1-PJBA75
EXPNO 1
PROCNO 1
F2 - Acquisition Parameters
Date_ 20220920
Time 11.57 h
INSTRUM avg400
PROBHD Z8400_0179 (PH
PULPROG zg60
TD 65536
SOLVENT DMSO
NS 16
DS 2
SWH 8012.820 Hz
FIDRES 0.244532 Hz
AQ 4.0894465 sec
RG 206.87
DW 62.400 usec
DE 6.50 usec
TE 293.5 K
D1 1.00000000 sec
TD0 1
SFO1 400.2024012 MHz
NUC1 1H
P1 11.00 usec
PLW1 14.00000000 W
F2 - Processing parameters
SI 32768
SF 400.2000036 MHz
WDW EM
SSB 0
LB 0.30 Hz
GB 0
PC 1.00
2-(2,6-Dioxopiperidin-3-yl)-5-hydroxyisoindoline-1,3-dione (5)
O
O
NH
HO
N
O
O
11
10
9
8
7
6
5
4
3
2
1
ppm
1.91
0.95
1.96
1.00
1.07
1.69
1.00

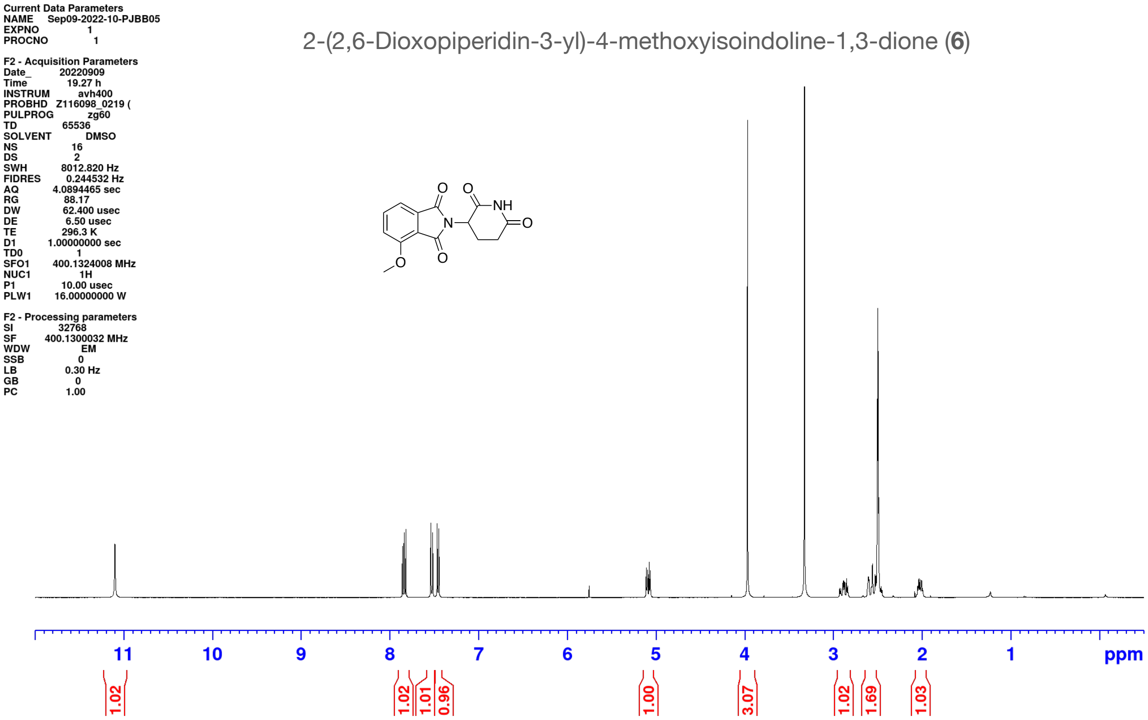

2-(2,6-Dioxopiperidin-3-yl)-4-methoxyisoindoline-1,3-dione (**6**)

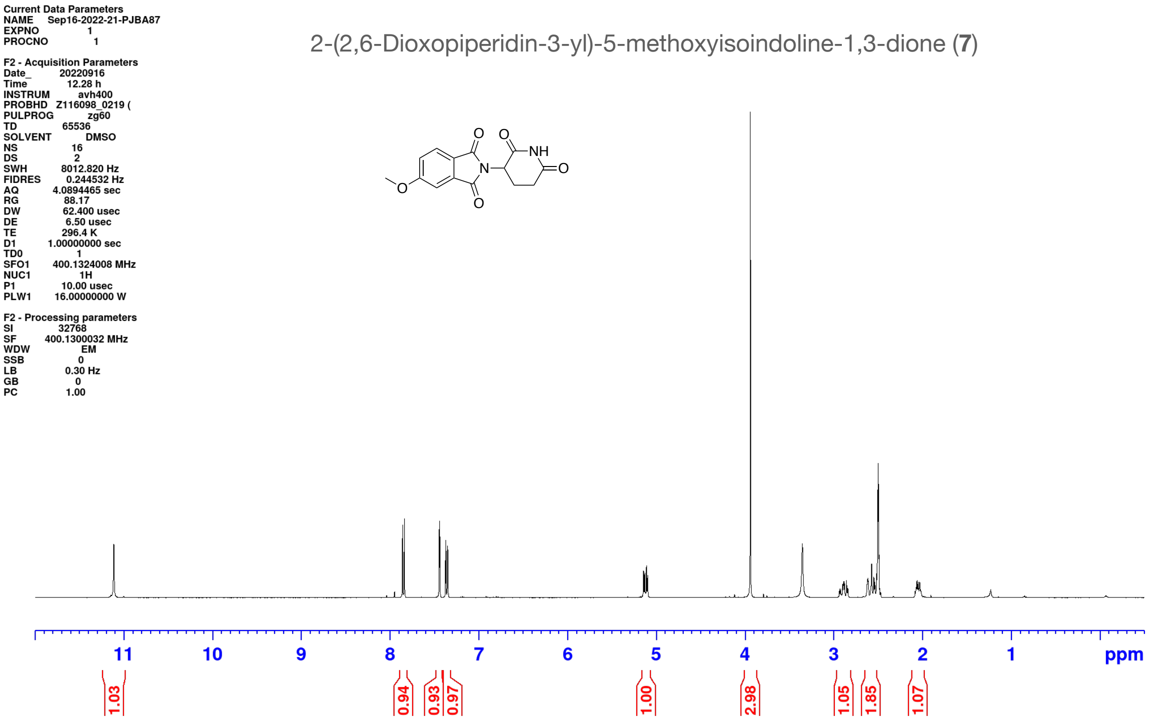
Current Data Parameters
NAME Sep16-2022-21-PJBA87
EXPNO 1
PROCNO 1
F2 - Acquisition Parameters
Date_ 20220916
Time 12.28 h
INSTRUM avh400
PROBHD Z116098_0219 (
PULPROG zg60
TD 65536
SOLVENT DMSO
NS 16
DS 2
SWH 8012.820 Hz
FIDRES 0.244532 Hz
AQ 4.0894465 sec
RG 88.17
DW 62.400 usec
DE 6.50 usec
TE 296.4 K
D1 1.00000000 sec
TD0 1
SFO1 400.1324008 MHz
NUC1 1H
P1 10.00 usec
PLW1 16.00000000 W
F2 - Processing parameters
SI 32768
SF 400.1300032 MHz
WDW EM
SSB 0
LB 0.30 Hz
GB 0
PC 1.00
2-(2,6-Dioxopiperidin-3-yl)-5-methoxyisoindoline-1,3-dione (7)
O
O
NH
N
O
O
O
11
10
9
8
7
6
5
4
3
2
1
ppm
1.03
0.94
0.93
0.97
1.00
2.98
1.05
1.85
1.07

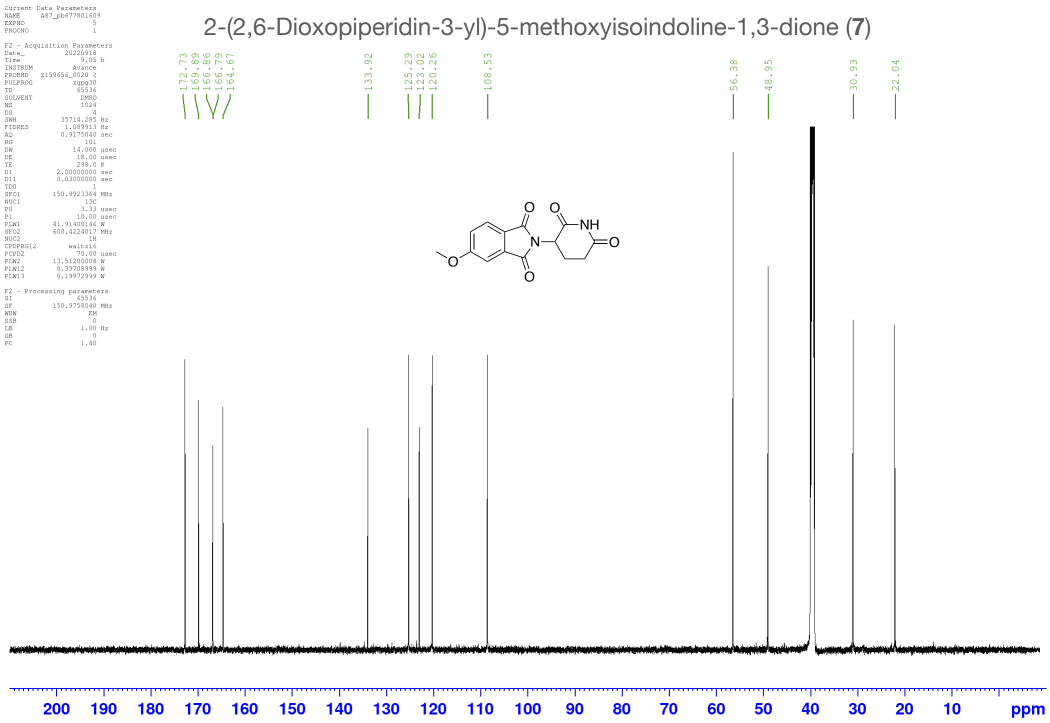
Current Data Parameters
NAME A87_pb677801609
EXPNO 5
PROCNO 1
F2 - Acquisition Parameters
Date_ 20220918
Time 9.05 h
INSTRUM Avance
PROBHD Z159656_0020 (
PULPROG zgpg30
TD 65536
SOLVENT DMSO
NS 1024
DS 4
SWH 35714.285 Hz
FIDRES 1.089913 Hz
AQ 0.9175040 sec
RG 101
DW 14.000 usec
DE 18.00 usec
TE 298.0 K
D1 2.00000000 sec
D11 0.03000000 sec
TD0 1
SFO1 150.9923364 MHz
NUC1 13C
P0 3.33 usec
P1 10.00 usec
PLW1 41.91400146 W
SFO2 600.4224017 MHz
NUC2 1H
CPDPRG[2 waltz16
PCPD2 70.00 usec
PLW2 13.51200008 W
PLW12 0.39708999 W
PLW13 0.19972999 W
F2 - Processing parameters
SI 65536
SF 150.9758040 MHz
WDW EM
SSB 0
LB 1.00 Hz
GB 0
PC 1.40
2-(2,6-Dioxopiperidin-3-yl)-5-methoxyisoindoline-1,3-dione (7)
172.73
169.89
166.86
166.79
164.67
133.92
125.29
123.02
120.26
108.53
56.38
48.95
30.93
22.04
O
O
NH
N
O
O
O
200
190
180
170
160
150
140
130
120
110
100
90
80
70
60
50
40
30
20
10
ppm

2-(2,6-Dioxopiperidin-3-yl)-4-fluoroisoindoline-1,3-dione (**8**)

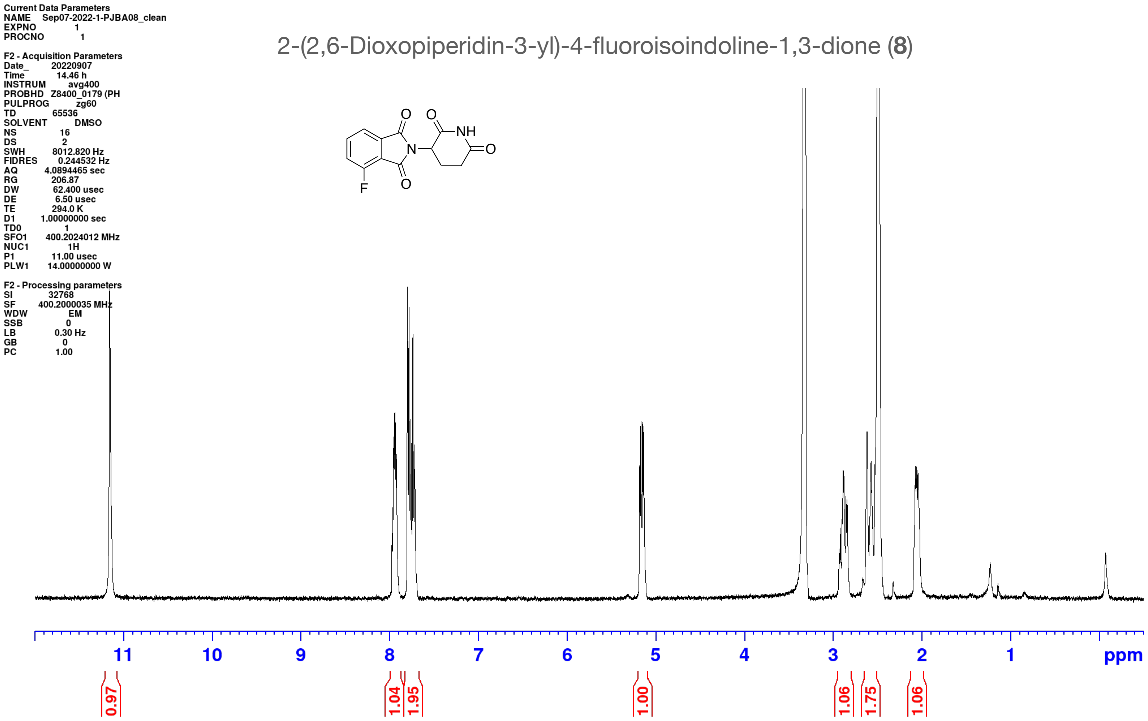

2-(2,6-Dioxopiperidin-3-yl)-5-fluoroisoindoline-1,3-dione (**9**)

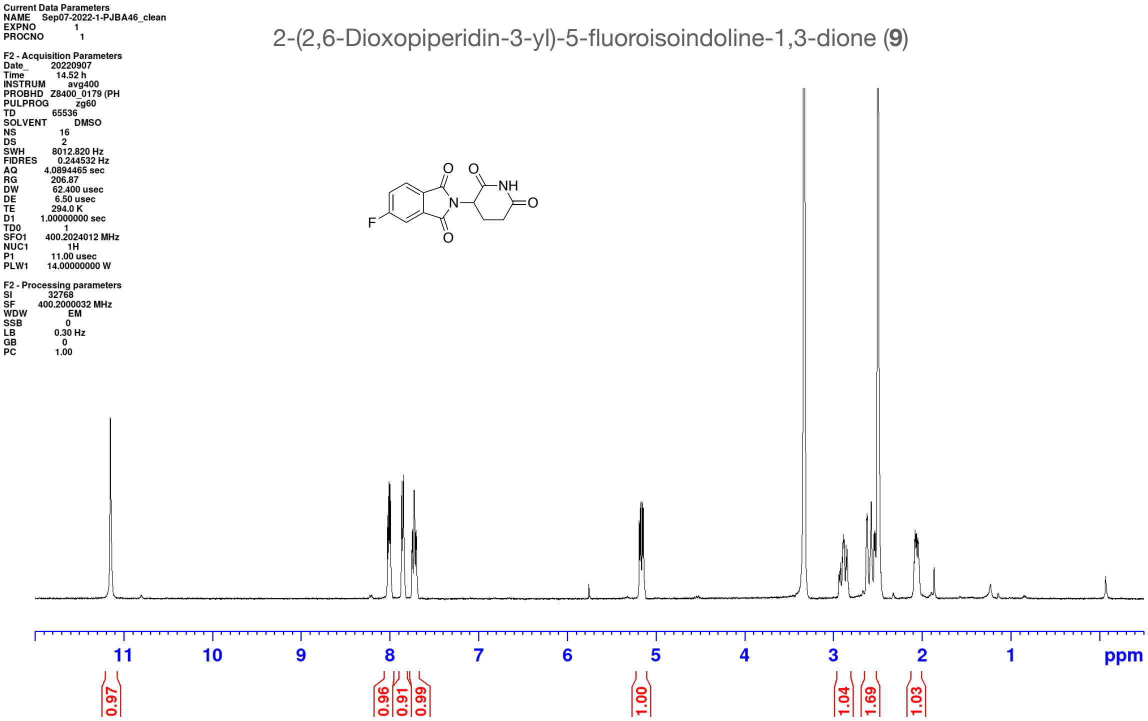

4-Chloro-2-(2,6-dioxopiperidin-3-yl)isoindoline-1,3-dione (**10**)

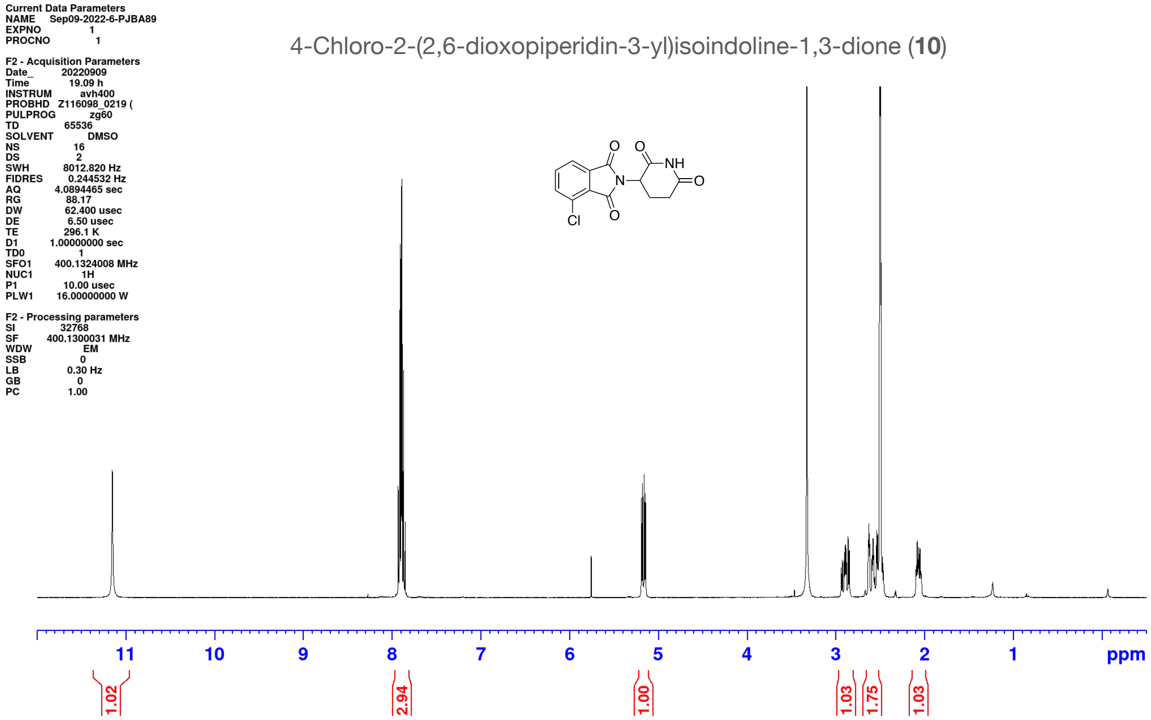

5-Chloro-2-(2,6-dioxopiperidin-3-yl)isoindoline-1,3-dione (**11**)

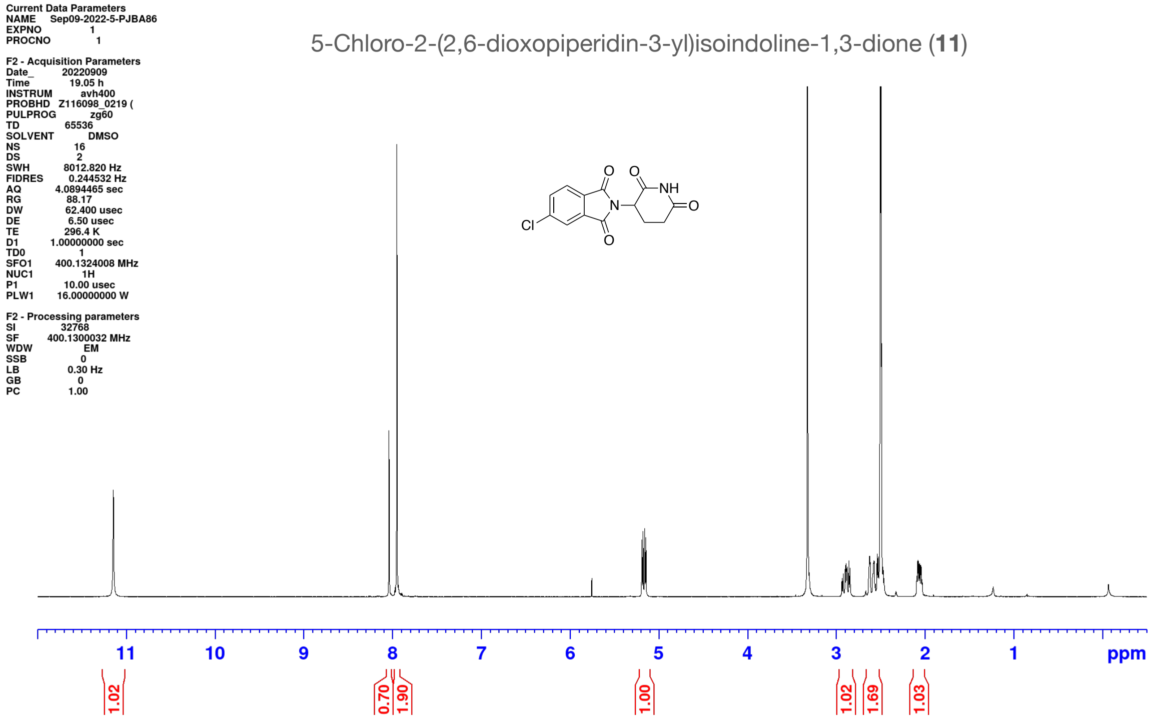

4-Bromo-2-(2,6-dioxopiperidin-3-yl)isoindoline-1,3-dione (**12**)

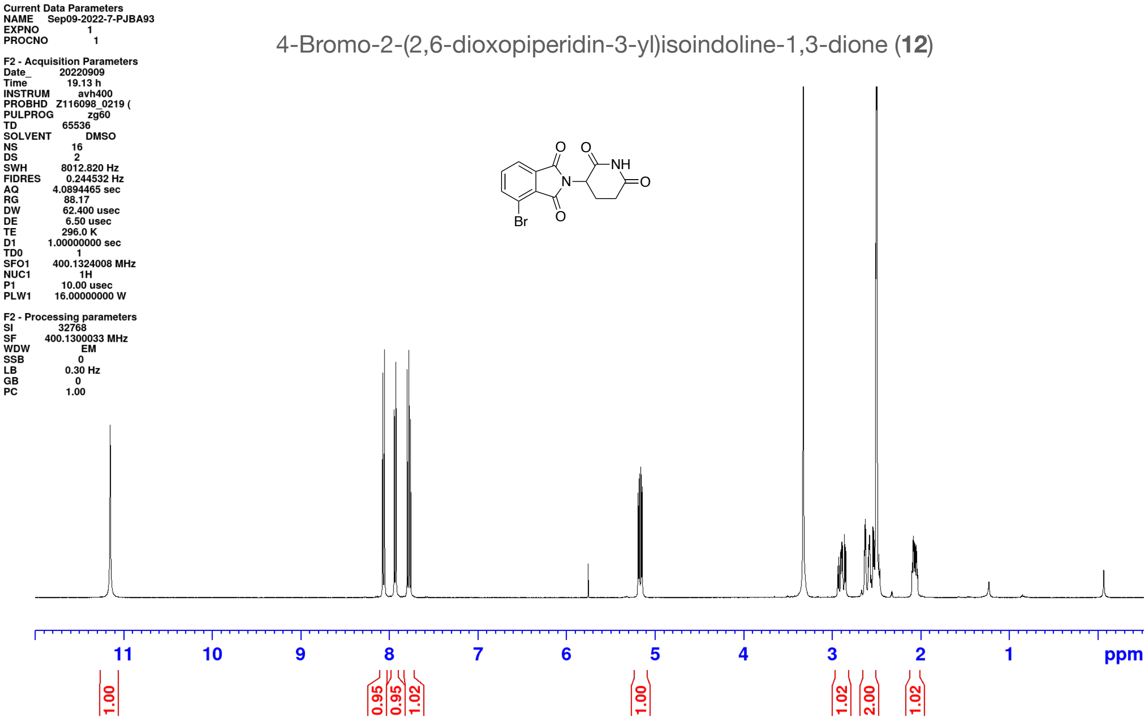

5-Bromo-2-(2,6-dioxopiperidin-3-yl)isoindoline-1,3-dione (**13**)

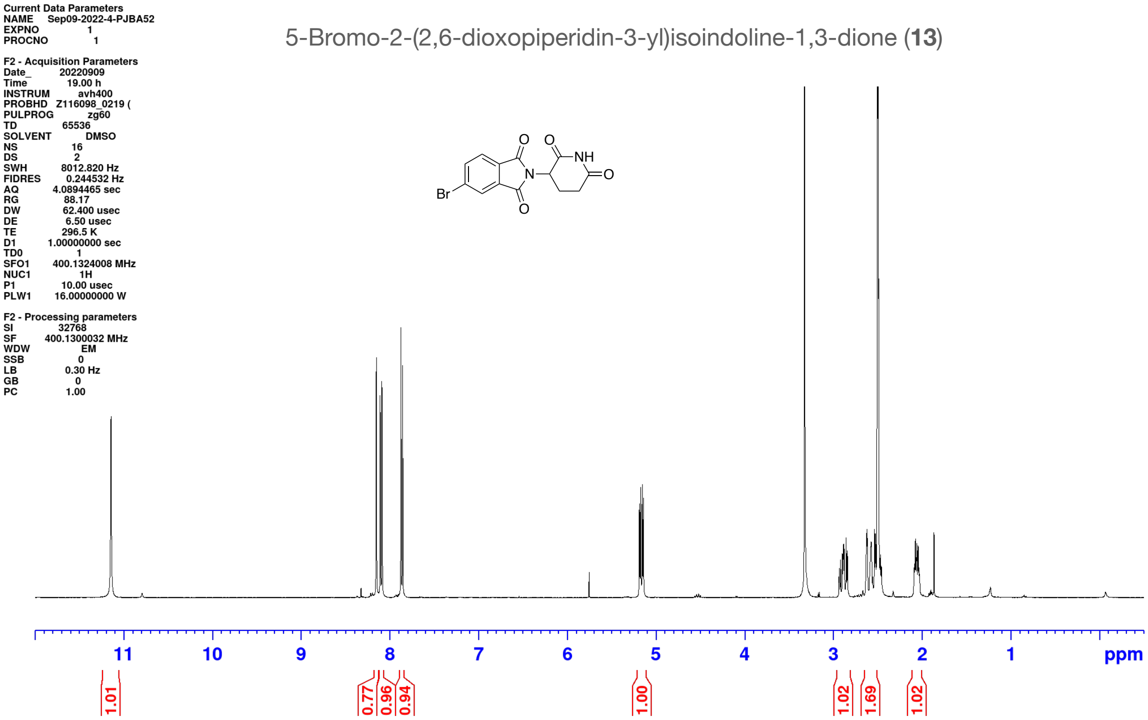

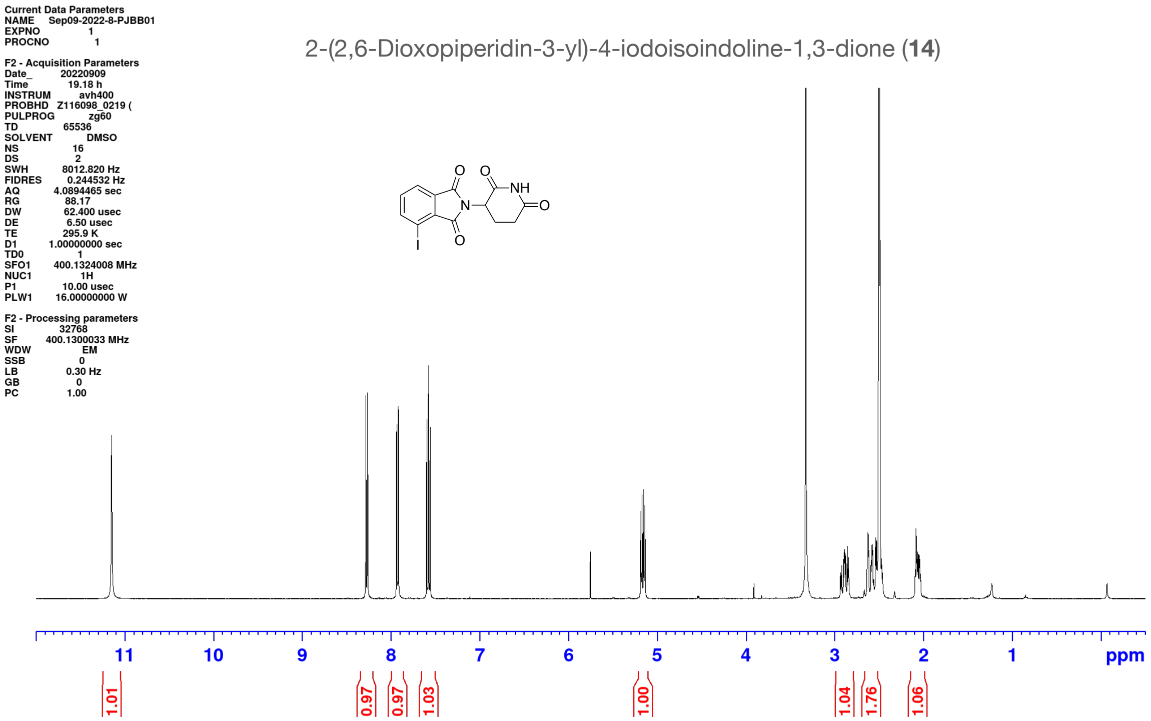
Current Data Parameters
NAME Sep09-2022-8-PJBB01
EXPNO 1
PROCNO 1
F2 - Acquisition Parameters
Date_ 20220909
Time 19.18 h
INSTRUM avh400
PROBHD Z116098_0219 (
PULPROG zg60
TD 65536
SOLVENT DMSO
NS 16
DS 2
SWH 8012.820 Hz
FIDRES 0.244532 Hz
AQ 4.0894465 sec
RG 88.17
DW 62.400 usec
DE 6.50 usec
TE 295.9 K
D1 1.00000000 sec
TD0 1
SFO1 400.1324008 MHz
NUC1 1H
P1 10.00 usec
PLW1 16.00000000 W
F2 - Processing parameters
SI 32768
SF 400.1300033 MHz
WDW EM
SSB 0
LB 0.30 Hz
GB 0
PC 1.00
2-(2,6-Dioxopiperidin-3-yl)-4-iodoisoindoline-1,3-dione (**14**)
O
O
NH
N
O
O
I
11
10
9
8
7
6
5
4
3
2
1
ppm
1.01
0.97
0.97
1.03
1.00
1.04
1.76
1.06

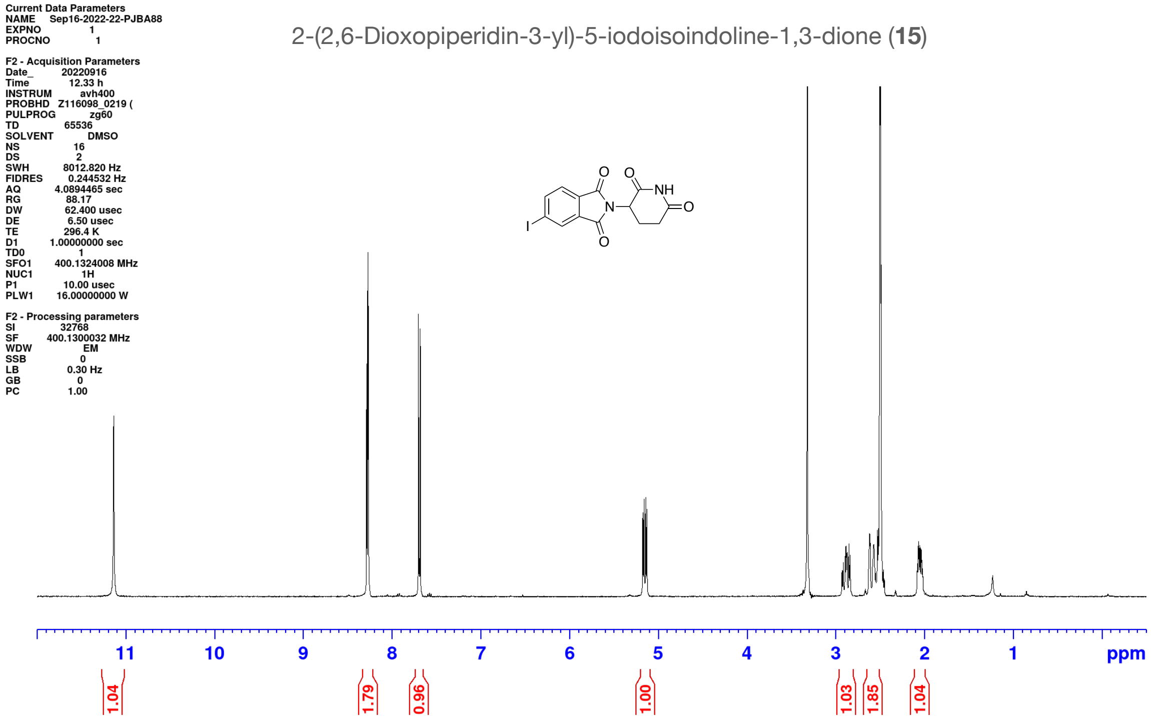
Current Data Parameters
NAME Sep16-2022-22-PJBA88
EXPNO 1
PROCNO 1
F2 - Acquisition Parameters
Date_ 20220916
Time 12.33 h
INSTRUM avh400
PROBHD Z116098_0219 (
PULPROG zg60
TD 65536
SOLVENT DMSO
NS 16
DS 2
SWH 8012.820 Hz
FIDRES 0.244532 Hz
AQ 4.0894465 sec
RG 88.17
DW 62.400 usec
DE 6.50 usec
TE 296.4 K
D1 1.00000000 sec
TD0 1
SFO1 400.1324008 MHz
NUC1 1H
P1 10.00 usec
PLW1 16.00000000 W
F2 - Processing parameters
SI 32768
SF 400.1300032 MHz
WDW EM
SSB 0
LB 0.30 Hz
GB 0
PC 1.00
2-(2,6-Dioxopiperidin-3-yl)-5-iodoisoindoline-1,3-dione (15)
11 10 9 8 7 6 5 4 3 2 1 ppm
1.04
1.79
0.96
1.00
1.03
1.85
1.04

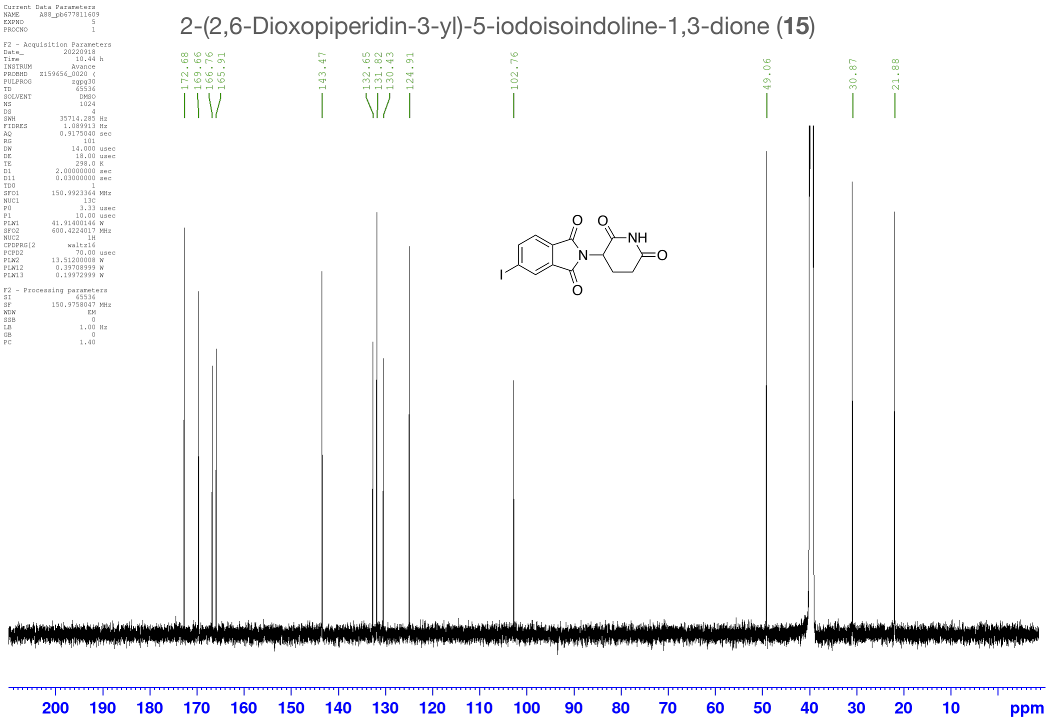
2-(2,6-Dioxopiperidin-3-yl)-5-iodoisoindoline-1,3-dione (15)
172.68
169.66
166.76
165.91
143.47
132.65
131.82
130.43
124.91
102.76
49.06
30.87
21.88
200 190 180 170 160 150 140 130 120 110 100 90 80 70 60 50 40 30 20 10 ppm

Current Data Parameters
NAME Feb04-2020-1-PJB-postcol1-DMSO__A13
EXPNO 1
PROCNO 1

F2 - Acquisition Parameters
Date_ 20200204
Time 12.15 h
INSTRUM avh400
PROBHD Z108618_0873 (
PULPROG zg60
TD 65536
SOLVENT DMSO
NS 16
DS 2
SWH 8012.820 Hz
FIDRES 0.244532 Hz
AQ 4.0894465 sec
RG 88.17
DW 62.400 usec
DE 6.50 usec
TE 298.8 K
D1 1.00000000 sec
TD0 1
SFO1 400.1324008 MHz
NUC1 1H
P1 14.00 usec
PLW1 14.36999989 W

F2 - Processing parameters
SI 32768
SF 400.1300031 MHz
WDW EM
SSB 0
LB 0.30 Hz
GB 0
PC 1.00

2-(2,6-Dioxopiperidin-3-yl)-4-methylisoindoline-1,3-dione (**16**)

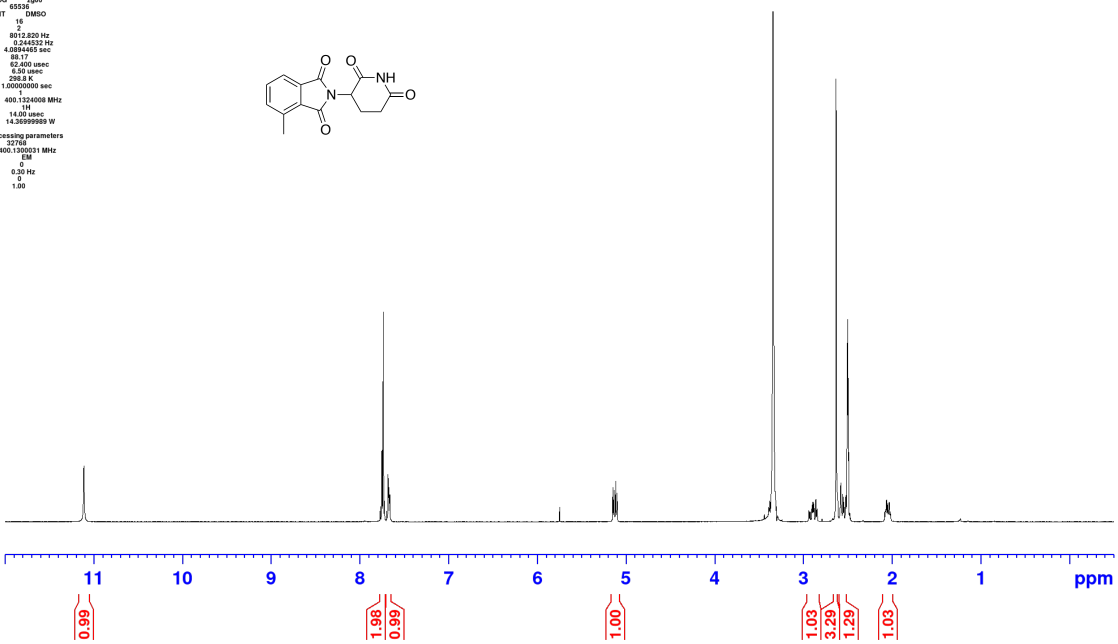

Current Data Parameters
NAME Sep09-2022-2-PJBA45
EXPNO 1
PROCNO 1

F2 - Acquisition Parameters
Date_ 20220909
Time 18.51 h
INSTRUM avh400
PROBHD Z116098_0219 (
PULPROG zg60
TD 65536
SOLVENT DMSO
NS 16
DS 2
SWH 8012.820 Hz
FIDRES 0.244532 Hz
AQ 4.0894465 sec
RG 88.17
DW 62.400 usec
DE 6.50 usec
TE 296.4 K
D1 1.00000000 sec
TD0 1
SFO1 400.1324008 MHz
NUC1 1H
P1 10.00 usec
PLW1 16.00000000 W

F2 - Processing parameters
SI 32768
SF 400.1300034 MHz
WDW EM
SSB 0
LB 0.30 Hz
GB 0
PC 1.00

2-(2,6-Dioxopiperidin-3-yl)-5-methylisoindoline-1,3-dione (**17**)

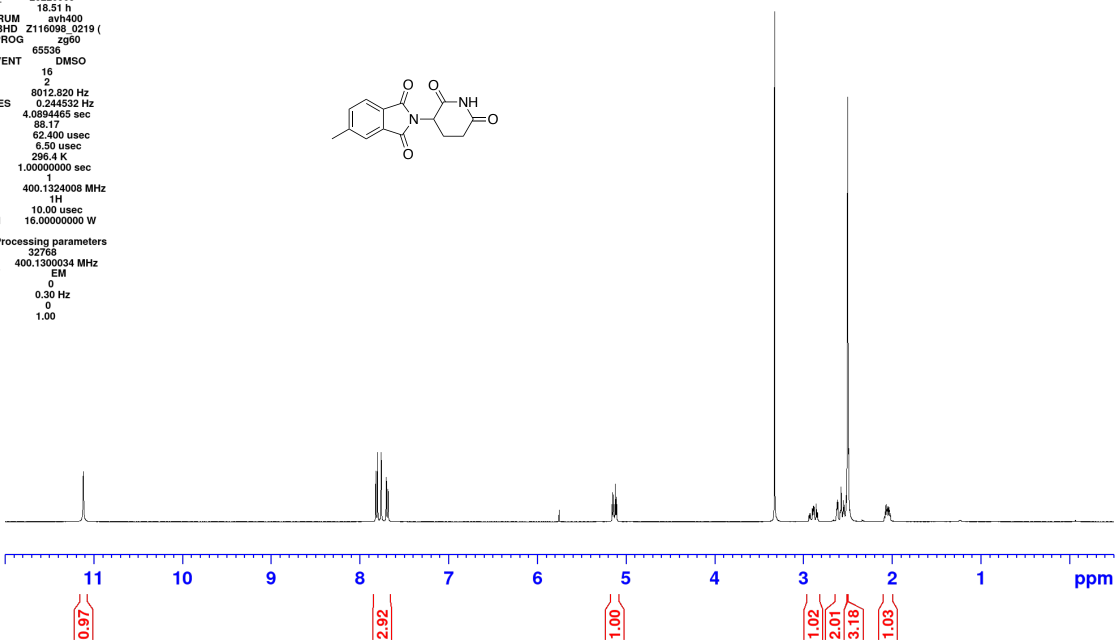

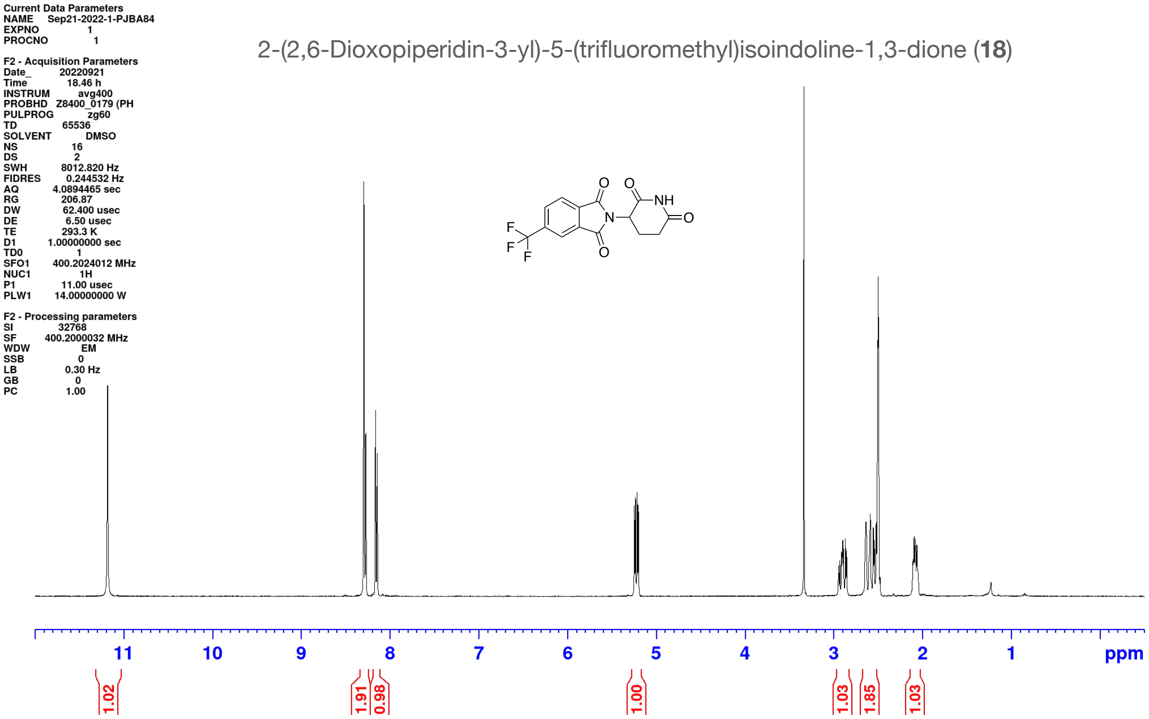
2-(2,6-Dioxopiperidin-3-yl)-5-(trifluoromethyl)isoindoline-1,3-dione (18)

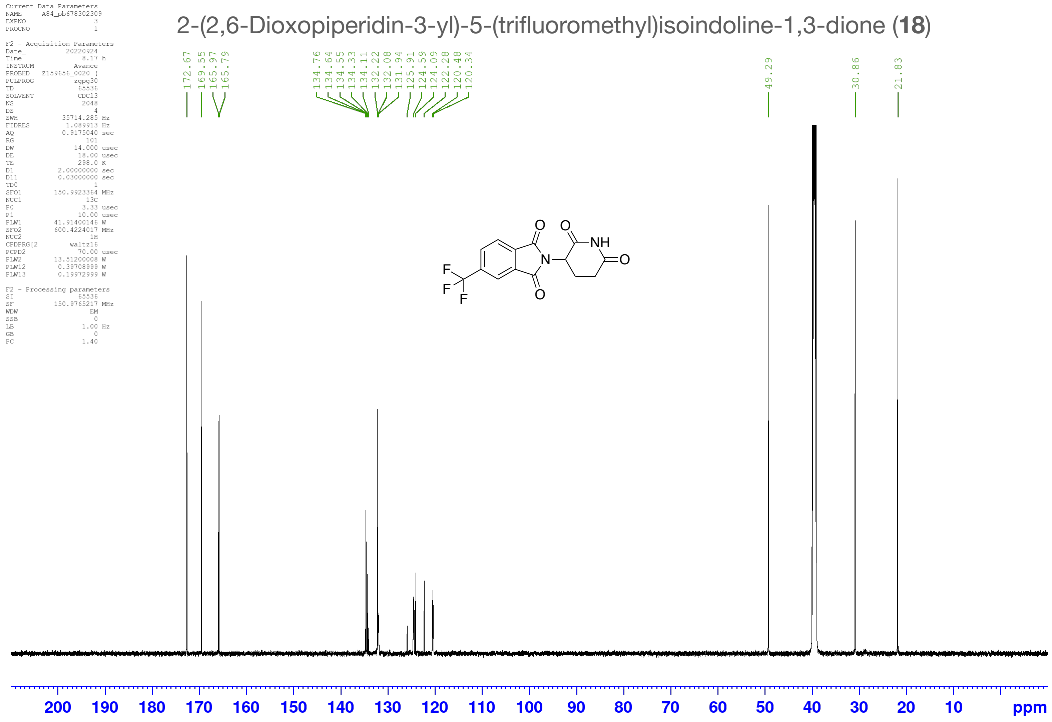
2-(2,6-Dioxopiperidin-3-yl)-5-(trifluoromethyl)isoindoline-1,3-dione (18)

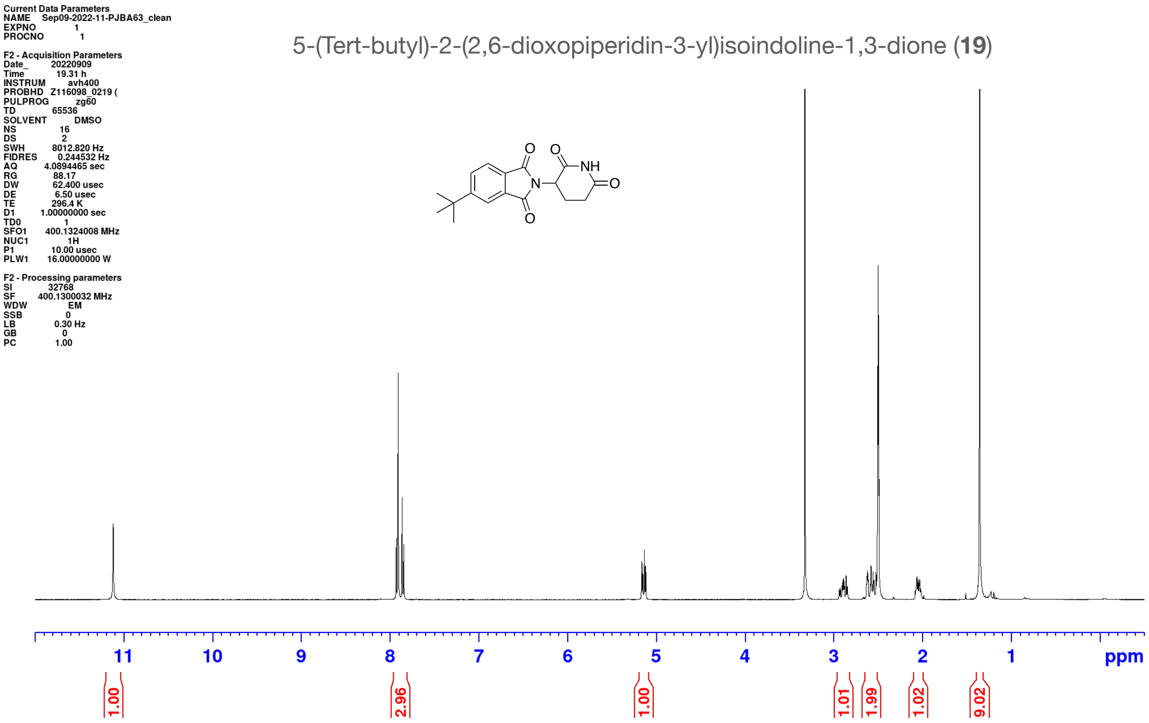
Current Data Parameters
NAME Sep09-2022-11-PJBA63_clean
EXPNO 1
PROCNO 1
F2 - Acquisition Parameters
Date_ 20220909
Time 19.31 h
INSTRUM avh400
PROBHD Z116098_0219 (
PULPROG zg60
TD 65536
SOLVENT DMSO
NS 16
DS 2
SWH 8012.820 Hz
FIDRES 0.244532 Hz
AQ 4.0894465 sec
RG 88.17
DW 62.400 usec
DE 6.50 usec
TE 296.4 K
D1 1.00000000 sec
TD0 1
SFO1 400.1324008 MHz
NUC1 1H
P1 10.00 usec
PLW1 16.00000000 W
F2 - Processing parameters
SI 32768
SF 400.1300032 MHz
WDW EM
SSB 0
LB 0.30 Hz
GB 0
PC 1.00
5-(Tert-butyl)-2-(2,6-dioxopiperidin-3-yl)isoindoline-1,3-dione (19)
O
O
NH
N
O
O
11
10
9
8
7
6
5
4
3
2
1
ppm
1.00
2.96
1.00
1.01
1.99
1.02
9.02

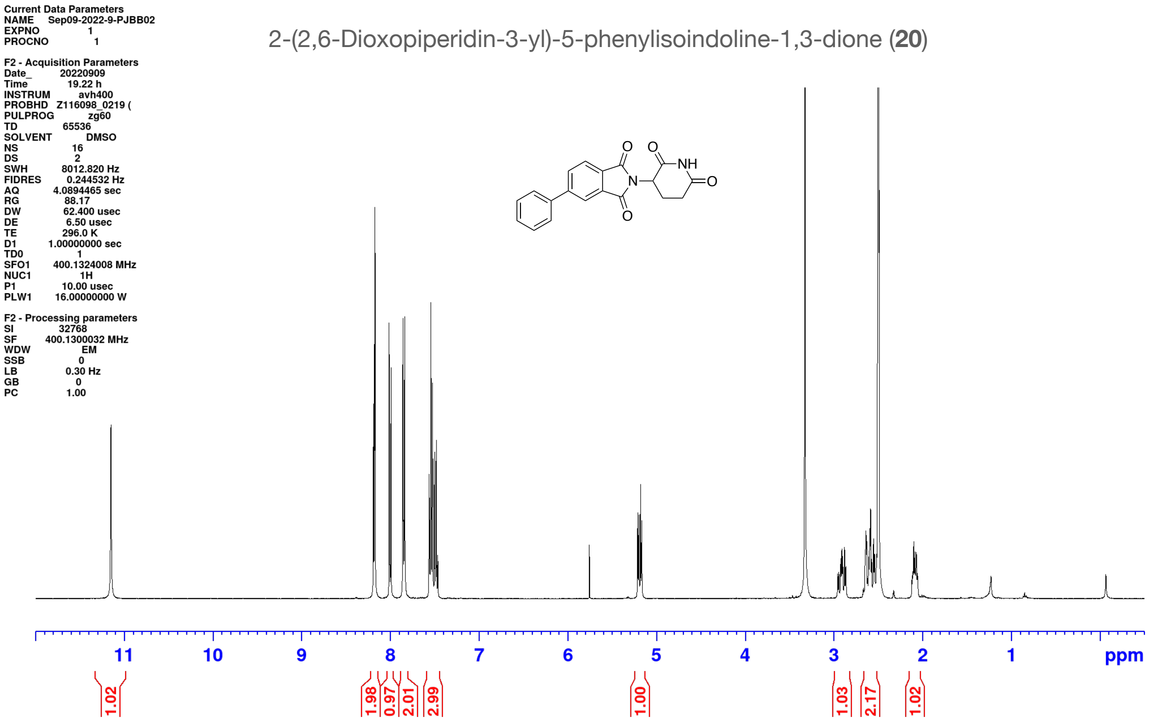
Current Data Parameters
NAME Sep09-2022-9-PJBB02
EXPNO 1
PROCNO 1
F2 - Acquisition Parameters
Date_ 20220909
Time 19.22 h
INSTRUM avh400
PROBHD Z116098_0219 (
PULPROG zg60
TD 65536
SOLVENT DMSO
NS 16
DS 2
SWH 8012.820 Hz
FIDRES 0.244532 Hz
AQ 4.0894465 sec
RG 88.17
DW 62.400 usec
DE 6.50 usec
TE 296.0 K
D1 1.00000000 sec
TD0 1
SFO1 400.1324008 MHz
NUC1 1H
P1 10.00 usec
PLW1 16.00000000 W
F2 - Processing parameters
SI 32768
SF 400.1300032 MHz
WDW EM
SSB 0
LB 0.30 Hz
GB 0
PC 1.00
2-(2,6-Dioxopiperidin-3-yl)-5-phenylisoindoline-1,3-dione (20)
O
O
NH
N
O
O
11
10
9
8
7
6
5
4
3
2
1
ppm
1.02
1.98
0.97
2.01
2.99
1.00
1.03
2.17
1.02

Current Data Parameters
NAME Dec17-2020-3-PJBA62-postcol1
EXPNO 1
PROCNO 1

F2 - Acquisition Parameters
Date_ 20201217
Time 18.34 h
INSTRUM avh400
PROBHD Z108618_0873 (
PULPROG zg60
TD 65536
SOLVENT CDCl3
NS 16
DS 2
SWH 8012.820 Hz
FIDRES 0.244532 Hz
AQ 4.0894465 sec
RG 88.17
DW 62.400 usec
DE 6.50 usec
TE 299.8 K
D1 1.00000000 sec
TD0 1
SFO1 400.1324008 MHz
NUC1 1H
P1 14.00 usec
PLW1 14.36999989 W

F2 - Processing parameters
SI 32768
SF 400.1300101 MHz
WDW EM
SSB 0
LB 0.30 Hz
GB 0
PC 1.00

5-(Dimethylamino)-2-(2,6-dioxopiperidin-3-yl)isoindoline-1,3-dione (**21**)

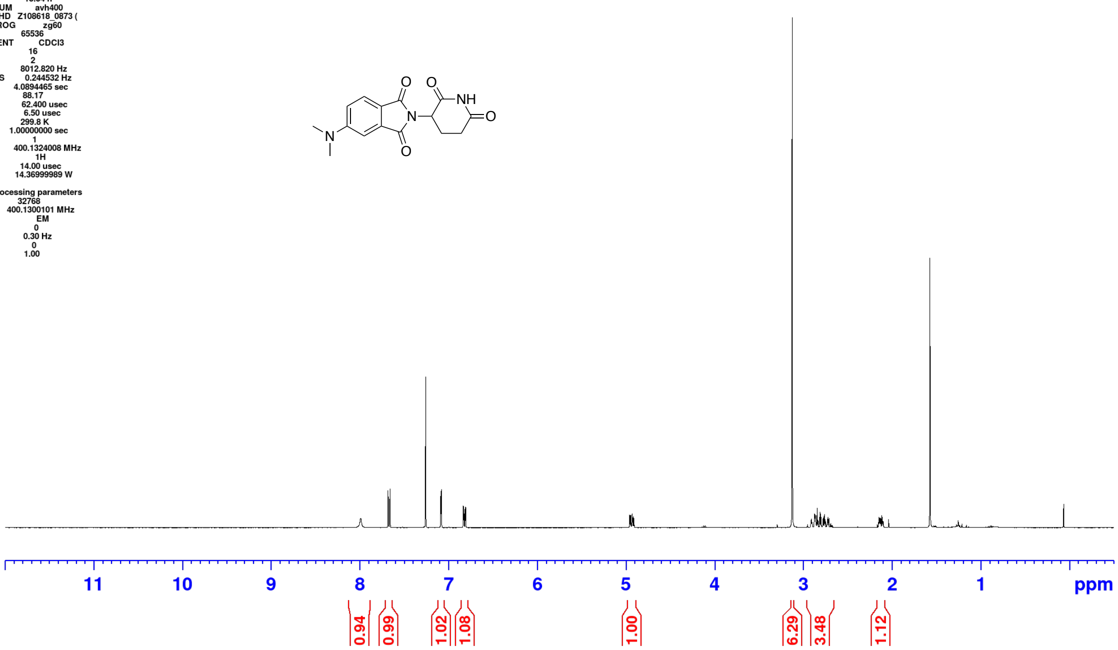

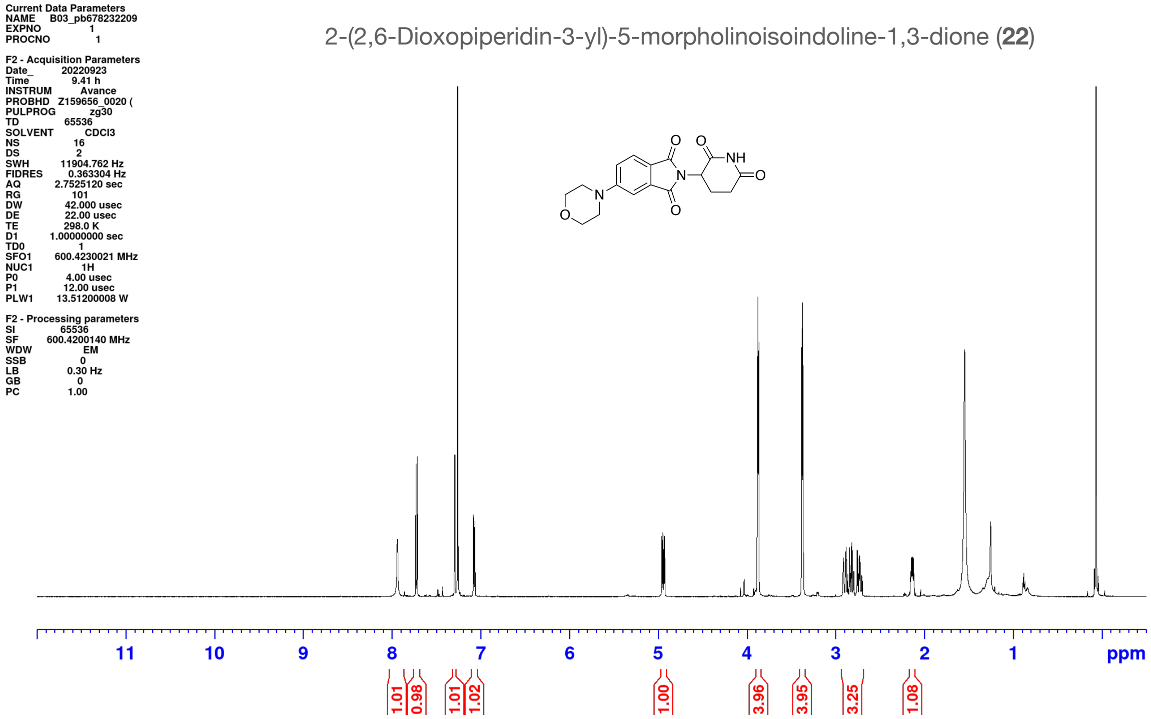
Current Data Parameters
NAME B03_pb678232209
EXPNO 1
PROCNO 1
F2 - Acquisition Parameters
Date_ 20220923
Time 9.41 h
INSTRUM Avance
PROBHD Z159656_0020 (
PULPROG zg30
TD 65536
SOLVENT CDCl3
NS 16
DS 2
SWH 11904.762 Hz
FIDRES 0.363304 Hz
AQ 2.7525120 sec
RG 101
DW 42.000 usec
DE 22.00 usec
TE 298.0 K
D1 1.00000000 sec
TD0 1
SFO1 600.4230021 MHz
NUC1 1H
P0 4.00 usec
P1 12.00 usec
PLW1 13.51200008 W
F2 - Processing parameters
SI 65536
SF 600.4200140 MHz
WDW EM
SSB 0
LB 0.30 Hz
GB 0
PC 1.00
2-(2,6-Dioxopiperidin-3-yl)-5-morpholinoisoindoline-1,3-dione (**22**)
11 10 9 8 7 6 5 4 3 2 1 ppm
1.01 0.98 1.01 1.02 1.00 3.96 3.95 3.25 1.08

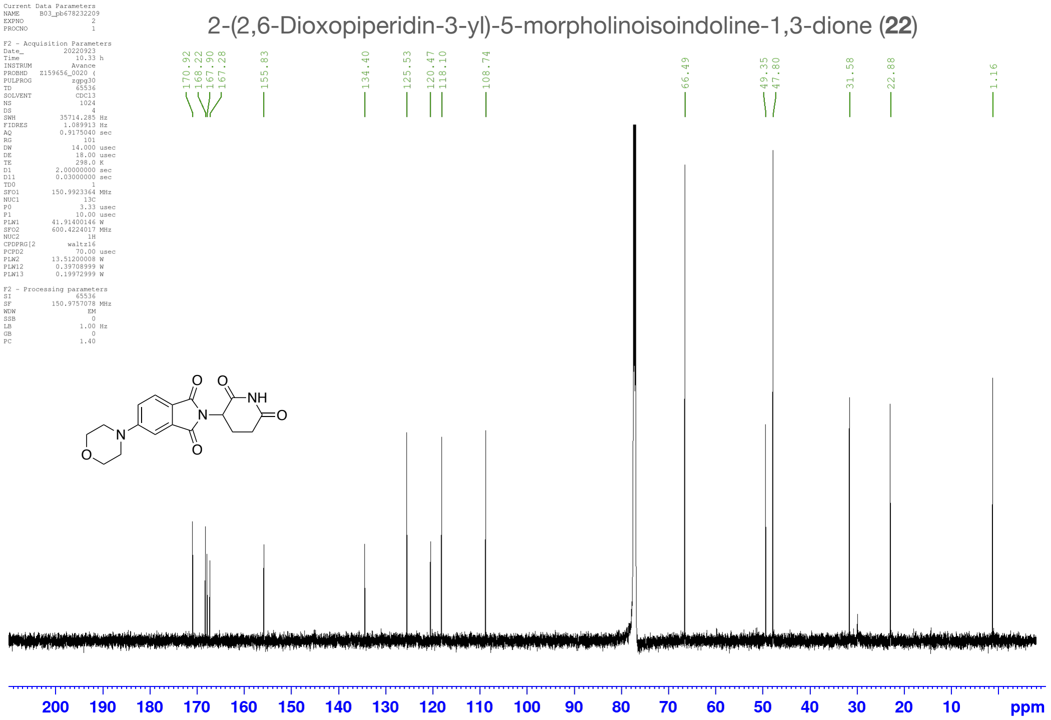
2-(2,6-Dioxopiperidin-3-yl)-5-morpholinoisoindoline-1,3-dione (**22**)
170.92 168.22 167.90 167.28 155.83 134.40 125.53 120.47 118.10 108.74 66.49 49.35 47.80 31.58 22.88 1.16
200 190 180 170 160 150 140 130 120 110 100 90 80 70 60 50 40 30 20 10 ppm

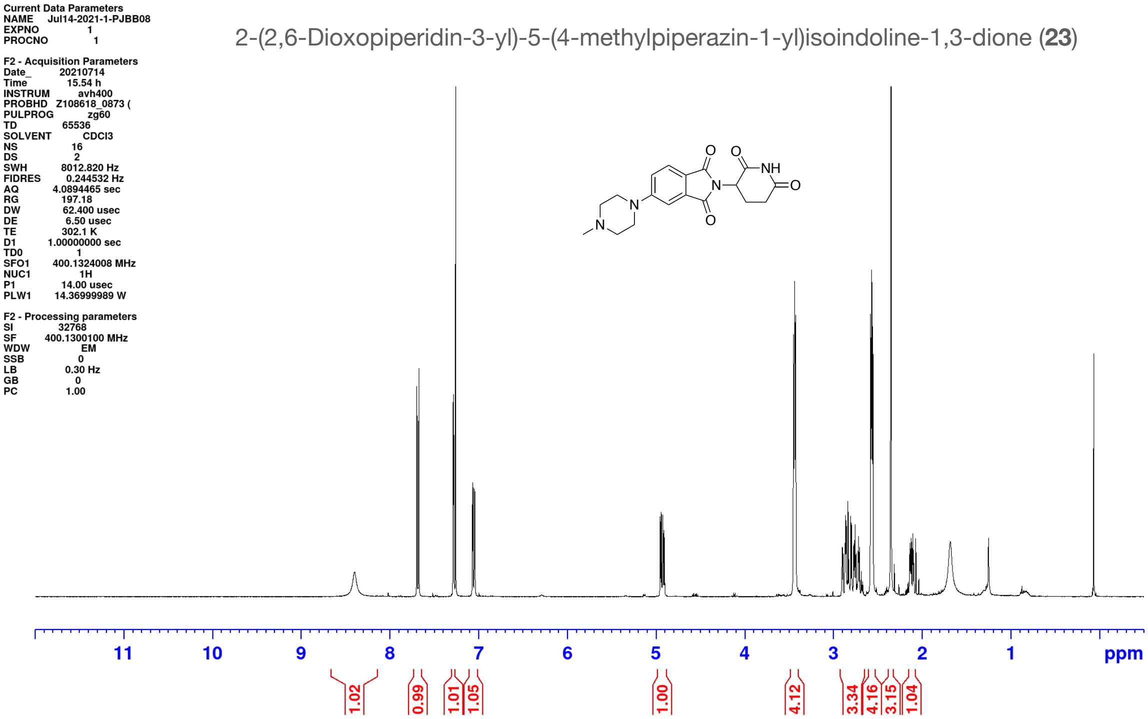
Current Data Parameters
NAME Jul14-2021-1-PJBB08
EXPNO 1
PROCNO 1
F2 - Acquisition Parameters
Date_ 20210714
Time 15.54 h
INSTRUM avh400
PROBHD Z108618_0873 (
PULPROG zg60
TD 65536
SOLVENT CDCl3
NS 16
DS 2
SWH 8012.820 Hz
FIDRES 0.244532 Hz
AQ 4.0894465 sec
RG 197.18
DW 62.400 usec
DE 6.50 usec
TE 302.1 K
D1 1.00000000 sec
TD0 1
SFO1 400.1324008 MHz
NUC1 1H
P1 14.00 usec
PLW1 14.36999989 W
F2 - Processing parameters
SI 32768
SF 400.1300100 MHz
WDW EM
SSB 0
LB 0.30 Hz
GB 0
PC 1.00
2-(2,6-Dioxopiperidin-3-yl)-5-(4-methylpiperazin-1-yl)isoindoline-1,3-dione (23)
11 10 9 8 7 6 5 4 3 2 1 ppm
1.02 0.99 1.01 1.05 1.00 4.12 3.34 4.16 3.15 1.04

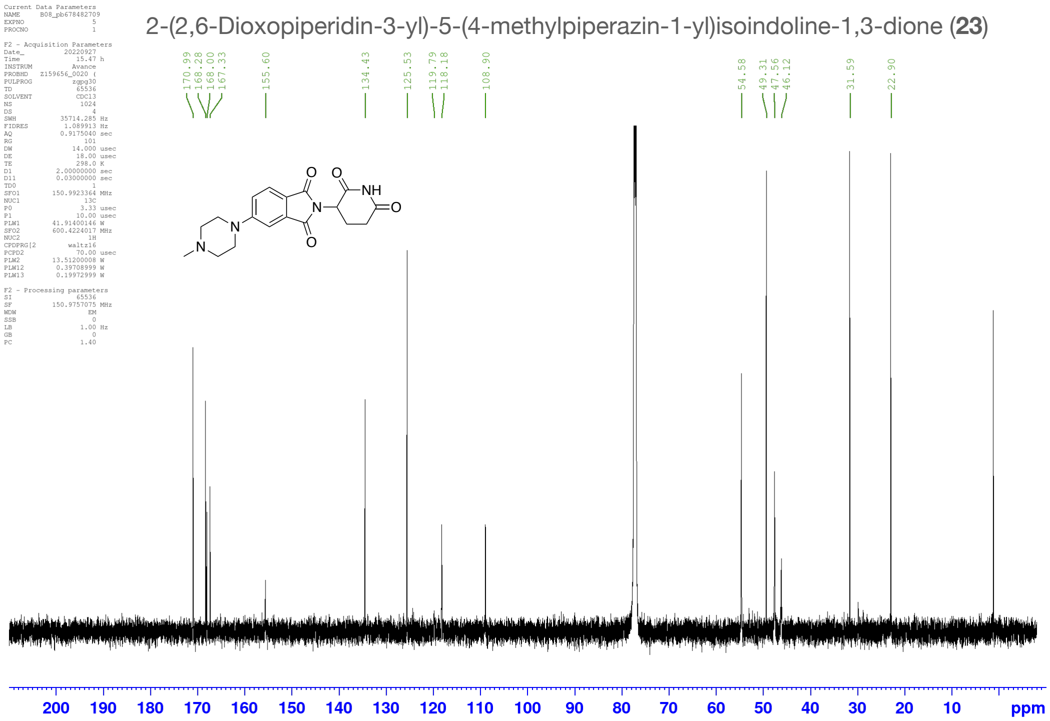
2-(2,6-Dioxopiperidin-3-yl)-5-(4-methylpiperazin-1-yl)isoindoline-1,3-dione (23)
170.99
168.28
168.00
167.33
155.60
134.43
125.53
119.79
118.18
108.90
54.58
49.31
47.56
46.12
31.59
22.90
200 190 180 170 160 150 140 130 120 110 100 90 80 70 60 50 40 30 20 10 ppm

2-(2,6-Dioxopiperidin-3-yl)-5-(4-methylpiperazin-1-yl)isoindoline-1,3-dione (**23**)

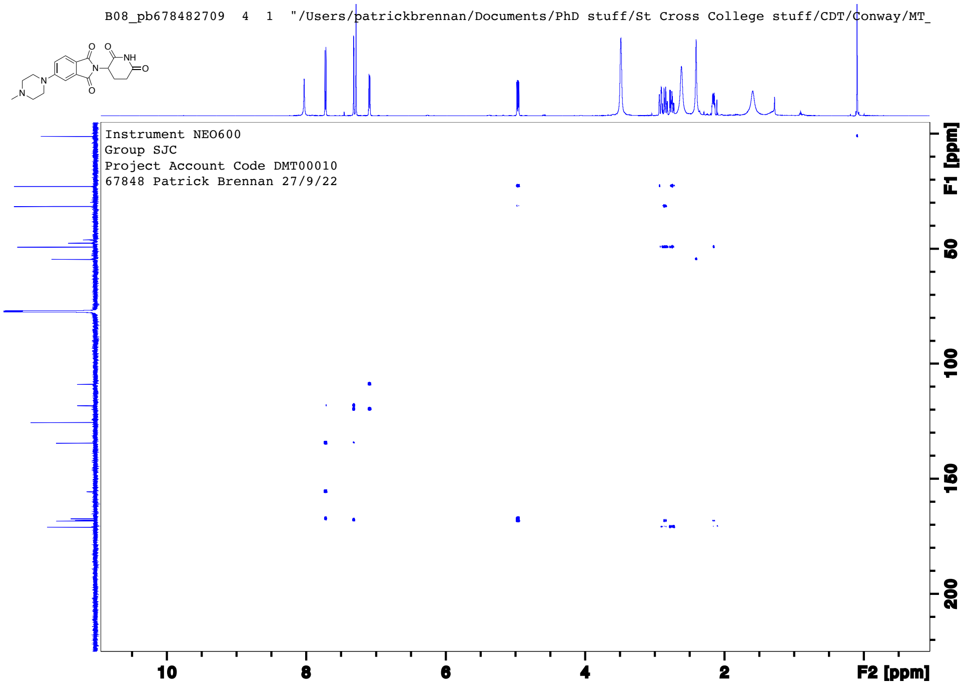

2-(2,6-Dioxopiperidin-3-yl)-5-(4-methylpiperazin-1-yl)isoindoline-1,3-dione (**23**)

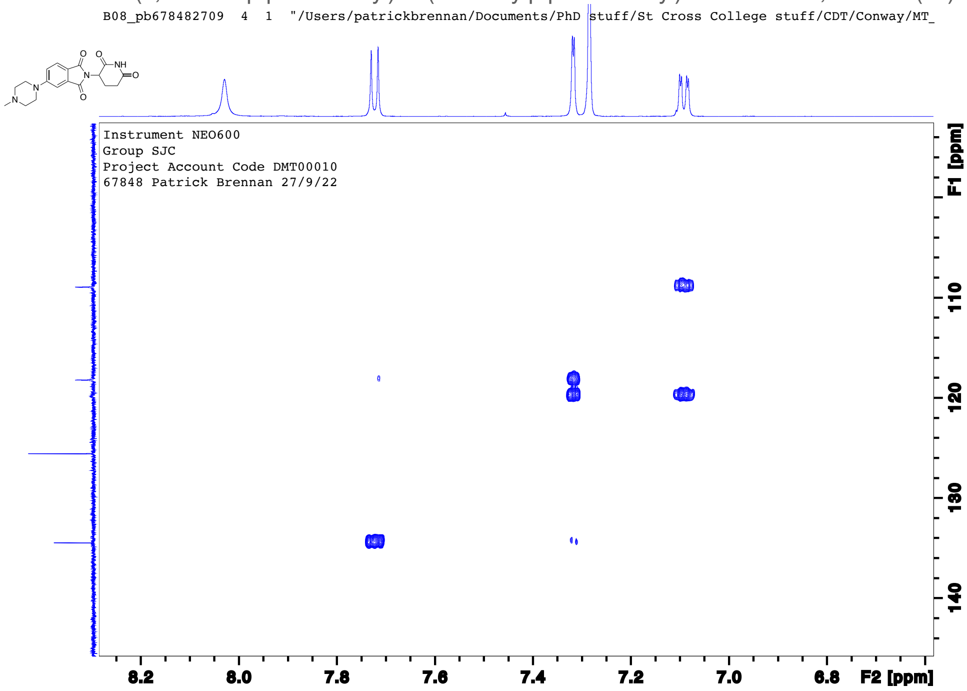

2-(2,6-Dioxopiperidin-3-yl)-4-(ethylamino)isoindoline-1,3-dione (**24**)

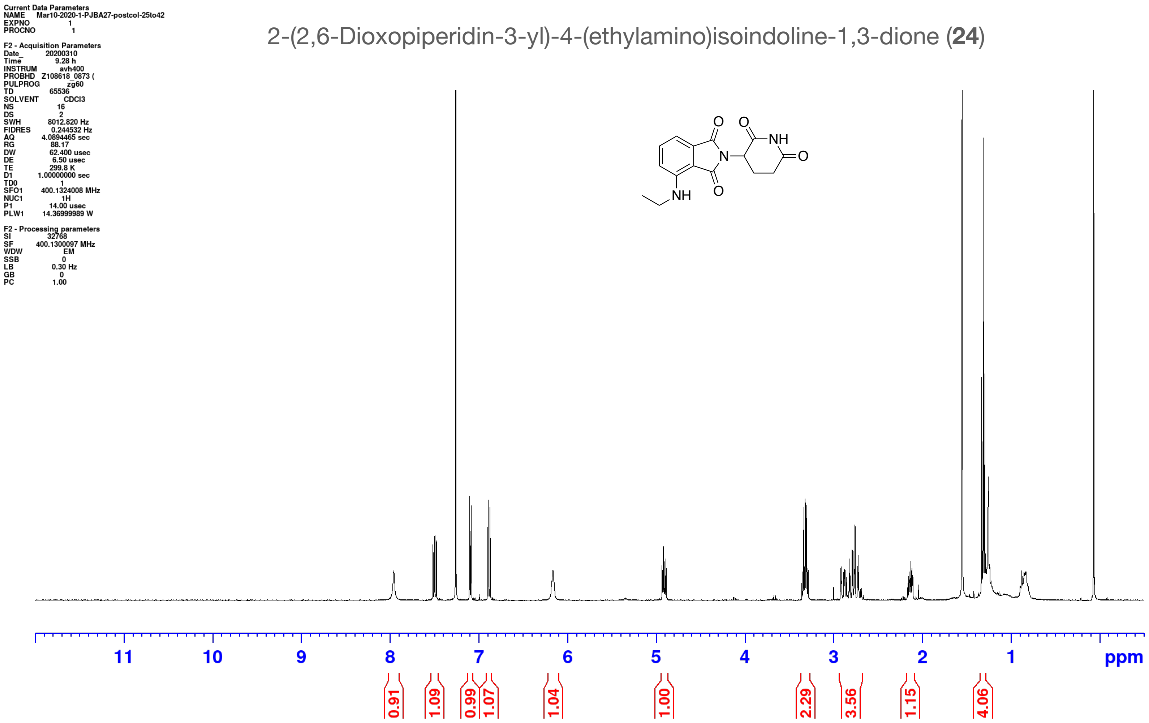

2-(2,6-Dioxopiperidin-3-yl)-4-(prop-2-yn-1-ylamino)isoindoline-1,3-dione (**25**)

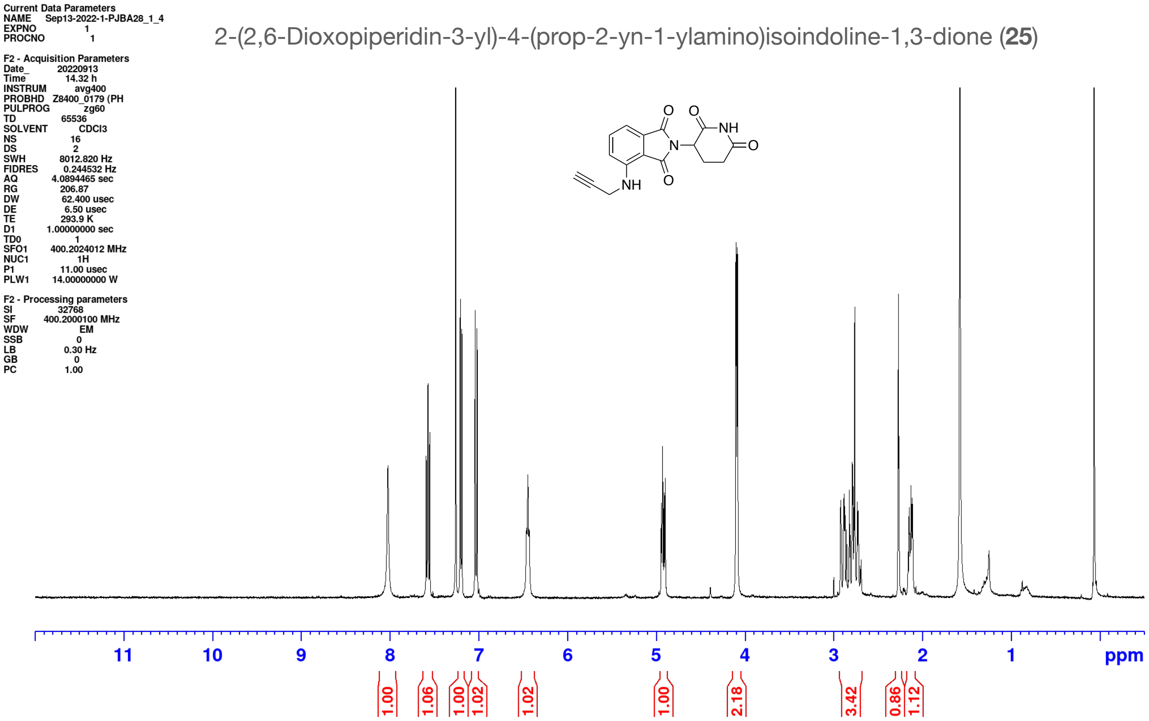

4-(Benzylamino)-2-(2,6-dioxopiperidin-3-yl)isoindoline-1,3-dione (**26**)

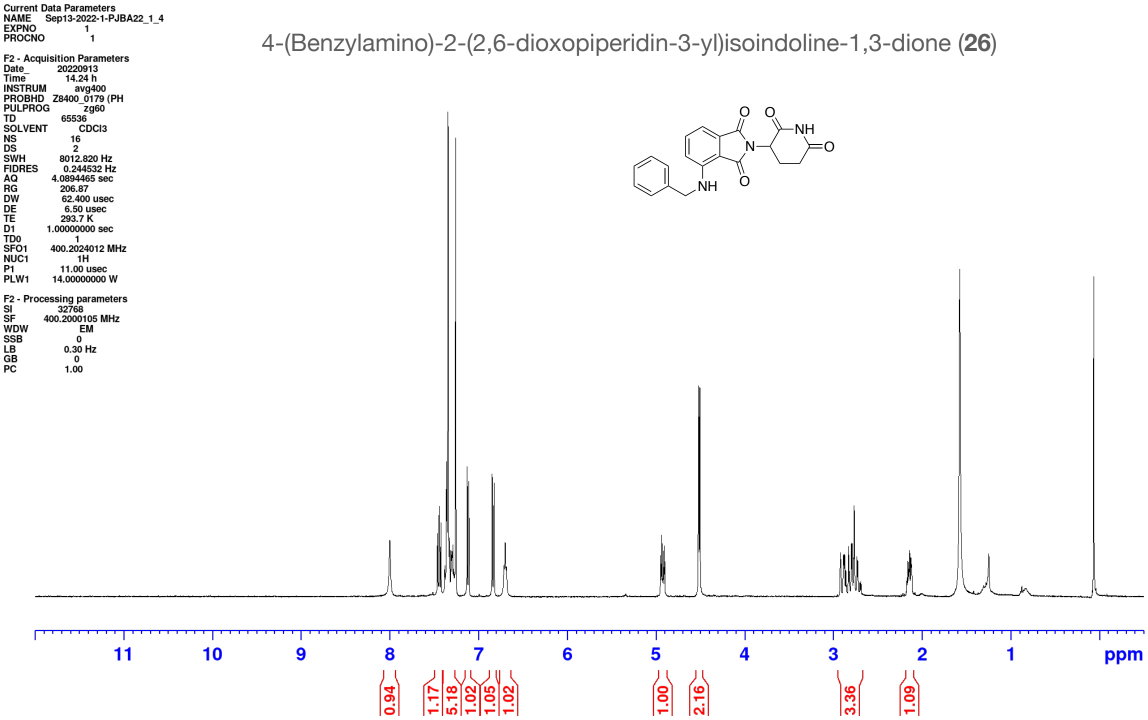

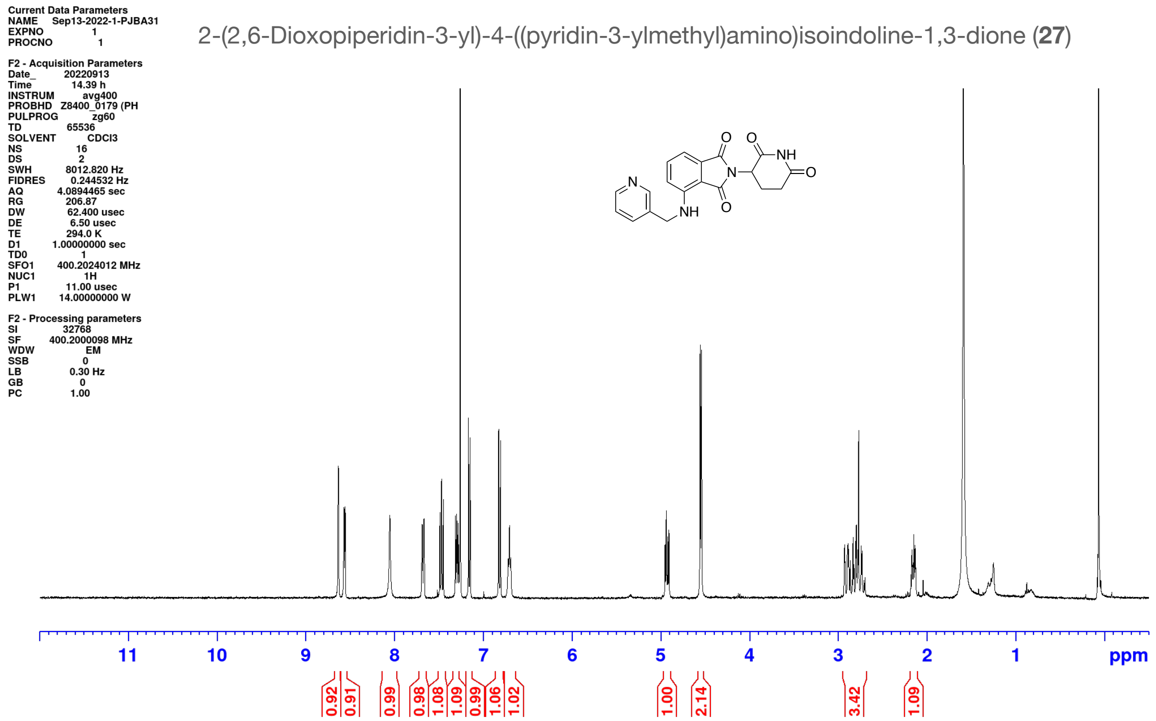
Current Data Parameters
NAME Sep13-2022-1-PJBA31
EXPNO 1
PROCNO 1
F2 - Acquisition Parameters
Date_ 20220913
Time 14.39 h
INSTRUM avg400
PROBHD Z8400_0179 (PH
PULPROG zg60
TD 65536
SOLVENT CDCl3
NS 16
DS 2
SWH 8012.820 Hz
FIDRES 0.244532 Hz
AQ 4.0894465 sec
RG 206.87
DW 62.400 usec
DE 6.50 usec
TE 294.0 K
D1 1.00000000 sec
TD0 1
SFO1 400.2024012 MHz
NUC1 1H
P1 11.00 usec
PLW1 14.00000000 W
F2 - Processing parameters
SI 32768
SF 400.2000098 MHz
WDW EM
SSB 0
LB 0.30 Hz
GB 0
PC 1.00
2-(2,6-Dioxopiperidin-3-yl)-4-((pyridin-3-ylmethyl)amino)isoindoline-1,3-dione (27)
N
NH
O
O
O
N
NH
O
11
10
9
8
7
6
5
4
3
2
1
ppm
0.92
0.91
0.99
0.98
1.08
1.09
0.99
1.06
1.02
1.00
2.14
3.42
1.09

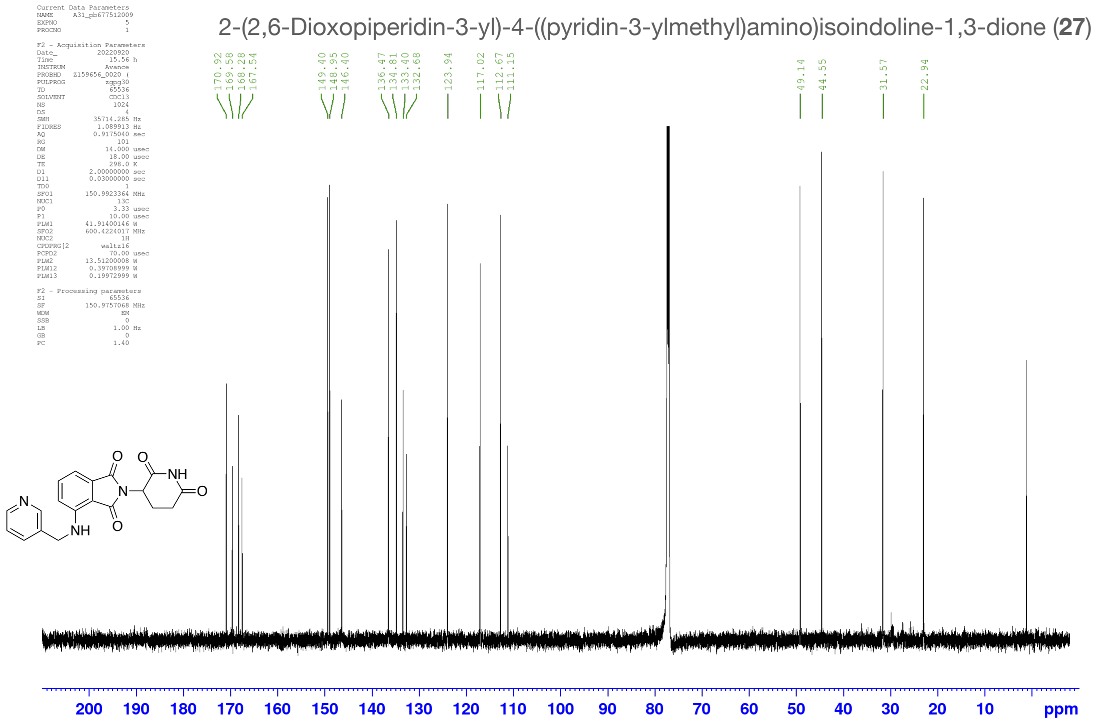
Current Data Parameters
NAME A31_pb677512009
EXPNO 5
PROCNO 1
F2 - Acquisition Parameters
Date_ 20220920
Time 15.56 h
INSTRUM Avance
PROBHD Z159656_0020 (
PULPROG zgpg30
TD 65536
SOLVENT CDCl3
NS 1024
DS 4
SWH 35714.285 Hz
FIDRES 1.089913 Hz
AQ 0.9175040 sec
RG 101
DW 14.000 usec
DE 18.00 usec
TE 298.0 K
D1 2.00000000 sec
D11 0.03000000 sec
TD0 1
SFO1 150.9923364 MHz
NUC1 13C
P0 3.33 usec
P1 10.00 usec
PLW1 41.91400146 W
SFO2 600.4224017 MHz
NUC2 1H
CPDPRG[2 waltz16
PCPD2 70.00 usec
PLW2 13.51200008 W
PLW12 0.39708999 W
PLW13 0.19972999 W
F2 - Processing parameters
SI 65536
SF 150.9757068 MHz
WDW EM
SSB 0
LB 1.00 Hz
GB 0
PC 1.40
2-(2,6-Dioxopiperidin-3-yl)-4-((pyridin-3-ylmethyl)amino)isoindoline-1,3-dione (27)
170.92
169.58
168.28
167.54
149.40
148.95
146.40
136.47
134.81
133.40
132.68
123.94
117.02
112.67
111.15
49.14
44.55
31.57
22.94
N
NH
O
O
O
N
NH
O
200
190
180
170
160
150
140
130
120
110
100
90
80
70
60
50
40
30
20
10
ppm

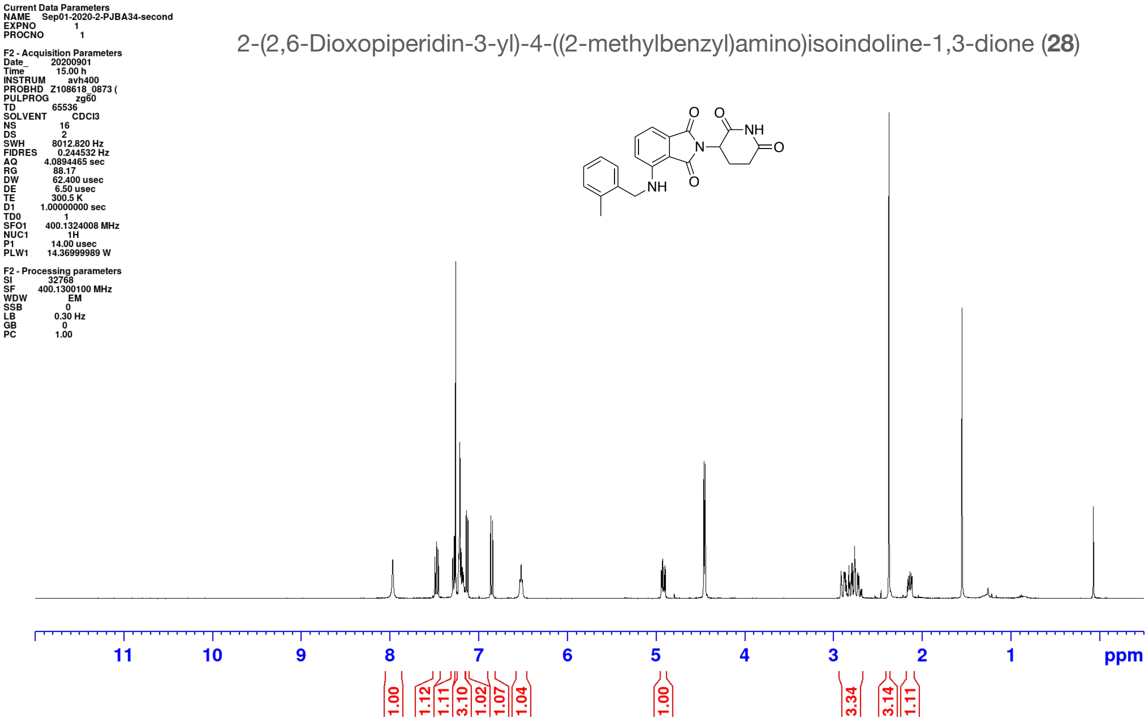
Current Data Parameters
NAME Sep01-2020-2-PJBA34-second
EXPNO 1
PROCNO 1
F2 - Acquisition Parameters
Date_ 20200901
Time 15.00 h
INSTRUM avh400
PROBHD Z108618_0873 (
PULPROG zg60
TD 65536
SOLVENT CDCl3
NS 16
DS 2
SWH 8012.820 Hz
FIDRES 0.244532 Hz
AQ 4.0894465 sec
RG 88.17
DW 62.400 usec
DE 6.50 usec
TE 300.5 K
D1 1.00000000 sec
TD0 1
SFO1 400.1324008 MHz
NUC1 1H
P1 14.00 usec
PLW1 14.36999989 W
F2 - Processing parameters
SI 32768
SF 400.1300100 MHz
WDW EM
SSB 0
LB 0.30 Hz
GB 0
PC 1.00
2-(2,6-Dioxopiperidin-3-yl)-4-((2-methylbenzyl)amino)isoindoline-1,3-dione (28)
O
O
NH
N
O
NH
O
11
10
9
8
7
6
5
4
3
2
1
ppm
1.00
1.12
1.11
3.10
1.02
1.07
1.04
1.00
3.34
3.14
1.11

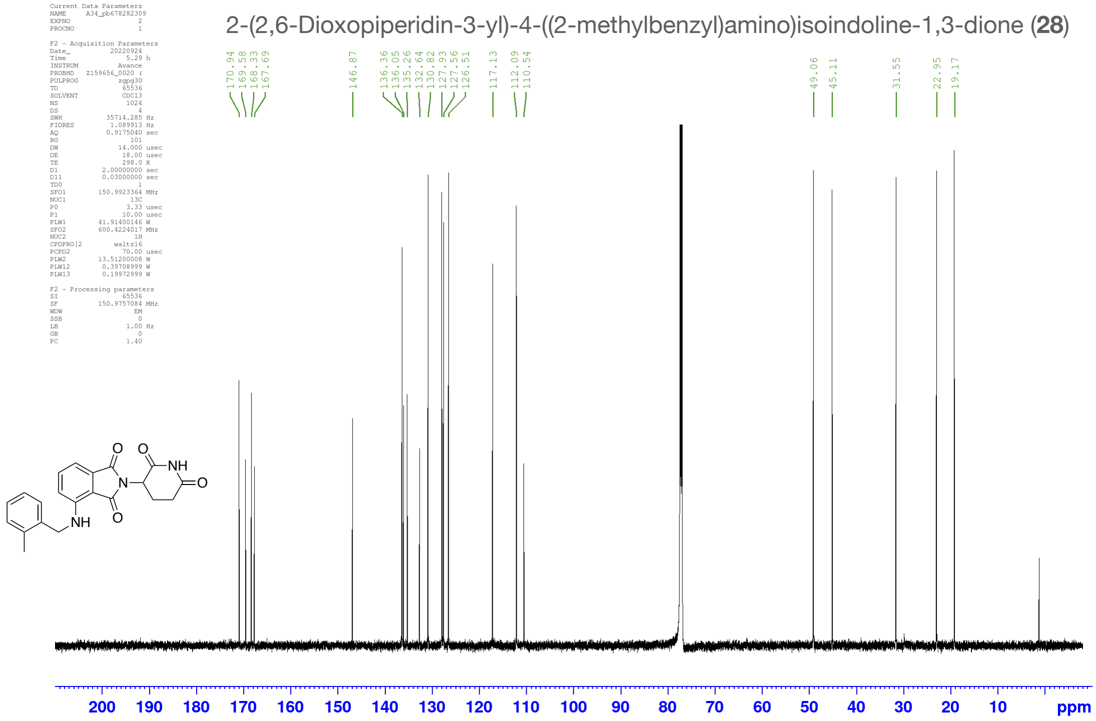
Current Data Parameters
NAME A34_pb678282309
EXPNO 2
PROCNO 1
F2 - Acquisition Parameters
Date_ 20220924
Time 5.29 h
INSTRUM Avance
PROBHD Z159656_0020 (
PULPROG zgpg30
TD 65536
SOLVENT CDCl3
NS 1024
DS 4
SWH 35714.285 Hz
FIDRES 1.089913 Hz
AQ 0.9175040 sec
RG 101
DW 14.000 usec
DE 18.00 usec
TE 298.0 K
D1 2.00000000 sec
D11 0.03000000 sec
TD0 1
SFO1 150.9923364 MHz
NUC1 13C
P0 3.33 usec
P1 10.00 usec
PLW1 41.91400146 W
SFO2 600.4224017 MHz
NUC2 1H
CPDPRG[2 waltz16
PCPD2 70.00 usec
PLW2 13.51200008 W
PLW12 0.39708999 W
PLW13 0.19972999 W
F2 - Processing parameters
SI 65536
SF 150.9757084 MHz
WDW EM
SSB 0
LB 1.00 Hz
GB 0
PC 1.40
2-(2,6-Dioxopiperidin-3-yl)-4-((2-methylbenzyl)amino)isoindoline-1,3-dione (28)
170.94
169.58
168.33
167.69
146.87
136.36
136.05
135.26
132.64
130.82
127.93
127.56
126.51
117.13
112.09
110.54
49.06
45.11
31.55
22.95
19.17
O
O
NH
N
O
NH
O
200
190
180
170
160
150
140
130
120
110
100
90
80
70
60
50
40
30
20
10
ppm

2-(2,6-Dioxopiperidin-3-yl)-4-(((4-methylpyridin-3-yl)methyl)amino)isoindoline-1,3-dione (**29**)

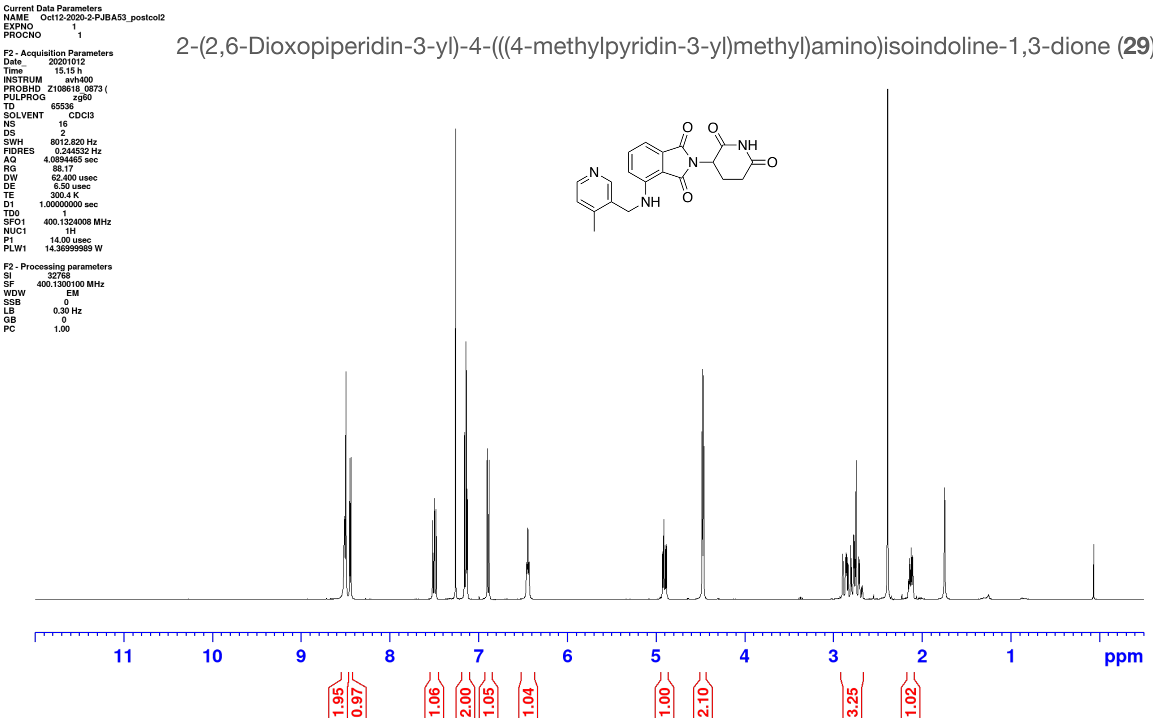

2-(2,6-Dioxopiperidin-3-yl)-4-(((4-methylpyridin-3-yl)methyl)amino)isoindoline-1,3-dione (**29**)

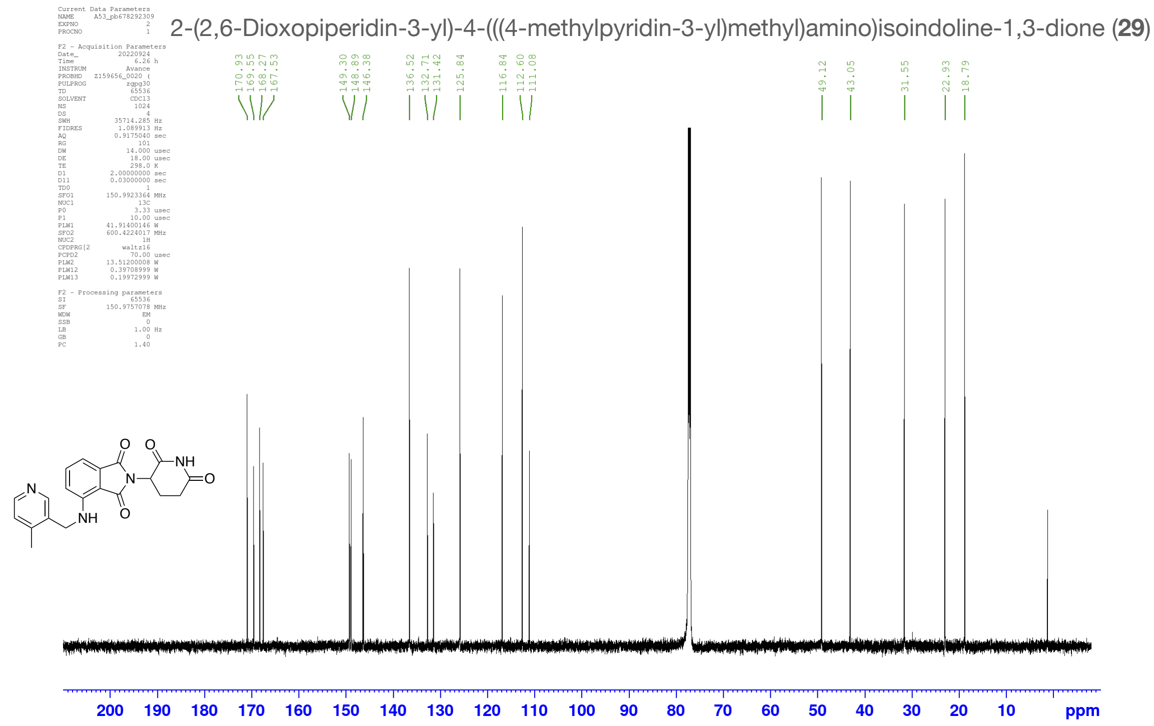

4-(Benzyloxy)-2-(2,6-dioxopiperidin-3-yl)isoindoline-1,3-dione (**30**)

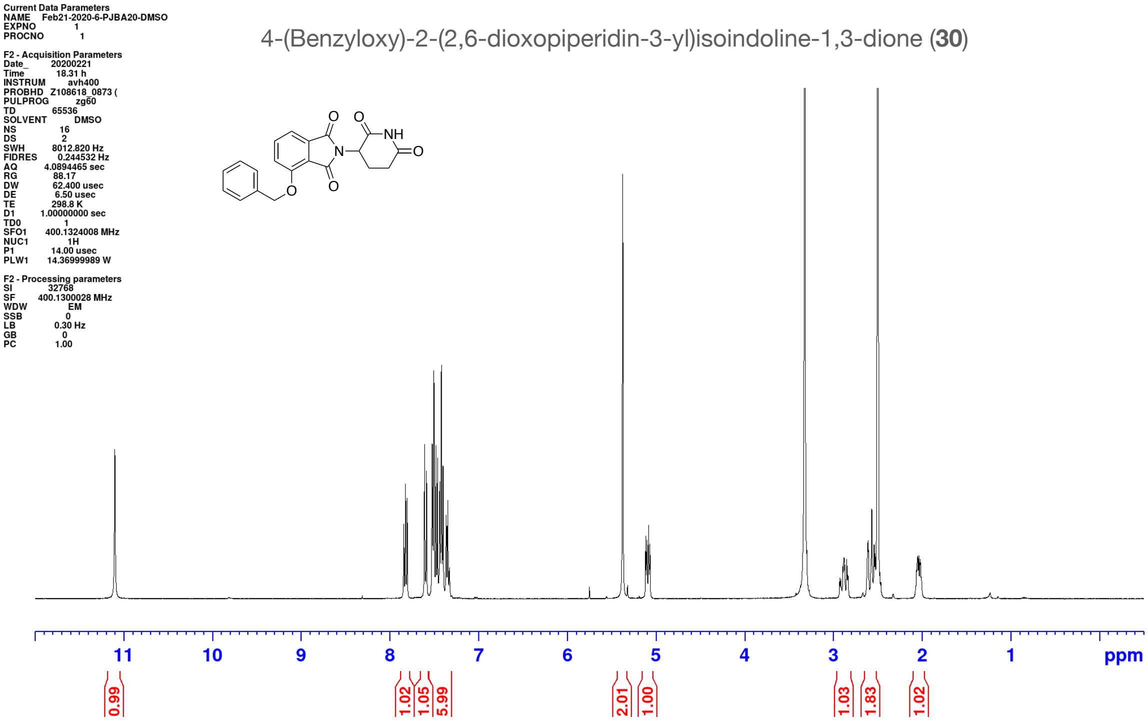

2-(2,6-Dioxopiperidin-3-yl)-4-(pyridin-3-ylmethoxy)isoindoline-1,3-dione (**31**)

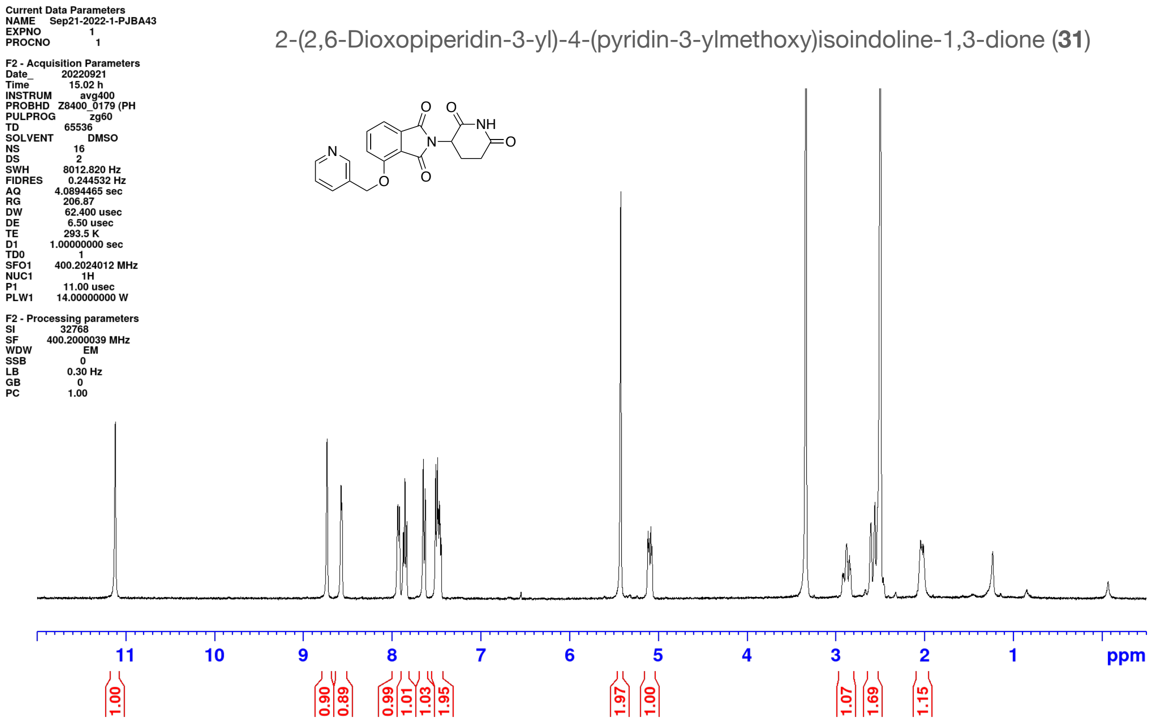

2-(2,6-Dioxopiperidin-3-yl)-4-(pyridin-3-ylmethoxy)isoindoline-1,3-dione (**31**)

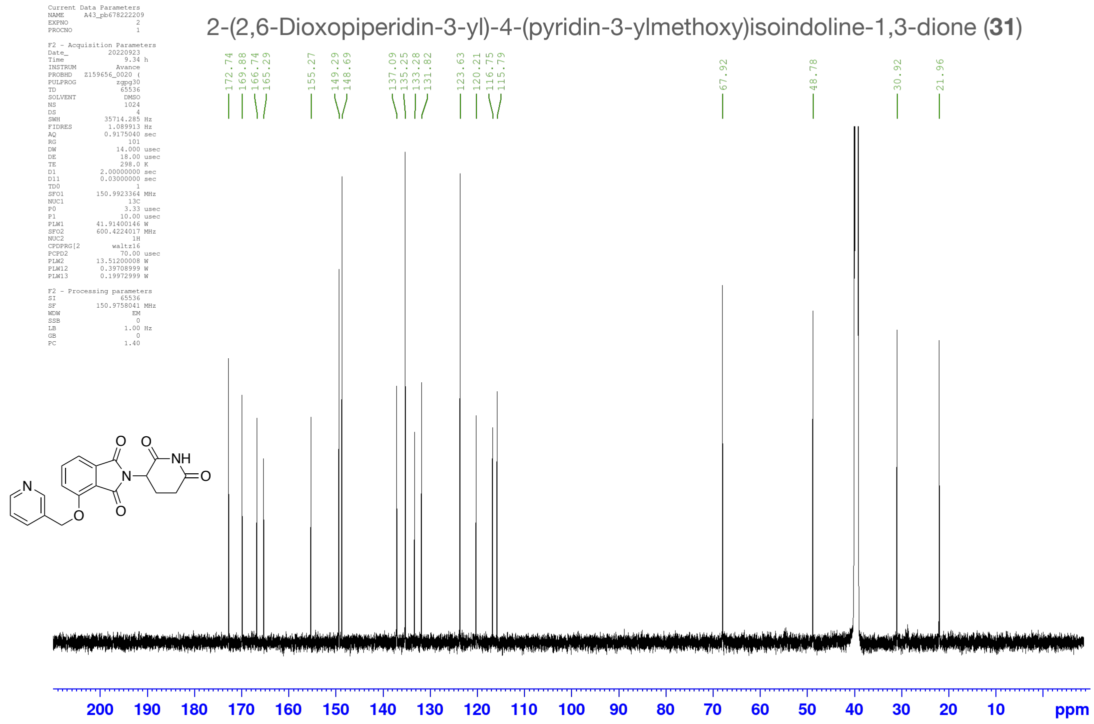

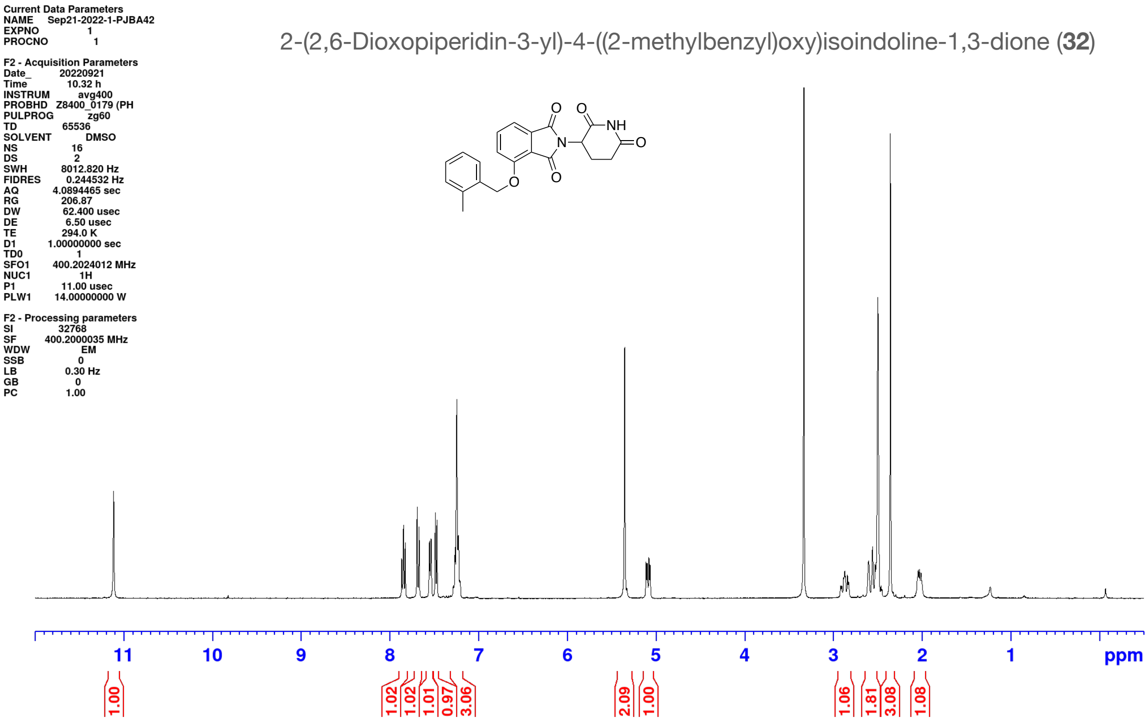
Current Data Parameters
NAME Sep21-2022-1-PJBA42
EXPNO 1
PROCNO 1
F2 - Acquisition Parameters
Date_ 20220921
Time 10.32 h
INSTRUM avg400
PROBHD Z8400_0179 (PH
PULPROG zg60
TD 65536
SOLVENT DMSO
NS 16
DS 2
SWH 8012.820 Hz
FIDRES 0.244532 Hz
AQ 4.0894465 sec
RG 206.87
DW 62.400 usec
DE 6.50 usec
TE 294.0 K
D1 1.00000000 sec
TD0 1
SFO1 400.2024012 MHz
NUC1 1H
P1 11.00 usec
PLW1 14.00000000 W
F2 - Processing parameters
SI 32768
SF 400.2000035 MHz
WDW EM
SSB 0
LB 0.30 Hz
GB 0
PC 1.00
2-(2,6-Dioxopiperidin-3-yl)-4-((2-methylbenzyl)oxy)isoindoline-1,3-dione (**32**)
O
O
NH
N
O
O
O
11 10 9 8 7 6 5 4 3 2 1 ppm
1.00
1.02 1.02 1.01 0.97 3.06
2.09 1.00
1.06 1.81 3.08 1.08

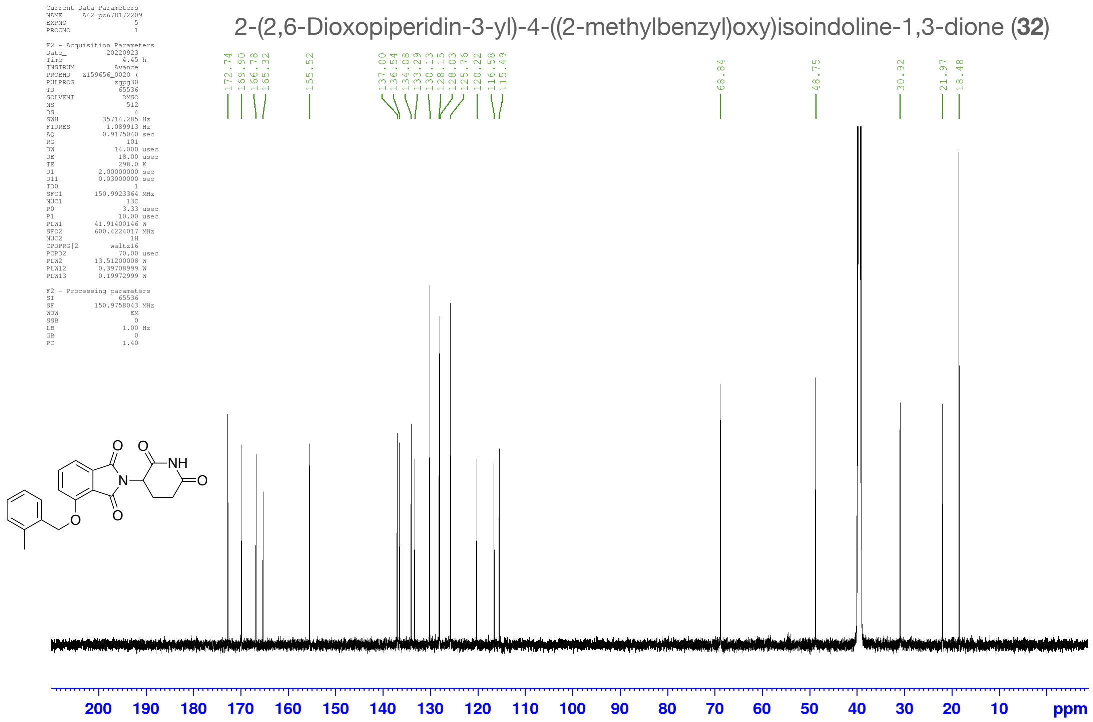
Current Data Parameters
NAME A42_pb678172209
EXPNO 5
PROCNO 1
F2 - Acquisition Parameters
Date_ 20220923
Time 4.45 h
INSTRUM Avance
PROBHD Z159656_0020 (
PULPROG zgpg30
TD 65536
SOLVENT DMSO
NS 512
DS 4
SWH 35714.285 Hz
FIDRES 1.089913 Hz
AQ 0.9175040 sec
RG 101
DW 14.000 usec
DE 18.00 usec
TE 298.0 K
D1 2.00000000 sec
D11 0.03000000 sec
TD0 1
SFO1 150.9923364 MHz
NUC1 13C
P0 3.33 usec
P1 10.00 usec
PLW1 41.91400146 W
SFO2 600.4224017 MHz
NUC2 1H
CPDPRG[2 waltz16
PCPD2 70.00 usec
PLW2 13.51200008 W
PLW12 0.39708999 W
PLW13 0.19972999 W
F2 - Processing parameters
SI 65536
SF 150.9758043 MHz
WDW EM
SSB 0
LB 1.00 Hz
GB 0
PC 1.40
2-(2,6-Dioxopiperidin-3-yl)-4-((2-methylbenzyl)oxy)isoindoline-1,3-dione (**32**)
172.74 169.90 166.78 165.32
155.52
137.00 136.54 134.08 133.29 130.13 128.15 128.03 125.76 120.22 116.58 115.49
68.84
48.75
30.92
21.97 18.48
O
O
NH
N
O
O
O
200 190 180 170 160 150 140 130 120 110 100 90 80 70 60 50 40 30 20 10 ppm

4-(Benzhydrylamino)-2-(2,6-dioxopiperidin-3-yl)isoindoline-1,3-dione (**33**)

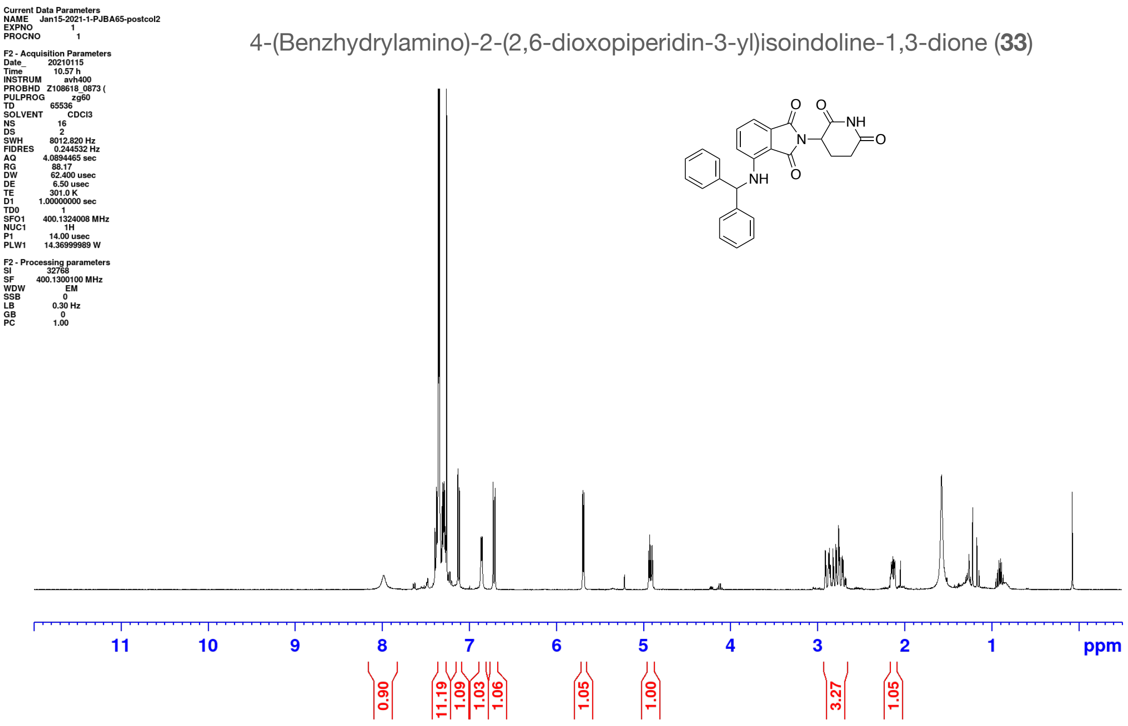

4-(Benzhydrylamino)-2-(2,6-dioxopiperidin-3-yl)isoindoline-1,3-dione (**33**)

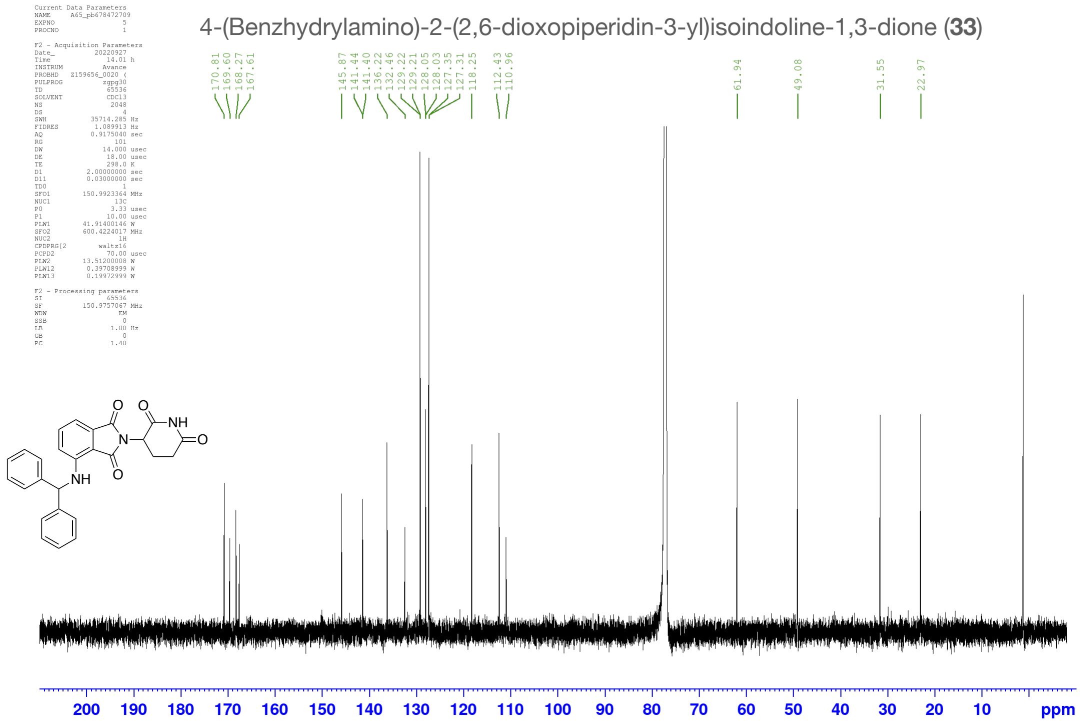

4-(Benzhydryloxy)-2-(2,6-dioxopiperidin-3-yl)isoindoline-1,3-dione (**34**)

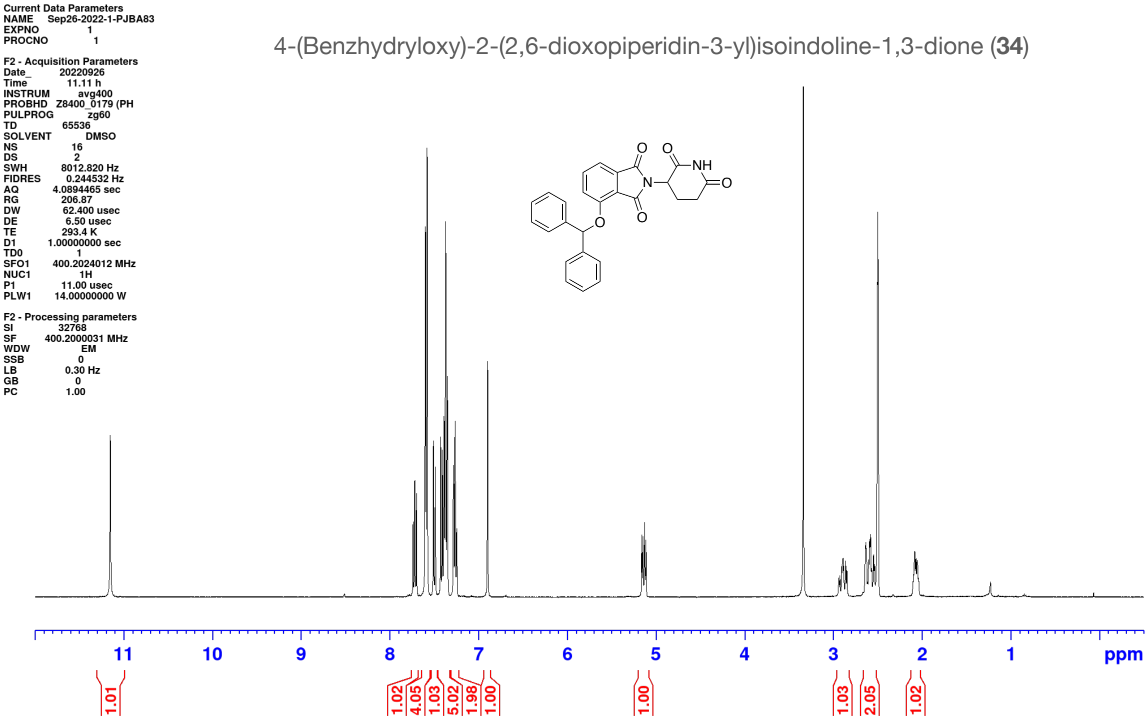

4-(Benzhydryloxy)-2-(2,6-dioxopiperidin-3-yl)isoindoline-1,3-dione (**34**)

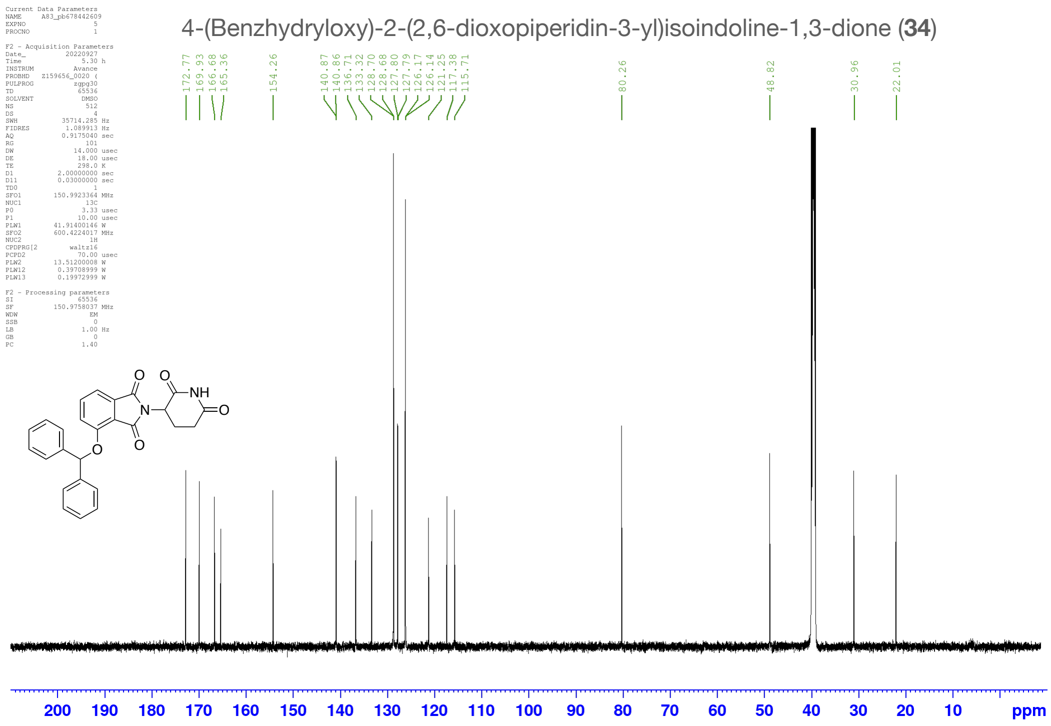

6-(2,6-Dioxopiperidin-3-yl)-5H-pyrrolo[3,4-b]pyridine-5,7(6H)-dione (**35**)

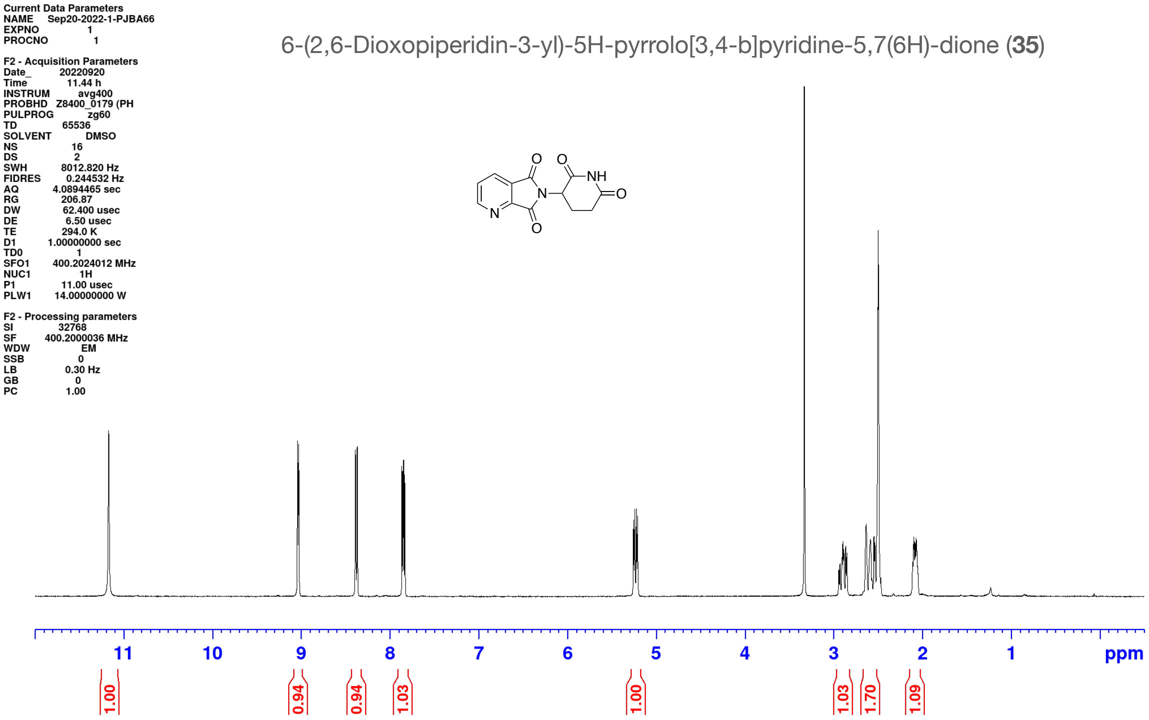

2-(2,6-Dioxopiperidin-3-yl)-1H-pyrrolo[3,4-c]pyridine-1,3(2H)-dione (**36**)

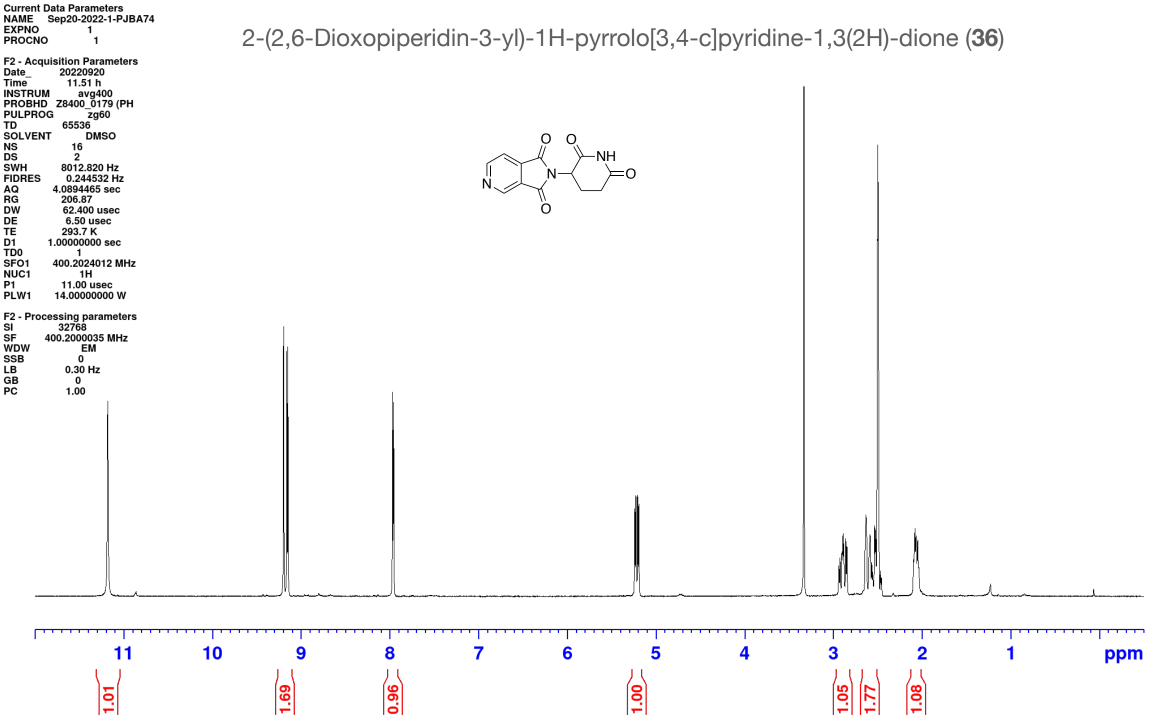

#### HPLC Traces for Compounds

2-(2,6-Dioxopiperidin-3-yl)isoindoline-1,3-dione **(1)**

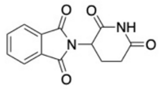

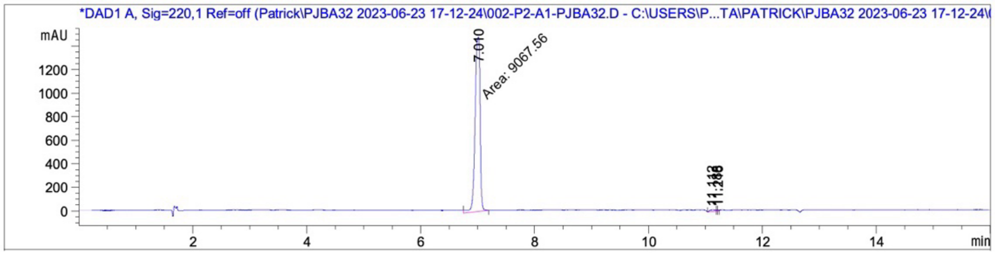

| Peak # | RetTime [min] | Type | Width [min] | Area [mAU*s] | Height [mAU] | Area % |
|---|---|---|---|---|---|---|
| 1 | 7.010 | MM | 0.1021 | 9067.55762 | 1480.10632 | 98.7473 |
| 2 | 11.112 | BV | 0.0775 | 89.24358 | 13.60625 | 0.9719 |
| 3 | 11.188 | VV | 0.0158 | 11.46414 | 9.67118 | 0.1248 |
| 4 | 11.216 | VV | 0.0207 | 14.32457 | 8.69674 | 0.1560 |

Totals : 9182.58990 1512.08049

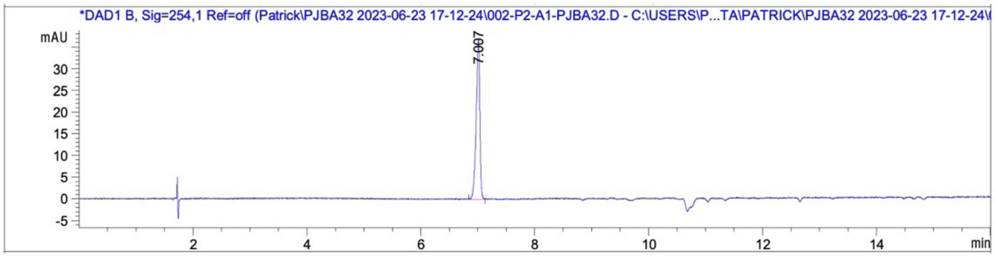

| Peak # | RetTime [min] | Type | Width [min] | Area [mAU*s] | Height [mAU] | Area % |
|---|---|---|---|---|---|---|
| 1 | 7.007 | BB | 0.0659 | 165.96967 | 36.92858 | 100.0000 |

Totals : 165.96967 36.92858

4-Amino-2-(2,6-dioxopiperidin-3-yl)isoindoline-1,3-dione **(2)**

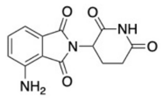

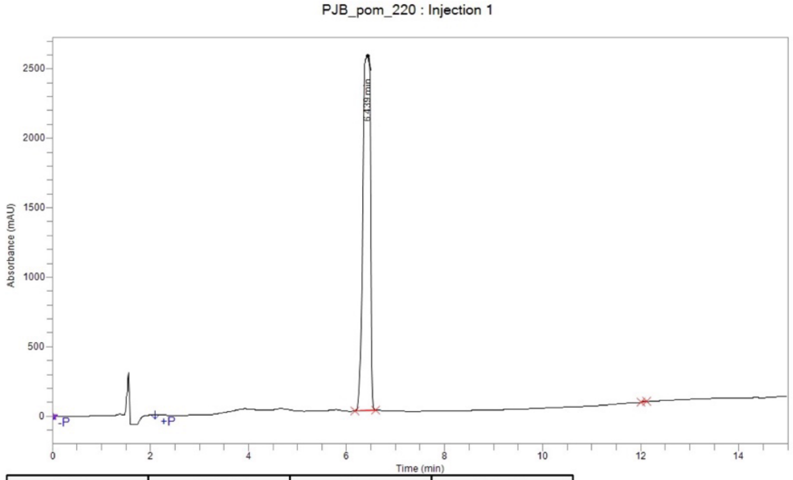

| Time | Height | Area | Area % |
|---|---|---|---|
| 6.439 | 2,553,608.8 | 27,408,154.1 | 99.93 |
| 12.077 | 4,916.0 | 19,776.4 | 0.07 |
| **Total** | | 27,427,930.4 | 100.00 |

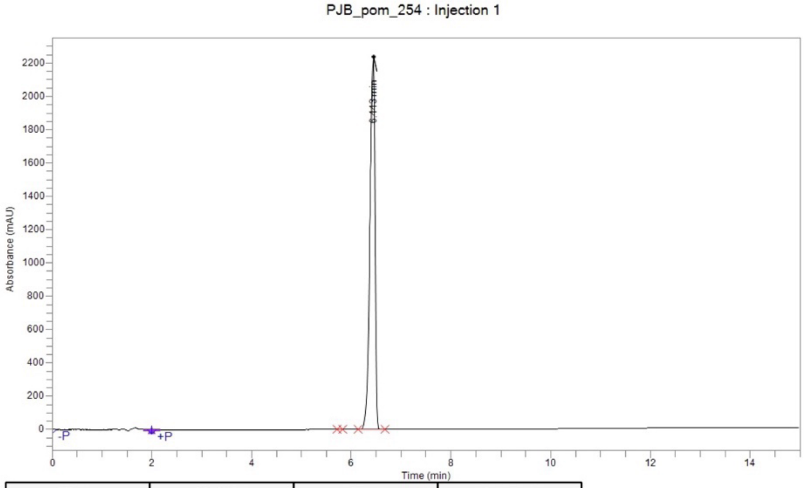

| Time | Height | Area | Area % |
|---|---|---|---|
| 5.789 | 340.4 | 1,487.8 | 0.01 |
| 6.443 | 2,239,247.0 | 15,788,772.2 | 99.99 |
| **Total** | | 15,790,260.0 | 100.00 |

5-Amino-2-(2,6-dioxopiperidin-3-yl)isoindoline-1,3-dione **(3)**

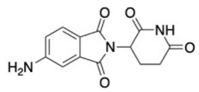

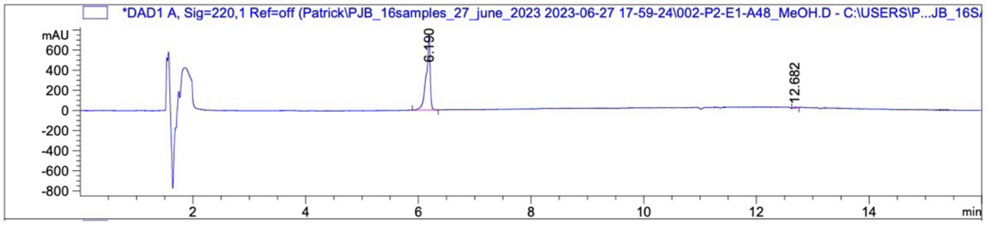

| Peak # | RetTime [min] | Type | Width [min] | Area [mAU*s] | Height [mAU] | Area % |
|---|---|---|---|---|---|---|
| 1 | 6.190 | BB | 0.0657 | 3589.78223 | 744.89728 | 98.7363 |
| 2 | 12.682 | BV | 0.0706 | 45.94411 | 7.98587 | 1.2637 |
| Totals : | | | | 3635.72633 | 752.88315 | |

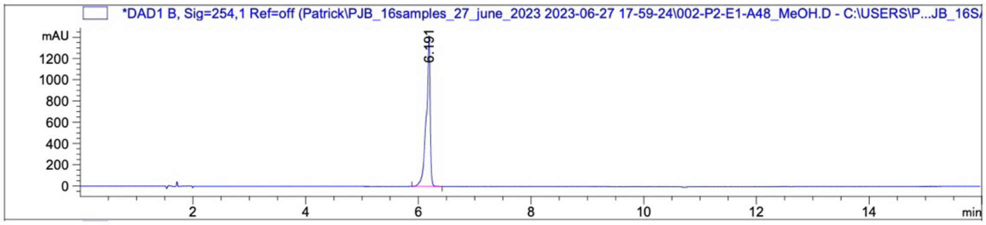

| Peak # | RetTime [min] | Type | Width [min] | Area [mAU*s] | Height [mAU] | Area % |
|---|---|---|---|---|---|---|
| 1 | 6.191 | BB | 0.0653 | 6732.34521 | 1419.69299 | 100.0000 |
| Totals : | | | | 6732.34521 | 1419.69299 | |

2-(2,6-Dioxopiperidin-3-yl)-4-hydroxyisoindoline-1,3-dione **(4)**

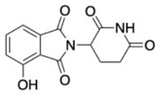

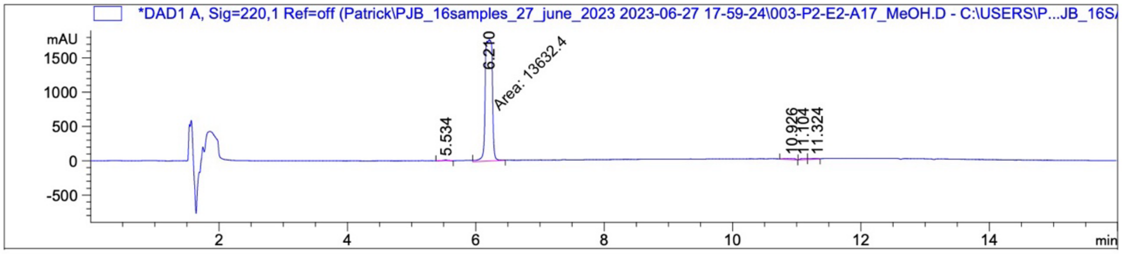

| Peak # | RetTime [min] | Type | Width [min] | Area [mAU*s] | Height [mAU] | Area % |
|---|---|---|---|---|---|---|
| 1 | 5.534 | BB | 0.0646 | 65.99696 | 12.70768 | 0.4697 |
| 2 | 6.210 | MM | 0.1278 | 1.36324e4 | 1777.73010 | 97.0149 |
| 3 | 10.926 | BB | 0.1220 | 141.60933 | 13.88067 | 1.0078 |
| 4 | 11.104 | BV | 0.0815 | 105.82807 | 15.42453 | 0.7531 |
| 5 | 11.324 | VV | 0.1264 | 106.02243 | 9.86394 | 0.7545 |
| Totals : | | | | 1.40519e4 | 1829.60693 | |

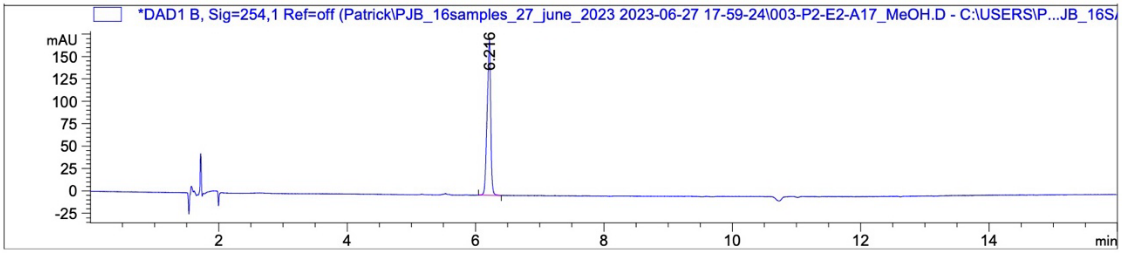

| Peak # | RetTime [min] | Type | Width [min] | Area [mAU*s] | Height [mAU] | Area % |
|---|---|---|---|---|---|---|
| 1 | 6.216 | BB | 0.0554 | 675.15735 | 173.32211 | 100.0000 |
| Totals : | | | | 675.15735 | 173.32211 | |

2-(2,6-Dioxopiperidin-3-yl)-5-hydroxyisoindoline-1,3-dione **(5)**

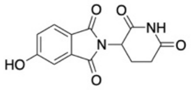

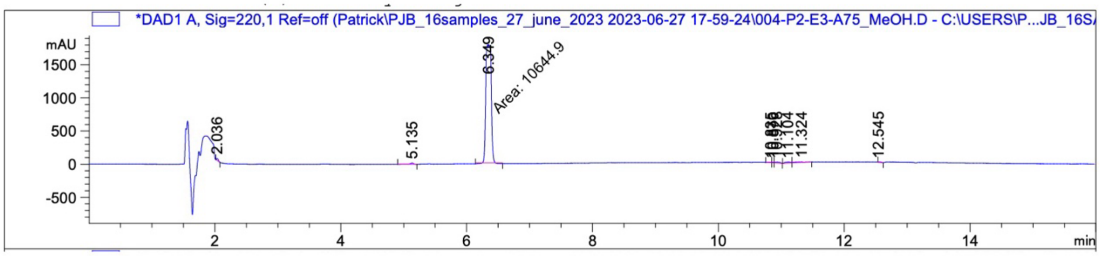

| Peak # | RetTime [min] | Type | Width [min] | Area [mAU*s] | Height [mAU] | Area % |
|---|---|---|---|---|---|---|
| 1 | 2.036 | BB | 0.0318 | 63.73929 | 32.55661 | 0.5708 |
| 2 | 5.135 | BB | 0.0519 | 80.56284 | 20.23950 | 0.7214 |
| 3 | 6.349 | MM | 0.0991 | 1.06449e4 | 1791.00110 | 95.3208 |
| 4 | 10.835 | BV | 0.0522 | 27.88848 | 7.70735 | 0.2497 |
| 5 | 10.870 | VV | 0.0245 | 18.61944 | 9.87828 | 0.1667 |
| 6 | 10.928 | VB | 0.0819 | 87.11492 | 13.66248 | 0.7801 |
| 7 | 11.104 | BV | 0.0830 | 105.74609 | 15.13966 | 0.9469 |
| 8 | 11.324 | VV R | 0.1459 | 110.80653 | 9.01607 | 0.9922 |
| 9 | 12.545 | VB | 0.0465 | 28.06925 | 7.27520 | 0.2513 |
| Totals : | | | | 1.11674e4 | 1906.47626 | |

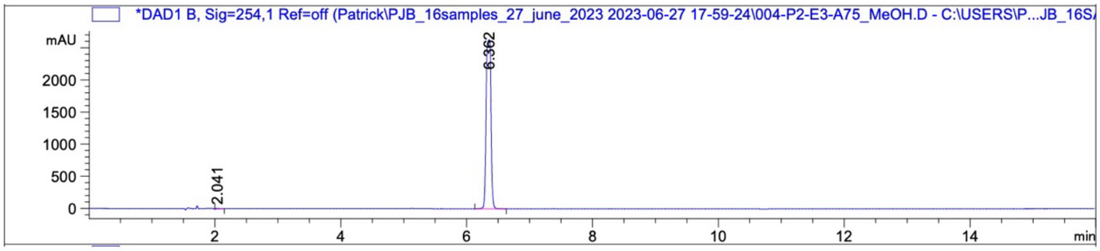

| Peak # | RetTime [min] | Type | Width [min] | Area [mAU*s] | Height [mAU] | Area % |
|---|---|---|---|---|---|---|
| 1 | 2.041 | BB | 0.0587 | 47.90619 | 10.50316 | 0.3428 |
| 2 | 6.362 | BB | 0.0638 | 1.39278e4 | 2630.26196 | 99.6572 |
| Totals : | | | | 1.39757e4 | 2640.76512 | |

2-(2,6-Dioxopiperidin-3-yl)-4-methoxyisoindoline-1,3-dione **(6)**

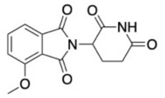

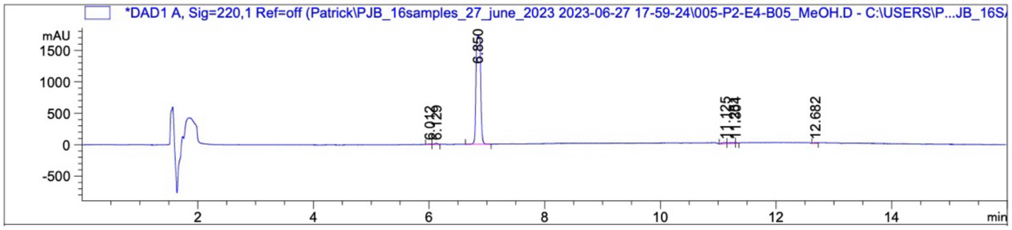

| Peak # | RetTime [min] | Type | Width [min] | Area [mAU*s] | Height [mAU] | Area % |
|---|---|---|---|---|---|---|
| 1 | 6.012 | BV | 0.0440 | 25.54466 | 7.36038 | 0.2574 |
| 2 | 6.129 | VB | 0.0474 | 57.92704 | 15.93125 | 0.5838 |
| 3 | 6.850 | BB | 0.0661 | 9592.31934 | 1717.97754 | 96.6734 |
| 4 | 11.125 | BV | 0.0756 | 93.57810 | 14.94160 | 0.9431 |
| 5 | 11.281 | VV | 0.1129 | 89.21191 | 9.51512 | 0.8991 |
| 6 | 11.304 | VB | 0.0268 | 18.89025 | 8.87971 | 0.1904 |
| 7 | 12.682 | BV | 0.0587 | 44.93058 | 9.40365 | 0.4528 |
| Totals : | | | | 9922.40189 | 1784.00926 | |

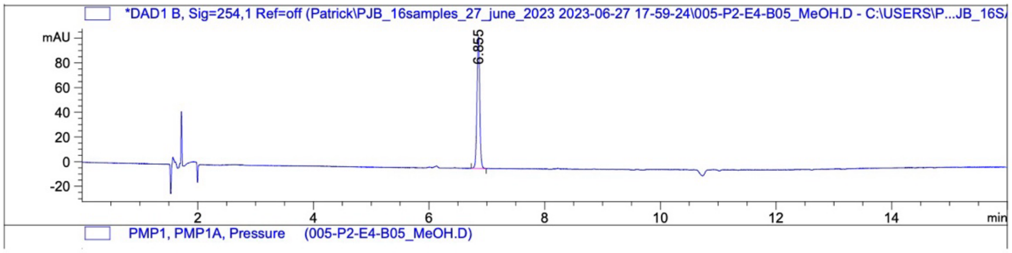

Signal 2: DAD1 B, Sig=254,1 Ref=off
Signal has been modified after loading from rawdata file!

| Peak # | RetTime [min] | Type | Width [min] | Area [mAU*s] | Height [mAU] | Area % |
|---|---|---|---|---|---|---|
| 1 | 6.855 | BB | 0.0527 | 358.38605 | 106.73189 | 100.0000 |
| Totals : | | | | 358.38605 | 106.73189 | |

2-(2,6-Dioxopiperidin-3-yl)-5-methoxyisoindoline-1,3-dione **(7)**

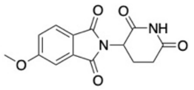

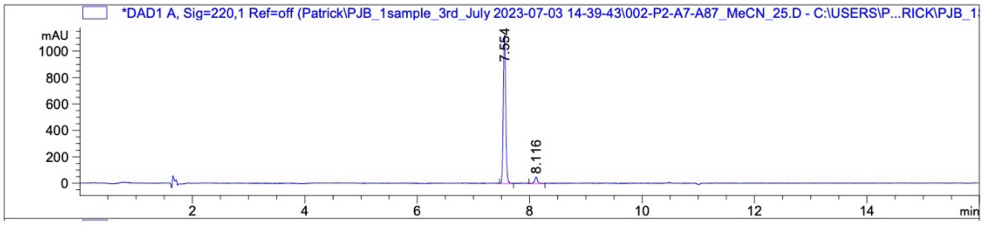

| Peak # | RetTime [min] | Type | Width [min] | Area [mAU*s] | Height [mAU] | Area % |
|---|---|---|---|---|---|---|
| 1 | 7.554 | VB R | 0.0463 | 3345.32910 | 1122.03931 | 95.5459 |
| 2 | 8.116 | BB | 0.0463 | 155.95166 | 48.19432 | 4.4541 |

Totals : 3501.28076 1170.23363

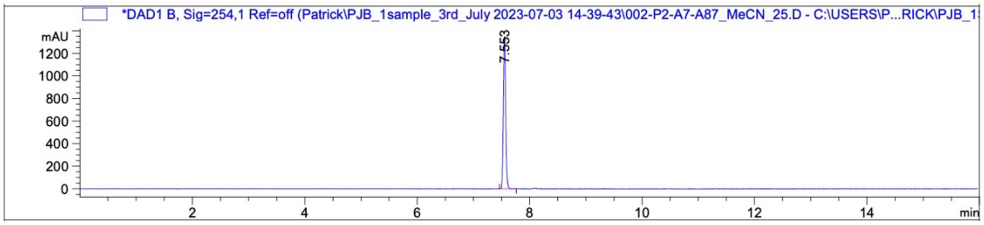

| Peak # | RetTime [min] | Type | Width [min] | Area [mAU*s] | Height [mAU] | Area % |
|---|---|---|---|---|---|---|
| 1 | 7.553 | BB | 0.0436 | 3819.21582 | 1347.94446 | 100.0000 |

Totals : 3819.21582 1347.94446

2-(2,6-Dioxopiperidin-3-yl)-4-fluoroisoindoline-1,3-dione **(8)**

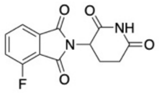

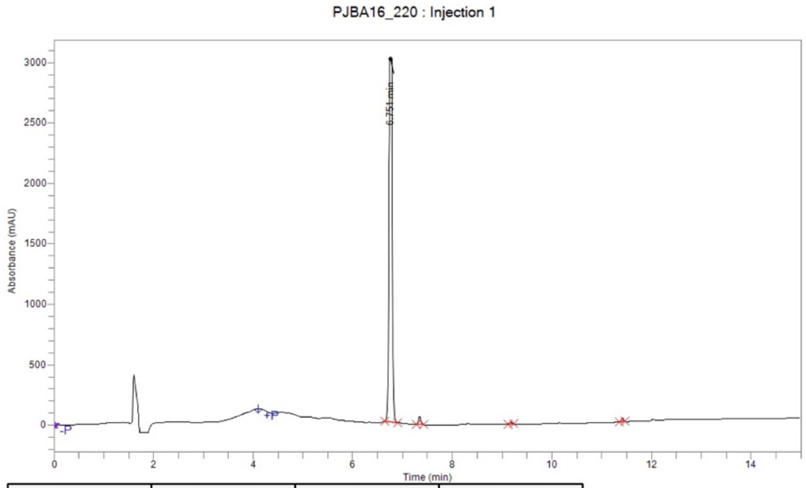

| Time | Height | Area | Area % |
|---|---|---|---|
| 6.751 | 3,028,627.9 | 14,117,851.2 | 97.52 |
| 7.346 | 63,271.4 | 219,745.1 | 1.52 |
| 9.173 | 16,466.0 | 52,777.1 | 0.36 |
| 11.416 | 28,380.4 | 86,527.3 | 0.60 |
| **Total** | | 14,476,900.7 | 100.00 |

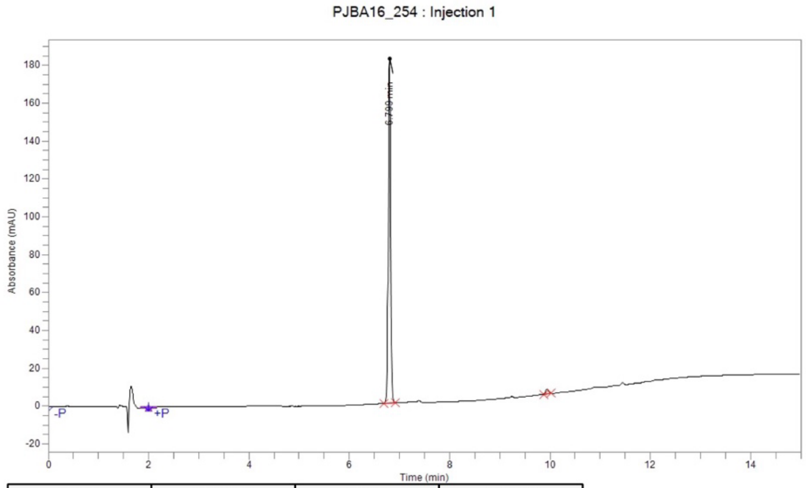

| Time | Height | Area | Area % |
|---|---|---|---|
| 6.799 | 182,217.2 | 630,377.0 | 98.51 |
| 9.945 | 2,444.7 | 9,551.0 | 1.49 |
| **Total** | | 639,928.0 | 100.00 |

2-(2,6-Dioxopiperidin-3-yl)-5-fluoroisoindoline-1,3-dione **(9)**

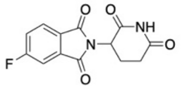

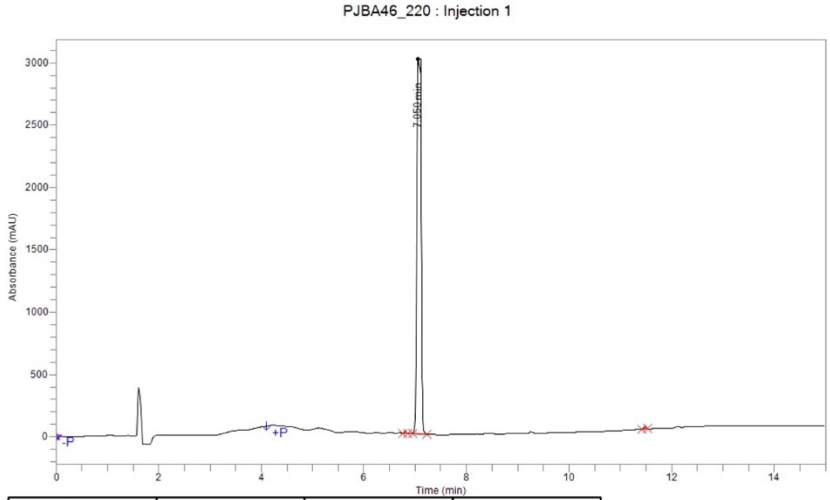

| Time | Height | Area | Area % |
|---|---|---|---|
| 6.809 | 13,303.9 | 39,412.8 | 0.22 |
| 7.050 | 3,039,466.0 | 18,002,760.7 | 99.45 |
| 11.482 | 20,006.3 | 60,644.8 | 0.34 |
| **Total** | | 18,102,818.3 | 100.00 |

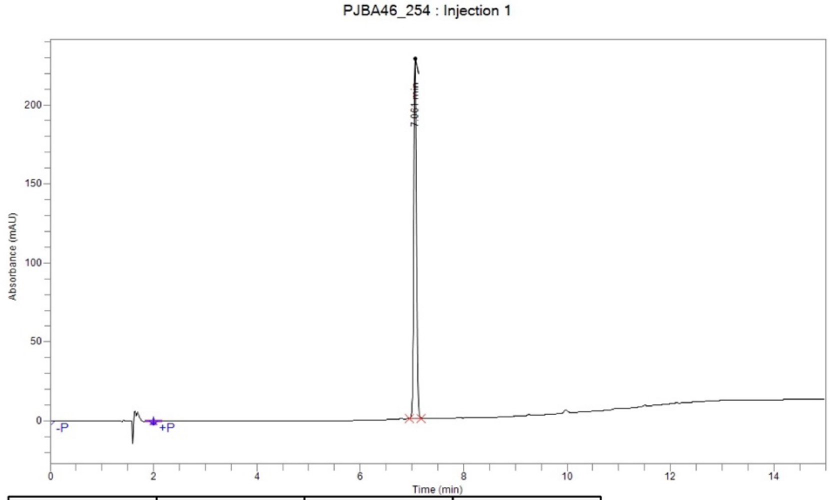

| Time | Height | Area | Area % |
|---|---|---|---|
| 7.061 | 227,924.9 | 796,290.9 | 100.00 |
| **Total** | | 796,290.9 | 100.00 |

4-Chloro-2-(2,6-dioxopiperidin-3-yl)isoindoline-1,3-dione **(10)**

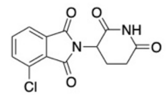

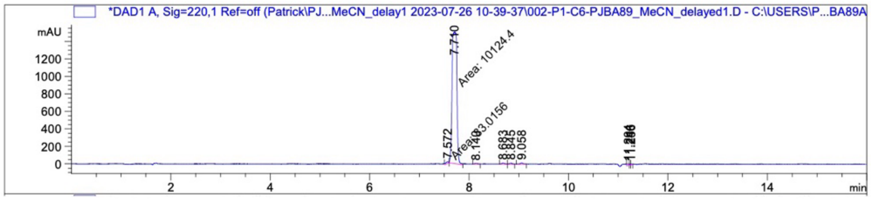

| Peak # | RetTime [min] | Type | Width [min] | Area [mAU*s] | Height [mAU] | Area % |
|---|---|---|---|---|---|---|
| 1 | 7.572 | MM | 0.0716 | 83.01558 | 19.31462 | 0.7948 |
| 2 | 7.710 | MM | 0.1120 | 1.01244e4 | 1506.12683 | 96.9367 |
| 3 | 8.140 | BB | 0.0456 | 28.66743 | 7.59032 | 0.2745 |
| 4 | 8.683 | BV | 0.0456 | 39.57912 | 10.96288 | 0.3790 |
| 5 | 8.845 | VB | 0.0483 | 35.57435 | 10.30676 | 0.3406 |
| 6 | 9.058 | BV R | 0.0457 | 46.73374 | 12.61280 | 0.4475 |
| 7 | 11.204 | VV | 0.0401 | 47.28849 | 14.10421 | 0.4528 |
| 8 | 11.246 | VV | 0.0244 | 19.43041 | 9.87937 | 0.1860 |
| 9 | 11.256 | VV | 0.0257 | 19.65645 | 9.27698 | 0.1882 |
| Totals : | | | | 1.04443e4 | 1600.17477 | |

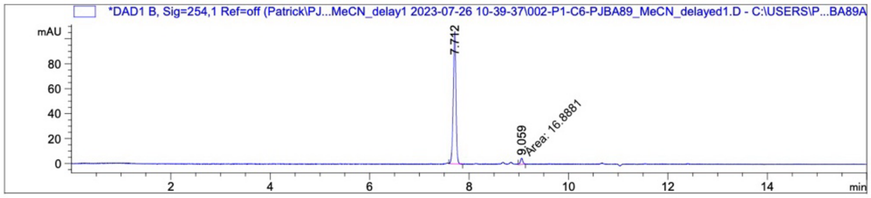

| Peak # | RetTime [min] | Type | Width [min] | Area [mAU*s] | Height [mAU] | Area % |
|---|---|---|---|---|---|---|
| 1 | 7.712 | BB | 0.0580 | 400.58704 | 106.22004 | 95.9547 |
| 2 | 9.059 | MM | 0.0571 | 16.88810 | 4.93309 | 4.0453 |
| Totals : | | | | 417.47514 | 111.15313 | |

5-Chloro-2-(2,6-dioxopiperidin-3-yl)isoindoline-1,3-dione **(11)**

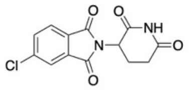

PJBA86_220_actual : Injection 1

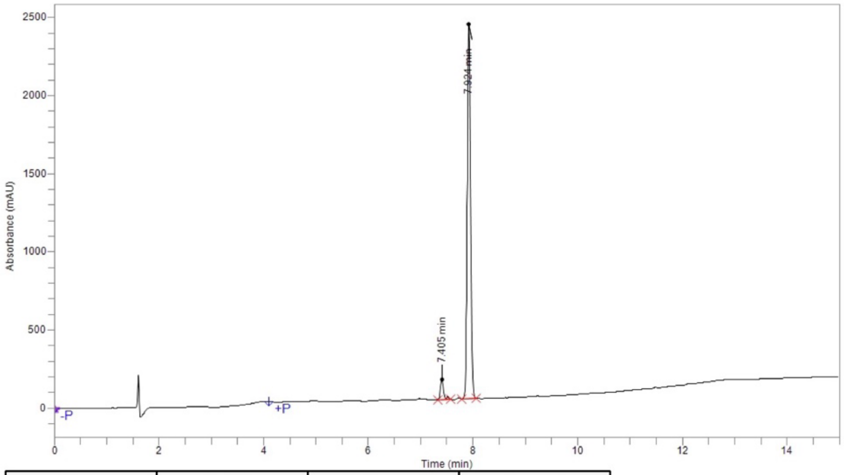

| Time | Height | Area | Area % |
|---|---|---|---|
| 7.405 | 129,530.5 | 482,204.2 | 4.35 |
| 7.518 | 16,496.3 | 52,657.9 | 0.48 |
| 7.924 | 2,400,093.6 | 10,540,490.2 | 95.17 |
| **Total** | | 11,075,352.3 | 100.00 |

PJBA86_254 : Injection 1

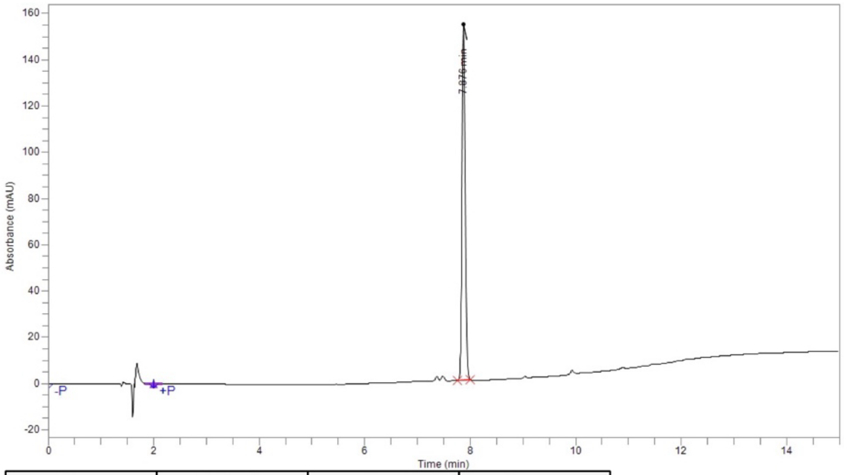

| Time | Height | Area | Area % |
|---|---|---|---|
| 7.876 | 153,998.7 | 642,358.4 | 100.00 |
| **Total** | | 642,358.4 | 100.00 |

4-Bromo-2-(2,6-dioxopiperidin-3-yl)isoindoline-1,3-dione **(12)**

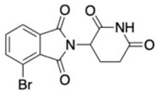

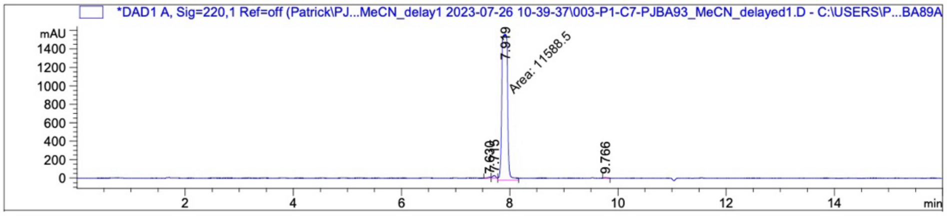

| Peak # | RetTime [min] | Type | Width [min] | Area [mAU*s] | Height [mAU] | Area % |
|---|---|---|---|---|---|---|
| 1 | 7.630 | BV | 0.0488 | 47.52709 | 11.97518 | 0.4039 |
| 2 | 7.715 | VB | 0.0466 | 107.22497 | 28.36489 | 0.9112 |
| 3 | 7.919 | MM | 0.1219 | 1.15885e4 | 1584.42163 | 98.4798 |
| 4 | 9.766 | BB | 0.0395 | 24.13687 | 7.50915 | 0.2051 |
| Totals : | | | | 1.17674e4 | 1632.27086 | |

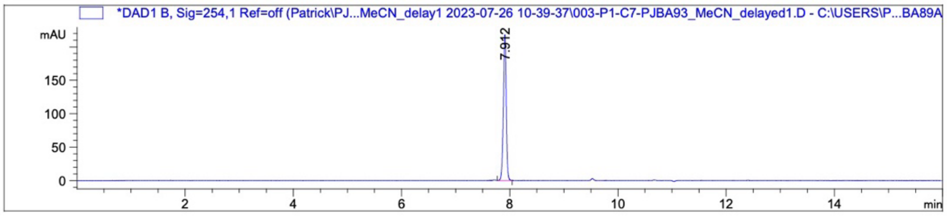

| Peak # | RetTime [min] | Type | Width [min] | Area [mAU*s] | Height [mAU] | Area % |
|---|---|---|---|---|---|---|
| 1 | 7.912 | BB | 0.0561 | 796.89526 | 218.08075 | 100.0000 |
| Totals : | | | | 796.89526 | 218.08075 | |

5-Bromo-2-(2,6-dioxopiperidin-3-yl)isoindoline-1,3-dione **(13)**

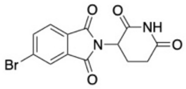

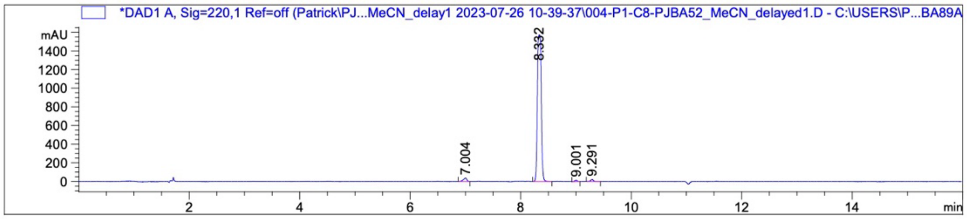

| Peak # | RetTime [min] | Type | Width [min] | Area [mAU*s] | Height [mAU] | Area % |
|---|---|---|---|---|---|---|
| 1 | 7.004 | VB R | 0.0601 | 169.77148 | 37.28113 | 2.1269 |
| 2 | 8.332 | BB | 0.0589 | 7684.37988 | 1575.89148 | 96.2706 |
| 3 | 9.001 | BB | 0.0400 | 47.47767 | 14.40140 | 0.5948 |
| 4 | 9.291 | BV R | 0.0449 | 80.43294 | 22.11587 | 1.0077 |
| Totals : | | | | 7982.06198 | 1649.68988 | |

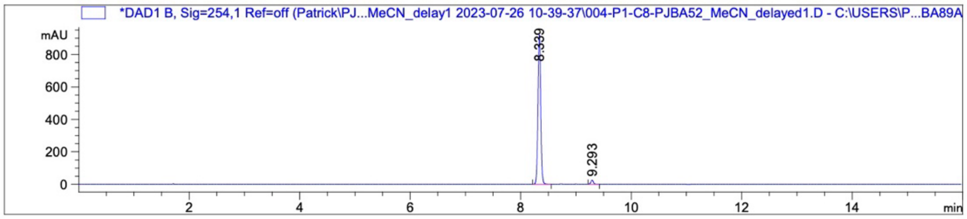

| Peak # | RetTime [min] | Type | Width [min] | Area [mAU*s] | Height [mAU] | Area % |
|---|---|---|---|---|---|---|
| 1 | 8.339 | BB | 0.0550 | 3232.09131 | 920.24182 | 97.5726 |
| 2 | 9.293 | BB | 0.0479 | 80.40845 | 25.42469 | 2.4274 |
| Totals : | | | | 3312.49976 | 945.66651 | |

2-(2,6-Dioxopiperidin-3-yl)-4-iodoisoindoline-1,3-dione **(14)**

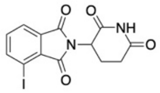

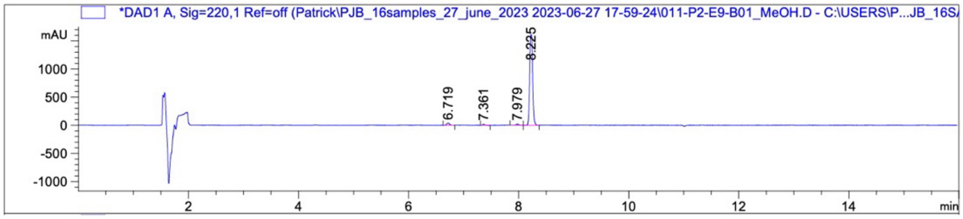

| Peak # | RetTime [min] | Type | Width [min] | Area [mAU*s] | Height [mAU] | Area % |
|---|---|---|---|---|---|---|
| 1 | 6.719 | BB | 0.0445 | 133.82944 | 37.15325 | 2.0950 |
| 2 | 7.361 | BB | 0.0403 | 49.76325 | 15.35464 | 0.7790 |
| 3 | 7.979 | BB | 0.0524 | 77.25655 | 20.54155 | 1.2094 |
| 4 | 8.225 | BB | 0.0468 | 6127.28809 | 1613.76160 | 95.9167 |

Totals : 6388.13733 1686.81104

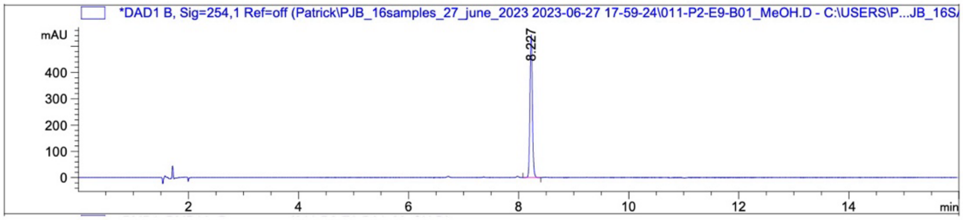

| Peak # | RetTime [min] | Type | Width [min] | Area [mAU*s] | Height [mAU] | Area % |
|---|---|---|---|---|---|---|
| 1 | 8.227 | BB | 0.0480 | 1676.24133 | 543.46442 | 100.0000 |

Totals : 1676.24133 543.46442

2-(2,6-Dioxopiperidin-3-yl)-5-iodoisoindoline-1,3-dione **(15)**

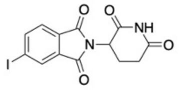

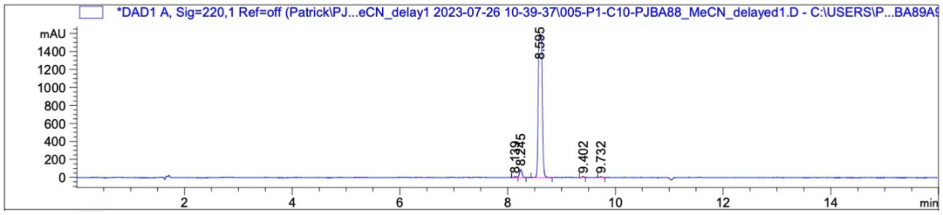

| Peak # | RetTime [min] | Type | Width [min] | Area [mAU*s] | Height [mAU] | Area % |
|---|---|---|---|---|---|---|
| 1 | 8.139 | BV E | 0.0379 | 31.36179 | 10.05772 | 0.3685 |
| 2 | 8.245 | VV R | 0.0507 | 303.46432 | 89.25816 | 3.5662 |
| 3 | 8.595 | VB R | 0.0608 | 8107.98242 | 1594.07239 | 95.2812 |
| 4 | 9.402 | BV | 0.0366 | 35.78897 | 11.90493 | 0.4206 |
| 5 | 9.732 | VB | 0.0454 | 30.93200 | 8.13432 | 0.3635 |
| Totals : | | | | 8509.52951 | 1713.42752 | |

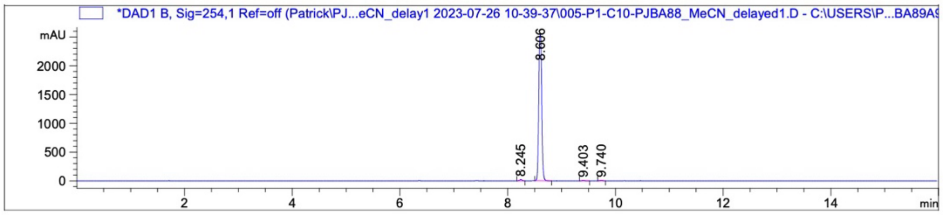

| Peak # | RetTime [min] | Type | Width [min] | Area [mAU*s] | Height [mAU] | Area % |
|---|---|---|---|---|---|---|
| 1 | 8.245 | BB | 0.0493 | 65.21466 | 19.61108 | 0.6621 |
| 2 | 8.606 | BB | 0.0543 | 9735.32910 | 2534.51929 | 98.8364 |
| 3 | 9.403 | BB | 0.0462 | 24.64587 | 7.71999 | 0.2502 |
| 4 | 9.740 | BB | 0.0482 | 24.74964 | 7.86426 | 0.2513 |
| Totals : | | | | 9849.93928 | 2569.71462 | |

2-(2,6-Dioxopiperidin-3-yl)-4-methylisoindoline-1,3-dione **(16)**

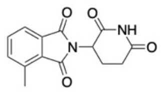

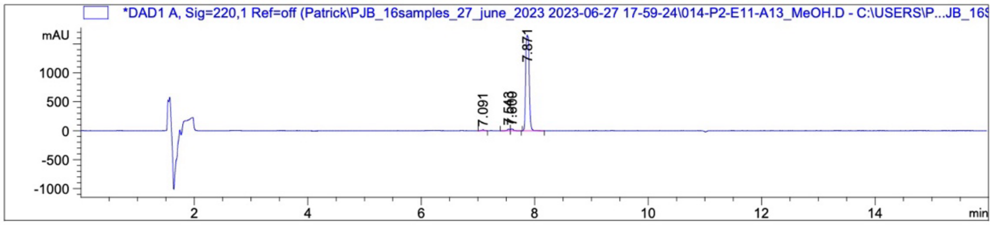

| Peak # | RetTime [min] | Type | Width [min] | Area [mAU*s] | Height [mAU] | Area % |
|---|---|---|---|---|---|---|
| 1 | 7.091 | BB | 0.0601 | 48.40011 | 12.24416 | 0.6610 |
| 2 | 7.543 | BV | 0.0466 | 84.10289 | 26.80878 | 1.1486 |
| 3 | 7.600 | VB | 0.0487 | 106.54169 | 32.09800 | 1.4550 |
| 4 | 7.871 | BB | 0.0705 | 7083.42627 | 1636.31177 | 96.7355 |
| Totals : | | | | 7322.47095 | 1707.46271 | |

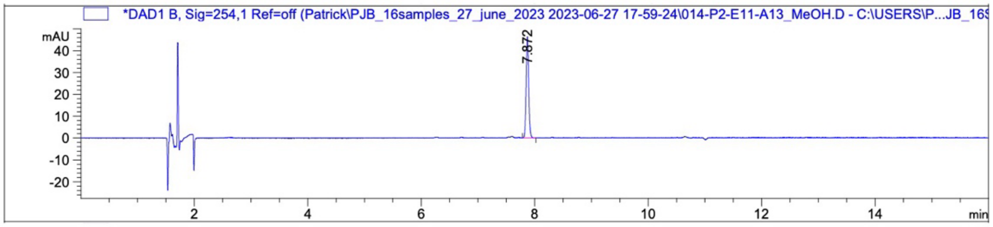

| Peak # | RetTime [min] | Type | Width [min] | Area [mAU*s] | Height [mAU] | Area % |
|---|---|---|---|---|---|---|
| 1 | 7.872 | BB | 0.0472 | 144.07738 | 46.43835 | 100.0000 |
| Totals : | | | | 144.07738 | 46.43835 | |

2-(2,6-Dioxopiperidin-3-yl)-5-methylisoindoline-1,3-dione **(17)**

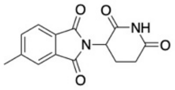

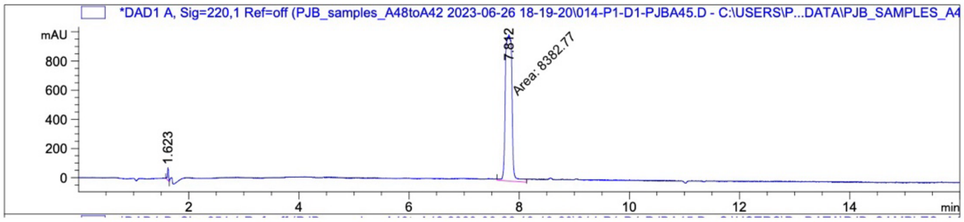

| Peak # | RetTime [min] | Type | Width [min] | Area [mAU*s] | Height [mAU] | Area % |
|---|---|---|---|---|---|---|
| 1 | 1.623 | BB | 0.0208 | 108.92145 | 85.71591 | 1.2827 |
| 2 | 7.812 | MM | 0.1389 | 8382.77344 | 1005.74152 | 98.7173 |

Totals : 8491.69489 1091.45743

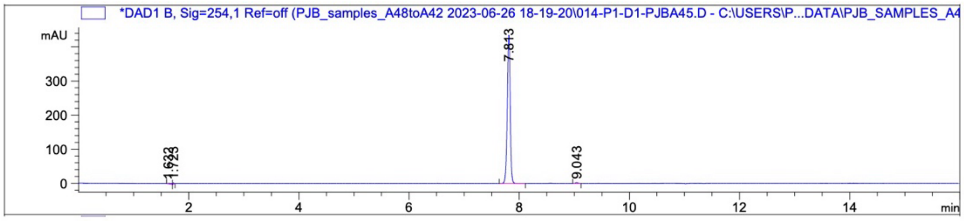

| Peak # | RetTime [min] | Type | Width [min] | Area [mAU*s] | Height [mAU] | Area % |
|---|---|---|---|---|---|---|
| 1 | 1.632 | BB | 0.0811 | 11.08154 | 1.72119 | 0.6859 |
| 2 | 1.723 | BB | 0.0229 | 5.64052 | 3.67605 | 0.3491 |
| 3 | 7.813 | BB | 0.0567 | 1589.70142 | 434.66150 | 98.3937 |
| 4 | 9.043 | BB | 0.0501 | 9.23091 | 2.82392 | 0.5713 |

Totals : 1615.65439 442.88267

2-(2,6-Dioxopiperidin-3-yl)-5-(trifluoromethyl)isoindoline-1,3-dione **(18)**

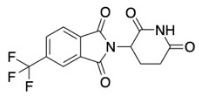

A84_220_12_7_2021 : Injection 1

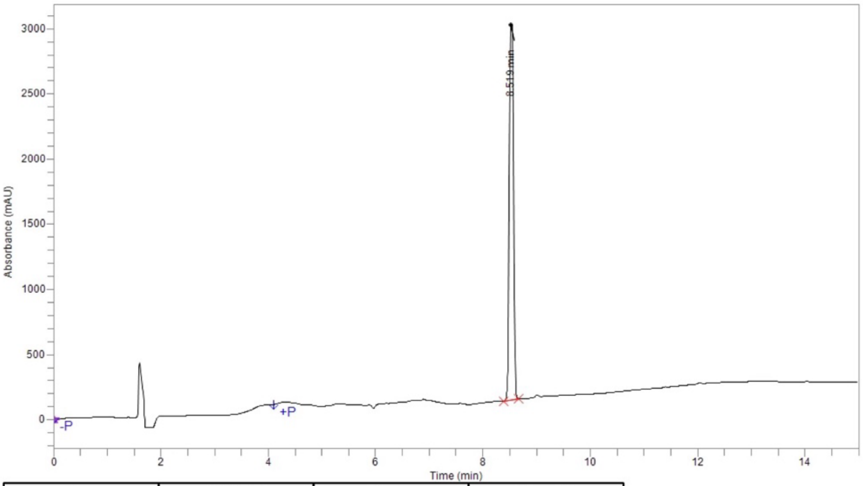

| Time | Height | Area | Area % |
|---|---|---|---|
| 8.519 | 2,921,498.9 | 15,168,796.2 | 100.00 |
| **Total** | | 15,168,796.2 | 100.00 |

A84_254_12_7_2021 : Injection 1

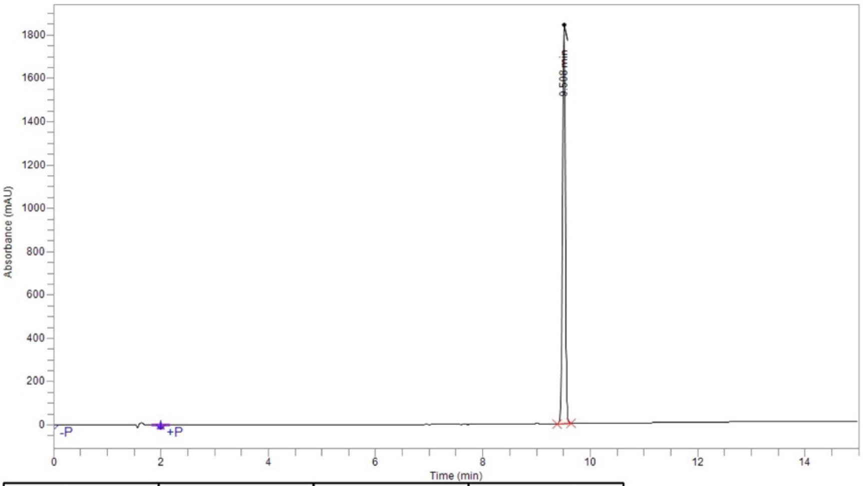

| Time | Height | Area | Area % |
|---|---|---|---|
| 9.508 | 1,844,947.1 | 6,668,456.4 | 100.00 |
| **Total** | | 6,668,456.4 | 100.00 |

5-(Tert-butyl)-2-(2,6-dioxopiperidin-3-yl)isoindoline-1,3-dione **(19)**

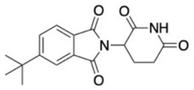

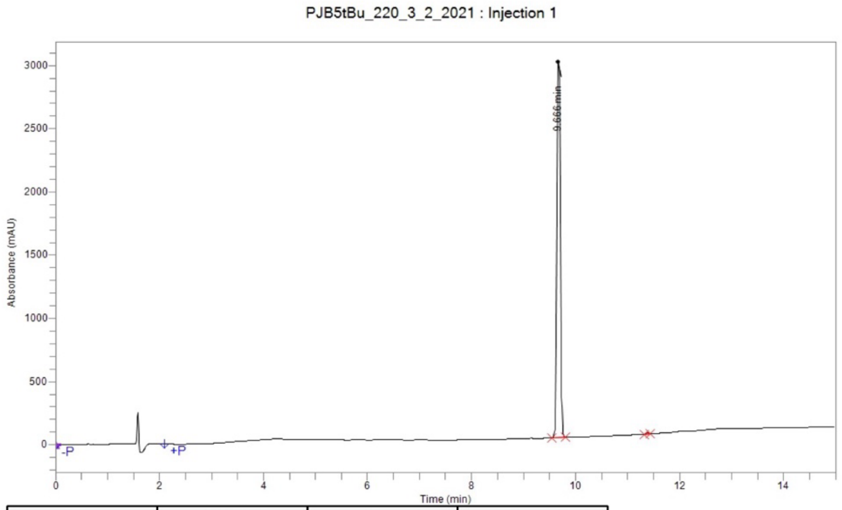

| Time | Height | Area | Area % |
|---|---|---|---|
| 9.666 | 3,015,607.0 | 13,255,833.1 | 99.79 |
| 11.388 | 8,856.5 | 28,068.3 | 0.21 |
| **Total** | | 13,283,901.5 | 100.00 |

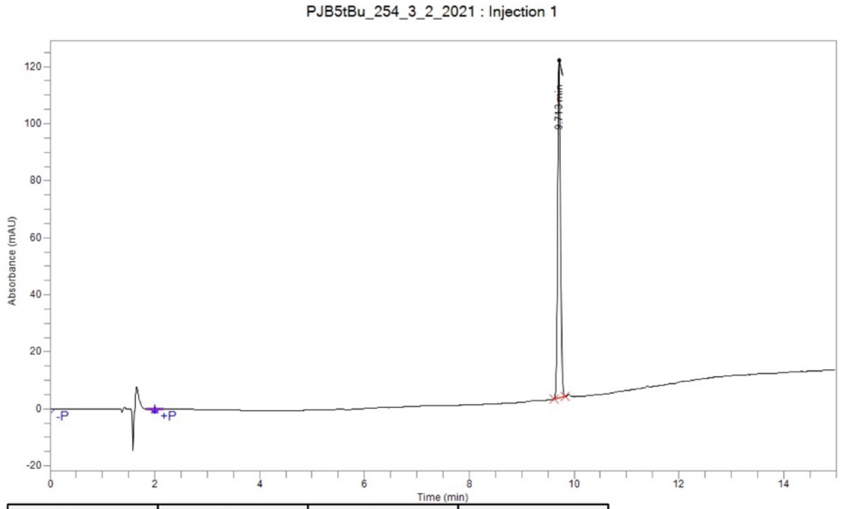

| Time | Height | Area | Area % |
|---|---|---|---|
| 9.713 | 118,463.5 | 453,471.6 | 100.00 |
| **Total** | | 453,471.6 | 100.00 |

2-(2,6-Dioxopiperidin-3-yl)-5-phenylisoindoline-1,3-dione **(20)**

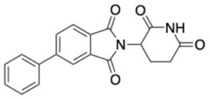

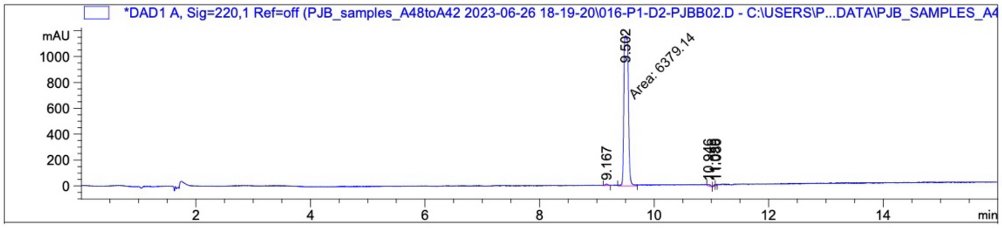

| Peak # | RetTime [min] | Type | Width [min] | Area [mAU*s] | Height [mAU] | Area % |
|---|---|---|---|---|---|---|
| 1 | 9.167 | VV | 0.0397 | 38.47590 | 12.08027 | 0.5924 |
| 2 | 9.502 | MM | 0.0916 | 6379.13721 | 1160.77014 | 98.2165 |
| 3 | 10.946 | VB | 0.0532 | 41.28960 | 10.78413 | 0.6357 |
| 4 | 11.059 | BV | 0.0189 | 16.44551 | 11.05400 | 0.2532 |
| 5 | 11.086 | VV | 0.0242 | 19.62358 | 10.30767 | 0.3021 |
| Totals : | | | | 6494.97180 | 1204.99620 | |

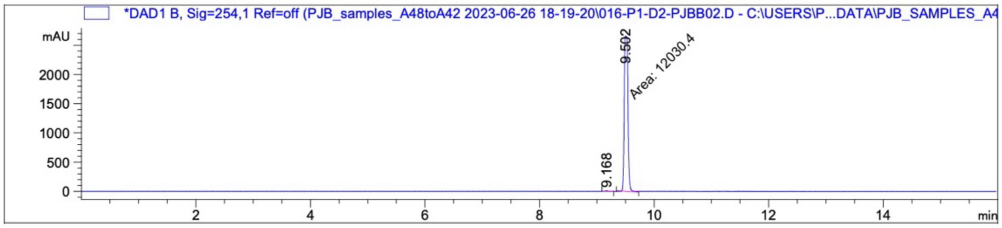

| Peak # | RetTime [min] | Type | Width [min] | Area [mAU*s] | Height [mAU] | Area % |
|---|---|---|---|---|---|---|
| 1 | 9.168 | BB | 0.0474 | 32.15612 | 10.45554 | 0.2666 |
| 2 | 9.502 | MM | 0.0756 | 1.20304e4 | 2653.79126 | 99.7334 |
| Totals : | | | | 1.20626e4 | 2664.24680 | |

5-(Dimethylamino)-2-(2,6-dioxopiperidin-3-yl)isoindoline-1,3-dione **(21)**

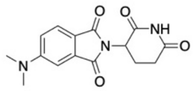

A62_220_13_7_2021 : Injection 1

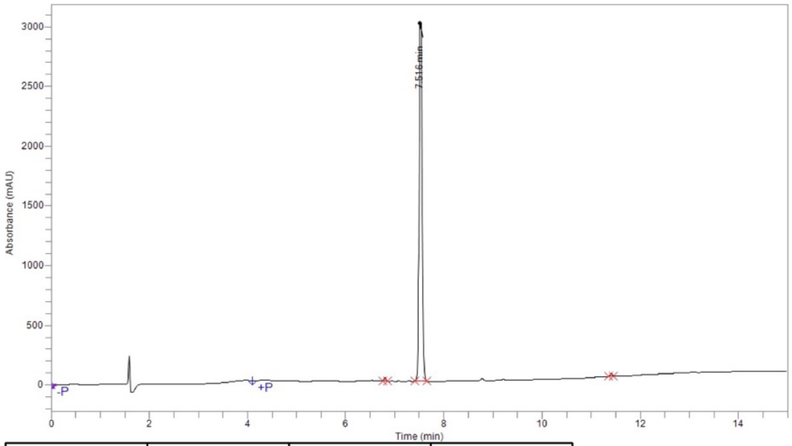

| Time | Height | Area | Area % |
|---|---|---|---|
| 6.801 | 31,486.5 | 93,221.0 | 0.68 |
| 7.516 | 3,044,345.1 | 13,613,467.1 | 98.89 |
| 11.409 | 20,678.6 | 59,497.1 | 0.43 |
| **Total** | | 13,766,185.2 | 100.00 |

A62_254_13_7_2021 : Injection 1

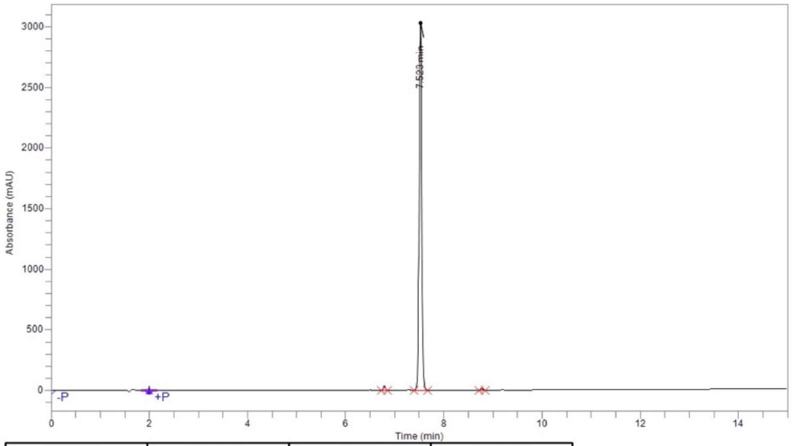

| Time | Height | Area | Area % |
|---|---|---|---|
| 6.790 | 37,642.5 | 119,915.4 | 1.03 |
| 7.523 | 3,057,655.3 | 11,472,276.8 | 98.60 |
| 8.777 | 11,796.8 | 42,782.4 | 0.37 |
| **Total** | | 11,634,974.6 | 100.00 |

2-(2,6-Dioxopiperidin-3-yl)-5-morpholinoisoindoline-1,3-dione **(22)**

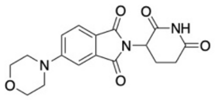

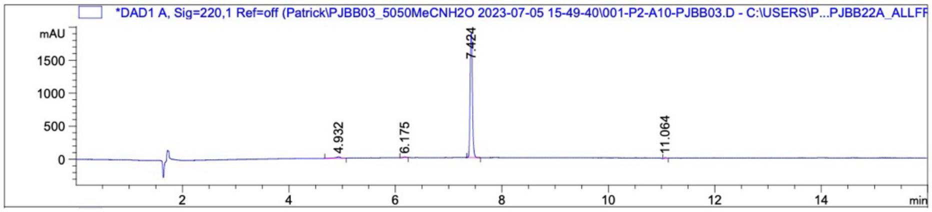

| Peak # | RetTime [min] | Type | Width [min] | Area [mAU*s] | Height [mAU] | Area % |
|---|---|---|---|---|---|---|
| 1 | 4.932 | BB | 0.0854 | 133.68457 | 19.52182 | 2.2990 |
| 2 | 6.175 | BB | 0.0472 | 36.72113 | 10.41101 | 0.6315 |
| 3 | 7.424 | BB | 0.0469 | 5612.18750 | 1878.12195 | 96.5140 |
| 4 | 11.064 | BB | 0.0438 | 32.30088 | 10.53894 | 0.5555 |
| Totals : | | | | 5814.89408 | 1918.59372 | |

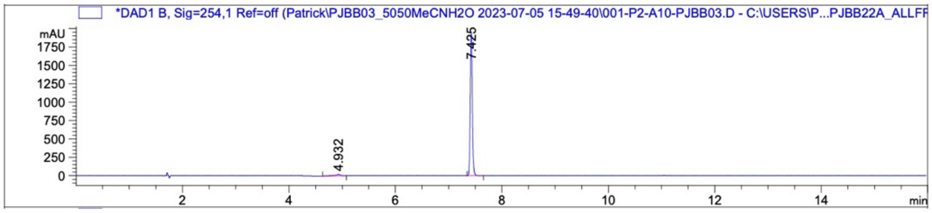

| Peak # | RetTime [min] | Type | Width [min] | Area [mAU*s] | Height [mAU] | Area % |
|---|---|---|---|---|---|---|
| 1 | 4.932 | BB | 0.0914 | 132.48828 | 19.23407 | 2.5003 |
| 2 | 7.425 | BB | 0.0413 | 5166.39648 | 1927.11658 | 97.4997 |
| Totals : | | | | 5298.88477 | 1946.35064 | |

2-(2,6-Dioxopiperidin-3-yl)-5-(4-methylpiperazin-1-yl)isoindoline-1,3-dione **(23)**

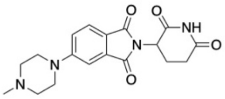

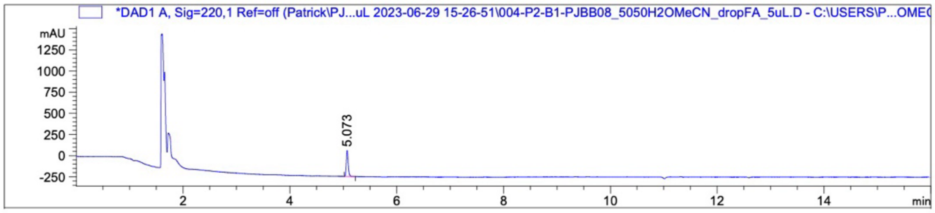

| Peak # | RetTime [min] | Type | Width [min] | Area [mAU*s] | Height [mAU] | Area % |
|---|---|---|---|---|---|---|
| 1 | 5.073 | BB | 0.0382 | 760.88721 | 309.12369 | 100.0000 |
| Totals : | | | | 760.88721 | 309.12369 | |

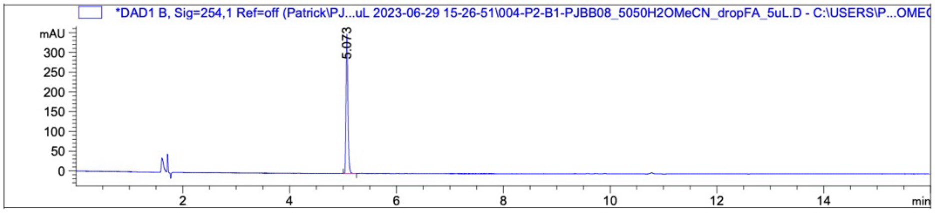

| Peak # | RetTime [min] | Type | Width [min] | Area [mAU*s] | Height [mAU] | Area % |
|---|---|---|---|---|---|---|
| 1 | 5.073 | BB | 0.0380 | 864.64014 | 353.40967 | 100.0000 |
| Totals : | | | | 864.64014 | 353.40967 | |

2-(2,6-Dioxopiperidin-3-yl)-4-(ethylamino)isoindoline-1,3-dione **(24)**

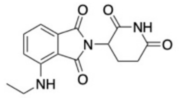

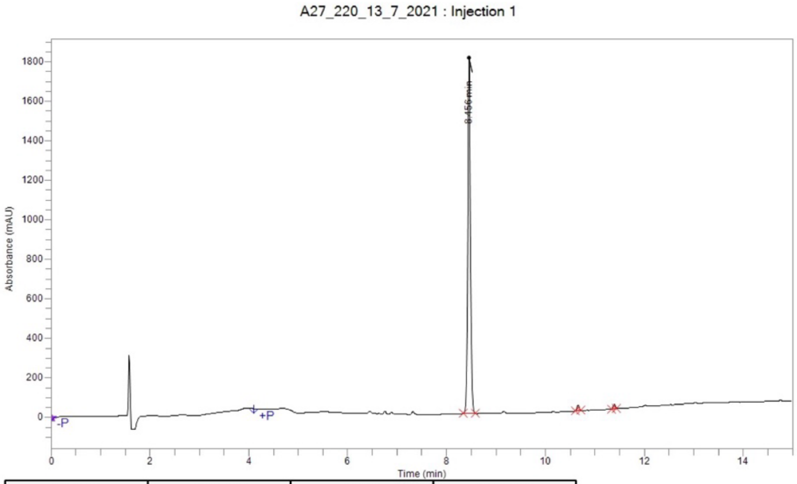

| Time | Height | Area | Area % |
|---|---|---|---|
| 8.456 | 1,800,428.2 | 7,219,094.3 | 97.92 |
| 10.660 | 27,073.5 | 88,235.7 | 1.20 |
| 11.390 | 22,352.5 | 65,484.8 | 0.89 |
| **Total** | | 7,372,814.8 | 100.00 |

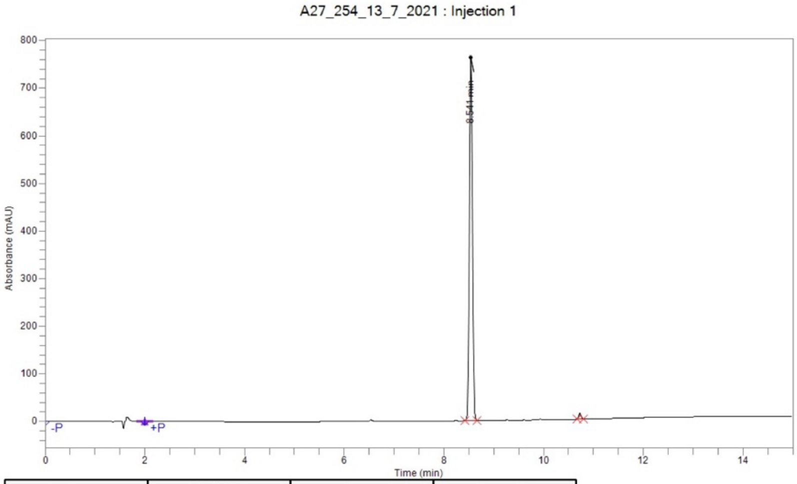

| Time | Height | Area | Area % |
|---|---|---|---|
| 8.541 | 763,584.8 | 3,012,532.3 | 98.61 |
| 10.731 | 12,178.1 | 42,392.6 | 1.39 |
| **Total** | | 3,054,924.9 | 100.00 |

2-(2,6-Dioxopiperidin-3-yl)-4-(prop-2-yn-1-ylamino)isoindoline-1,3-dione **(25)**

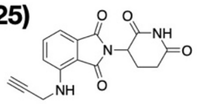

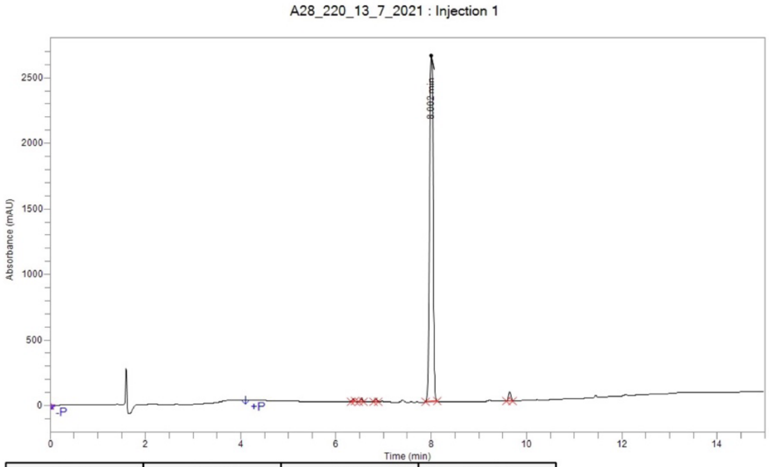

| Time | Height | Area | Area % |
|---|---|---|---|
| 6.357 | 21,251.7 | 61,668.2 | 0.45 |
| 6.531 | 27,158.9 | 83,361.4 | 0.61 |
| 6.838 | 28,107.5 | 86,355.7 | 0.63 |
| 8.002 | 2,641,521.8 | 13,311,843.4 | 96.69 |
| 9.649 | 69,316.8 | 224,418.0 | 1.63 |
| Total | | 13,767,646.8 | 100.00 |

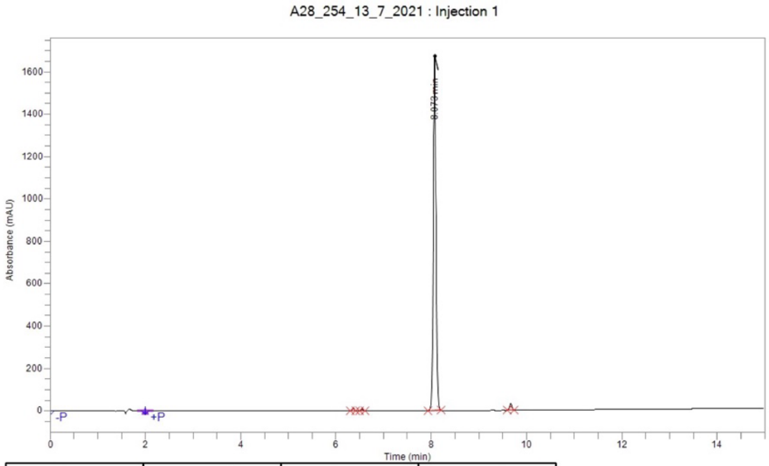

| Time | Height | Area | Area % |
|---|---|---|---|
| 6.365 | 10,841.5 | 35,164.2 | 0.49 |
| 6.547 | 8,636.8 | 27,331.6 | 0.38 |
| 8.073 | 1,674,556.2 | 7,007,741.9 | 97.68 |
| 9.670 | 31,892.5 | 103,799.2 | 1.45 |
| Total | | 7,174,036.9 | 100.00 |

4-(Benzylamino)-2-(2,6-dioxopiperidin-3-yl)isoindoline-1,3-dione **(26)**

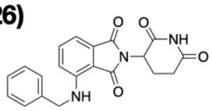

PJBA22_220 : Injection 1

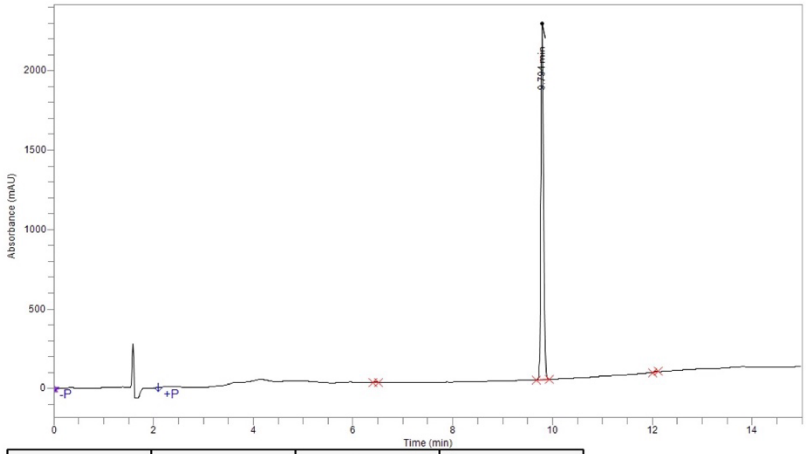

| Time | Height | Area | Area % |
|---|---|---|---|
| 6.453 | 5,600.9 | 19,763.4 | 0.22 |
| 9.794 | 2,243,243.4 | 8,880,223.9 | 99.53 |
| 12.077 | 5,477.1 | 22,020.7 | 0.25 |
| **Total** | | 8,922,008.0 | 100.00 |

PJBA22_254 : Injection 1

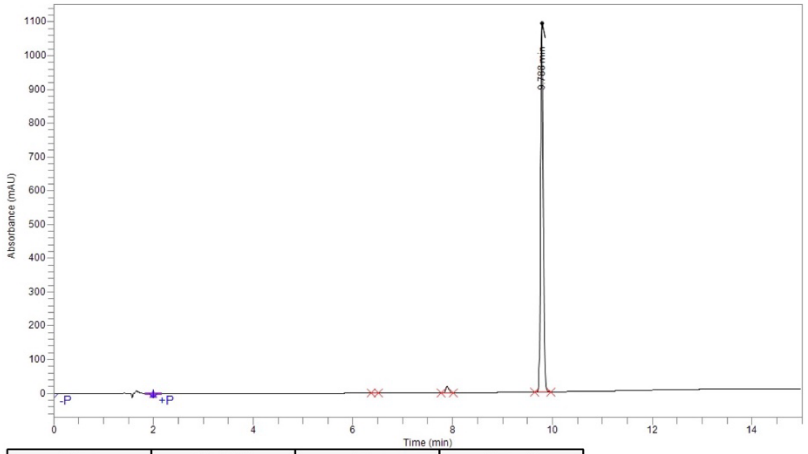

| Time | Height | Area | Area % |
|---|---|---|---|
| 6.444 | 1,859.3 | 7,084.6 | 0.16 |
| 7.894 | 19,164.5 | 93,740.9 | 2.17 |
| 9.788 | 1,093,396.3 | 4,221,720.4 | 97.67 |
| **Total** | | 4,322,546.0 | 100.00 |

2-(2,6-Dioxopiperidin-3-yl)-4-((pyridin-3-ylmethyl)amino)isoindoline-1,3-dione **(27)**

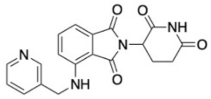

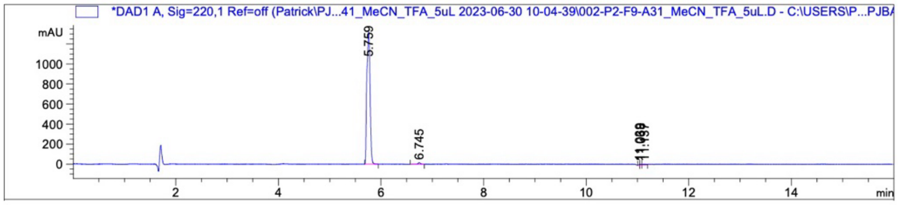

| Peak # | RetTime [min] | Type | Width [min] | Area [mAU*s] | Height [mAU] | Area % |
|---|---|---|---|---|---|---|
| 1 | 5.759 | BB | 0.0630 | 5757.53711 | 1316.86975 | 97.6576 |
| 2 | 6.745 | VV R | 0.0474 | 57.94011 | 14.87543 | 0.9828 |
| 3 | 11.039 | BV | 0.0169 | 10.67051 | 8.06211 | 0.1810 |
| 4 | 11.068 | VV | 0.0292 | 21.24766 | 9.12582 | 0.3604 |
| 5 | 11.137 | VV | 0.0710 | 48.23938 | 8.21837 | 0.8182 |
| Totals : | | | | 5895.63477 | 1357.15148 | |

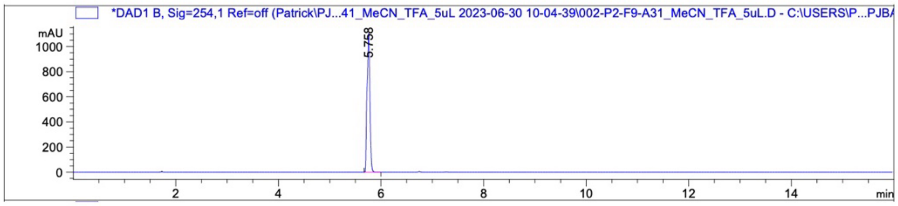

| Peak # | RetTime [min] | Type | Width [min] | Area [mAU*s] | Height [mAU] | Area % |
|---|---|---|---|---|---|---|
| 1 | 5.758 | BB | 0.0542 | 4278.00879 | 1104.08264 | 100.0000 |
| Totals : | | | | 4278.00879 | 1104.08264 | |

2-(2,6-Dioxopiperidin-3-yl)-4-((2-methylbenzyl)amino)isoindoline-1,3-dione **(28)**

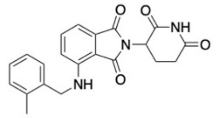

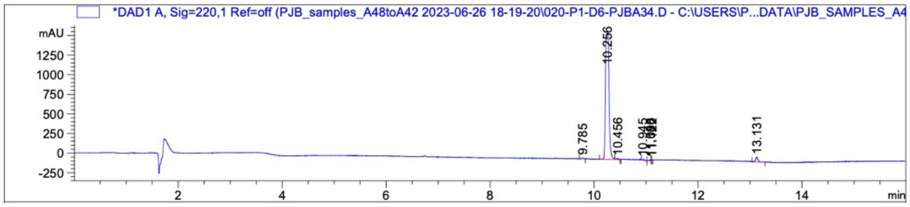

| Peak # | RetTime [min] | Type | Width [min] | Area [mAU*s] | Height [mAU] | Area % |
|---|---|---|---|---|---|---|
| 1 | 9.785 | VV | 0.0444 | 45.78736 | 12.75951 | 0.5763 |
| 2 | 10.256 | BV R | 0.0579 | 7577.73291 | 1609.09106 | 95.3733 |
| 3 | 10.456 | VV E | 0.0356 | 30.46284 | 10.73134 | 0.3834 |
| 4 | 10.945 | VB | 0.0530 | 46.46904 | 10.63425 | 0.5849 |
| 5 | 11.086 | BV | 0.0376 | 31.33801 | 10.12394 | 0.3944 |
| 6 | 11.096 | VV | 0.0175 | 9.68551 | 9.24094 | 0.1219 |
| 7 | 11.122 | VV | 0.0131 | 6.09629 | 7.05964 | 0.0767 |
| 8 | 13.131 | BV R | 0.0402 | 197.76942 | 61.97969 | 2.4891 |
| Totals : | | | | 7945.34139 | 1731.62037 | |

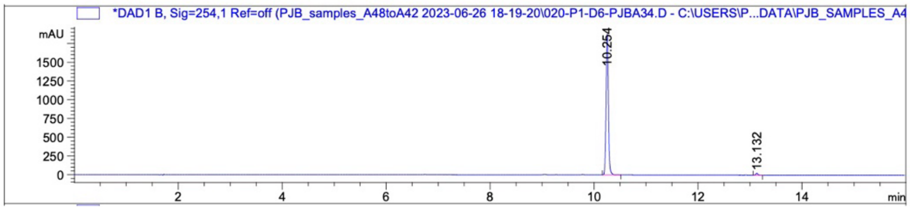

| Peak # | RetTime [min] | Type | Width [min] | Area [mAU*s] | Height [mAU] | Area % |
|---|---|---|---|---|---|---|
| 1 | 10.254 | BV R | 0.0481 | 5814.16699 | 1877.96265 | 98.7680 |
| 2 | 13.132 | BB | 0.0450 | 72.52560 | 24.90003 | 1.2320 |
| Totals : | | | | 5886.69259 | 1902.86268 | |

2-(2,6-Dioxopiperidin-3-yl)-4-(((4-methylpyridin-3-yl)methyl)amino)isoindoline-1,3-dione **(29)**

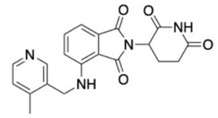

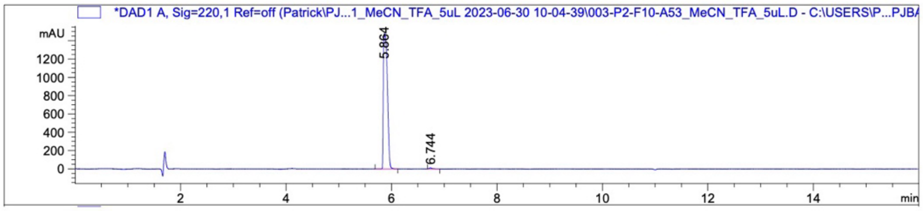

| Peak # | RetTime [min] | Type | Width [min] | Area [mAU*s] | Height [mAU] | Area % |
|---|---|---|---|---|---|---|
| 1 | 5.864 | VB R | 0.0630 | 7864.15283 | 1479.57996 | 99.4521 |
| 2 | 6.744 | BB | 0.0488 | 43.32656 | 10.80010 | 0.5479 |

Totals : 7907.47939 1490.38006

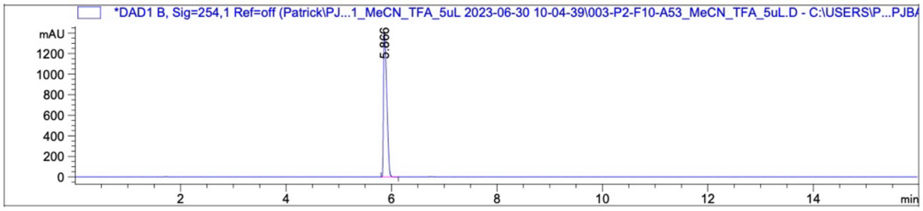

| Peak # | RetTime [min] | Type | Width [min] | Area [mAU*s] | Height [mAU] | Area % |
|---|---|---|---|---|---|---|
| 1 | 5.866 | BB | 0.0655 | 6021.60986 | 1392.17969 | 100.0000 |

Totals : 6021.60986 1392.17969

4-(Benzyloxy)-2-(2,6-dioxopiperidin-3-yl)isoindoline-1,3-dione **(30)**

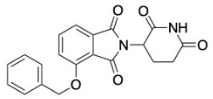

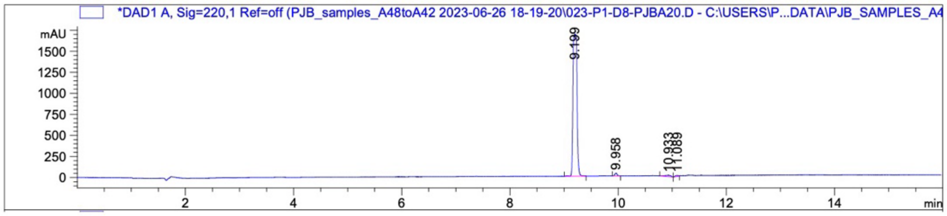

| Peak # | RetTime [min] | Type | Width [min] | Area [mAU*s] | Height [mAU] | Area % |
|---|---|---|---|---|---|---|
| 1 | 9.199 | VV R | 0.0616 | 8759.21680 | 1687.26782 | 96.7374 |
| 2 | 9.958 | BB | 0.0428 | 104.18281 | 33.90955 | 1.1506 |
| 3 | 10.933 | BB | 0.0947 | 129.00211 | 16.15368 | 1.4247 |
| 4 | 11.089 | BV | 0.0602 | 62.22880 | 12.37132 | 0.6873 |

Totals : 9054.63051 1749.70237

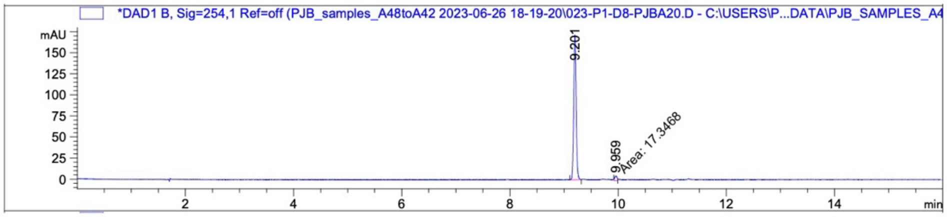

| Peak # | RetTime [min] | Type | Width [min] | Area [mAU*s] | Height [mAU] | Area % |
|---|---|---|---|---|---|---|
| 1 | 9.201 | BB | 0.0474 | 529.15289 | 171.89610 | 96.8258 |
| 2 | 9.959 | MM | 0.0531 | 17.34685 | 5.44532 | 3.1742 |

Totals : 546.49974 177.34143

2-(2,6-Dioxopiperidin-3-yl)-4-(pyridin-3-ylmethoxy)isoindoline-1,3-dione **(31)**

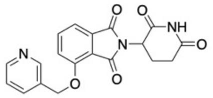

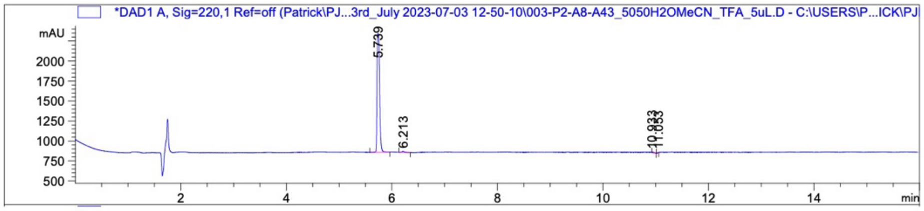

| Peak # | RetTime [min] | Type | Width [min] | Area [mAU*s] | Height [mAU] | Area % |
|---|---|---|---|---|---|---|
| 1 | 5.739 | BB | 0.0422 | 5070.66211 | 1490.58447 | 98.3312 |
| 2 | 6.213 | VV R | 0.0412 | 47.04042 | 14.54279 | 0.9122 |
| 3 | 10.933 | VB | 0.0443 | 27.20971 | 7.49671 | 0.5277 |
| 4 | 11.053 | BV | 0.0186 | 11.80451 | 8.07088 | 0.2289 |
| Totals : | | | | 5156.71675 | 1520.69486 | |

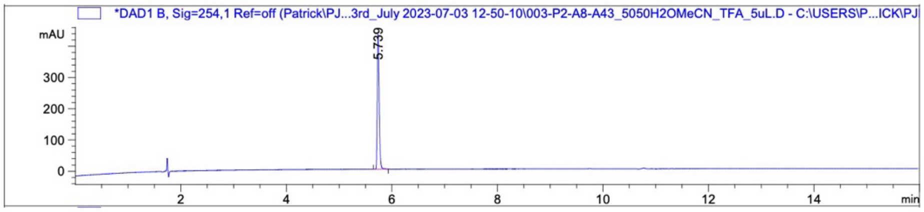

| Peak # | RetTime [min] | Type | Width [min] | Area [mAU*s] | Height [mAU] | Area % |
|---|---|---|---|---|---|---|
| 1 | 5.739 | BV R | 0.0364 | 1051.10193 | 431.90494 | 100.0000 |
| Totals : | | | | 1051.10193 | 431.90494 | |

2-(2,6-Dioxopiperidin-3-yl)-4-((2-methylbenzyl)oxy)isoindoline-1,3-dione **(32)**

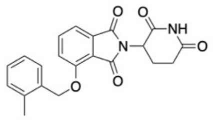

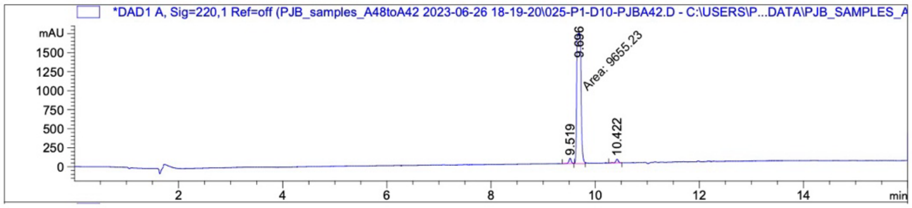

| Peak # | RetTime [min] | Type | Width [min] | Area [mAU*s] | Height [mAU] | Area % |
|---|---|---|---|---|---|---|
| 1 | 9.519 | BV | 0.0462 | 220.06575 | 71.82719 | 2.1918 |
| 2 | 9.696 | MM | 0.0935 | 9655.23340 | 1721.78210 | 96.1649 |
| 3 | 10.422 | VB R | 0.0463 | 164.99365 | 49.65711 | 1.6433 |
| Totals : | | | | 1.00403e4 | 1843.26640 | |

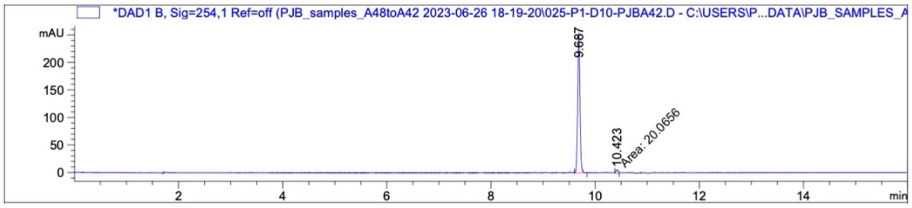

| Peak # | RetTime [min] | Type | Width [min] | Area [mAU*s] | Height [mAU] | Area % |
|---|---|---|---|---|---|---|
| 1 | 9.687 | BB | 0.0462 | 766.29730 | 253.89758 | 97.4483 |
| 2 | 10.423 | MM | 0.0522 | 20.06562 | 6.41060 | 2.5517 |
| Totals : | | | | 786.36293 | 260.30818 | |

4-(Benzhydrylamino)-2-(2,6-dioxopiperidin-3-yl)isoindoline-1,3-dione **(33)**

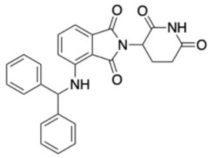

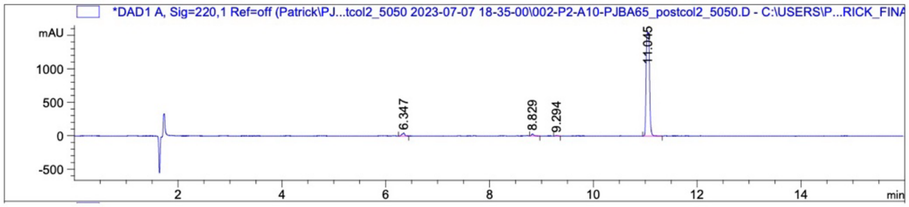

| Peak # | RetTime [min] | Type | Width [min] | Area [mAU*s] | Height [mAU] | Area % |
|---|---|---|---|---|---|---|
| 1 | 6.347 | BB | 0.0499 | 170.95700 | 46.51646 | 2.3214 |
| 2 | 8.829 | BB | 0.0469 | 96.15903 | 30.37664 | 1.3057 |
| 3 | 9.294 | BB | 0.0451 | 28.13963 | 9.34437 | 0.3821 |
| 4 | 11.045 | BB | 0.0635 | 7069.25488 | 1541.79138 | 95.9908 |
| Totals : | | | | 7364.51055 | 1628.02885 | |

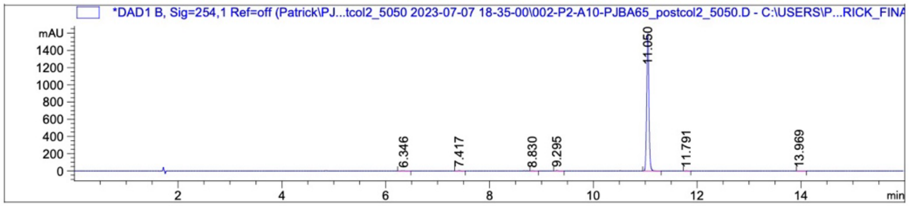

| Peak # | RetTime [min] | Type | Width [min] | Area [mAU*s] | Height [mAU] | Area % |
|---|---|---|---|---|---|---|
| 1 | 6.346 | BB | 0.0537 | 8.49456 | 2.12141 | 0.1766 |
| 2 | 7.417 | BB | 0.0469 | 9.34295 | 3.12326 | 0.1943 |
| 3 | 8.830 | BB | 0.0450 | 3.33332 | 1.05111 | 0.0693 |
| 4 | 9.295 | BB | 0.0453 | 11.41339 | 3.77203 | 0.2373 |
| 5 | 11.050 | BB | 0.0458 | 4765.96533 | 1597.53052 | 99.1084 |
| 6 | 11.791 | BB | 0.0543 | 3.98341 | 9.63068e-1 | 0.0828 |
| 7 | 13.969 | BB | 0.0718 | 6.30632 | 1.08459 | 0.1311 |
| Totals : | | | | 4808.83929 | 1609.64599 | |

4-(Benzhydryloxy)-2-(2,6-dioxopiperidin-3-yl)isoindoline-1,3-dione **(34)**

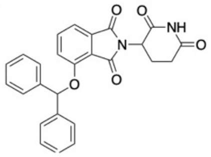

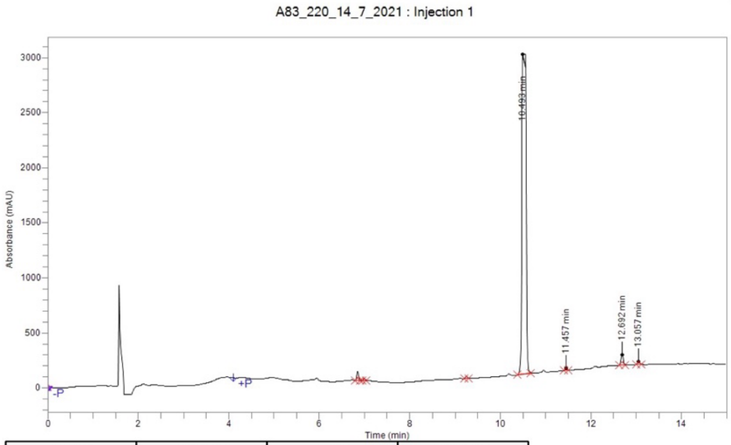

| Time | Height | Area | Area % |
|---|---|---|---|
| 6.850 | 84,966.7 | 237,594.8 | 1.19 |
| 6.994 | 19,324.9 | 50,593.3 | 0.25 |
| 9.255 | 18,338.3 | 54,587.1 | 0.27 |
| 10.493 | 2,942,967.8 | 19,196,493.1 | 95.94 |
| 11.457 | 20,516.2 | 58,680.1 | 0.29 |
| 12.692 | 95,320.7 | 316,917.6 | 1.58 |
| 13.057 | 27,577.0 | 93,787.6 | 0.47 |
| **Total** | | 20,008,653.7 | 100.00 |

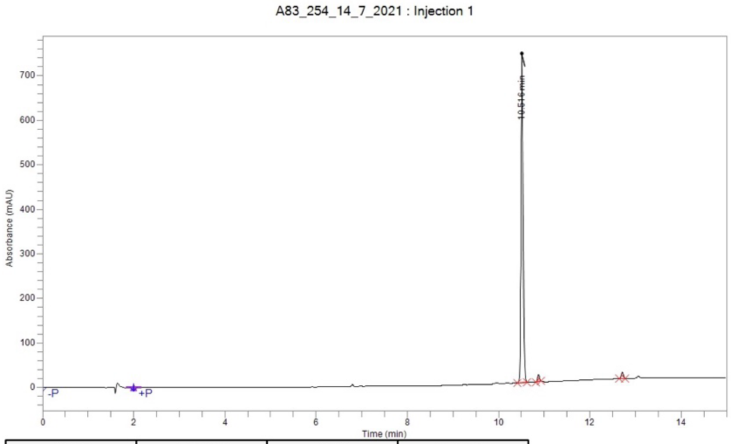

| Time | Height | Area | Area % |
|---|---|---|---|
| 10.516 | 739,977.7 | 2,443,080.6 | 96.27 |
| 10.876 | 15,065.8 | 48,078.3 | 1.89 |
| 12.714 | 14,528.5 | 46,657.1 | 1.84 |
| **Total** | | 2,537,816.0 | 100.00 |

6-(2,6-Dioxopiperidin-3-yl)-5H-pyrrolo[3,4-b]pyridine-5,7(6H)-dione **(35)**

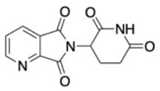

A66_220_13_7_2021 : Injection 1

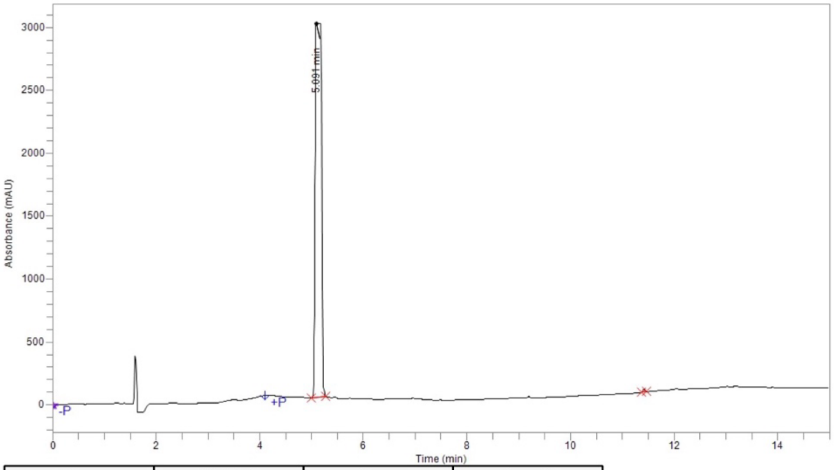

| Time | Height | Area | Area % |
|---|---|---|---|
| 5.091 | 3,000,256.3 | 24,431,285.3 | 99.74 |
| 11.418 | 22,738.6 | 64,527.0 | 0.26 |
| **Total** | | 24,495,812.2 | 100.00 |

A66_254_13_7_2021 : Injection 1

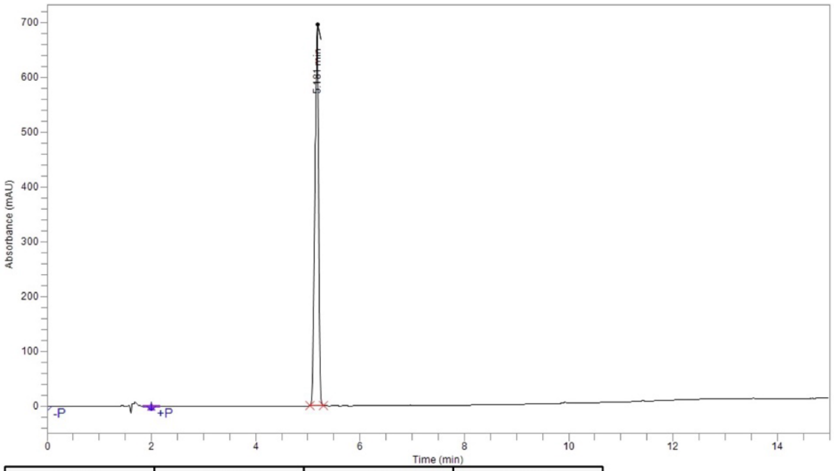

| Time | Height | Area | Area % |
|---|---|---|---|
| 5.181 | 697,023.0 | 3,671,621.6 | 100.00 |
| **Total** | | 3,671,621.6 | 100.00 |

2-(2,6-Dioxopiperidin-3-yl)-1H-pyrrolo[3,4-c]pyridine-1,3(2H)-dione **(36)**

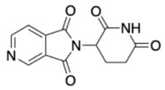

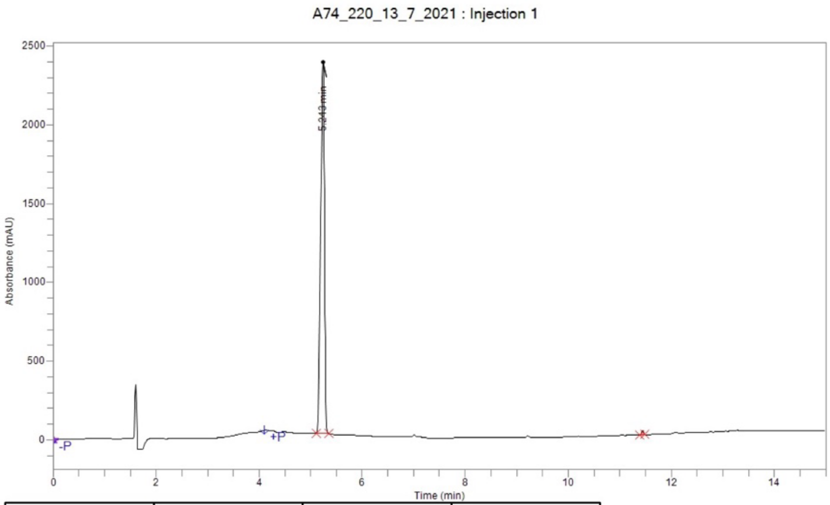

| Time | Height | Area | Area % |
|---|---|---|---|
| 5.243 | 2,361,847.5 | 12,178,523.5 | 99.41 |
| 11.440 | 23,934.0 | 72,061.0 | 0.59 |
| **Total** | | 12,250,584.5 | 100.00 |

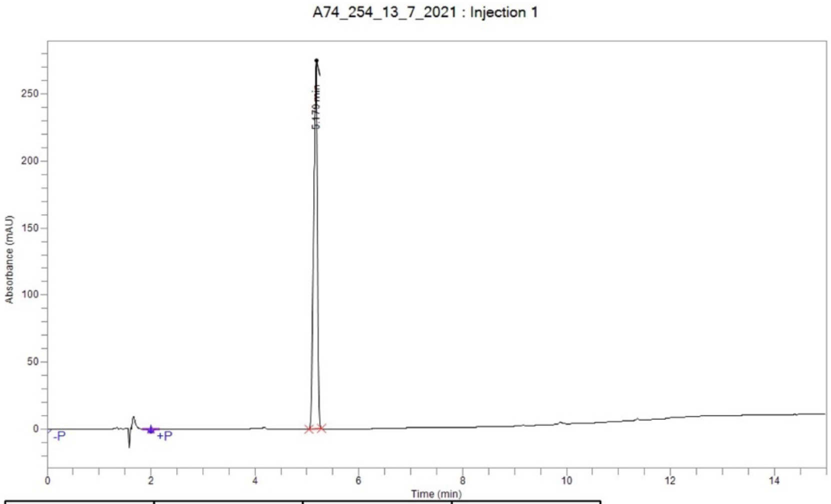

| Time | Height | Area | Area % |
|---|---|---|---|
| 5.179 | 275,514.1 | 1,450,941.8 | 100.00 |
| **Total** | | 1,450,941.8 | 100.00 |
